# A brain-penetrant P2X7R antagonist mitigates Alzheimer’s disease pathology

**DOI:** 10.64898/2026.04.06.716621

**Authors:** Andreea L. Turcu, Adam C. Oken, Christian Griñán-Ferré, Anna Durner, Juan Sierra-Marquez, Sonja Hinz-Kowalik, Jessica Nagel, So-Deok Lee, Efpraxia Tzortzini, Kyriakos Georgiou, Mamina Bhol, Zuriñe Baz, Jordi Llop, Marion Schneider, Carla Barbaraci, Ga-Ram Kim, Marta Barniol-Xicota, Cristina Val, José Brea, M. Isabel Loza, Belén Pérez, Lieve Naesens, Antonios Kolocouris, Yong-Chul Kim, Christa E. Müller, Annette Nicke, Mercè Pallàs, Steven E. Mansoor, Santiago Vázquez

## Abstract

The ATP-gated P2X7 receptor (P2X7R) activates inflammatory signaling pathways in the central nervous system. In particular, P2X7Rs drive chronic glia-mediated neuroinflammation, which is increasingly recognized as a key contributor to Alzheimer’s disease, a neurodegenerative disorder that lacks effective disease-modifying therapies. Here we identify a potent and selective negative allosteric modulator of P2X7Rs with therapeutic potential. We synthesize a series of small molecules based on a polycyclic scaffold and confirm blood-brain barrier penetration by testing a radiolabeled analogue using positron emission tomography imaging. Through a structure-guided medicinal chemistry campaign centered on our scaffold, we identify four promising P2X7R antagonists. Of these, UB-ALT-P2 exhibits the most favorable safety profile, high oral bioavailability and robust brain penetration. High-resolution cryo-EM structures of UB-ALT-P2 bound to human, mouse, and rat P2X7Rs reveal a conserved antagonist binding mode with steric features that favor prolonged binding to human receptors. In the 5xFAD mouse model of AD, oral UB-ALT-P2 blunts weight loss, improves short- and long-term memory, reduces amyloid-β plaque burden, lowers hyperphosphorylated tau, and diminishes oxidative and inflammatory markers. These results establish UB-ALT-P2 as a potent and safe P2X7R antagonist that can mitigate core AD pathologies, providing a compelling foundation for further development.

## Introduction

Alzheimer’s disease (AD) is a progressive neurodegenerative disorder of the central nervous system (CNS) and represents the leading cause of dementia worldwide. The disease currently affects more than 50 million individuals, a number that is projected to rise as the population ages^1^. Pathologically, brains affected by AD are characterized by the accumulation of extracellular Aβ plaques and intracellular neurofibrillary tangles of hyperphosphorylated tau, particularly in the hippocampus and entorhinal cortex^2–3^. These cellular changes disrupt synaptic signaling, promote neurotoxic cascades, and result in a progressive breakdown of memory systems, including encoding and consolidation of episodic information^4^. Despite decades of research to understand and treat AD, disease-modifying therapies remain limited, underscoring an urgent need to identify new and effective therapeutic strategies.

Chronic neuroinflammation is increasingly recognized as a major driver of AD pathology. Activation of microglia, sustained release of pro-inflammatory mediators, and prolonged inflammatory responses have been shown to contribute to synaptic dysfunction, neuronal loss, and cognitive decline^5–6^. A major trigger of neuroinflammation is extracellular ATP, a danger-associated molecular pattern molecule. ATP activates purinergic receptors, among which, P2X7Rs on microglia and astrocytes are emerging as key regulators of inflammation^7–9^. Their sustained activation drives pro-inflammatory pathways that link immune activation to impaired Aβ clearance, tau pathology, and synaptic dysfunction^10–14^. Indeed, upregulation of P2X7R expression is consistently observed in postmortem AD brains^15–16^. Transgenic modification, genetic deletion, or inhibition of P2X7R activity reduces AD pathology^17–22^. However, progress towards clinical translation has been hampered by the scarcity of selective P2X7R antagonists with a high degree of brain penetration^23^.

In this study, we address this translational hurdle by uncovering a highly selective P2X7R antagonist with therapeutic promise for AD. Utilizing a systematic medicinal chemistry campaign centered around a brain-penetrant scaffold, we advanced four lead compounds through structural, pharmacokinetic, and metabolic assessments. High-resolution cryo-EM structures of our lead compound, UB-ALT-P2, revealed species-specific interaction networks underlying its diverse pharmacological effects on rat, mouse and human P2X7Rs. The structures further provided a mechanistic explanation for the slow dissociation and prolonged residence time of UB-ALT-P2, particularly on the human receptor. Notably, oral administration of UB-ALT-P2 to 5xFAD mice blunted weight loss, reduced amyloid burden, improved tau pathology, and preserved short- and long-term memory. Together, these findings define a potent, selective, and well-tolerated scaffold for the development of novel AD therapeutics and establish P2X7R inhibition as a viable strategy for targeting neuroinflammation and neurodegeneration.

### Polycyclic P2X7R antagonists penetrate the CNS

Inspired by prior studies of adamantane-based inhibitors and our identification of a P2X7R antagonist with a tetracyclo[4.4.0.0^3,9^.0^4,8^]decane core (UB-MBX-46)^24–26^, we synthesized a series of small molecules based on this polycyclic scaffold (Extended Data Table 1). In these analogs, the substituted aryl moiety and its linker to the polycyclic core were systematically varied to define structure–activity relationships (SARs) for the human P2X7R (hP2X7R) and to optimize metabolic stability and pharmacokinetic properties (Fig. 1A, Extended Data Table 1). Synthetic procedures are described in the Supplementary Information, and compound characterization is provided in Supplementary Figs. 1–25 and Supplementary Table 1. Half-maximal inhibitory concentrations (IC₅₀ values) were determined using an ethidium bromide uptake assay in HEK293 cells expressing hP2X7R (Extended Data Table 1).

**Figure 1.**
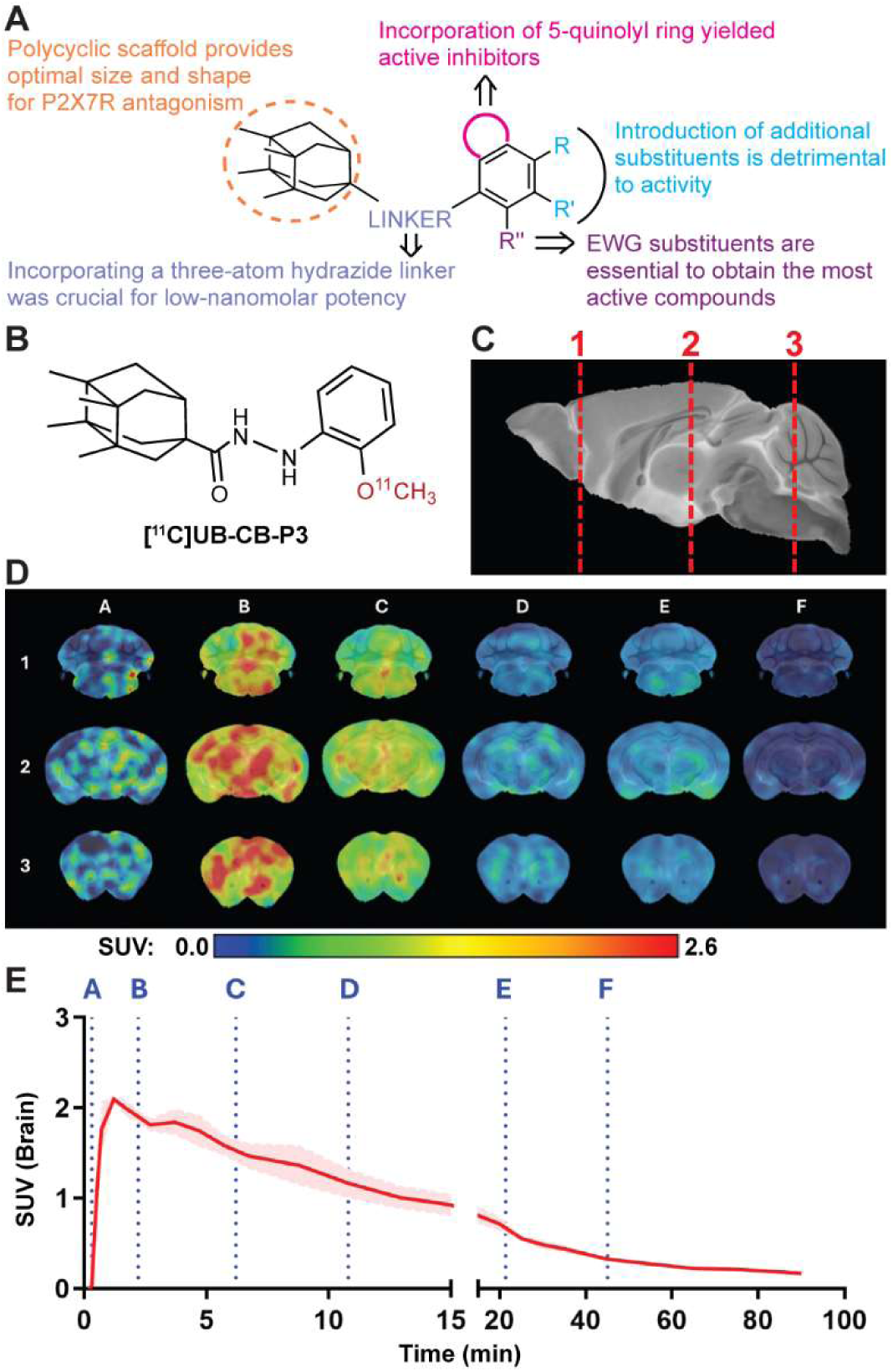
Compounds based on a unique polycyclic scaffold show rapid CNS uptake during PET imaging. (**A**) Overview of SARs for this compound series^24^. The tetramethyl core with a short hydrazide linker provides a favorable steric framework for receptor antagonism, especially with a carbonyl proximal to the polycyclic core. In the intermediate aryl ring, *o*-electron-withdrawing (EWG) substituents maximize potency whereas additional substituents are detrimental. Incorporation of a 5-quinolinyl ring yields the highest activity. (**B**) Chemical structure of the radiolabeled tracer [¹¹C]UB-CB-P3 for PET studies, with carbon-11 incorporation as indicated. (**C**) Representative sagittal section of a mouse brain. Regions of interest for quantitative PET analysis indicated by dashed vertical lines (1, 2, and 3). (**D**) Representative coronal PET images of a mouse brain following intravenous injection of [¹¹C]UB-CB-P3. Standardized uptake value (SUV) maps for six acquisition frames (A–F) show rapid and widespread brain uptake at early time points (A–C) and progressive washout at later times (D–F). Color scale (0.0 – 2.6 SUV) is shown below and time points (A–F) are as in **E**. (**E**) Time–activity curve (SUV *vs.* time) for [¹¹C]UB-CB-P3 brain uptake following intravenous injection. Rapid brain uptake, consistent with BBB penetration, is followed by gradual washout. Shading represents variability across measurements. Vertical dashed lines labeled A – F indicate time points corresponding to images in **D**.

In the first series of SAR studies, we repositioned the *ortho*-chlorine on the aryl ring of UB-MBX-46 to the *meta-* or *para-*positions (UB-ALT-P39 and UB-ALT-P40, respectively) (Fig. 1A, Extended Data Table 1). These modifications increased the IC_50_ for UB-MBX-46 from 1.0 ± 0.2 nM to more than 1 µM, abolishing activity. Similarly, the addition of a second chlorine to the aryl ring (UB-ALT-P41 and UB-ALT-P42) markedly reduced potency (Extended Data Table 1), confirming a strict requirement for a single *ortho*-substituent for optimal receptor engagement. The introduction of hydroxy (UB-CB-P4) or methoxy (UB-CB-P3) groups at the *ortho*-position also decreased activity (Extended Data Table 1), likely due to their polarity disrupting predominantly hydrophobic contacts between the aryl ring and side chains that stabilize the ligand in the allosteric pocket (F88, M105, and I310)^24^. Electron-withdrawing substituents exhibited size-dependent effects; fluorine (UB-CB-P1) abolished activity while bulkier pentafluorosulfanyl (UB-ALT-P6) or iodo (UB-ALT-P35) substitutions retained moderate potency (IC_50_ = 39 ± 4 nM and 50 ± 20 nM, respectively) (Extended Data Table 1). Indeed, a trifluoromethyl substituent (UB-MBX-47), a size-bioisostere of chlorine^27^, preserved activity (IC_50_ = 4.5 ± 0.7 nM, Extended Data Table 1) to levels near UB-MBX-46. These results suggest that, although the allosteric pocket disfavors excessive polarity or steric bulk, it tolerates substituents that enhance lipophilicity without altering the spatial alignment of the aryl ring. We next explored the heteroaromatic replacements 2,6-dichloro-4-pyridinyl (UB-ALT-P1) and 5-quinolinyl (UB-ALT-P2), which were approximately as potent as our initial compound (IC_50_ = 1.6 ± 0.8 nM and 1.3 ± 0.5 nM, respectively) (Extended Data Table 1), indicating that the allosteric pocket can accommodate electron-poor heteroaromatic systems. However, the addition of a 2-chloro substituent to the 5-quinolinyl ring (UB-ALT-P21) reduced potency (IC_50_ = 90 ± 40 nM), likely due to a deviation of the heteroaromatic system from an optimal orientation inside the allosteric pocket (Extended Data Table 1).

In the second series of SAR studies, we modified the linker between the polycyclic scaffold and the quinoline moiety of our most potent derivative, UB-ALT-P2 (Extended Data Table 1). Replacement of the hydrazide bridge with less flexible or electronically different linkers, including amides, ureas, and thioureas, typically abolished activity (Fig. 1A and Extended Data Table 1). Although two amide derivatives deviated from this trend by retaining some potency (UB-ALT-P19, IC_50_ = 42 ± 9 nM; UB-ALT-P20, IC_50_ = 31 ± 7 nM), these results highlight the limited tolerance for modifications of the bridge (Fig. 1A and Extended Data Table 1). This is likely due to the hydrogen bonds that form between the hydrazide linker and the backbone carbonyl of D92 as well as the side chain hydroxyl of Y298^24^. Together, these SAR studies define the steric and electronic constraints of the allosteric pocket in the hP2X7R and reveal the minimal structural features required for potent inhibition, underscoring the stringent spatial and electronic demands of this binding site.

Because ligand access to the CNS is a prerequisite for AD therapies^28^ and to support development of a companion diagnostic to assess *in vivo* target engagement, we evaluated the ability of our polycyclic P2X7R antagonist scaffold to cross the blood-brain barrier (BBB) using a radiolabeled analog of compound UB-CB-P3 (Fig. 1B and Extended Data Table 1). UB-CB-P3 was selected as a representative compound because its precursor, UB-CB-P4, contains an accessible methylation site that enables radiolabeling with [¹¹C]CH₃I, yielding a tracer ([¹¹C]UB-CB-P3) with an optimal balance of lipophilicity and hydrophilicity (logP = 2.61) for passive diffusion across the BBB (Supplementary Fig. 26)^29^. Dynamic positron emission tomography (PET) imaging in healthy mice confirmed that intravenous injection of [¹¹C]UB-CB-P3 resulted in rapid brain uptake (peak standardized uptake value ∼2.0), consistent with efficient BBB transit (Fig. 1C-E, Extended Data Fig. 1A-B)^30^. Although brain radioactivity declined gradually thereafter, indicating non-specific distribution and washout, these data show that ligands derived from our polycyclic scaffold can penetrate the CNS. During the pseudo-equilibrium window (15–60 min), the brain-to-blood partition coefficient (K_p,brain_) of [¹¹C]UB-CB-P3 was ∼0.45, indicating a lower concentration in the brain than the blood during this interval. Several centrally active, clinically effective drugs have similar K_p,brain_ values^31^, including meprobamate (∼0.42), midazolam (∼0.23) and morphine (∼0.46). Although others, such as risperidone (∼0.78) and caffeine (∼1.0), span slightly higher ranges^31^, this diversity highlights that K_p,brain_ values below 1 are compatible with meaningful CNS pharmacology, often reflecting differences in non-specific binding rather than limitations in BBB permeability. Elsewhere, transient uptake was observed in the heart, liver, and kidneys, consistent with perfusion and systemic clearance (Extended Data Fig. 1C-H). Progressive accumulation in the bladder established urinary excretion as the primary route of elimination (Extended Data Fig. 1F). Together, these PET and SAR studies confirmed the accessibility of our polycyclic scaffold to brain tissue and identified four compounds with low nanomolar potency (UB-MBX-46, UB-MBX-47, UB-ALT-P1, and UB-ALT-P2) for further evaluation (Fig. 2A-D, Extended Data Table 1).

**Figure 2:**
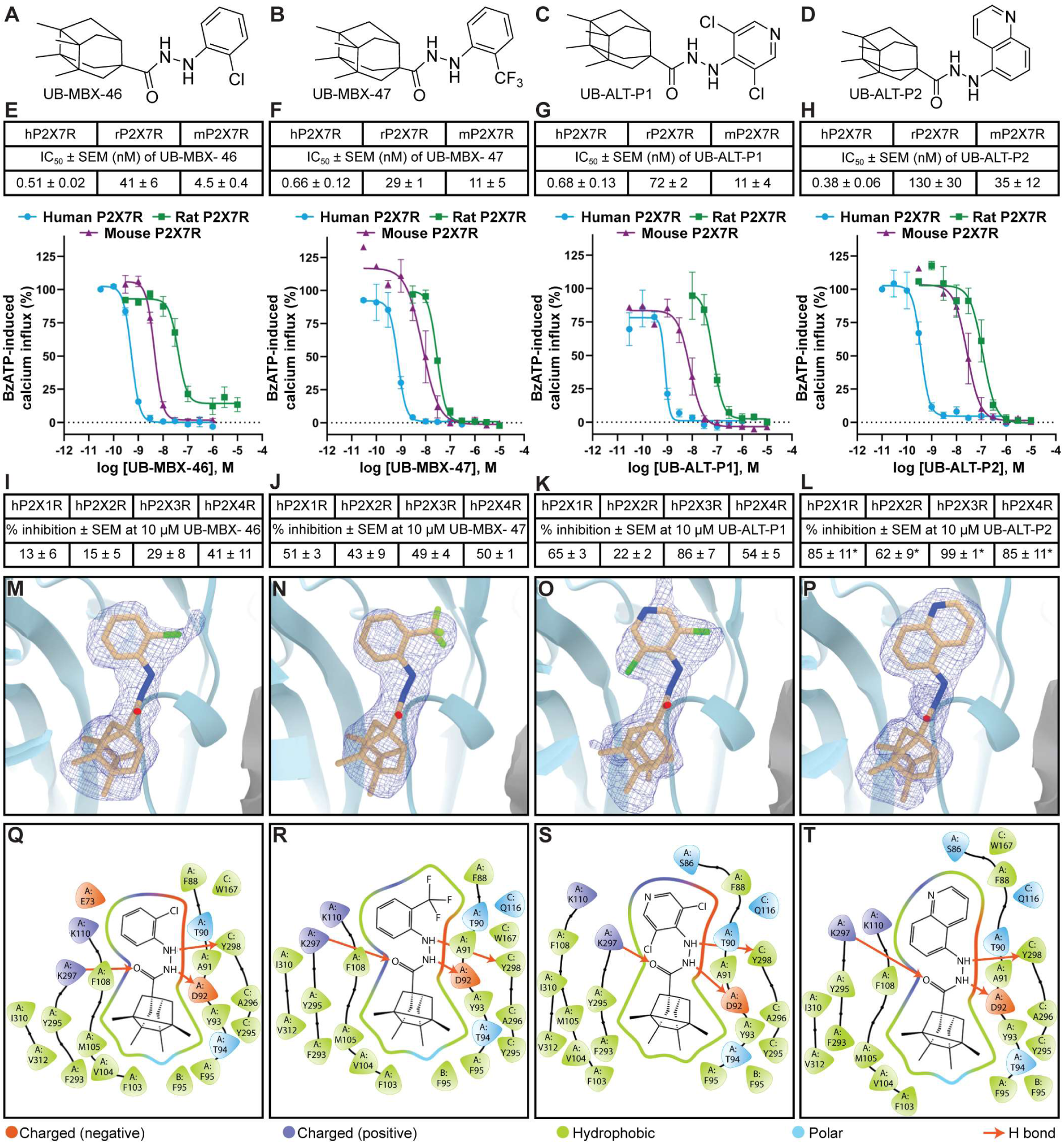
Tetracyclo[4.4.0.0^3,9^.0^4,8^]decane-based antagonist scaffolds effectively inhibit the hP2X7R. (**A-D**) 2D chemical structures of UB-MBX-46 (**A**), UB-MBX-47 (**B**), UB-ALT-P1 (**C**), and UB-ALT-P2 (**D**). (**E-H**) Concentration–response curves for inhibition of agonist-induced calcium influx through human (blue circles), rat (green squares), and mouse (purple triangles) P2X7Rs by UB-MBX-46 (**E**), UB-MBX-47 (**F**), UB-ALT-P1 (**G**), and UB-ALT-P2 (**H**). Recombinant P2X7Rs were expressed in 1321N1 astrocytoma cells, activated by an 80% maximal effective concentration (EC_80_) of BzATP (hP2X7R: 12.0 µM, mP2X7R: 17.3 µM, rP2X7R: 1.58 µM), and responses normalized to that from an EC_80_ concentration in the absence of antagonist. (**I-L**) Inhibition of calcium influx through hP2X1Rs, hP2X2Rs, hP2X3Rs, and hP2X4Rs by 10 µM UB-MBX-46 (**I**), UB-MBX-47 (**J**), UB-ALT-P1 (**K**), and UB-ALT-P2 (**L**). Receptors were stably expressed in 1321N1 astrocytoma cells and activated by an EC_80_ concentration of ATP (hP2X1R, 100 nM; hP2X2R, 1000 nM; hP2X3R, 100 nM; hP2X4R, 300 nM). *UB-ALT-P2 inhibited hP2X1Rs, hP2X2Rs, hP2X3Rs, and hP2X4Rs with IC_50_ values of 4.6 ± 0.6 µM, 5.2 ± 1.2 µM, 1.7 ± 0.5 µM, and 4.9 ± 0.9 µM, respectively. Data in **E-L** are mean ± SEM of at least three independent experiments performed in duplicate. (**M-P**) Ribbon representation of the classical allosteric pocket in hP2X7R at the interface of two protomers (chain A in light blue and chain C in grey) bound to one molecule of UB-MBX-46 (**M**), UB-MBX-47 (**N**), UB-ALT-P1 (**O**), and UB-ALT-P2 (**P**) shown in tan with corresponding cryo-EM density in blue mesh at 2.5 Å, 2.8 Å, 2.9 Å, and 2.6 Å, respectively. (**Q-T**) Schematic representation of ligand–receptor interactions between UB-MBX-46 (**Q**), UB-MBX-47 (**R**), UB-ALT-P1 (**S**), and UB-ALT-P2 (**T**) and the classical allosteric pocket of hP2X7R. Protomer chain IDs (A, B and C) are assigned in a counterclockwise orientation when viewed from the extracellular domain. Hydrogen bonds shown as red arrows and residue interactions in green (hydrophobic), blue (positively charged), red (negatively charged), and light blue (polar). Interaction plots generated using Maestro^36^. Data for UB-MBX-46 from ref. 24.

### Four compounds target the classical allosteric pocket

To ensure consistent exposure–response relationships and accurate dose predictions between preclinical animal models and humans^32^, we measured the potency of our four lead compounds on different P2XR subtypes and P2X7R orthologs (Fig. 2A-L). In calcium influx assays, all four compounds displayed sub-nanomolar potency at the hP2X7R (IC_50_ = 0.4 nM - 0.7 nM) (Fig. 2E-H). Although potency was lower at both the rP2X7R (IC_50_ = 29 nM - 130 nM) and mP2X7R (IC_50_ = 5 nM - 35 nM) (Fig. 2E-H)^24^, all four compounds acted in the low-nanomolar range on the mouse ortholog, demonstrating their suitability for *in vivo* efficacy studies in murine models of AD. We subsequently tested the specificity of the four compounds for other P2XR subtypes using a concentration at which 100% inhibition of the hP2X7R was expected (10 μM). All four compounds were much less potent at the hP2X1R (>10,000-fold reduction), hP2X2R (>10,000-fold reduction), hP2X3R (>4,000-fold reduction), and hP2X4R (>10,000-fold reduction) (Fig. 2I-L). Thus, despite the co-expression of multiple P2XR subtypes across many cell types, the lower activity at the other P2XR subtypes suggests that our compounds will selectively target hP2X7R *in vivo*^33^.

Before advancing the compounds for further screening, we sought to validate their mechanism of action by determining the interactions of each compound with hP2X7R. High-resolution cryo-EM structures of UB-MBX-47- (2.8 Å), UB-ALT-P1- (2.9 Å), and UB-ALT-P2-bound hP2X7Rs (2.6 Å) were compared to our previous structure of UB-MBX-46-bound hP2X7R (2.5 Å)^24^ (Fig. 2M-T, Extended Data Fig. 2-4, Supplementary Table 2). Each structure represents an antagonist-bound inhibited state of the receptor with sufficient resolution to describe the pose of the ligand within the classical allosteric pocket (Fig. 2M-P and Extended Data Fig. 4)^34^. Each compound has a similar pose, interacting with residues known to be important for ligand binding^34–35^. In particular, the interaction between the hydrazide linker and the backbone carbonyl of D92 is present in all four structures (Fig. 2Q-T and Extended Data Fig. 4)^24,34^. Molecular dynamics simulations at the hP2X7R show that UB-MBX-46^24^, UB-MBX-47, UB-ALT-P1, and UB-ALT-P2 remain stably bound within the allosteric pocket, exhibiting a similar overall interaction pattern characterized by hydrogen bonds, aromatic interactions (π–π and cation–π), hydrophobic contacts, and water-mediated interactions throughout 500 ns simulations (Extended Data Fig. 5A-C). Given the high potency and selectivity for the hP2X7R, robust cross-species activity, and similar binding poses, all four compounds were advanced to *in vitro* drug metabolism and pharmacokinetic (DMPK) profiling.

### UB-ALT-P2 has a favorable drug-like profile

To investigate the safety and potential utility of our four lead compounds, we conducted further *in vitro* studies. First, cytotoxicity testing across four mammalian cell lines representing diverse tissue origins and biological contexts (HEL, HeLa, Vero, and MT4) revealed no detectable adverse effects for any of the compounds (half-maximal cytotoxic concentration (CC_50_) > 50 µM in MT4 cells and > 100 µM in HEL, HeLa, and Vero cells) (Supplementary Table 3). In addition, inhibition of the hERG potassium channel—an early predictor of cardiotoxicity for ion-channel modulators^37^—was assessed at a concentration at which 100% inhibition of the hP2X7R is expected (10 μM), and none of the compounds produced significant hERG inhibition (Table 1).

**Table 1.** DMPK and safety profiling of the *in vivo* candidates UB-MBX-46, UB-MBX-47, UB-ALT-P1, and UB-ALT-P2.

| Code | hERG channel<br>% inh. ±<br>SD at<br>10 μM<sup>a</sup> | PAMPA<br>BBB<sup>b</sup> | Caco-2<br>permeability<sup>c</sup> | | Cytochrome (CYP) inhibition<br>% inh. ± SD at 10 μM or IC<sub>50</sub><sup>d</sup> | | | | |
| --- | --- | --- | --- | --- | --- | --- | --- | --- | --- |
|  |  |  | Papp<br>(nm/s)<br>AB/BA | ER | 1A2 | 2C9 | 2C19 | 2D6 | 3A4<br>(DBF) |
| **UB-MBX-46** | 15 ± 2 | CNS+ | 1.5/0.3 | 0.2 | 17 ± 1 | 33 ± 3 | 52 ± 1 | 54 ± 2 | IC<sub>50</sub>=<br>106 nM |
| **UB-MBX-47** | 8 ± 3 | CNS+ | 24/7 | 0.3 | 22 ± 3 | 43 ± 1 | 44 ± 2 | 57 ± 1 | IC<sub>50</sub>=<br>74 nM |
| **UB-ALT-P1** | 29 ± 6 | CNS+ | 38/18 | 0.5 | 33 ± 2 | 33 ± 1 | IC<sub>50</sub>=<br>137 nM | 69 ± 1 | IC<sub>50</sub>=<br>132 nM |
| **UB-ALT-P2** | 14 ± 2 | CNS+ | 137/21 | 0.2 | 21 ± 6 | IC<sub>50</sub>=<br>1.6 μM | 39 ± 1 | 20 ± 4 | IC<sub>50</sub>=<br>2.9 μM |

<sup>a</sup>Inhibition of the hERG channel was evaluated in CHO cells expressing the hERG channel using a potassium flux assay. Fluorescence recordings ( $\lambda_{\text{ex}}$ = 490 nm, $\lambda_{\text{em}}$ = 525 nm) showed minimal inhibition for all compounds ( $\leq 29 \pm 6\%$ ) when measured at 10 μM, suggesting a low risk of cardiotoxicity.
<sup>b</sup>Blood–brain barrier permeability predicted using the PAMPA-BBB assay with porcine brain lipid extract and PBS:EtOH (70:30) as buffers. Assay validation was confirmed through linear correlation with permeability values reported for 14 commercial reference drugs ( $R^2 = 0.9421$ ). All compounds were classified as CNS-permeable (CNS+; $P_e > 5.26 \times 10^{-6} \text{ cm} \cdot \text{s}^{-1}$ )<sup>38</sup> in line with the PET imaging data obtained for the UB-CB-P3 scaffold.
<sup>c</sup>Bidirectional Caco-2 permeability expressed as apparent permeability coefficients (Papp; A→B and B→A) and efflux ratio (ER = $P_{\text{appB} \rightarrow \text{A}}/P_{\text{appA} \rightarrow \text{B}}$ ). All assays were performed using a fixed concentration of 10 μM for both the test compounds and the reference standards (colchicine and E3S).
<sup>d</sup>Cytochrome P450 inhibition profile across the major human isoforms CYP1A2, CYP2C9, CYP2C19, CYP2D6 and CYP3A4 (DBF substrate) at 10 μM, using fluorescence-based probe reactions in the presence of NADPH-regenerating system. IC<sub>50</sub> values were determined when inhibition exceeded 50%.

In agreement with our PET imaging data (Figure 1B-E), effective BBB permeability was confirmed for all four compounds using a BBB-specific parallel artificial membrane permeability assay (PAMPA-BBB)^38^ (Table 1 and Supplementary Table 4). Intestinal absorption was then tested using the Caco-2 assay and all compounds exhibited low efflux ratios (0.2–0.5), indicative of negligible transporter-mediated efflux (Table 1). However, only UB-ALT-P2 exhibited a high apical-to-basolateral apparent permeability coefficient (Papp = 136.5 nm/s) together with a low basolateral-to-apical coefficient (Papp = 21.3 nm/s) across these Caco-2 epithelial monolayers, indicating rapid epithelial transit with minimal efflux liability and likely efficient *in vivo* absorption^39^. This superior permeability across Caco-2 epithelial monolayers indicates favorable intestinal absorption and identifies UB-ALT-P2 as a candidate suitable for oral administration (Table 1).

The metabolic safety profiles of our lead compounds were evaluated by measuring inhibition of five major cytochrome P450 isoforms. None of the four compounds showed any noteworthy inhibition of CYP1A2, CYP2C9, CYP2C19, or CYP2D6 (Table 1). In contrast, UB-MBX-46, UB-MBX-47, and UB-ALT-P1 each inhibited CYP3A4 with IC_50_ values between 74 nM and 132 nM. UB-ALT-P2 was significantly less active against CYP3A4 (IC_50_ = 2.9 µM), revealing a wide safety margin that reduces the risk of drug–drug interactions. Overall, these favorable DMPK properties, combined with superior potency and selectivity, positioned UB-ALT-P2 as the most promising candidate for *in vivo* studies.

### UB-ALT-P2 shows sustained binding to hP2X7Rs

Initial mutagenesis studies of hP2X7R residues involved in antagonist binding (F88A and K110A)^40^ revealed an altered IC_50_ for UB-ALT-P2 (Fig. 3D, H, I; Extended Data Fig. 6 and Supplementary Fig. 27), consistent with its action as a negative allosteric modulator. Notably, UB-ALT-P2 exhibited reduced inhibitory potency at both rat and mouse P2X7Rs in calcium influx assays (∼339-fold lower at rP2X7R than at hP2X7R and ∼92-fold lower at mP2X7R than at hP2X7R; Fig. 2H). Given these species-dependent differences in the antagonistic potency of UB-ALT-P2, we surmised that delineating ortholog-specific pharmacology would be critical for interpreting and predicting drug efficacy across animal models. We thus sought to investigate the molecular basis of this variance. We determined cryo-EM structures of UB-ALT-P2 in complex with the mP2X7R (2.3 Å) and rP2X7R (2.6 Å) and compared them to UB-ALT-P2-bound to the hP2X7R structure (2.6 Å) (Fig. 3A-C, Extended Data Fig. 2-4,7, Supplementary Table 2). Each UB-ALT-P2-bound structure resembled the apo closed state of the corresponding ortholog, with root-mean-square deviations (RMSDs) between Cα carbons of 0.5 Å for hP2X7R, 0.3 Å for mP2X7R, and 0.8 Å for rP2X7R (Extended Data Fig. 7A-C)^24,41^. The cryo-EM density for each reconstruction was sufficiently robust to confidently model UB-ALT-P2 within the classical allosteric pocket of each ortholog (Extended Data Fig. 4), classifying UB-ALT-P2 as a shallow binding ligand like other polycyclic P2X7R antagonists^24,34^.

**Figure 3:**
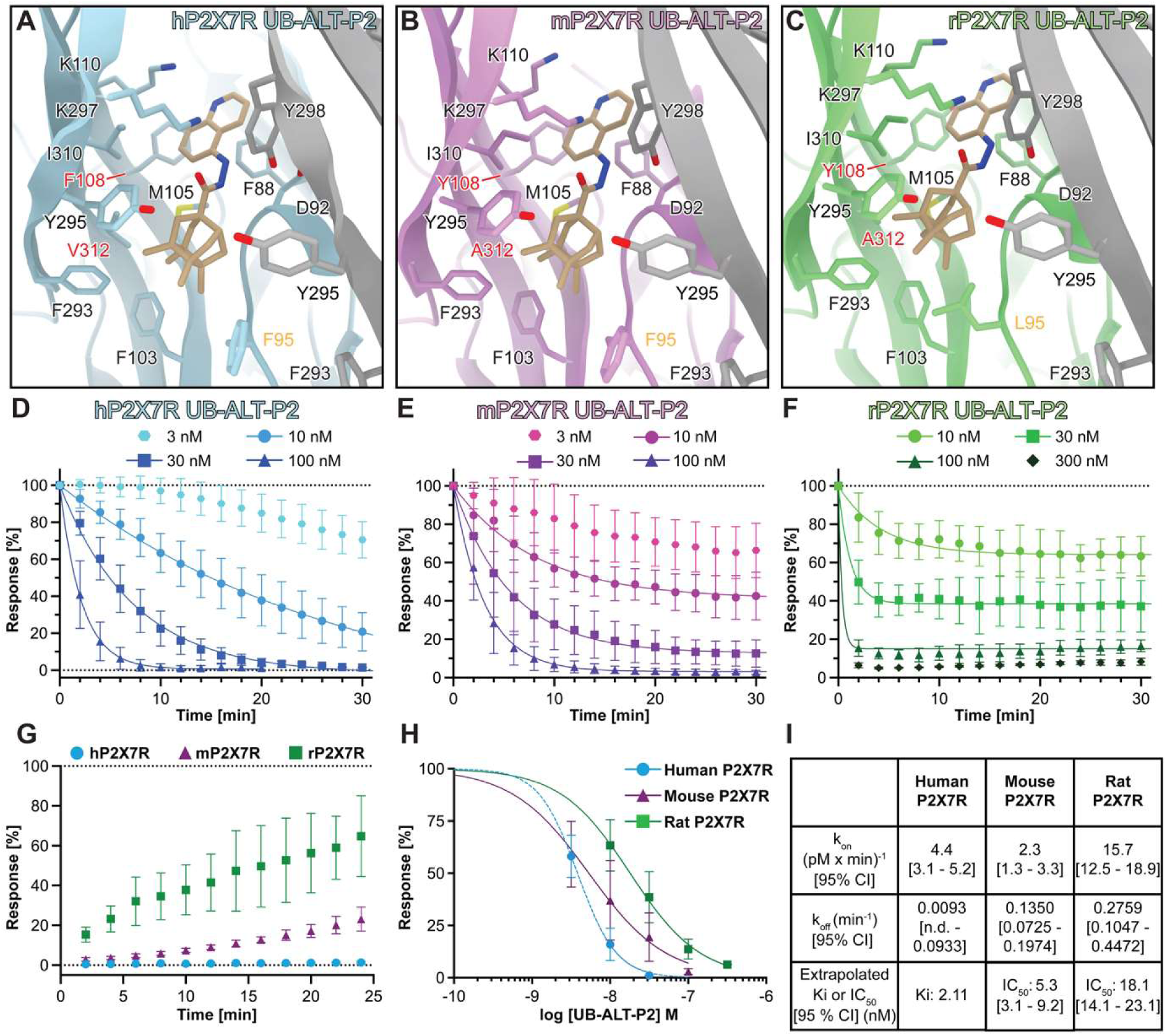
UB-ALT-P2 has distinct interactions with human, mouse, and rat P2X7Rs. (**A-C**) Ribbon representation of the classical allosteric pocket at the interface of two protomers, labeled chain A (in color) and chain C (in grey). Residues interacting with UB-ALT-P2 (carbon framework in brown sticks) shown for hP2X7R (**A**, light blue and gray), mP2X7R (**B**, light purple and gray), and rP2X7R (**C**, light green and gray). Residues 95 (gold), 108 (red), and 312 (red) are different in each ortholog and affect the pose of UB-ALT-P2. The rotameric position of L95 in rP2X7R forces UB-ALT-P2 into a distinct binding pose. (**D-I**) Time course and kinetics of UB-ALT-P2 binding to and dissociation from human, mouse, and rat orthologs. *Xenopus laevis* oocytes expressing P2X7Rs were held at −70 mV using TEVC and subjected to 5-s pulses of 300 µM ATP every 2 min. Current responses at 0 min (before continuous antagonist superfusion) were normalized to 100%. Datasets shown without fits were not included in the quantitative analysis. (**G**) Time course of antagonist dissociation. Normalized responses after a 90-100% block by the antagonists are shown. Current responses at 0 min represent responses immediately after a 2-min antagonist application. (**H**) Dose inhibition curves for UB-ALT-P2 at human and rodent P2X7Rs. Values for rodent receptors derived from equilibrium responses in (**D-F**). Values for hP2X7Rs (dotted line) were taken at 30 min due to the lack of equilibrium responses. (**I**) Rate constants and binding affinities for UB-ALT-P2 (K_i_) on human, mouse, and rat P2X7Rs. In the absence of equilibration, the K_i_ for hP2X7R was estimated by plotting experimentally determined on-rates (k_obs_) against antagonist concentrations (F) to obtain estimates for the dissociation rate constant (k_off_, y-intercept), and association rate constant (k_on_), according to the formula k_obs_=k_on_*F+k_off_. The K_i_ (2.1 nM) was then determined according to K_i_=k_off_/k_on_. Note that the estimated k_off_ (0.009 min^-1^) does not reflect the very slow dissociation from the hP2X7R, thus K_i_ is overestimated. Numbers in square brackets are 95% confidence intervals (CI). N.D., not determined. Data are mean ± SD from 3-7 measurements.

Sequence alignment of the human, mouse, and rat P2X7R orthologs revealed antagonist binding site differences at residues 95, 108, and 312 (Supplementary Table 5)^24^. Prior structural analysis of the apo closed state has suggested that a valine at position 312 significantly influences the shape of the apo binding pocket in hP2X7R by sterically forcing the side chain of Y295 to adopt an alternate rotamer (Extended Data Fig. 7D)^24^. In addition, our UB-ALT-P2-bound structures indicate that residue 95 is responsible for the lower potency at the rat receptor (Fig. 3A-C, Extended Data Fig. 7E). In contrast to the phenylalanine at position 95 in the human and mouse receptors (F95), the leucine in rP2X7R (L95) forces UB-ALT-P2 into a distinctly different pose (Extended Data Fig. 7F). If UB-ALT-P2 were to occupy the same pose as observed in hP2X7Rs or mP2X7Rs, the side chain of L95 would be just 1.3 Å away from the ligand, creating steric clashes between the ligand and receptor (Extended Data Fig. 7F).

We subsequently performed MD simulations to further investigate the comparative potency of UB-ALT-P2 across orthologs. Throughout 500 ns simulations starting from the cryo-EM structures of UB-ALT-P2 bound to hP2X7R, rP2X7R and mP2X7R, the ligand remained stably engaged within the allosteric pocket, forming similar hydrogen bonds, π–π, and hydrophobic interactions in all three complexes (Extended Data Fig. 5C-E). We therefore turned to two-electrode voltage clamp (TEVC) recordings in *Xenopus laevis* oocytes recombinantly expressing the receptor of interest to investigate potential differences in UB-ALT-P2 binding kinetics between P2X7R orthologs (Fig. 3D–I; Extended Data Fig. 6 and Supplementary Fig. 28). We observed a slow but complete block of hP2X7R currents within ∼30 min in the presence of 30 nM UB-ALT-P2 (Fig. 3D). In contrast, inhibition developed more rapidly but was less complete for the rodent orthologs, reaching ∼90% block after ∼25 min in mP2X7Rs and ∼60% after ∼5 min in rP2X7Rs (Fig. 3E-F). Recovery from UB-ALT-P2 block revealed pronounced differences for the three orthologs; rP2X7Rs showed partial recovery from UB-ALT-P2, mP2X7Rs more limited recovery, and hP2X7Rs essentially no recovery within the recording period (Fig. 3G-I).

Thus, although rodent receptors are more rapidly inhibited by UB-ALT-P2, this is offset by faster dissociation and therefore a shorter time bound to the receptor, resulting in less sustained inhibition. In contrast, the extremely slow dissociation of UB-ALT-P2 from the human receptor leads to prolonged target engagement and a near-irreversible block, providing a likely explanation for its species-dependent potency. Together, these results demonstrate that the subnanomolar potency of UB-ALT-P2 at the human receptor arises from its extraordinarily slow dissociation from the binding pocket. This is desirable for therapeutic applications, in particular for chronic diseases such as AD, because it ensures prolonged functional antagonism during dynamic fluctuations of extracellular ATP, stabilizing ion channel blockade, and providing durable suppression of downstream inflammatory signaling^42^.

### UB-ALT-P2 improves AD hallmarks *in vivo*

Having established the potency and selectivity of UB-ALT-P2 for hP2X7Rs, we began to investigate its potential utility *in vivo*. Oral administration of UB-ALT-P2 hydrochloride (3 mg/kg) to CD1 mice resulted in rapid systemic absorption and robust brain exposure (Fig. 4A). The compound reached a maximum concentration (Cmax) of 124.5 ng/mL (0.3 µM) in blood plasma within 0.5 h of administration, decreasing with an elimination half-life of 0.79 h. Brain levels reached a Cmax of 833.6 ng/mL (2.2 µM) at 0.5 h (brain-to-plasma ratio = 6.7) then decreased with an elimination half-life of 0.83 h. Despite these short systemic and brain half-lives, the slow dissociation of UB-ALT-P2 from hP2X7Rs (Fig. 3G-I) would be likely to sustain receptor occupancy beyond the period of detectable free drug, extending its pharmacodynamic effects relative to its pharmacokinetic exposure window. Moreover, because the concentration achieved in brain tissue greatly exceeds its sub-nanomolar potency, UB-ALT-P2 has the potential for full receptor occupancy *in vivo*.

**Figure 4:**
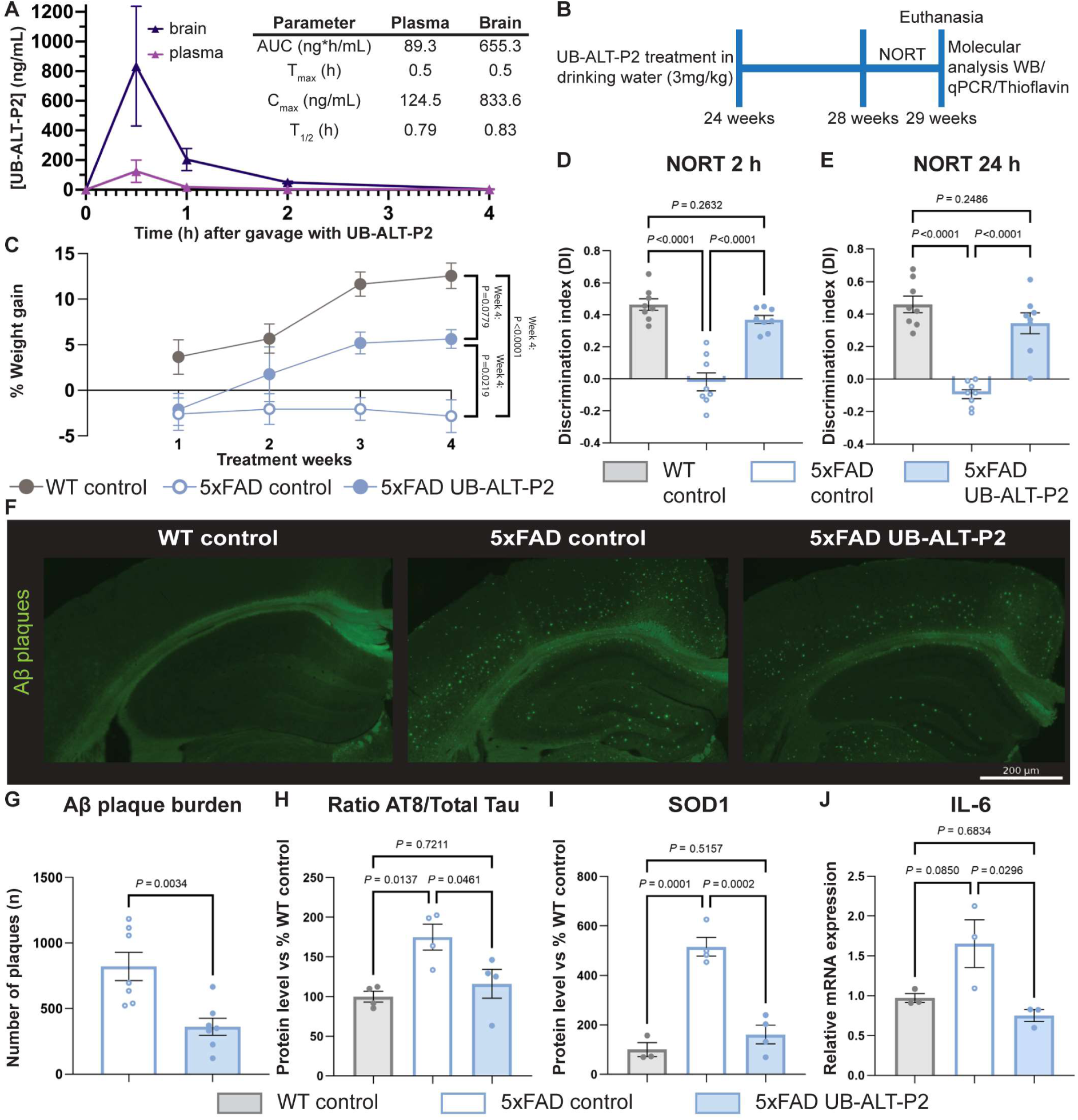
UB-ALT-P2 mitigates weight loss, memory impairment, and molecular pathology in 5xFAD mice. (**A**) Plasma and brain pharmacokinetic profile following a single dose of UB-ALT-P2 in male CD1 mice. UB-ALT-P2 (3 mg/kg) administrated by oral gavage (10% (2-hydroxypropyl)-β-cyclodextrin in physiological saline). Blood and brain samples collected at 0.5, 1, 2, 4, 8 and 10 h post-dose (n = 3 mice per time point). Plasma and brain concentrations of UB-ALT-P2 are mean ± SD. (**B**) For chronic treatment, 5xFAD mice and WT littermates received UB-ALT-P2 orally (3 mg/kg/day in drinking water) between 24-29 weeks of age. Cognitive testing (NORT) was conducted at week 28, and molecular, qPCR and neurohistological analyses followed euthanasia at week 29. (**C**) Body weight gain during UB-ALT-P2 treatment. Percent body weight change relative to baseline (body weight at treatment onset) is shown for WT control, 5xFAD control, and UB-ALT-P2–treated 5xFAD mice over the 4-week treatment. Statistical analyses performed using percent weight gain at week 4. (**D-E**) NORT performance at 2 h and 24 h post-training, reflecting short and long-term memory. UB-ALT-P2 treatment preserved memory performance of 5XFAD mice relative to the performance of WT mice at both time points. (**F-G**) Representative thioflavin S-stained hippocampal sections (**F**) show marked reduction in amyloid-β plaque burden in UB-ALT-P2-treated 5xFAD mice compared to transgenic controls (scale bar, 200 µm). Quantification (**G**) shows significant decrease in plaques in UB-ALT-P2-treated 5xFAD animals. (**H**) Tau pathology quantified as the ratio of phosphorylated tau (AT8) to total tau and values were expressed relative to the ratio in WT mice. UB-ALT-P2 treatment significantly reduced tau phosphorylation (lower AT8/total tau ratio) compared to untreated 5xFAD mice. (**I**) Oxidative stress assessed by measuring the SOD1 protein levels, which were quantified and expressed relative to the SOD1 expression level in WT mice. 5xFAD mice showed a pronounced upregulation of SOD1 levels that was markedly reduced by UB-ALT-P2 treatment. (**J**) Neuroinflammation evaluated by expression of IL-6, which showed a non-significant increase in 5xFAD mice compared to WT controls that was significantly reduced by UB-ALT-P2 treatment. Data in (**D–J**) are mean ± SEM with individual values overlaid. Statistical significance in each panel was determined by ANOVA followed by Tukey’s post hoc test (**D, E, H, I, J**) or two-tailed Student’s t-test (**G**).

To evaluate the therapeutic potential of UB-ALT-P2 *in vivo*, we leveraged the 5xFAD mouse model of AD, which recapitulates key pathological and behavioral aspects of the human disease, including severe amyloid pathology, synaptic dysfunction, and progressive cognitive decline, within approximately six months^43^. UB-ALT-P2 was administered orally via drinking water (3 mg/kg) between 24 and 28 weeks of age, after which behavioural and molecular endpoints were assessed (Fig. 4B). Consistent with previous reports^44^, untreated 5xFAD mice exhibited attenuated weight gain relative to wild-type (WT) control mice, reflecting a systemic disease-associated phenotype. Notably, UB-ALT-P2 treatment significantly improved weight gain at the end of the four-week dosing period, restoring a growth trajectory comparable to that of WT mice (Fig. 4C).

We next examined whether this systemic benefit extended to cognitive function. As expected, vehicle-treated 5xFAD mice displayed a marked reduction of the discrimination index (DI) in the novel object recognition test (NORT)^45^ compared with WT controls (*P* < 0.0001), indicative of impaired recognition memory (Fig. 4D,E and Extended Data Fig. 8A,B). In contrast, UB-ALT-P2–treated 5xFAD mice showed a significant improvement in DI in both short-term (2 h) and long-term (24 h) memory tests (*P* < 0.0001), consistent with a slowing or prevention of memory decline (Fig. 4D,E). Strikingly, the performance of 5xFAD mice treated with UB-ALT-P2 was statistically indistinguishable from that of WT controls at both time points, indicating near-complete preservation of recognition memory.

Because extracellular amyloid-β (Aβ) plaque deposition is an early hallmark of AD pathology, and thioflavin S is widely used to visualize fibrillar amyloid, we employed thioflavin S staining to quantify amyloid deposition in the cortex and hippocampus of 5xFAD mice^46^. We observed a significantly reduced number of thioflavin S-positive plaques after UB-ALT-P2 treatment (*P* < 0.01), indicating a reduction in amyloid pathology (Fig. 4F,G). A reduction in plaque burden is mechanistically relevant and may be clinically meaningful, as decreased Aβ plaque deposition has been associated with improvements in synaptic plasticity, reduced neuronal loss and attenuation of chronic neuroinflammation^16^.

Although 5xFAD mice do not develop significant endogenous tau pathology or neurofibrillary tangles characteristic of human AD or tau transgenic models^47–48^, we evaluated the effect of UB-ALT-P2 on tau pathology by quantifying the ratio of hyperphosphorylated tau (at S202 and T205) to total tau in hippocampal tissue using the AT8 monoclonal antibody in western blots^48^. Interestingly, the 5xFAD control group had a significantly elevated ratio of phosphorylated tau compared to WT controls (*P* < 0.05) (Fig. 4H). Treatment with UB-ALT-P2 (3 mg/kg) significantly reduced the AT8:total tau ratio in 5xFAD mice (*P* < 0.05) to levels that were not significantly different from WT mice (Fig. 4H and Extended Data Fig. 8C). We observed a non-significant trend towards increased tau phosphorylation at S404 (p-S404/total tau ratio) in 5xFAD mice relative to WT controls, along with a trend towards reduced S404 phosphorylation following UB-ALT-P2 treatment (Extended Data Fig. 8D). Given that this study was conducted at an established yet still progressive stage of disease in the 5xFAD model, characterized by robust amyloid pathology but limited secondary tau changes, the absence of a significant increase in S404 phosphorylation was not unexpected^49^.

Finally, we investigated the effect of UB-ALT-P2 on cellular stress *in vivo*, first by measuring the expression levels of superoxide dismutase 1 (SOD1), a hallmark of oxidative stress and AD pathophysiology^50^. As expected, 5xFAD control mice had significantly elevated (*P* < 0.001) SOD1 expression (Fig. 4I and Extended Data Fig. 8E). Treatment with UB-ALT-P2 for five weeks significantly reduced (*P* < 0.001) SOD1 levels to values that were statistically indistinguishable from WT, indicating normalization of the oxidative environment (Fig. 4I). Similarly, hippocampal mRNA levels of the pro-inflammatory cytokines IL-6 and IL-1β were increased in 5xFAD mice relative to WT controls (Fig. 4J and Extended Data Fig. 8F), consistent with the development of chronic neuroinflammatory signalling^6^. IL-6 mRNA expression exhibited a non-significant trend towards elevation in vehicle-treated 5xFAD mice compared to WT mice, which was significantly reduced (*P* < 0.05) by UB-ALT-P2 treatment (Fig. 4J). In contrast, IL-1β mRNA levels were significantly increased in 5xFAD mice (*P* < 0.05) relative to WT controls, and UB-ALT-P2 treatment reduced this expression but not significantly. Nonetheless, IL-1β mRNA expression in UB-ALT-P2-treated 5xFAD mice did not differ significantly from levels in WT controls (Extended Data Fig. 8F). Together, these data indicate that UB-ALT-P2 attenuates inflammatory signalling in the hippocampus of 5xFAD mice, consistent with a reduced neuroinflammatory burden.

## Conclusion

Our structure-based drug design pipeline, from medicinal chemistry to *in vivo* efficacy, has led to the development of UB-ALT-P2, a hP2X7R antagonist with sub-nanomolar potency, high selectivity, prolonged target residence time, favorable DMPK properties, and BBB penetrance (Fig. 5). Cryo-EM and TEVC analysis suggested a basis for cross-species potency of UB-ALT-P2 and defined the molecular determinants of efficacious pharmacophores from preclinical animal models to humans. Oral treatment of 5xFAD mice with UB-ALT-P2 ameliorated multiple AD-associated hallmarks, including weight loss, cognitive impairment, Aβ plaque deposition, tau hyperphosphorylation, oxidative stress, and neuroinflammation (Fig. 5). Together, these data reveal the therapeutic potential of UB-ALT-P2 for modulating the molecular pathways associated with AD pathology. Given the aging population and limited treatment options for AD, this study positions UB-ALT-P2 as a promising therapeutic with potential for further preclinical development towards clinical translation.

**Figure 5:**
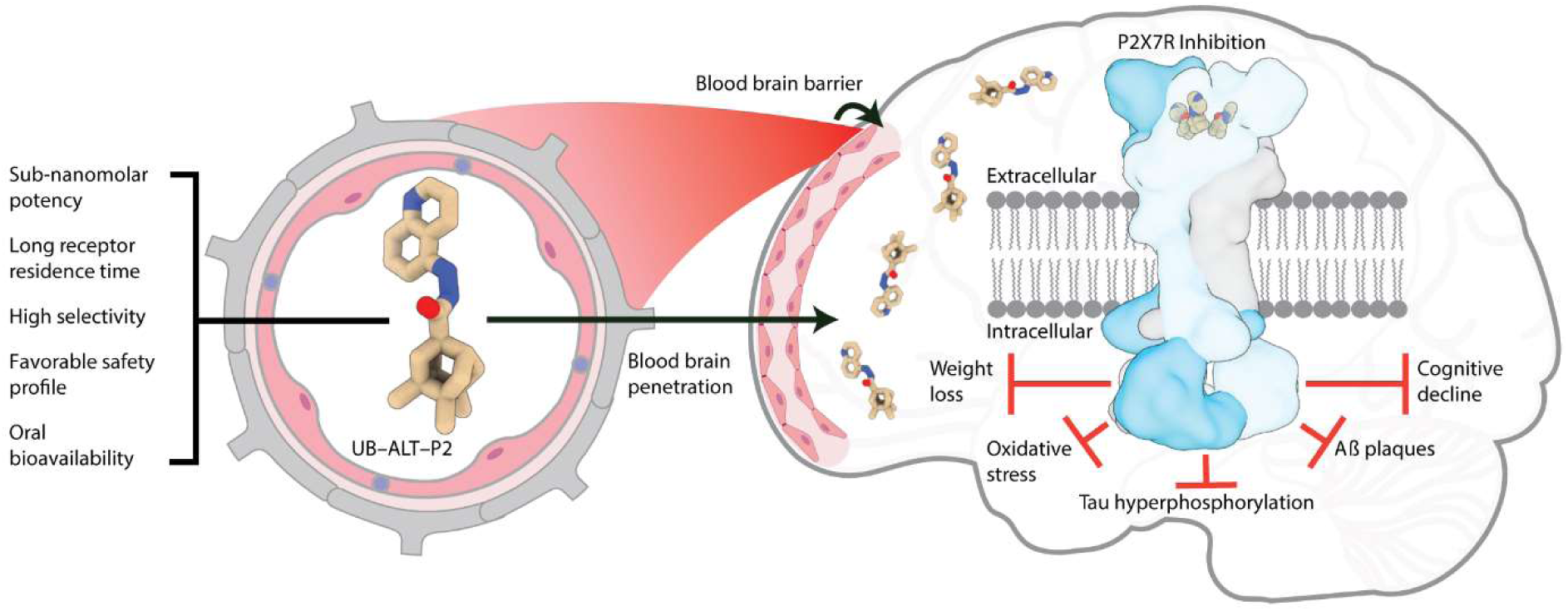
Potent, selective, and safe antagonism of the P2X7R is a promising therapeutic approach for AD. Structure-guided medicinal chemistry led to the development of UB-ALT-P2, a hP2X7R antagonist with sub-nanomolar potency, long receptor residence time, high selectivity, and favorable DMPK properties including oral bioavailability and blood–brain barrier penetrance. Oral administration of UB-ALT-P2 to 5xFAD mice ameliorated multiple AD–associated hallmarks, including weight loss, cognitive deficits, amyloid-β plaque burden, tau hyperphosphorylation, oxidative stress and neuroinflammation, supporting the therapeutic potential of P2X7R inhibition for AD.

## Methods

### Chemistry. General methods

Chemicals and solvents were purchased from commercial suppliers (Merck, Fluorochem, Fisher, Sigma-Aldrich) and used without further purification unless otherwise specified. 400 MHz ^1^H NMR and 100.6 MHz ^13^C NMR were recorded on a Varian Mercury 400. 500 MHz ^1^H NMR and 125.7 MHz ^13^C NMR were recorded on a Bruker Avance Neo 500. The chemical shifts are reported in ppm (δ scale) relative to internal tetramethylsilane, or to solvent peak, and coupling constants are reported in Hertz (Hz). Assignments given for the NMR spectra of the new compounds have been carried out based on homocorrelation ^1^H/^1^H (COSY) and/or heterocorrelation ^1^H/^13^C (HSQC) experiments. The used abbreviations are: s, singlet; d, doublet; t, triplet; q, quadruplet; m, multiplet; cs, complex signal; broad s, broad singlet, or combinations thereof. IR spectra were run on a FTIR Perkin-Elmer Spectrum RX I or a Perkin-Elmer Spectrum TWO spectrophotometers, using sodium chloride (NaCl) pellets or attenuated total reflectance (ATR) techniques. Absorption values are expressed as wavenumbers (cm^-1^); only significant absorption bands are given. Column chromatography was performed on silica gel 60 Å (Sigma Aldrich, 40 - 63 μm, 230-400 mesh) or with a CombiFlash Rf 150 Teledine ISCO provided with a UV-vis detector, with RediSep^®^ Rf Normal-phase Silica Flash Columns (silica gel 60 Ǻ, 35-70 μm, 230-400 mesh). Thin Layer Chromatography (TLC) was performed with aluminium-backed sheets with silica gel 60 F254 (Merck, ref 1.05554), and spots were visualized with UV light, 1% aqueous solution of KMnO_4_, iodine or ninhydrin. Melting points were determined in open capillary tubes with an MFB 59510M Gallenkamp melting point apparatus. Accurate mass spectra were recorded with ESI techniques on a Hewlett-Packard 5988a LC/MSD-TOF instrument at *Unitat d’Espectrometria de Masses dels Centres Científics i Tecnològics de la Universitat de Barcelona* (CCiTUB). The analytical samples of all the new compounds, which were subjected to pharmacological evaluation, possessed purity ≥95% as evidenced by their HPLC-UV or their elemental analyses. HPLC-UV analyses were performed using an Agilent 1260 Infinity II LC/MSD coupled to a photodiode array and a mass spectrometer. Samples (5 μL, 0.5 mg/mL) in a 1:1 mixture of water with 0.05% formic acid (A) and acetonitrile with 0.05% formic acid (B) were injected using an Agilent Poroshell 120 EC-C18 (2.7 μm, 50 mm × 4.6 mm) column at 40 °C. The mobile phase was a mixture of A and B, with a flow 0.6 mL/min, using the following gradients: from 95% A–5% B to 100% B in 3 min; 100% B for 3 min; from 100% B to 95% A–5% B in 1 min; and 95% A–5% B for 3 min. Purity is given as % of absorbance at 254 nm.

### Radiochemistry

All procedures involving the manipulation of radioactive substances were performed in an authorised radiation-controlled facility (IRA 2916) under the required radiological protection and safety conditions. Ultrapure water was produced using a Milli-Q^®^ purification system (Millipore^®^, Merck KGaA, Darmstadt, Germany). Solid-phase extraction (SPE) cartridge (Sep-Pak^®^ Light C18 Plus) was obtained from Waters (Waters Corporation, MA, USA). Prior to use, the C18 cartridges were preconditioned with 10 mL of ethanol followed by 10 mL of ultrapure water.

### Measurement of ethidium bromide accumulation in hP2X7R-expressing HEK293 cells

HEK293 cells expressing hP2X7R were cultured in a humidified atmosphere of 5% CO_2_ at 37 °C in Dulbecco’s Modification of Eagle’s Medium (DMEM; Corning) supplemented with 10% (v/v) fetal bovine serum (Corning) and 1% (v/v) antibiotic–antimycotic (Gibco)^51^. To perform the assay, the DMEM was removed, and the HEK293 cells were washed with 3 mL of 1X DPBS (Dulbecco’s Phosphate-Buffered Saline; Corning). After the removal of DPBS solution, the cells were detached from the dish with 1 mL of trypsin/EDTA (Gibco), and the cells were collected by centrifugation. The cells (density of 2.25 × 10^7^ cells/mL) were re-suspended in 4-(2-hydroxyethyl)-1-piperazineethanesulfonic acid (HEPES)-buffered salt solution consisting of 140 mM potassium chloride, 1 mM ethylenediaminetetraacetic acid (EDTA), 5 mM glucose, 20 mM HEPES and 0.1 mM ethidium bromide (pH 7.4). The vehicles or appropriate range of concentrations of compounds (10 μL, pre-diluted in 10% (v/v) DMSO in DW from 10 mM stock) were added to each well of 96-well black plate (Corning), and 80 μL of the cell suspension was subsequently added to each well. BzATP (10 μL, pre-diluted in DW from 10 mM stock) was then added, and the final assay volume was maintained at 100 μL. After the incubation in 5% CO_2_ at 37 °C for 2 hours, the ethidium dye uptake was detected by measuring the fluorescence (excitation wavelength of 530 nm and emission wavelength of 590 nm) of each well using a CHAMELEON™ Multi-Technology Plate Reader. The antagonistic activities of compounds are expressed as the percentages relative to the maximum accumulation of ethidium bromide observed in the control group with the stimulation of the hP2X7R by an agonist, BzATP. The IC_50_ values of compounds were calculated using nonlinear regression analysis using OriginPro 9.1 software.

### PET biodistribution studies

Animals (B6129PF2/J mice; n=2; age = 24 months) were maintained and handled in accordance with European Council Directive 2010/63/UE at CIC biomaGUNE (San Sebastián, Spain) facilities. Experimental procedures were approved by the CIC biomaGUNE Ethical Committee and the corresponding local authorities (authorisation number PRO-AE-SS-207). Animals were housed in ventilated cages and provided with a standard diet.

Biodistribution studies were carried out via positron emission tomography (PET) imaging. Mice were anesthetized with 4% isoflurane in pure O₂ for induction and maintained under 1–2% isoflurane during imaging. Body temperature and respiration were continuously monitored throughout the scans. [¹¹C]UB-CB-P3 (2.09 ± 0.17 MBq in 10% ethanol/saline) was intravenously administered via one of the lateral tail-veins using a pre-inserted cannula, concomitantly with the start of a dynamic PET acquisition using a β-CUBE imaging system (Molecubes, Ghent, Belgium). Immediately after the PET session, a 5-min whole-body computerised tomography (CT) scan was acquired on the X-CUBE platform of the same vendor. PET data were reconstructed using a 3D-OSEM iterative image reconstruction algorithm and analysed with PMOD software (PMOD Technologies, Switzerland). Volumes of interest (VOIs) were drawn in the brain, heart, lungs, liver, kidneys, bladder and gallbladder. Dynamic time–activity curves (TACs) were generated over the entire scan duration with GraphPad Prism 9 (version 9.5.0). Radioactivity concentration values were expressed as Bq/cm³ and normalized to injected amount of radioactivity and body weight to calculate standard uptake values (SUVs).

### Calcium influx assays

Calcium influx assays for UB-MBX-46, UB-MBX-47, UB-ALT-P1 and UB-ALT-P2 at human P2X1, P2X2, P2X3, P2X4 and P2X7 receptors, and additionally at rat and mouse P2X7 receptors, were performed as previously described^24,52,53^. Recombinant 1321N1 astrocytoma cells stably expressing the respective receptor were employed^24,52,53^.

For the slowly or non-desensitizing P2X2R, P2X4R, and P2X7R subtypes, we used Fluo-4 acetoxymethyl ester (Fluo-4 AM, Thermo Fisher Scientific, Dreieich, Germany) for calcium measurements. For P2X1R and P2X3R, the FLIPR^®^ Calcium 5 Assay Kit (Molecular Devices, San José, CA, USA) was utilized. Cells (30,000–45,000 per well) were seeded in a volume of 200 µL into black, flat-bottom 96-well polystyrene microplates and incubated at 37 °C and 10% CO_2_ for ca. 12 h. Then, the medium was removed, and the cells were incubated for 1h with dye solution (3 µM of Fluo-4 AM and 1% Pluronic F-127 at room temperature, or Calcium 5 dye at 37°C, respectively). The assays were performed in Hanks’ balanced salt solution (HBSS, Thermo Fisher Scientific, Dreieich, Germany), except for the P2X7R assays (buffer composition: 150 mM sodium-glutamate, 5 mM KCl, 0.5 mM CaCl_2_, 0.1 mM MgCl_2_, 10 mM D-glucose, 25 mM 4-(2-hydroxyethyl)-1-piperazineethanesulfonic acid (HEPES), pH 7.4). Then, the dye solution was removed, and the buffer solution was added. Antagonists were dissolved in DMSO (final concentration: 0.5-1%) and initially screened at 10 µM concentration. Cells were preincubated with antagonist for 30 min before stimulation with agonist at its EC_80_ concentration, and fluorescence was measured using a plate reader (NOVOstar, BMG Labtech GmbH, Offenburg, Germany, excitation at 485 nm, emission at 520 nm, for 30 s with 0.4 s intervals). Concentration-dependent inhibition was determined at human, rat and mouse P2X7 receptors, and IC_50_ values were calculated. For selected compounds that showed significant inhibition at the other human P2XR subtypes, IC_50_ values were also determined. Data were analyzed using Microsoft Excel and GraphPad Prism (Version 8.0, San Diego, CA, USA). The IC_50_ value for each compound was calculated by nonlinear regression with a sigmoidal dose-response equation.

### Parallel Artificial Membrane Permeation Assays-Blood-Brain Barrier (PAMPA-BBB)

To evaluate the brain penetration of the different compounds, a parallel artificial membrane permeation assay for blood-brain barrier was used, following the method described by Di *et al*.^38^ The *in vitro* permeability (*P*_e_) of fourteen commercial drugs through lipid extract of porcine brain membrane together with the test compounds was determined. Commercial drugs and assayed compounds were tested using a mixture of PBS:EtOH (70:30). The assay was validated by comparing the experimentally determined permeability (*P*_e_) values of fourteen commercial drugs with permeability values reported in the literature. The relationship between experimental and reported *P*_e_ values showed a strong linear correlation in the PAMPA-BBB assay (y = 1.5714x − 1.1076; R² = 0.9421). From this equation and taking into account the limits established by Di et al. for BBB permeation, we established the following ranges of permeability: compounds of high BBB permeation (CNS +): *P*e > 5.178 x 10^-6^ cm s^-1^; compounds of low BBB permeation (CNS ‒): *P*e < 2.035 x 10^-6^ cm s^-1^; and compounds of uncertain BBB permeation (CNS ±): 5.178 x 10^-6^ cm s^-1^) > *P*e > 2.035 x 10^-6^ cm s^-1^ (see Supplementary Table 4).

### Cytochrome P450 inhibition assay

The objective of this study was to screen the potential of each compound to inhibit cytochrome P450 using recombinant human cytochrome P450 enzymes (CYP1A2, CYP2C9, CYP2C19, CYP2D6 and CYP3A4 from BioIVT, https://bioivt.com) and probe substrates with fluorescent detection; CYP3A4 was measured using dibenzylfluorescein (DBF). Incubations were conducted in a 200 µL volume in 96-well microtiter plates (COSTAR 3915). The addition of the mixture buffer-cofactor (KH_2_PO_4_ buffer, 1.3 mM NADP, 3.3 mM MgCl_2_, 3.3 mM glucose-6-phosphate and 0.4 U/mL glucose-6-phosphate dehydrogenase), control microsomes, standard inhibitors (furafylline, tranylcypromine, ketoconazole, sulfaphenazole and quinidine; Sigma Aldrich), and previously diluted compound to plates was carried out by a liquid handling station (Zephyr Caliper). The plate was then preincubated at 37 °C for 5 min, and the reaction was initiated by the addition of prewarmed enzyme/substrate (E/S) mix. The E/S mix contained buffer (KH_2_PO_4_), c-DNA-expressed P450 in insect cell microsomes, substrate (3-cyano-7-ethoxycoumarin, for CYP1A2 and CYP2C19; 7-methoxy-4-(trifluoromethyl)coumarin for CYP2C9; 3-[2-(N,N-diethyl-N-methylammonium)ethyl]-7-methoxy-4-methylcoumarin for CYP2D6; and dibenzylfluorescein (DBF) for CYP3A4) in a reaction volume of 200 µL. Reactions were terminated after various times (a specific time for each cytochrome) by addition of STOP solution (ACN/TrisHCl 0.5 M 80:20, and 2 N NaOH for CYP3A4). Fluorescence per well was measured using a fluorescence plate reader (Tecan Infinity M1000 pro or EnVision 210 multilabel Reader) and percentage of inhibition was calculated.

### hERG inhibition assay

The assay was carried out using a CHO cell line transfected with the hERG potassium channel (CHO hERG-Duo Cell Line, B’SYS GmbH, Switzerland). At 72 h prior to running the assay, 2500 cells were seeded on a 384-well black plate (Greiner 781091). The cell line was maintained at 37°C in a 5% CO_2_ atmosphere for 24h and at 30°C in a 5% CO_2_ atmosphere for an additional 48 h. hERG activity was measured by using the FluxorTM Potassium Ion Channel Assay Kit (Thermo Fisher F10016). Medium was replaced with 20 μl Loading Buffer and the cells were incubated for 60 min at RT, protected from direct light. After incubation, Loading Buffer was replaced with Assay Buffer and the compounds were incubated for 30 min at RT. Stimulus Buffer (5μl) was added to each well and the fluorescence was read (λ_ex_=490 nm, λ_em_=525nm) using an imaging plate reader system (FDSS7000EX, Hamamatsu^®^) every second after the establishment of a baseline signal. Loading buffer, Assay Buffer and Stimulus buffer were prepared from the reagents provided in the kit.

### Caco-2 cells permeability

The Caco-2 cells were cultured to confluency, trypsinized and seeded onto a filter transwell inserted at a density of ∼10,000 cells/well in DMEM cell culture medium. Confluent Caco-2 cells were sub-cultured at passages 58-62 and grown in a humidified atmosphere of 5% CO_2_ at 37°C. Following an overnight attachment period (24 h after seeding), the cell medium was replaced with fresh medium in both the apical and basolateral compartments every other day. The cell monolayers were used for transport studies 21 days post seeding. The monolayer integrity was checked by measuring the transepithelial electrical resistance (TEER) obtaining values ≥ 500 Ω/cm^2^. On the day of the study, after the TEER measurement, the medium was removed and the cells were washed twice with pre-warmed (37°C) Hank’s Balanced Salt Solution (HBSS) buffer to remove traces of medium. Stock solutions were made in dimethyl sulfoxide (DMSO) and further diluted in HBSS (final DMSO concentration 1%). Each compound and reference compounds (Colchicine, E3S) were all tested at a final concentration of 10 μM. For A → B directional transport, the donor working solution was added to the apical (A) compartment and the transport media as receiver working solution was added to the basolateral (B) compartment. For B → A directional transport, the donor working was added to the basolateral (B) compartment and transport media as a receiver working solution was added to the apical (A) compartment. The cells were incubated at 37°C for 2 hours with gentle stirring. At the end of the incubation, samples were taken from both donor and receiver compartments and transferred into 384-well plates and analyzed by UPLC-MS/MS. The detection was performed using an ACQUITY UPLC/Xevo TQD System. After the assay, Lucifer yellow (LY) was used to further validate the cell monolayer integrity, cells were incubated with LY 10μM in HBSS for 1hour at 37°C, obtaining permeability (Papp) values for LY of ≤ 10 nm/s confirming the well-established Caco-2 monolayer.

### Mammalian cell toxicity

Four mammalian cell lines were used to evaluate the potential cytotoxicity of the compounds: human embryonic lung (HEL) fibroblast cells, human cervix carcinoma-derived HeLa cells, human T-cell leukemia-derived MT4 cells, and African Green monkey kidney-derived Vero cells. Semi-confluent cell cultures in 96-well plates were exposed to serial dilutions of the compounds or to medium (= no compound control), then incubated at 37°C. Four days later, the cells were inspected by microscopy to determine the MCC, that is, the compound concentration that causes a microscopically detectable alteration of normal cell morphology. Next, the MTS cell viability reagent (CellTiter 96 AQueous MTS Reagent from Promega) was added. After 4 hours of incubation at 37°C, the optical density (OD) at 490 nm was recorded in a microplate reader. For each concentration, the percentage cytotoxicity was calculated as: [1 − (OD_Cpd_)/(OD_Contr_)] × 100, in which OD_Cpd_ equals the OD value at the corresponding compound concentration and OD_Contr_ equals the OD value for the no compound control. Next, the half-maximal cytotoxic concentration (CC_50_) was derived by extrapolation, assuming a semi-log dose response.

### Cell lines for large-scale receptor expression

SF9 cells were cultured in SF-900 III SFM (Fisher Scientific) at 27°C. Cells of female origin were used for the expression of baculovirus. HEK293S GnTI^-^ cells were cultured using Gibco Freestyle 293 Expression Medium (Fisher Scientific) at 37°C supplemented with 2% v/v fetal bovine serum (FBS). HEK293S GnTI^-^ cells of female origin were used to express P2X7Rs.

### Receptor constructs

The full-length wild-type hP2X7R, mP2X7R, and rP2X7R constructs used for protein expression contain a C-terminal HRV 3-C protease site, GFP, and a histidine affinity tag for purification. For structural determination, all tags were removed during purification, and no mutations or truncations were made to the receptor.

### Receptor expression and purification

The full-length wild-type hP2X7R, mP2X7R, and rP2X7R constructs were expressed by baculovirus mediated gene transfection (BacMam) using similar protocols as previously outlined for the hP2X7R, the mP2X7R, and the rP2X7R^24,34,41,54,55^. Briefly, HEK293S GnTI^-^ cells were grown in suspension to a sufficient density and infected with P2 BacMam virus. Following overnight expression at 37 °C, sodium butyrate was added to a final concentration of 10 mM, and the cells were shifted to 30 °C for an additional two days. Next, the cells were harvested by centrifugation, washed with PBS buffer (137 mM NaCl, 2.7 mM KCl, 8 mM Na_2_HPO_4_, 2 mM KH_2_PO_4_), suspended in TBS (50 mM Tris pH 8.0, 150 mM NaCl) containing protease inhibitors (1 mM PMSF, 0.05 mg/mL aprotinin, 2 µg/mL Pepstatin A, 2 µg/mL leupeptin), and broken via sonication. Intact cells and cellular debris were removed by centrifugation. The membranes were isolated by ultracentrifugation and snap frozen and stored at −80 °C until use.

When ready, membranes were thawed, resuspended in TBS buffer containing 15% glycerol, dounce homogenized, and solubilized. For the mP2X7R and the rP2X7R, the homogenized membranes were solubilized in 40 mM dodecyl-β-D-maltopyranoside (DDM or C12M) and 8 mM cholesterol hemisuccinate tris salt (CHS) while the hP2X7R was solubilized in 40 mM DDM and ∼17 mM CHS. After ultracentrifugation, the soluble fraction was incubated with TALON resin in the presence of 10 mM imidazole at 4 °C for 1–2 h. Next, the resin was loaded into an XK-16 column and washed with 2 column volumes of buffer (TBS plus 5% glycerol, 1 mM C12M, 0.2 mM CHS at pH 8.0) containing 20 mM imidazole, 10 column volumes containing 30 mM imidazole, and eluted with buffer containing 250 mM imidazole. Peak fractions containing the fusion protein were concentrated and digested with HRV 3-C protease (1:25, w/w) overnight at 4 °C to remove GFP and the histidine affinity tag. To isolate the trimeric receptor from GFP and HRV 3-C protease, the digested protein was ultracentrifuged and injected onto a Superose 6 10/300 GL column for size exclusion chromatography (SEC) that was equilibrated with 20 mM HEPES, pH 7.0, 100 mM NaCl, and 0.5 mM C12M. Fractions were analyzed by fluorescence size exclusion chromatography (FSEC) and selected fractions were concentrated for cryo-EM grid preparation.

### Electron microscopy sample preparation

To prepare antagonist-bound samples, the purified receptor was incubated with antagonist at 3-4x the molar concentration of the P2X7R monomer. Following a one-hour incubation period at 4 °C, cryo-EM grids were prepared for each sample condition. For all samples, 2.5 μL of solution was applied to glow-discharged (15 mA, 1 min) Quantifoil R1.2/1.3 300 mesh gold holey carbon grids which were blotted for 1.5 s or 2.0 s under 100% humidity at 6 °C. The grids were flash frozen in liquid ethane using a FEI Vitrobot Mark IV and stored under liquid nitrogen until screening and large-scale data acquisition.

### Electron microscopy data acquisition

Cryo-EM data for all receptor orthologs and complexes were collected on Titan Krios microscopes (FEI) operated at 300 kV at the Pacific Northwest Center for Cryo-EM (PNCC). Datasets were acquired hardware-binned (hP2X7R UB-MBX-47) or in super-resolution mode (hP2X7R UB-ALT-P1, hP2X7R UB-ALT-P2, mP2X7R UB-ALT-P2, and rP2X7R UB-ALT-P2) on Gatan K3 direct-electron detectors using defocus values that ranged between -0.7 µm and -1.5 µm. Movies for each dataset contain between 43 and 50 frames (hP2X7R UB-MBX-47: 50 frames, hP2X7R UB-ALT-P1: 50 frames, hP2X7R UB-ALT-P2: 43 frames, mP2X7R UB-ALT-P2: 50 frames, rP2X7R UB-ALT-P2: 48 frames) and a total dose ranging between 42 to 46 e^-^/ Å^2^. All datasets used an energy filter and nominal magnification of 130,000x, corresponding to a physical pixel size of ∼0.648 Å/pixel. Each dataset utilized ‘multi-shot’ and ‘multi-hole’ collection schemes driven by serial EM^56^.

### Electron microscopy data processing

Movies were imported and patch motion corrected in cryoSPARC, outputting micrographs at the physical pixel size (∼0.648 Å/pixel) (Extended Data Fig. 2)^57^. Following estimation of the contrast transfer function (CTF) parameters, bad micrographs were culled and particles picked using 2D templates generated from a 3D volume. Next, particles were inspected, extracted, and sent directly to 3D classification consisting of *ab initio* jobs to generate reference volumes and heterogeneous classifications to remove bad particles (Extended Data Fig. 2, Supplementary Table 2). No 2D classifications were performed for any of the datasets. Consensus cryo-EM reconstructions from the final particle stacks were generated by non-uniform refinements with global and local CTF refinements (Extended Data Fig. 2, 3, Supplementary Table 2)^58^. Sharpened maps were generated using either B-factor sharpening in cryoSPARC or DeepEMhancer^59^.

### Model building and structure determination

The cryo-EM structure of the hP2X7R in the UB-MBX-46-bound inhibited state was used for the initial model for each antagonist-bound hP2X7R structure (PDB code: 9E3P)^24^. The cryo-EM structure of the mP2X7R in the apo closed state was used for the initial mode of the mP2X7R in the UB-ALT-P2-bound inhibited state (PDB code: 9E3Q)^24^. The cryo-EM structure of the rP2X7R in the AZD9056-bound inhibited state was used for the initial mode of the rP2X7R in the UB-ALT-P2-bound inhibited state (PDB code: 8TR8)^34^. Each initial model was then docked into the corresponding cryo-EM reconstruction using ChimeraX and model building was performed using COOT, PHENIX, and ISOLDE^60–64^. All ligands were built and refined using eLBOW with protonation states corresponding to approximately pH 7^65^. Limited glycosylation, acylation, and palmitoylation were included in models when justified by the corresponding cryo-EM density. Model quality was evaluated by MolProbity (Supplementary Table 2)^66^.

### Protein preparation for Molecular Dynamics (MD) simulations

Cryo-EM structures of the hP2X7R, mP2X7R, and rP2X7R in complex with UB-ALT-P2 allosteric ligand were prepared for simulations using the Protein Preparation Wizard implementation in Schrödinger suite (Protein Preparation Wizard 2015-2; Schrödinger, LLC, New York, 2015). In this process, the bond orders and disulfide bonds were assigned, and missing hydrogen atoms and loops were added using Prime. Additionally, N- and C-termini of the protein model were capped by acetyl and N-methyl-amino groups, respectively. The ionization states of the compounds at pH 7.5 were confirmed based on the Epik^67^ utility. The protein complex was subjected to an all-atom minimization using the OPLS2005 force field^68^ with heavy atom RMSD values constrained to 0.30 Å.

### System setup for MD simulations with Desmond

The experimental structure of the hP2X7R, mP2X7R, or rP2X7R with or without bound cholesterols was inserted in a pre-equilibrated hydrated 1-palmitoyl-2-oleoyl-*sn*-glycero-3-phosphocholine (POPC) bilayer expanding from the furthermost vertex of the protein to the simulation orthorhombic box edge 30 Å to all axes. The protein was positioned with respect to the membrane plane (x,y plane) as suggested by the server “Orientations of Proteins in Membranes (OPM)” database^69^ using the System setup utility in Schrödinger Maestro software (Schrödinger Release 2020-2: Maestro, Schrödinger, LLC, New York, 2020) using the System Builder Wizard in Maestrο software (Schrödinger Release 2020-4: Desmond Molecular Dynamics System, D. E. Shaw Research, New York, NY, 2021). Using the same utility, the TIP3P^70^ solvation model was applied while sodium and chloride ions were added randomly in the water phase to neutralize the system and reach the experimental salt concentration of 0.150 M NaCl. The resulting lipid buffer contained approximately ∼ 542,000 atoms, consisting of 711 POPC lipids and ∼ 139,300 water molecules. Periodic boundary conditions were applied, and the dimensions of the simulation box was 162×162×217 Ǻ^3^. The viparr tool in Desmond v5.3 software (Schrödinger Release 2020-2: Desmond Molecular Dynamics System, D. E. Shaw Research, New York, NY, 2020. Maestro-Desmond Interoperability Tools, Schrödinger, New York, 2020)^71^ was used to assign amber99sb^72^ force field parameters to protein, CHARMM36 force field to lipids^73–74^, while the Generalized Amber Force Field (GAFF)^75^ was used for the ligand parameterization with the antechamber module^76^ of AMBER18 software^77^; intermolecular interactions were calculated with amber99sb force field^72^.

### MD simulations with Desmond

MD simulations of the three protein-ligand complexes embedded in hydrated POPC bilayers were performed with Desmond v5.3 (Schrödinger Release 2020-2: Desmond Molecular Dynamics System, D. E. Shaw Research, New York, NY, 2020. Maestro-Desmond Interoperability Tools, Schrödinger, New York, 2020)^71^, to investigate the interactions between ligand and hP2X7R, mP2X7R, rP2X7R and the stability of each complex. Systems were equilibrated using a six-step protocol provided in Desmond v5.3. First, a NVT (constant number of particles, volume, temperature) ensemble with Brownian dynamics at 10 K was simulated for 100 ps with small time steps while the solute non-hydrogen atoms were restrained by a force constant of 50 kcal mol^-1^ Å^-2^. Then, the NPT (constant number of particles, pressure, temperature) ensemble was simulated for 12 ps at 10 K, using a Berendsen thermostat^77^ with a fast temperature relaxation constant, velocity re-sampling every 1 ps and keeping the non-hydrogen solute atoms restrained using a force constant of 50 kcal mol Å^-2^. Next, NPT ensemble was simulated for 12 ps at 10 K and 1 atm, using a Berendsen thermostat and a Berendsen barostat with a fast temperature and a slow pressure relaxation constant^78^, re-sampling velocity every 1 ps and keeping non-hydrogen solute atoms restrained by a force constant of 50 kcal mol Å^-2^ restraint. Then another 24 ps simulation of the ensemble was performed using the same protocol as the previous step after raising temperature to 300 K. Final step of the equilibration protocol was 24 ps long simulation of the NPT ensemble.

The particle mesh Ewald method (PME)^79–80^ was employed to calculate the long-range electrostatic interactions with a grid spacing of 0.8 Å. Van der Waals and short-range electrostatic interactions were smoothly truncated at 9.0 Å. The equations of motion were integrated using the multistep reversible reference system propagation *algorithm* (r-RESPA)^81^ integrator with an inner time step of 2 fs for bonded interactions and non-bonded interactions within a cutoff of 12 Å. An outer time step of 6 fs was used for non-bonded interactions beyond the cut-off. The SHAKE method was used to constrain heavy atom-hydrogen bonds at ideal lengths and angles^82^. In the production phase, the relaxed systems were simulated in the NPT ensemble using the Nose-Hoover chain thermostat^83^ and Martyna-Tobias-Klein barostat^84^ at 310 K and 1 atm with a fast temperature relaxation constant and a normal pressure relaxation constant for 500 ns.

We observed that the RMSD of protein’s Cα carbons for hP2X7 and rP2X7 reached a plateau after 50 ns while for mP2X7 after 25ns and thus in the rest of ∼400 ns total simulation time the systems were considered equilibrated and suitable for statistical analysis.

### Analysis of MD simulations trajectories

The visualization of MD simulation trajectories was carried out using the GUI of Maestro (Schrödinger, Schrödinger Release 2020-2) and the protein-ligand interaction analysis was done with the Simulation Interaction Diagram (SID) tool, available with Schrödinger Desmond v.5.3 (Schrödinger Release 2020-2: Desmond Molecular Dynamics System, D. E. Shaw Research, New York, NY, 2020. Maestro-Desmond Interoperability Tools, Schrödinger, New York, 2020). For hydrogen bond interactions, a distance of 2.5 Å between donor and acceptor heavy atoms, and an angle ≥120° between donor-hydrogen-acceptor atoms and ≥ 90° between hydrogen-acceptor-bonded atom were considered. Non-specific hydrophobic contacts were measured if the side chain of a hydrophobic residue fell within 3.6 Å from a ligand’s aromatic or aliphatic carbon, while π-π interactions were measured if two aromatic groups are stacked face-to-face or face-to-edge. Water-mediated interactions were measured if the distance between donor and acceptor atoms is 2.7 Å, the angle between donor-hydrogen-acceptor atoms is ≥ 110° and the angle between hydrogen-acceptor-bonded atom is ≥ 80°. Frames were saved every 10ps. All the MD simulations were run on GTX 4090 GPUs in lab workstations using the GPU implementation of the MD simulations codes. All figures of frames from MD simulations were generated using PyMol (PyMOL Molecular Graphics System, Version 2.0 Schrödinger, LLC).

### Cloning, mutagenesis, and cRNA synthesis

A modified pUC19 vector^85^ with an N-terminally His-tagged rat^85^ and human^86^ P2X7R and the mP2X7R was generated via Gibson assembly according to the protocol of the manufacturer (Gibson Assembly^®^ Master Mix, New England Biolabs, Frankfurt am Main, Germany). F88A and K110A substitutions were introduced into hP2X7R using the Q5^®^ Site-Directed Mutagenesis Kit (New England Biolabs, Ipswich, MA, USA). Capped cRNA was synthesized using the mMESSAGE mMACHINE™ T7 Transcription Kit (ThermoFisher Scientific, Schwerte, Germany) and dissolved in nuclease-free water (500 ng/µl).

### Xenopus laevis oocytes

*Xenopus laevis* frogs were from Nasco International (Fort Atkinson, WI) and kept at the Core Facility Animal Models (CAM) of the Biomedical Center (BMC) of LMU Munich, Germany (Az:4.3.2–5682/LMU/BMC/CAM) in accordance with the EU Animal Welfare Act. Ovaries were surgically extracted (ROB_54-011-AZ 2532.Vet_02-23-166), treated with collagenase (Nordmark, Uetersen, Germany 1.0-1.5 mg/ml, ≥2 at RT) and defolliculated (15-min treatment in Ca^2+^-free oocyte Ringer (90 mM NaCl, 1 mM KCl, 2 mM MgCl_2_, 5 mM HEPES), respectively. 50 nl of cRNA (0.5 µg/µl) were injected using a Nanoject II injector (Science Products, Drummond) and oocytes were kept in ND96 (96 mM NaCl, 2 mM KCl, 1 mM MgCl_2_, 1 mM CaCl_2_, 5 mM HEPES, pH 7.4-7.5) at 16-18°C for 16 h (rP2X7R) or 2 days and then transferred to 4°C to control expression level.

### Two-electrode voltage clamp (TEVC) electrophysiology

Two-electrode voltage clamp recordings were performed at RT and a holding potential of -70 mV using a Turbo Tec-05X amplifier (npi electronic, Tamm, Germany), a magnetic valve system, and CellWorks E 5.5.1 software. Currents were filtered at 1 kHz and digitized at 200 Hz. Electrode resistances were below 1.2 MΩ. A fast and reproducible solution exchange (<300 ms) was achieved with a 50-μl funnel-shaped oocyte chamber combined with a fast vertical solution flow fed through a custom-made manifold mounted immediately above the oocyte.

Recordings were performed in ND96. To estimate association and dissociation rates, a standard concentration of 300 μM ATP in low divalent buffer (no MgCl_2_, 0.5 mM CaCl_2_) was applied for 2 or 5 sec in 2-min intervals. UB-ALT-P2 was dissolved and diluted in DMSO. The final concentration of DMSO in the recording solution was ≤1%. After stabilization of agonist-evoked responses, solutions were switched to antagonist-containing solutions and continuously superfused to determine association rates. Responses following antagonist application were normalized to the preceding, stabilized agonist responses. Dissociation was determined in 1- or 2 min intervals after current responses were inhibited to < 10%. Association curves were fit to the data by the equation % Response = (100-Plateau)*exp(-k_obs_*Time)+Plateau, where Plateau indicates the response at equilibrium and *k_obs_* the experimentally determined on-rate. Dose-inhibition curves for mouse and rat P2X7Rs were generated using the plateau values and IC_50_ values were calculated from a nonlinear fit of the Hill equation to the data : *% Response =Bottom + (Top-Bottom)/(1+10^((LogIC_50_-X)*n_H_*)), where *Top* and *Bottom* are constrained to 100% and 0%, respectively, *X* corresponds to the log of concentration of the antagonist, and *n_H_* corresponds to the Hill coefficient. All data were analyzed using Prism software (Graphpad Software, Inc., Version 8.3.0, San Diego, CA) and are presented as mean ± SD from at least three oocytes.

### Pharmacokinetic study

The pharmacokinetic profile of UB-ALT-P2 was evaluated in male CD1 mice (Envigo Laboratories), aged 8 weeks and weighing 40–50 g. Animals were randomly assigned to either the treatment group (n = 21) or the control group for time zero (t = 0; n = 3). Mice in the treatment group received a single oral dose of UB-ALT-P2 at 3 mg/kg, administered via gavage. The compound was dissolved in 10% (2-hydroxypropyl)-β-cyclodextrin (CAS 128446-35-5; Sigma-Aldrich, Ref. 332607-25G) in physiological saline, with a dosing volume of 10 mL/kg. Animals were monitored for signs of pain or distress throughout the study period. Blood and brain samples were collected at predetermined time points: 0, 0.5, 1, 2, 4, 8, and 10 h post-administration. Blood (0.3–0.5 mL) was obtained via cardiocentesis under anesthesia (90 mg/kg ketamine and 10 mg/kg xylazine) and collected into tubes containing 5% K₂-EDTA. Plasma was separated by centrifugation at 10,000 rpm for 10 minutes at 4 °C and stored at −80 °C until analysis. Brains were carefully extracted immediately following cardiocentesis and stored at −80 °C. All procedures were conducted in accordance with the European Communities Council Directive 86/609/EEC and approved by the Institutional Animal Care and Use Committee of Catalonia (Protocol No. 10291, approved on January 28, 2018).

Determination of UB-ALT-P2 concentrations by LC-MS/MS: An HPLC 1290 Infinity (Agilent, Waldbronn, Germany) with a Synergy Fusion column (Phenomenex), coupled to a QTrap mass 6500+ mass spectrometer (Sciex, Darmstadt, Germany) with an electrospray ionization (ESI) source was employed. Column temperature was 40°C. An HPLC gradient was employed, starting with 90% water containing 0.1% formic acid and 10% acetonitrile. After 30 s the gradient started reaching 100% acetonitrile within 4 min. Then, the column was flushed with acetonitrile for 2.5 min and subsequently equilibrated for 3 min. A sample solution (2 µL) was injected and a flow rate of 0.6 mL/min was applied. Papaverine was used as an internal standard. Calibration samples containing known concentrations of UB-ALT-P2 and the internal standard papaverine were run before, during and after sample measurements. Plasma samples were treated as follows: To 100 µL of plasma, 100 µL of cold acetonitrile containing 0.2% formic acid and 10 nM of papaverine was added. After homogenization and centrifugation (15 min at 15,000 rpm), the supernatant was transferred to HPLC vials and 2 µL were injected. Brain samples were weighed and per mg of brain, 1 mL of acetonitrile containing 0.2% formic acid and 20 nM papaverine was added. The mixture was then treated with a Tissuelyzer (Qiagen, Germany) for 2 min at 50 Hz. Then, the same amount of acetonitrile containing 0.1% of formic acid and 10 nM papaverine was added and the lysing procedure was repeated for 3 min. The samples were subsequently centrifuged at 15,000 rpm for 15 min, and aliquots of the supernatant (100 µL each) were transferred into HPLC vials. Plasma and brain concentrations of UB-ALT-P2 in the samples were subsequently calculated.

### Experimental animals and treatment

Six-month-old male 5xFAD transgenic mice (n = 16) and wild-type (WT) littermates (n = 8) were randomly assigned to two experimental groups: 5xFAD Control (vehicle-treated, n = 8) and 5xFAD + UB-ALT-P2 (n = 8). Mice were housed in groups of four per cage under standardized conditions (temperature: 22 ± 2 °C; light/dark cycle: 12 h/12 h), with ad libitum access to standard chow and water. UB-ALT-P2 was administered via drinking water at a dose of 3 mg/kg, dissolved in 1.8% (2-Hydroxypropyl)-β-cyclodextrin. Drug concentrations were adjusted weekly based on individual cage water consumption. The pharmacological treatment lasted four weeks and continued throughout the behavioral testing period until euthanasia (see Fig. 4B). All procedures were conducted in accordance with the ARRIVE guidelines and the European Communities Council Directive 2010/63/EU, as well as the Guidelines for the Care and Use of Mammals in Neuroscience and Behavioral Research (National Research Council, 2003). The protocol was approved by the Institutional Animal Care and Use Committee and the Generalitat de Catalunya (approval #10291, 15/01/2019). All efforts were made to minimize animal suffering and reduce the number of animals used.

### Novel object recognition test (NORT)

The NORT was used to assess recognition memory based on the natural tendency of rodents to explore novel objects over familiar ones. Experiments were conducted in a 90° two-arm black maze (25-cm-long, 20-cm-high, and 5-cm-wide) of black polyvinyl chloride. The test consisted of three phases: Habituation phase (Days 1–3): Mice were individually placed in the empty maze for 10 min/day. Training phase (Day 4): Two identical plastic objects were placed at the end of each arm. Mice explored the maze for 10 min. Testing phase: Short-term memory trial (2 h post-training): One familiar object was replaced with a novel object; mice explored for 10 min. Long-term memory trial (24 h post-training): A different novel object replaced the familiar one; mice explored for 10 min. Between trials, the maze was cleaned with 70% ethanol to eliminate olfactory cues. All sessions were video-recorded. Exploration was defined as the mouse directing its nose toward the object at ≤2 cm or rearing on it. Turning or sitting near the object was not considered exploratory behaviour. Cognitive performance was quantified using the Discrimination Index (DI), calculated as: DI=TN -TO /TN +TO where (TN) is the time spent exploring the novel object and (TO) is the time spent exploring the familiar object. A positive DI indicates a preference for the novel object, reflecting successful encoding and recall of the familiar object.

### Tissue collection and processing

For Western blot and qPCR analyses, mice were euthanized by cervical dislocation, and brains were rapidly extracted. The cortex and hippocampus were dissected, snap-frozen in powdered dry ice, and stored at −80 °C for protein, RNA, and DNA extraction. For protein isolation, tissues were homogenized in lysis buffer containing phosphatase and protease inhibitors (Cocktail II, Sigma-Aldrich). Protein concentrations were determined using the Bradford assay.

For immunofluorescence, mice were anesthetized with ketamine (100 mg/kg) and xylazine (10 mg/kg) intraperitoneally, followed by transcardial perfusion with 4% paraformaldehyde (PFA) in 0.1 M phosphate buffer. Brains were post-fixed in 4% PFA overnight at 4 °C, then transferred to 15% sucrose in PFA. Brains were frozen in powdered dry ice and stored at −80 °C. Coronal sections (30 μm) were obtained using a cryostat (Leica CM3050S, Wetzlar, Germany) and stored in cryoprotectant at −20 °C.

### Protein levels determination by Western blotting

Protein, histone extraction, and immunoblot analysis were performed as previously reported^87^. Proteins were separated by SDS-PAGE (8-20%) and transferred onto PVDF membranes. Afterward, membranes were blocked in 5% BSA in 0.1% Tween20-TBS (TBS-T) for 1 hour at room temperature (RT), followed by overnight incubation at 4 °C with the primary antibodies presented in Supplementary Table 6. Immunoreactive proteins were viewed with a chemiluminescence-based detection kit, following the manufacturer’s protocol and digital images were acquired using a ChemiDoc XRS+System. Semi-quantitative analyses were carried out using ImageLab Software and results were expressed in Arbitrary Units (AU), considering the control mice group as 100%. Protein loading controls were routinely monitored by immunodetection of GAPDH.

### RNA extraction and gene expression analysis

Total RNA was extracted from hippocampal tissue using TRIsure™ reagent (Bioline), following the manufacturer’s instructions. RNA concentration and purity were assessed using a NanoDrop™ ND-1000 spectrophotometer (Thermo Scientific), and integrity was confirmed with an Agilent 2100 Bioanalyzer (Agilent Technologies). Only samples with 260/280 ratios >1.9 and RNA integrity numbers (RIN) >7.5 were used. Reverse transcription was performed using 2 μg of mRNA and the High-Capacity cDNA Reverse Transcription Kit (Applied Biosystems). Quantitative real-time PCR (qPCR) was conducted using SYBR^®^ Green PCR Master Mix (Applied Biosystems) on a StepOnePlus™ Real-Time PCR System. Each 15 μL reaction contained 6.75 μL of cDNA (2 μg), 0.75 μL of each primer (100 nM), and 6.75 μL of SYBR® Green Master Mix. Gene expression of oxidative stress and synaptic plasticity markers (see Supplementary Table 7) was analyzed using the ΔΔCt method, with β-actin as the reference gene. All reactions were performed in duplicate.

### Amyloid-β plaque staining

Amyloid-β plaques were visualized using thioflavin S staining. Coronal brain sections (30 μm) were washed in 0.01 M PBS for 5 min, followed by sequential 1-min washes in 70% and 80% ethanol. Sections were incubated in 0.3% thioflavin S (Sigma-Aldrich) for 15 min in the dark at room temperature, then washed in 80%, 70%, and 50% ethanol (1 min each), followed by three 2-min washes in 0.1 M PBS. Sections were mounted using Fluoromount-G (EMS).

### Image acquisition and analysis

Fluorescence images were acquired using an Olympus BX51 microscope (Hamburg, Germany) with 4×, 10×, and 20× objectives. ImageJ software (RRID: SCR_003070) was used for analysis. For plaque quantification, comparable cortical and hippocampal regions were selected. Images were converted to 8-bit grayscale, adjusted to the appropriate threshold, and analyzed using the “Analyze Particles” function (size: 10–Infinity). The percentage of area covered by thioflavin S (20X objective) was calculated and averaged from two different sections from each animal.

### Statistical analysis

Statistical analyses were performed using GraphPad Prism (version 9.5.1). Data are presented as mean ± standard error of the mean (SEM). For Western blot, n = 4; for qPCR, n = 3. Comparisons between two groups were made using two-tailed Student’s t-tests. For comparisons for more than 2 groups, one-way ANOVA followed by Tukey’s post hoc test. Statistical significance was set at *P* < 0.05. Outliers were identified and excluded using Grubbs’ test.

## Data Availability

All cryo-EM density maps for full-length wild-type hP2X7R, mP2X7R, and rP2X7R in antagonist-bound inhibited states have been deposited in the Electron Microscopy Data Bank (EMDB) under accession codes: EMD-74135 (UB-MBX-47-bound inhibited hP2X7R), EMD-74139 (UB-ALT-P1-bound inhibited hP2X7R), EMD-74136 (UB-ALT-P2-bound inhibited hP2X7R), EMD-74138 (UB-ALT-P2-bound inhibited mP2X7R), and EMD-74134 (UB-ALT-P2-bound inhibited rP2X7R). The maps within these depositions include both half maps, sharpened/unsharpened maps, refinement masks, and any local refinements or locally sharpened maps that helped with model building. The corresponding coordinates for the structures have been deposited in Protein Data Bank under the PDB accession codes: 9ZF7 (UB-MBX-47-bound inhibited hP2X7R), 9ZFB (UB-ALT-P1-bound inhibited hP2X7R), 9ZF8 (UB-ALT-P2-bound inhibited hP2X7R), 9ZFA (UB-ALT-P2-bound inhibited mP2X7R), and 9ZF6 (UB-ALT-P2-bound inhibited rP2X7R). First and last snapshots from MD simulations of complexes between full-length WT hP2X7R, mP2X7R, rP2X7R and UB-ALT-P2, as well as WT hP2X7R and UB-ALT-P1 or UB-MBX-46 or UB-MBX-47 can be found in the following GitHub repository: https://github.com/georgioukyriakos/P2X7/tree/main/brain_penetrant_P2X7R_antago <u>nist</u>. Additional PDB files were used for analysis including 9E3M (apo closed hP2X7R), 9E3Q (apo closed mP2X7R), 8TR5 (apo closed rP2X7R), 8TR8 (AZD9056-bound inhibited rP2X7R), 9E3O (UB-ALT-P30-bound inhibited hP2X7R), and 9E3P (UB-MBX-46-bound inhibited hP2X7R)^24,34,41^.

## Acknowledgements

We thank O. Davulcu, C. Yoshioka, and C. López at PNCC for access and microscopy assistance. Electron microscopy grid screening was performed at the Multiscale Microscopy Core within Oregon Health & Science University (OHSU). The authors would like to acknowledge the contributions of the OHSU Biophysics Shared Resource Core (Research Resource ID: RRID: SCR_022744) in facilitating this work. We thank L. Anson for comments on the manuscript. A portion of this research was supported by NIH grant U24GM129547 and performed at the PNCC at OHSU and accessed through EMSL (grid.436923.9), a DOE Office of Science User Facility sponsored by the Office of Biological and Environmental Research. This research was supported by the National Heart, Lung and Blood Institute (R00HL138129, S.E.M.), the National Institute of General Medical Sciences (DP2GM149551, S.E.M.), and the American Heart Association (24PRE1195450, A.C.O.). Part of this work was funded by the Spanish *Ministerio de Ciencia, Innovación y Universidades*, MICIU/AEI/10.13039/501100011033: grants PID2023-147004OB-I00 (to S.V.) and PID2023-148642OB-I00 (to J.L.). A.K. thanks Chiesi Hellas for funding with computer resources. C.E.M. and A.N. were supported by grants from the Deutsche Forschungsgemeinschaft (DFG, German Research Association) Project-ID 335447717 - SFB 1328 (A11 and A15). The Mansoor Lab would like to thank the Silver family, Steve Janik and Sheryl Manning, Barbara Allen and Jim Batzer, and Randy and Barbara Lovre for their generous support. The Müller Lab would like to thank Christiane Bous (laboratory technician). A.L.T. and S.V. thank Laura Castro (laboratory technician).

## Author contributions

The manuscript was written with input from all authors. A.L.T., A.C.O., S.E.M. and S.V. designed the project. A.L.T., C.B., M.B.-X. and S.V. designed, synthesized, purified and chemically characterized the P2X7R antagonists. A.C.O. performed the cryo-EM sample preparation, data collection, data processing, and built the models. C.G.-F. and M.P. performed *in vivo* assays. A.D., J.S.-M. and A.N. performed and analyzed the TEVC electrophysiological experiments. S.H., J.N. and C.E.M. prepared stable cell lines expressing human P2X1-4Rs, and cell lines expressing human, rat and mouse P2X7Rs, and performed the calcium influx assays, and analyzed the data. M.B., Z.B. and J.L. performed the synthesis of the radiolabeled compound and PET studies. G.K., S.L., and Y.K. performed and analyzed the ethidium bromide accumulation experiments. M.S. and C.E.M. analyzed the pharmacokinetics samples. C.V., J.B. and M.I.L. performed in vitro DMPK profiling, including cytochrome P450 assays, hERG assessment and Caco-2 permeability experiments. B.P. performed the PAMPA-BBB assay. L.N. performed the cytotoxicity assays. K.G., E.T. and A.K. performed and analyzed the MD simulations. A.L.T., A.C.O., S.E.M and S.V. wrote the manuscript with feedback from A.N, C.E.M, A.K. and M.P. All authors approved the final version of the manuscript.

## Additional information

**Supplementary information** is available for this paper at

**Correspondence and requests for materials should be addressed** to Steven E. Mansoor and Santiago Vázquez.

---

**Extended Data Fig. 1:**
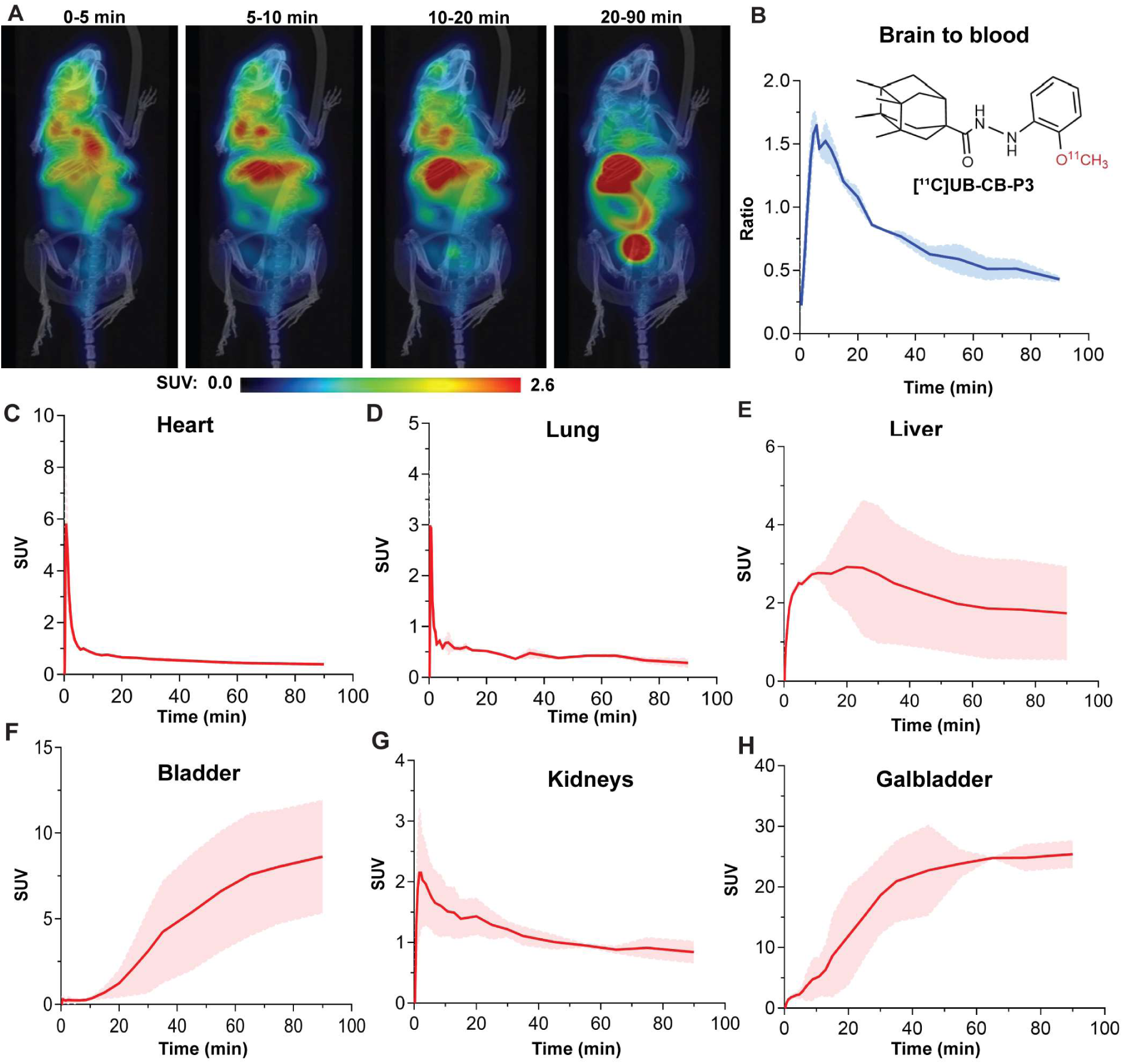
Dynamic PET/CT biodistribution of [¹¹C]UB-CB-P3 in healthy mice. (**A**) Representative maximum intensity projections (MIPs) of PET images co-registered with 3D-rendered CT images from the same animal at sequential post-injection windows (0–5 min, 5–10 min, 10–20 min, and 20–90 min). Frames within each window were averaged to generate the representative images. Early images show tracer uptake in the brain and other highly perfused organs, whereas later frames reveal progressive accumulation in the gallbladder and urinary bladder. The colour scale denotes standardized uptake values (SUVs). (**B**) Brain-to-blood ratio over time for [¹¹C]UB-CB-P3, shown together with the chemical structure of the radiotracer. Quantification during the pseudo-equilibrium interval (15–60 min) yielded a K_p,brain_ of ∼0.45, indicating that total brain concentrations remain lower than blood concentrations over this period. Total K_p,brain_ does not directly represent the pharmacologically relevant exposure. A more mechanistic assessment of BBB transport requires the unbound brain-to-plasma ratio (K_p,uu,brain_), which integrates total concentrations with the unbound fractions in plasma and brain tissue. Determining these fractions ex vivo, such as by equilibrium dialysis in plasma and brain homogenates or by brain-slice assays, would allow distinguishing between permeability-driven effects and tissue-binding artifacts. Although such neuroPK experiments were beyond the scope of the present imaging study, our PET data provide clear proof-of-principle that the P2X7R antagonist scaffold reaches the CNS *in vivo*. (**C-H**) Dynamic time–activity curves (SUV vs time) for heart (**C**), lungs (**D**), liver (**E**), urinary bladder (**F**), kidneys (**G**) and gallbladder (**H**) after intravenous administration of [¹¹C]UB-CB-P3. Peripheral uptake is evident in highly perfused organs, with increasing activity in the urinary bladder indicating renal clearance. A marked accumulation in the gallbladder suggests additional hepatobiliary contribution to tracer elimination.

**Extended data Fig. 2:**
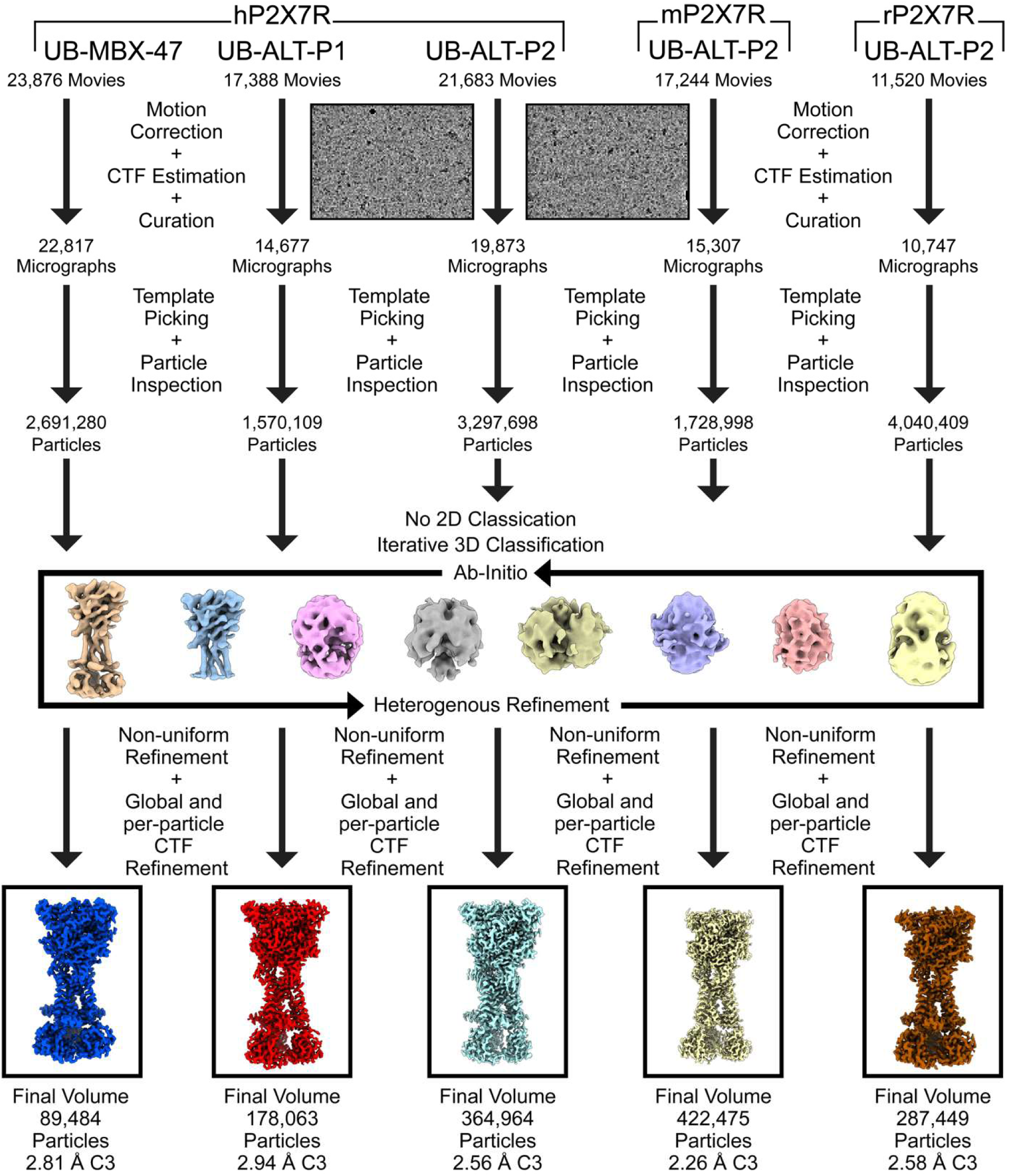
Cryo-EM processing pipeline for P2X7R reconstructions. Following large-scale data acquisition at the Pacific Northwest Center for Cryo-EM (PNCC), movies were motion corrected at the physical pixel size, and patch-based contrast transfer function (CTF) correction was performed in cryoSPARC^57^. Next, micrographs were inspected, and particle picking was performed using 2D templates generated from a low-resolution 3D reconstruction. Poor initial particle picks were removed, and the remaining picks were extracted and sent directly to iterative 3D classification in cryoSPARC using ab initio and heterogeneous refinement jobs. After sufficient classification, a final particle stack was re-extracted at the physical pixel size for non-uniform refinement performed with 3-fold (C3) symmetry as well as global and per-particle CTF refinements.

**Extended data Fig. 3:**
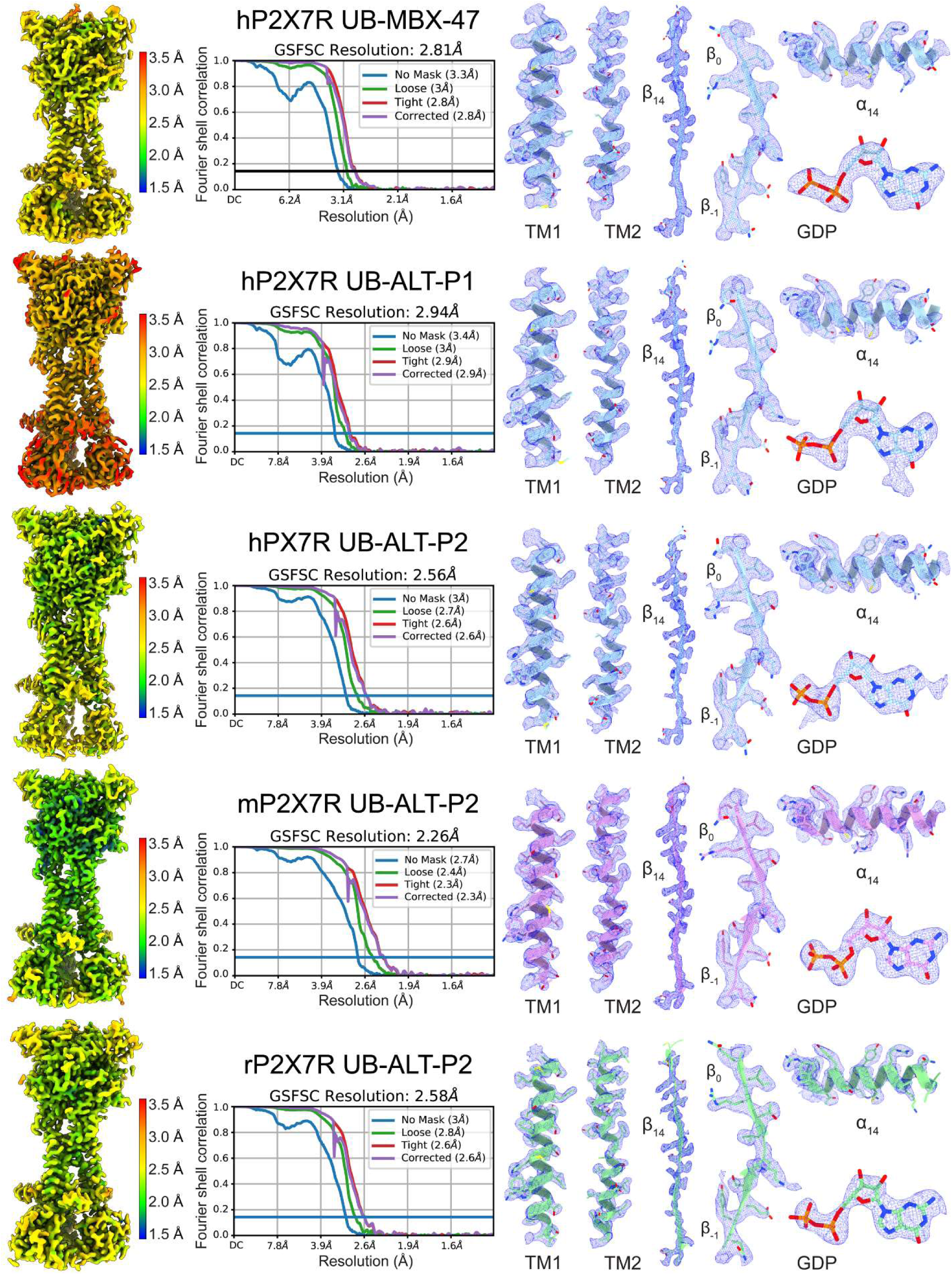
Fourier shell correlation (FSC), local resolution plots, and representative cryo-EM density for P2X7R reconstructions. The resolution stated is at an FSC = 0.143. All local resolution plots range between 1.5 Å (blue) and 3.5 Å (red). Representative regions (TM1, TM2, β_-1_, β_0_, β_14_, α_14_, and GDP) from the structures of the UB-MBX-47-bound inhibited hP2X7R, the UB-ALT-P1-bound inhibited hP2X7R, the UB-ALT-P2-bound inhibited hP2X7R, the UB-ALT-P2-bound inhibited mP2X7R, and the UB-ALT-P2-bound inhibited rP2X7R are shown within their respective cryo-EM densities (blue mesh) highlighting the strong map-to-model fits. Protein models are colored according to species: human (blue), mouse (purple), and rat (green).

**Extended data Fig. 4:**
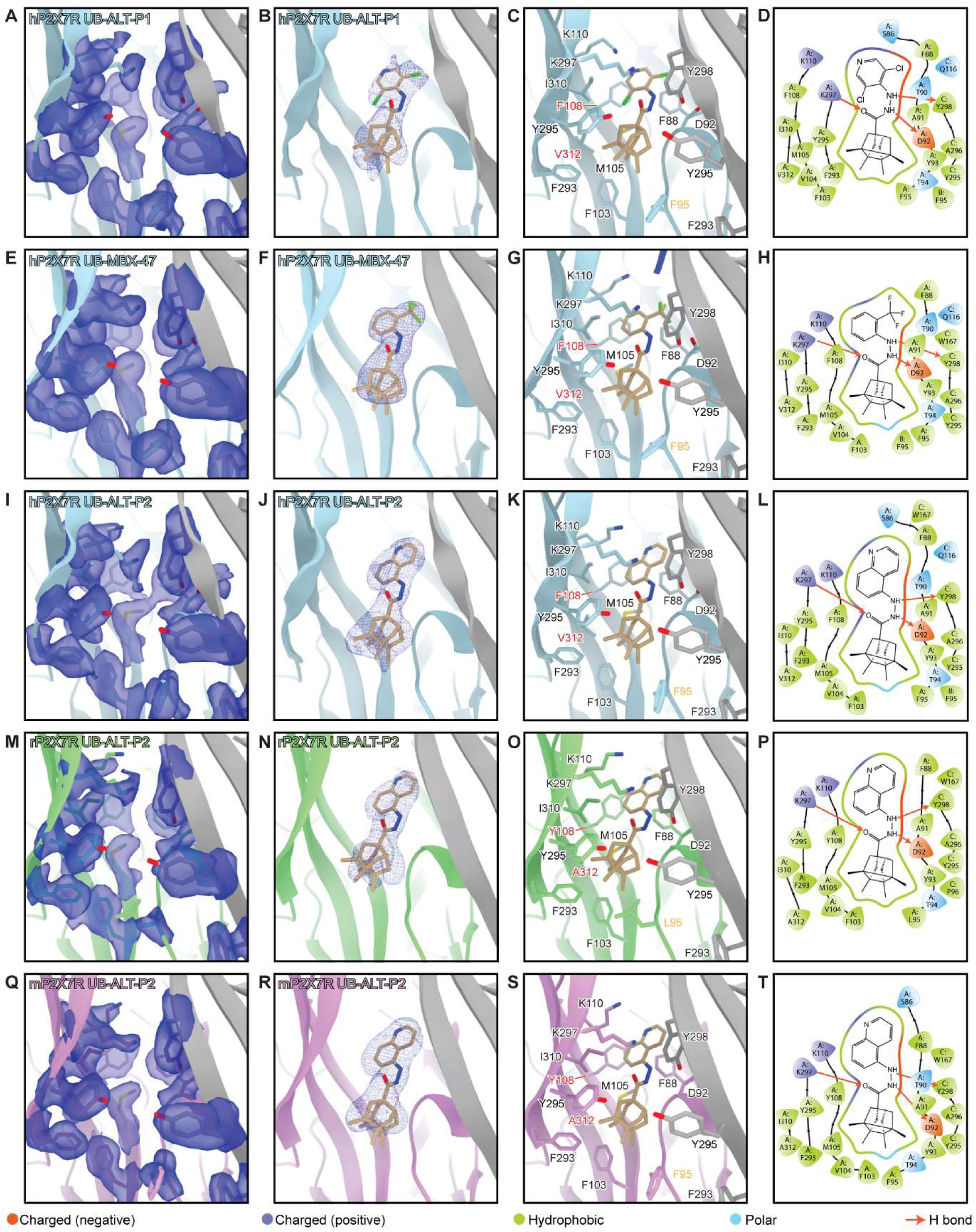
Cryo-EM analysis reveals structural features of the classical allosteric pocket in the P2X7R across five distinct antagonist-bound inhibited states. Ribbon representation of the classical allosteric ligand-binding site in human (light blue and grey), rat (light green and grey), and mouse (light purple and grey) P2X7Rs, located at the interface of two protomers, labeled chain A (in color) and chain C (in grey). (**A, E, I, M, Q**) View of the classical allosteric ligand-binding site, with the ligand hidden, showing the cryo-EM density for the side chains of residues within pocket for each of the five antagonist-bound structures. The cryo-EM density is shown in transparent blue, highlighting the high quality and resolution of each cryo-EM reconstruction. (**B, F, J, N, R)** Same view of the classical allosteric pocket as (A, E, I, M, Q) with one molecule of UB-ALT-P1 bound to the hP2X7R (**B**, tan), UB-MBX-47 bound to the hP2X7R (**F**, tan), UB-ALT-P2 bound to the hP2X7R, (**J**, tan), UB-ALT-P2 bound to the rP2X7R, (**N**, tan), and UB-ALT-P2 bound to the mP2X7R, (**R**, tan) shown with their corresponding cryo-EM density (blue mesh) at 2.9 Å, 2.8 Å, 2.5 Å, 2.2 Å, and 2.6 Å, respectively. (**C, G, K, O, S**) Same view of the classical allosteric pocket as (A, E, I, M, Q) showing residues that interact with UB-ALT-P1 in the hP2X7R (**C**, tan), UB-MBX-47 in the hP2X7R (**G**, tan), UB-ALT-P2 in the hP2X7R, (**K**, tan), UB-ALT-P2 in the rP2X7R, (**O**, tan), and UB-ALT-P2 in the mP2X7R (**S**, tan). The views highlight the differences between ligand-receptor interactions as well as the interactions of UB-ALT-P2 across orthologs. (**D, H, L, P, T**) Schematic representation of ligand-receptor interactions between UB-ALT-P1 and the hP2X7R (**D**), UB-MBX-47 and the hP2X7R (**H**), UB-ALT-P2 and the hP2X7R, (**L**), UB-ALT-P2 and the rP2X7R, (**P**), and UB-ALT-P2 and the mP2X7R (**T**). Protomer chain IDs (A, B and C) are assigned in a counterclockwise orientation when viewed from the extracellular domain. Hydrogen bonds shown as red arrows and residue interactions in green (hydrophobic), blue (positively charged), red (negatively charged), and light blue (polar). Interaction plots generated using Maestro^36^.

**Extended data Fig. 5:**
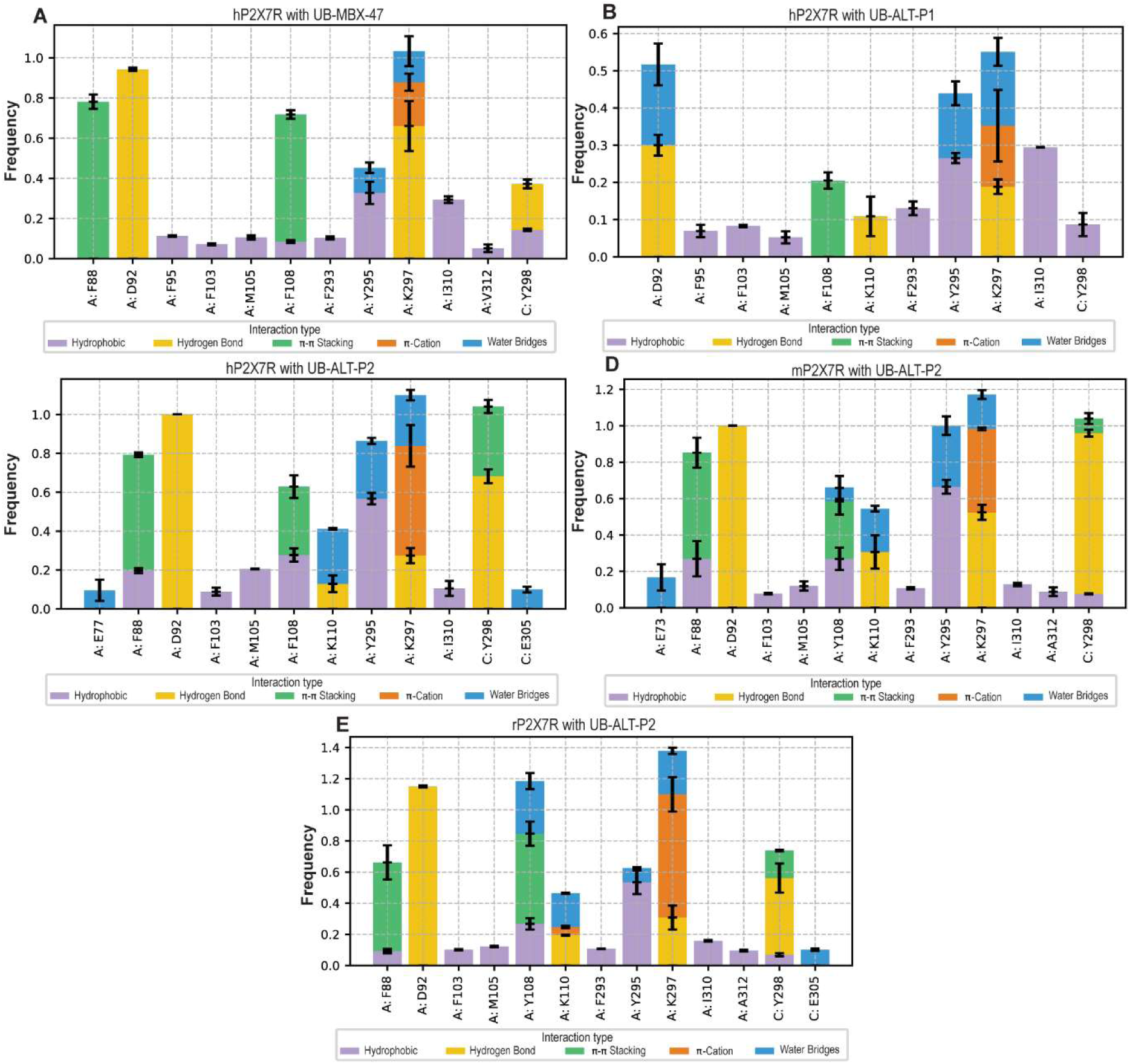
Interaction fingerprints from MD simulations of P2X7R–ligand complexes. Ligand–residue interaction frequency histograms derived from 500-ns molecular dynamics simulations of full-length P2X7Rs in complex with UB-MBX-47 bound to hP2X7R (**A**), UB-ALT-P1 bound to hP2X7R (**B**), and UB-ALT-P2 bound to human (**C**), mouse (**D**), and rat (**E**) P2X7Rs. Protomer chain IDs (A, B and C) are assigned in a counterclockwise orientation when viewed from the extracellular domain (top-down view). Bars represent the frequency of each interaction, classified by type: hydrophobic (purple), hydrogen bond (yellow), π–π stacking (green), cation–π (orange), and water bridges (blue. Error bars indicate the standard deviation across two independent simulation replicas.

**Extended data Fig. 6:**
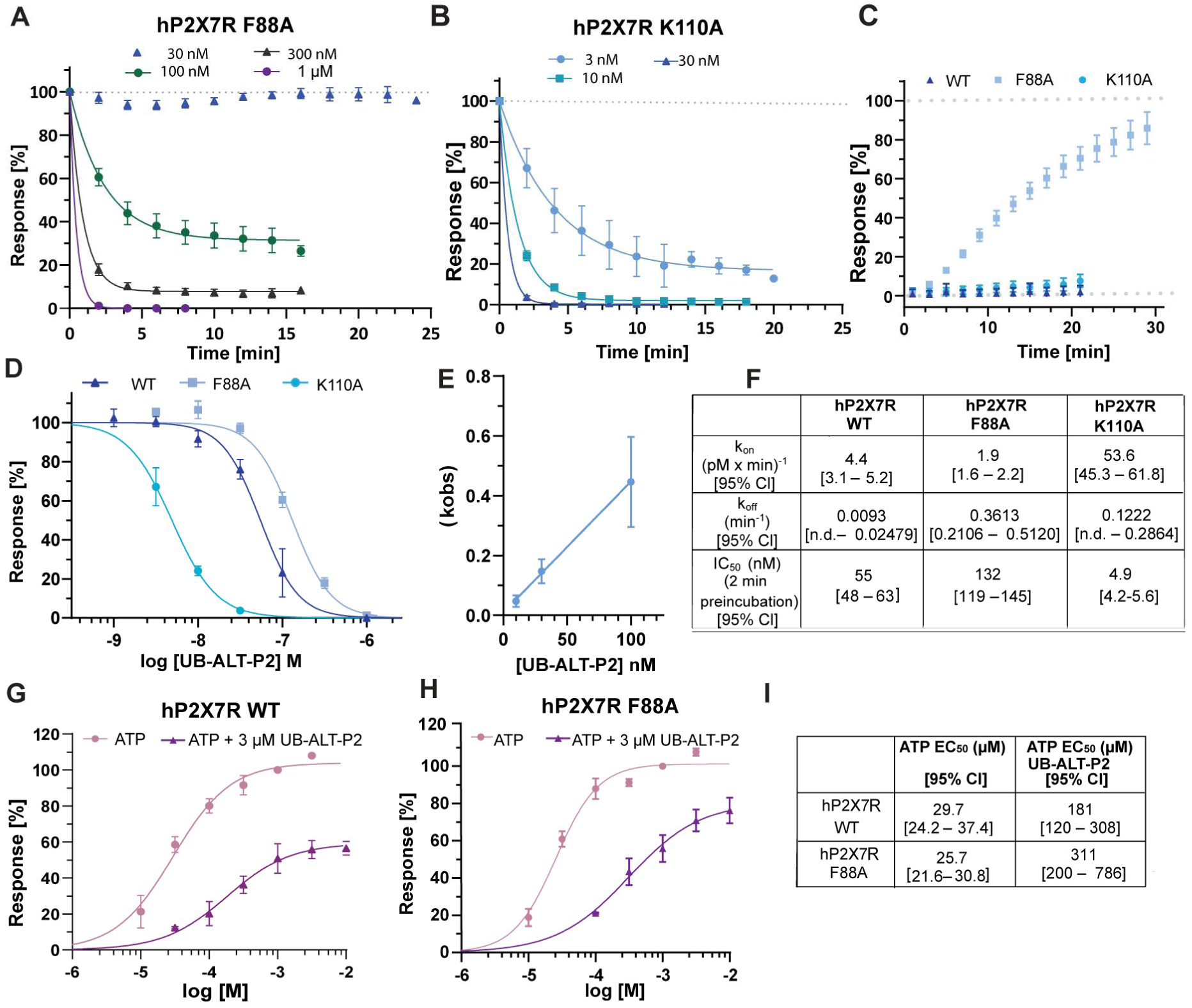
Functional and kinetic characterization of UB-ALT-P2 at hP2X7R and allosteric pocket mutants. (**A–B**) Time course of UB-ALT-P2 binding to hP2X7R F88A (**A**) and hP2X7R K110A (**B**). The F88A mutation has previously been shown to reduce the potency of P2X7R-specific inhibitors (A740003, A804598, AZ10606120, JNJ47965567) in a YO-PRO-1 uptake assay in HEK293 cells^40^. P2X7Rs were expressed in *Xenopus laevis* oocytes and analyzed by TEVC at −70 mV. A standard concentration of 300 µM ATP was applied for 2 sec in 2-min intervals, followed by a 1-min perfusion with buffer. After stabilization of agonist-evoked responses, solutions were switched to antagonist-containing solutions and continuously superfused with the indicated concentration of antagonist. Responses following the switch to antagonist-containing solutions were normalized to the preceding, stabilized agonist responses (represented as current responses at 0 min). (**C**) Time course of antagonist dissociation from WT and mutated hP2X7Rs. After stabilization of agonist-evoked responses, the antagonist was applied at a concentration that yielded about 90% inhibition (WT: 10 µM, F88A: 100 µM, K110A: 1 µM) and incubated for 1 min. The 90-100% block by the antagonist is indicated at 0 min and the following normalized responses to 2-sec pulses of ATP (300 µM) in 2-min intervals are shown. (**D**) Concentration–inhibition curves of UB-ALT-P2 on WT and mutated hP2X7R derived from responses obtained after a 2-min pre-incubation time of the antagonist. For IC_50_ values see **F**. As current responses were determined before antagonist binding was completed (compare A-B and Fig. 3D), the potency of UB-ALT-P2 in these measurements is largely underestimated, in particular for the WT hP2X7R. (**E**) Observed on-rates k_obs_ were determined for the WT hP2X7R by fitting the data to the function: % response = (100-Plateau)*exp(-k_obs_*time)+Plateau. The k_obs_ were then plotted against the respective antagonist concentration (F). Assuming the simplest case of receptor-ligand kinetics, a 1:1 binding model, the formulas k_obs_=k_on_*F+k_off_ and K_i_=k_off_/k_on_ were used to obtain an estimate for the theoretical off-rate constant k_off_ (y-intercept) and the corresponding K_i_ value, which is 2.1 nM (see Fig. 3I). It has to be noted, however, that the graphically determined k_off_ (0.009 min^-1^) does not reflect the very slow dissociation from the human P2X7R and this K_i_ value is therefore largely overestimated. (**F**) Rate constants and IC_50_ values of UB-ALT-P2 in WT and mutated receptors. The F88A mutation markedly increased the dissociation rate of UB-ALT-P2 and reduced its potency, indicating that F88 stabilizes high-affinity ligand binding. In contrast, the K110A mutation strongly increased the association rate while preserving slow dissociation, suggesting that K110 restricts ligand access to the binding site and primarily modulates the rate of antagonist association. (**G-H**) For binding mode analysis, the agonist dose-response relationship for ATP on WT hP2X7R (**G**) and the F88A mutant (**H**) were compared in the presence and absence of 3 µM UB-ALT-P2. (**I**) EC_50_ values for ATP with or without UB-ALT-P2 on hP2X7R and hP2X7R-F88A. Numbers in brackets are 95% confidence intervals in panel H and I. (**A-I**) If not otherwise noted, data are represented as mean ± SD from 3-9 measurements.

**Extended Data Fig. 7:**
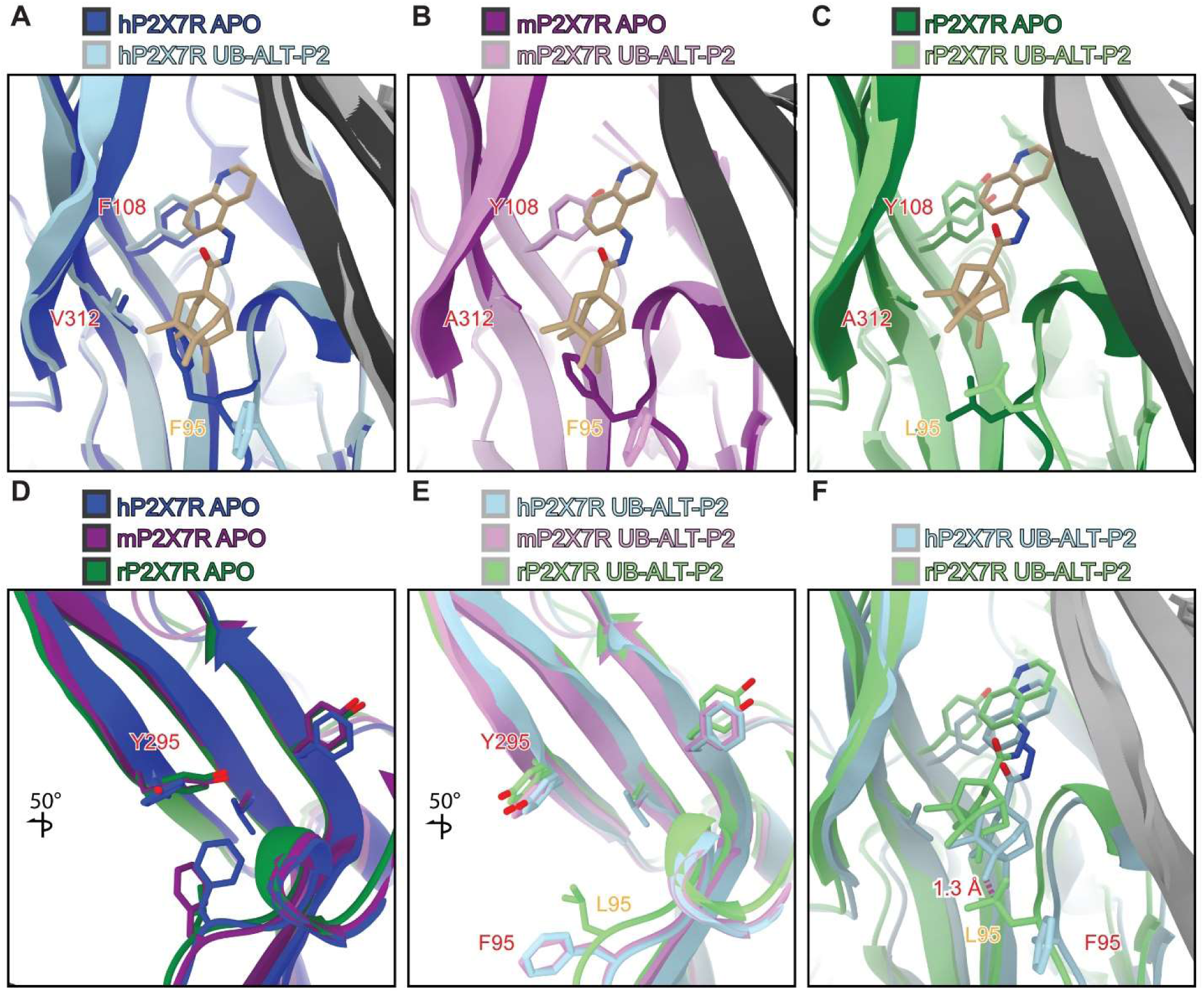
Species-specific structural changes in the classical allosteric pocket of P2X7 receptors upon UB-ALT-P2 binding. (**A-C**) Comparison of the apo closed (dark blue, dark purple, and dark green) and UB-ALT-P2 inhibited (light blue, light purple, and light green) state structures of the hP2X7R (**A**), the mP2X7R (**B**), and the rP2X7R (**C**), respectively, highlighting ligand-induced movements of subtype-specific residues within the classical allosteric pocket (PDB codes: 9E3M, 9E3Q, and 8TR5)^24,41^..Across all three orthologs, residues at positions 108 and 312 do not change conformations between the apo closed and UB-ALT-P2 inhibited states. In contrast, F95 in the hP2X7R and the mP2X7R are dramatically displaced upon ligand binding while L95 in the rP2X7R stays in a similar conformation. (**D**) Comparisons between the apo closed state structures of the hP2X7R (dark blue and dark grey, PDB code: 9E3M), the mP2X7R (dark purple and dark grey, PDB code: 9E3Q), and the rP2X7R (dark green and dark grey, PDB code: 8TR5), highlighting the position of subtype-specific residues^24,41^. In the apo closed state, residues at position 95, 108, and 312 all occupy similar rotamers between orthologs while the residues at position 295 occupy different rotameric conformations^24^. (**E**) Comparisons between the UB-ALT-P2-bound inhibited state structures of the hP2X7R (light blue and grey), the mP2X7R (light purple and grey), and the rP2X7R (light green and grey) with the ligand hidden, highlighting the position of subtype-specific residues. In the UB-ALT-P2-bound inhibited state, residues at position 108, 295, and 312 all occupy similar rotamers between orthologs while the residues at position 95 occupy different conformations. (**F**) Comparison between the UB-ALT-P2-bound inhibited state structures of the hP2X7R (light blue and grey) and the rP2X7R (light green and grey) highlighting the subtype-specific residue that alters the ligand pose. L95 in the rP2X7R points into the classical allosteric pocket while F95 in the hP2X7R points away from the pocket. The shortest distance between L95 in the rP2X7R and UB-ALT-P2 in the hP2X7R is only 1.3 Å, which would create steric clash. Thus, the ligand UB-ALT-P2 adopts a distinct binding pose in rP2X7R compared with its binding mode in hP2X7R and mP2X7R.

**Extended data Fig. 8:**
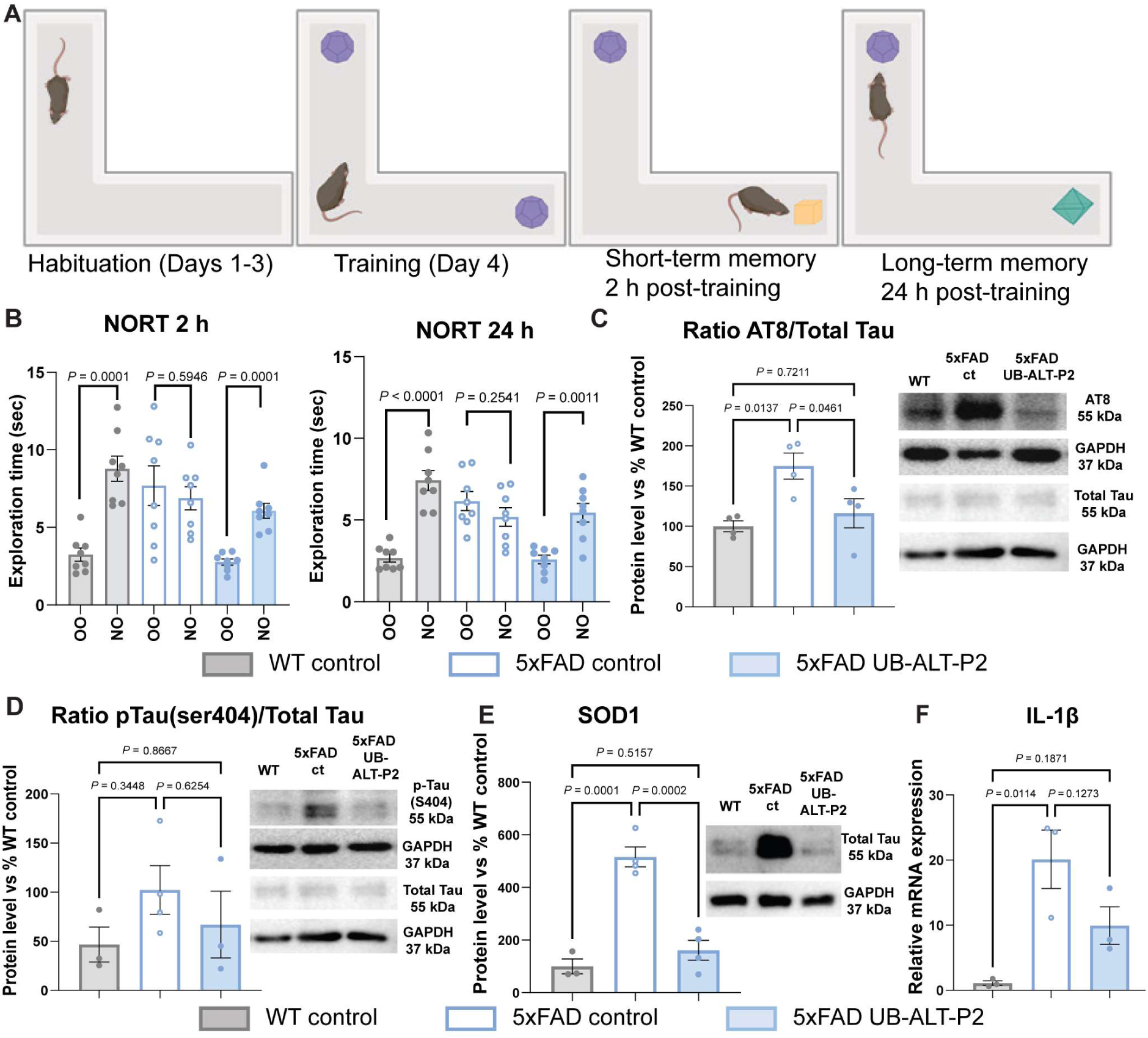
NORT performance and Western blot analysis of AT8/Total Tau, pTau(S404)/Total Tau, SOD1 and qPCR analysis of IL-1β in 5xFAD mice treated with UB-ALT-P2. (**A**) Schematic of the novel object recognition test (NORT). Mice underwent habituation (Days 1–3) and training (Day 4), followed by assessment of short-term memory (2 h post-training) and long-term memory (24 h post-training). (**B**) Exploration times for the familiar (OO) and novel object (NO) during the short-term (left) and long-term (right) memory trials. UB-ALT-P2–treated 5xFAD mice showed restored preference for the novel object compared to untreated 5xFAD controls. (**C-D**) Quantification and representative Western blots for tau pathology. UB-ALT-P2 treatment partially normalised the ratio of the protein levels AT8/Total Tau (**C**) and p-Tau(S404)/Total Tau (**D**) ratio. Tau pathology quantified as the ratio of phosphorylated (AT8 or S404) to total tau and values were expressed relative to corresponding ratio in WT mice. Immunoblots for AT8, p-Tau(S404), Total Tau and GAPDH (loading control) are shown alongside each graph. (**E**) Quantification and representative Western blots of the oxidative stress marker SOD1. 5xFAD controls showed robust induction of SOD1 protein levels, whereas UB-ALT-P2 treatment reduced SOD1 expression toward WT levels. (**F**) Relative hippocampal IL-1β mRNA expression in WT controls, 5xFAD controls and 5xFAD mice treated with UB-ALT-P2, showing the expected elevation in the 5xFAD group and a trend toward reduction following treatment with UB-ALT-P2. Across all panels, bars represent mean ± SEM; individual points indicate biological replicates. Statistical significance was determined by ANOVA followed by Tukey’s post hoc test (**C, D, E, F**) and two-tailed Student’s t-tests (**B**). Exact *P* values are reported in the figures.

**Extended Data Table 1:**
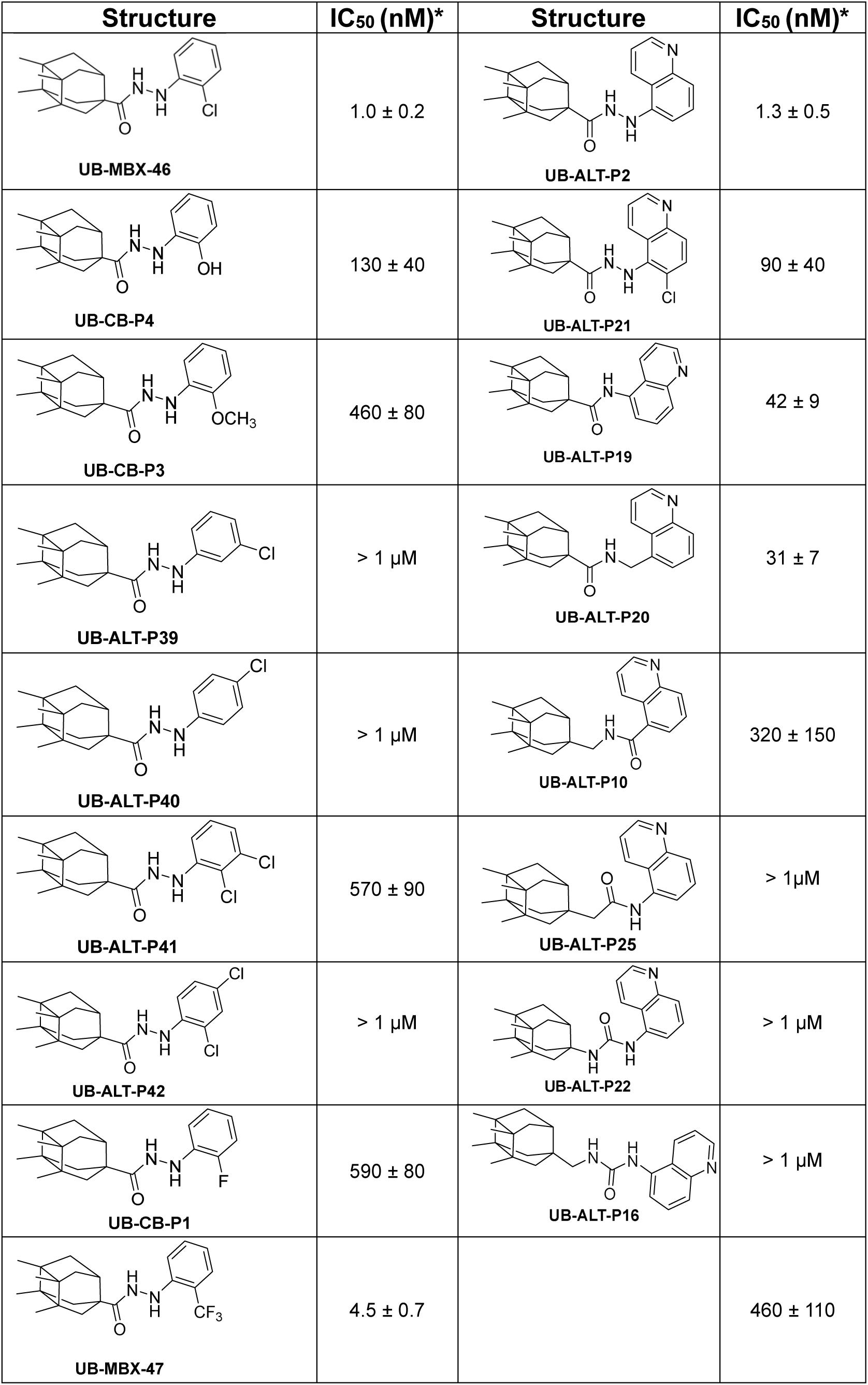
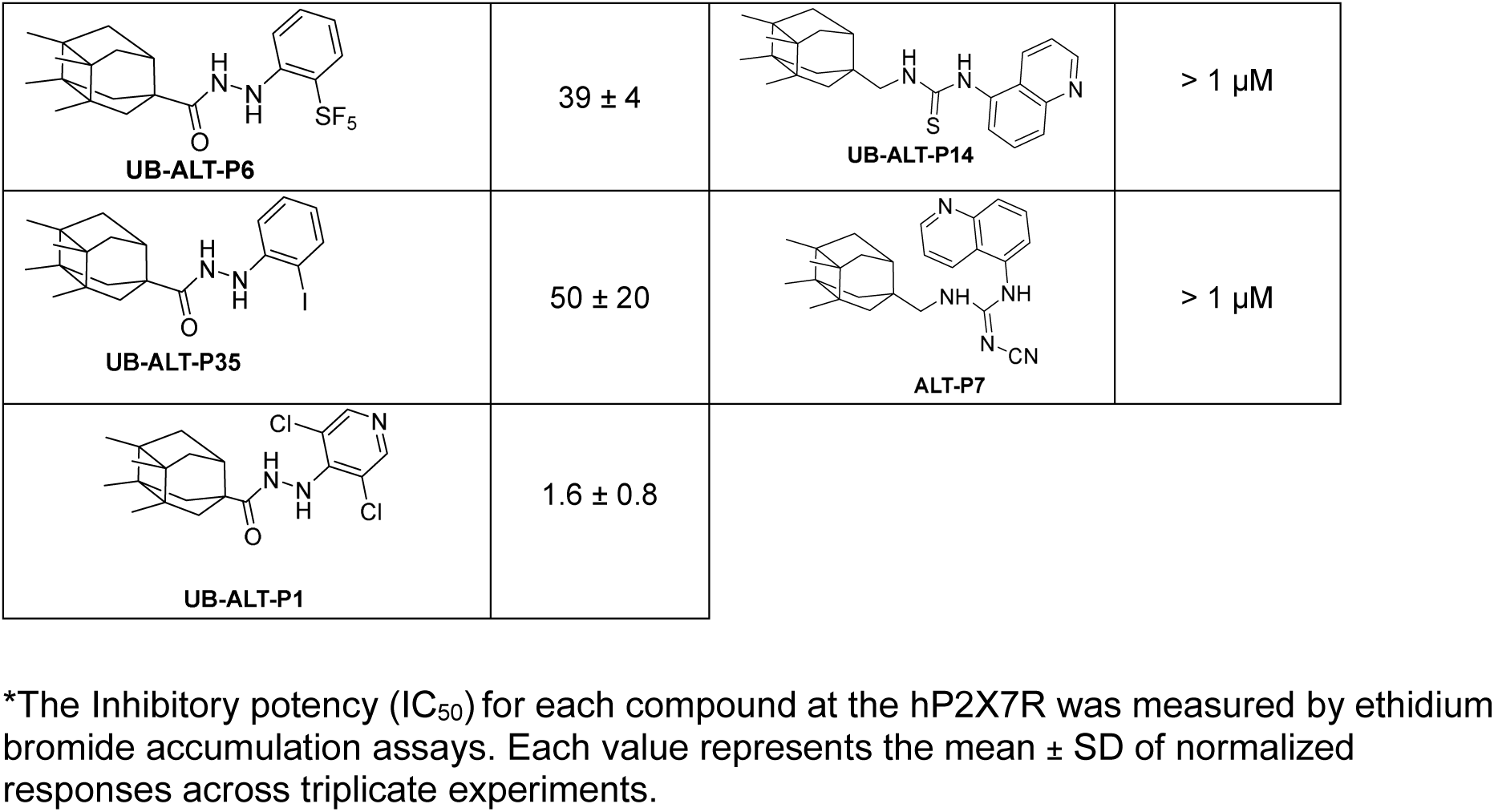
Structure and inhibitory potencies (IC_50_) of antagonists at the hP2X7R.

| Structure | IC<sub>50</sub> (nM)* | Structure | IC<sub>50</sub> (nM)* |
| --- | --- | --- | --- |
| UB-MBX-46 | 1.0 ± 0.2 | UB-ALT-P2 | 1.3 ± 0.5 |
| UB-CB-P4 | 130 ± 40 | UB-ALT-P21 | 90 ± 40 |
| UB-CB-P3 | 460 ± 80 | UB-ALT-P19 | 42 ± 9 |
| UB-ALT-P39 | > 1 μM | UB-ALT-P20 | 31 ± 7 |
| UB-ALT-P40 | > 1 μM | UB-ALT-P10 | 320 ± 150 |
| UB-ALT-P41 | 570 ± 90 | UB-ALT-P25 | > 1 μM |
| UB-ALT-P42 | > 1 μM | UB-ALT-P22 | > 1 μM |
| UB-CB-P1 | 590 ± 80 | UB-ALT-P16 | > 1 μM |
| UB-MBX-47 | 4.5 ± 0.7 |  | 460 ± 110 |
| UB-ALT-P6 | $39 \pm 4$ | UB-ALT-P14 | $> 1 \mu\text{M}$ |
| UB-ALT-P35 | $50 \pm 20$ | ALT-P7 | $> 1 \mu\text{M}$ |
| UB-ALT-P1 | $1.6 \pm 0.8$ | | |

\*The Inhibitory potency ( $\text{IC}_{50}$ ) for each compound at the hP2X7R was measured by ethidium bromide accumulation assays. Each value represents the mean ± SD of normalized responses across triplicate experiments.

