## Supporting information for "A brain-penetrant P2X7R antagonist mitigates Alzheimer’s disease pathology"

a. Laboratori de Química Farmacèutica, Facultat de Farmàcia i Ciències de l'Alimentació, Universitat de Barcelona, Barcelona, Spain
b. Institute of Biomedicine of the University of Barcelona, IBUB, Barcelona, Spain.
c. Department of Chemical Physiology & Biochemistry, Oregon Health & Science University, Portland, OR, USA.
d. Departament de Farmacologia, Toxicologia i Química Terapèutica, Institut de Neurociències-Universitat de Barcelona, Barcelona, Spain.
e. Centro de Investigación en Red, Enfermedades Neurodegenerativas (CIBERNED), Instituto de Salud Carlos III, Madrid, Spain.
f. Walther Straub Institute of Pharmacology and Toxicology, Faculty of Medicine, Ludwig-Maximilians-Universität München, Munich, Germany.
g. PharmaCenter Bonn & Pharmaceutical Institute, Pharmaceutical & Medicinal Chemistry, University of Bonn, Bonn, Germany.
h. Present address: Institute of Pharmaceutical and Medicinal Chemistry, University of Düsseldorf, Düsseldorf, Germany.
i. Present address: Department of Anesthesiology, Washington University Pain Center, St. Louis, MO, USA.
j. School of Life Sciences, Gwangju Institute of Science and Technology, 123 Cheomdangwagi-ro, Buk-gu, Gwangju, Republic of Korea.

k. Laboratory of Medicinal Chemistry, Section of Pharmaceutical Chemistry, Department of Pharmacy, National and Kapodistrian University of Athens, Panepistimiopolis-Zografou, Greece.
l. CIC biomaGUNE, Basque Research and Technology Alliance (BRTA), San Sebastián, Guipúzcoa, Spain.
m. Present address: Department of Medicine and Life Sciences, Biomedical Research Park (PRBB), Universitat Pompeu Fabra, Barcelona, Spain.
n. Innopharma Screening Platform, Biofarma Research Group, Centro de Investigación en Medicina Molecular y Enfermedades Crónicas (CIMUS), University of Santiago de Compostela, Santiago de Compostela, Spain.
o. Departament of Pharmacology, Pharmacy and Pharmaceutical Technology. School of Pharmacy. University of Santiago de Compostela. Santiago de Compostela. Spain.
p. Department of Pharmacology, Therapeutics and Toxicology, Institute of Neurosciences, Autonomous University of Barcelona, Bellaterra, Barcelona, Spain.
q. Rega Institute, Department of Microbiology, Immunology and Transplantation, KU Leuven, Leuven, Belgium.
r. Division of Cardiovascular Medicine, Knight Cardiovascular Institute, Oregon Health & Science University, Portland, OR, USA.

* To whom correspondence should be addressed:


**This PDF file includes:**

Synthesis and characterization methods
Radiochemistry methods
Supplementary Figures 1 to 28
Supplementary Tables 1 to 7

### CONTENT


**Synthesis and characterization methods** S6

General procedure A for the synthesis of the acyl chloride S6
General procedure B for the synthesis of hydrazides S6
General procedure C for the synthesis of amides S14
General procedure D for the synthesis of ureas S17
General procedure E for the synthesis of thioureas S19

**Radiochemistry: synthesis and characterization of [$^{11}$C]UB-CB-P3** S23

Supplementary figure 1:
General procedures for the synthesis of the new derivatives. S24

Supplementary figure 2:
*N′*-(2-Hydroxyphenyl)-3,4,8,9-tetramethyltetracyclo[4.4.0.0$^{3,9}$.0$^{4,8}$]decane-1-carbohydrazide (CB-P4). S26

Supplementary figure 3:
*N′*-(2-Methoxyphenyl)-3,4,8,9-tetramethyltetracyclo[4.4.0.0$^{3,9}$.0$^{4,8}$]decane-1-carbohydrazide (CB-P3). S28

Supplementary figure 4:
*N′*-(3-Chlorophenyl)-3,4,8,9-tetramethyltetracyclo[4.4.0.0$^{3,9}$.0$^{4,8}$]decane-1-carbohydrazide (ALT-P39). S30

Supplementary figure 5:
*N′*-(4-Chlorophenyl)-3,4,8,9-tetramethyltetracyclo[4.4.0.0$^{3,9}$.0$^{4,8}$]decane-1-carbohydrazide (ALT-P40). S32

Supplementary figure 6:
*N′*-(2,3-Dichlorophenyl)-3,4,8,9-tetramethyltetracyclo[4.4.0.0$^{3,9}$.0$^{4,8}$]decane-1-carbohydrazide (ALT-P41). S34

Supplementary figure 7:
*N′*-(2,4-Dichlorophenyl)-3,4,8,9-tetramethyltetracyclo[4.4.0.0$^{3,9}$.0$^{4,8}$]decane-1-carbohydrazide (ALT-P42). S36

Supplementary figure 8:
*N′*-(2-Fluorophenyl)-3,4,8,9-tetramethyltetracyclo[4.4.0.0$^{3,9}$.0$^{4,8}$]decane-1-carbohydrazide (CB-P1). S39

Supplementary figure 9:
*N′*-(2-Iodophenyl)-3,4,8,9-tetramethyltetracyclo[4.4.0.0$^{3,9}$.0$^{4,8}$]decane-1-carbohydrazide (ALT-P35). S41

Supplementary figure 10:
3,4,8,9-Tetramethyl-*N'*-(2-(trifluoromethyl)phenyl)tetracyclo[4.4.0.0$^{3,9}$.0$^{4,8}$] decane-1-carbohydrazide (MBX-47). S43

Supplementary figure 11:
3,4,8,9-Tetramethyl-*N'*-(2-(pentafluoro-λ$^{6}$-sulfanyl)phenyl)tetracyclo[4.4.0.0$^{3,9}$.0$^{4,8}$] decane-1-carbohydrazide (ALT-P6). S46

Supplementary figure 12:
*N'*-(3',5'-Dichloropyridin-4'-yl)-3,4,8,9-tetramethyltetracyclo[4.4.0.0$^{3,9}$.0$^{4,8}$]decane-1-carbohydrazide (ALT-P1). S48

Supplementary figure 13:
3,4,8,9-Tetramethyl-*N'*-(quinoline-5'-yl)tetracyclo[4.4.0.0$^{3,9}$0$^{4,8}$]decane-1-carbohydrazide (ALT-P2). S50

Supplementary figure 14:
*N'*-(6-Chloroquinolin-5-yl)-3,4,8,9-tetramethyltetracyclo[4.4.0.0$^{3,9}$.0$^{4,8}$]decane-1-carbohydrazide (ALT-P21). S52

Supplementary figure 15:
3,4,8,9-Tetramethyl-*N*-(quinolin-5-yl)tetracyclo[4.4.0.0$^{3,9}$.0$^{4,8}$]decane-1-carboxamide (ALT-P19). S54

Supplementary figure 16:
3,4,8,9-Tetramethyl-*N*-(quinolin-5-ylmethyl)tetracyclo[4.4.0.0$^{3,9}$.0$^{4,8}$]decane-1-carboxamide (ALT-P20). S56

Supplementary figure 17:
*N*-((3,4,8,9-Tetramethyltetracyclo[4.4.0.0$^{3,9}$.0$^{4,8}$]decan-1-yl)methyl)quinoline-5-carboxamide (ALT-P10). S58

Supplementary figure 18:
*N*-(quinolin-5-yl)-2-(3,4,8,9-tetramethyltetracyclo[4.4.0.0$^{3,9}$.0$^{4,8}$]decan-1-yl)acetamide (ALT-P25). S60

Supplementary figure 19:
1-(Quinolin-5-yl)-3-(3,4,8,9-tetramethyltetracyclo[4.4.0.0$^{3,9}$.0$^{4,8}$]decan-1-yl)urea (ALT-P22). S62

Supplementary figure 20:
1-(Quinolin-5-yl)-3-((3,4,8,9-tetramethyltetracyclo[4.4.0.0$^{3,9}$.0$^{4,8}$]decan-1-yl)methyl) urea hydrochloride (ALT-P16). S64

Supplementary figure 21:
1-(quinolin-5-yl)-3-(3,4,8,9-tetramethyltetracyclo[4.4.0.0$^{3,9}$.0$^{4,8}$]decan-1-yl)thiourea (ALT-P13). S66

Supplementary figure 22:
1-(quinolin-5-yl)-3-((3,4,8,9-tetramethyltetracyclo[4.4.0.0$^{3,9}$.0$^{4,8}$]decan-1-yl)methyl)thiourea (ALT-P14). S68

Supplementary figure 23:
2-cyano-1-(quinolin-5-yl)-3-((3,4,8,9-tetramethyltetracyclo[4.4.0.0$^{3,9}$.0$^{4,8}$]decan-1-yl)methyl)guanidine (ALT-P7). S70

Supplementary figure 24:
Methyl 2-(3,4,8,9-tetramethyltetracyclo[4.4.0.0$^{3,9}$.0$^{4,8}$]decan-1-yl)acetate (ALT-511). S72

Supplementary figure 25:
2-(3,4,8,9-tetramethyltetracyclo[4.4.0.0$^{3,9}$.0$^{4,8}$]decan-1-yl)acetic acid (ALT-560). S74

Supplementary Figure 26:
Synthesis and radiochemical analysis of [$^{11}$C]UB-CB-P3. S75

Supplementary Figure 27:
Kinetic analysis of UB-ALT-P2 binding to WT and mutants hP2X7R S76

Supplementary Figure 28:
Kinetic analysis of UB-ALT-P2 binding to P2X7 receptors across species. S77

Supplementary Table 1:
HPLC–UV chromatogram of the new series of compounds. S78

Supplementary Table 2:
Cryo-EM collection, refinement, and validation statistics. S83

Supplementary Table 3:
Cellular cytotoxicity in HEL, HeLa, Vero and MT4 cell lines. S84

Supplementary Table 4.
Permeability in the PAMPA-BBB assay from 14 commercial drugs and the assayed compounds and predictive penetration in the CNS. S85

Supplementary Table 5:
Species-specific amino acid differences at key positions of the P2X7 receptor classical allosteric pocket. S86

Supplementary Table 6:
Primary and secondary antibodies used for protein level determination by Western blotting in the in vivo UB-ALT-P2 study in 5xFAD murine model. S87

Supplementary Table 7:
Primers and probes used in qPCR studies for the in vivo UB-ALT-P2 assay. S88

**Synthesis and characterization methods**

**General procedure A for the synthesis of the acyl chloride**

To a solution of the acid (0.50 mmol) in neat thionyl chloride (13.5 mmol) is added a drop of DMF at room temperature and then refluxed for 2 h in a round bottom flask provided with a tube of anh. $CaCl_2$. Then, the excess of thionyl chloride is removed under vacuum. Toluene (5 mL) is added to the residue, and the remaining thionyl chloride is azeotropically removed under vacuum to give the acid chloride as a wax in quantitative yield.

**General procedure B for the synthesis of hydrazides**

Anh. triethylamine (1.03 mmol) is added to a solution of the required hydrazine (0.56 mmol) is anh. THF (2 mL) under nitrogen atmosphere. The mixture is stirred for 10 minutes at room temperature. Upon addition of a solution of the corresponding acyl chloride (0.51 mmol) in anh. THF (7 mL), the whole is left stirring overnight at room temperature. The purification procedure is specified below for each compound.

***N'*-(2-Hydroxyphenyl)-3,4,8,9-tetramethyltetracyclo[4.4.0.0$^{3,9}$.0$^{4,8}$]decane-1-carbohydrazide (CB-P4).** 3,4,8,9-Tetramethyltetracyclo[4.4.0.0$^{3,9}$.0$^{4,8}$]decane-1-carbonyl chloride (108 mg, 0.43 mmol) was obtained following the general procedure **A** and, without further purification, was reacted with 2-hydroxyphenylhydrazine·TsOH (139 mg, 0.47 mmol) in the presence of anh. triethylamine (118 µL, 0.85 mmol), following the general procedure **B**. The formed precipitate was filtered off and the organic layer was concentrated under vacuum, and subsequently washed with 4 mL of a mixture of DCM/hexane 1:9, yielding the desired product as a solid (50 mg, 34% yield), mp 213-219 °C. IR (ATR) v: 3329, 3168, 2955, 2859, 1634, 1603, 1504, 1465, 1428, 1383, 1369, 1277, 1239, 1196, 1139, 1101, 1080, 1032, 923, 872, 823, 792, 741, 666, 596, 571 cm$^{-1}$. $^1$H-NMR (400 MHz, DMSO-d$_6$) δ: 0.70 [dd, *J* = 11.6 Hz, *J'* = 2.8 Hz, 2H, 5(7)-H$_a$], 0.91 [d, *J* = 10.8 Hz, 2H, 2(10)-H$_a$], 0.94 [s, 12H, 3(9)-$CH_3$ and 4(8)-$CH_3$], 1.81 [dd, *J* = 11.6 Hz, *J'* = 1.6 Hz, 2H, 5(7)-H$_b$] 1.94 [d, *J* = 10.8 Hz, 2H, 2(10)-H$_b$], 2.45 (m, 1H, 6-H), 6.56-6.72 (cs, 5H, 3'-H, 4'-H, 5'-H, 6'-H, NH), 9.38 (s, 1H, OH), 9.50 (m, 1H, NH). $^{13}$C-NMR (100.6 MHz, DMSO-d$_6$) δ: 15.48 [$CH_3$, C3(9)-$\underline{C}H_3$ or C4(8)-$\underline{C}H_3$], 15.52 [$CH_3$, C4(8)-$\underline{C}H_3$ or C3(9)-$\underline{C}H_3$], 37.1 (CH, C6), 37.8 [$CH_2$, C5(7)], 40.5 [$CH_2$, C2(10)], 44.5 [C, C3(9) or C4(8)], 44.9 [C, C4(8) or C3(9)], 47.7 (C, C1), 112.3 (CH, C3' or C6'), 114.2 (CH, C6' or C3'), 119.2 (CH, C4' or C5'), 119.3 (CH, C5' or C4'), 137.8 (C, C1'), 144.4 (C, C2'), 175.3 (C, CO). HRMS-ESI+ *m/z* [M+H]$^+$ calcd for $[C_{21}H_{29}N_2O_2]^+$: 341.2224, found: 341.2225.

Elemental analysis: Calcd for $C_{21}H_{28}N_2O_2·0.3CH_2Cl_2$: C 69.91, H 7.88, N 7.66. Found: C 70.04, H 7.88, N 7.49. HPLC-UV purity at 254 nm = 97.73%.

***N'*-(2-Methoxyphenyl)-3,4,8,9-tetramethyltetracyclo[4.4.0.0$^{3,9}$.0$^{4,8}$]decane-1-carbohydrazide (CB-P3).** 3,4,8,9-Tetramethyltetracyclo[4.4.0.0$^{3,9}$.0$^{4,8}$]decane-1-carbonyl chloride (108 mg, 0.43 mmol) was obtained following the general procedure **A** and, without further purification, was reacted with 2-methoxyphenylhydrazine hydrochloride (80mg, 0.7 mmol) in the presence of anh. triethylamine (118 µL, 0.85 mmol), following the general procedure **B**. The solved was evaporated and DCM (10 mL) was added. The organic phase was washed with 2 N HCl solution (2 x 5 mL), 2 N NaOH solution (2 x 5 mL) and brine (2 x 5 mL) and dried over anhydrous $Na_2SO_4$, filtered, and concentrated under vacuum to give a residue. An analytical sample of the title compound was obtained by crystallization from DCM/Pentane (10 mg, 7% yield), mp 170-171 °C. IR (ATR) ν: 3347, 3324, 2945, 2859, 1642, 1600, 1581, 1498, 1452, 1426, 1316, 1252, 1215, 1182, 1108, 1084, 1047, 1029, 915, 856, 740, 686, 600 cm$^{-1}$. $^1$H-NMR (400 MHz, $CDCl_3$) δ: 0.79 [dd, *J* = 11.2 Hz, *J'* = 2.8 Hz, 2H, 5(7)-$H_a$], 0.96 [m, 2H, 2(10)-$H_a$], 0.97 [s, 6H, 3(9)-$CH_3$ or 4(8)-$CH_3$], 0.98 [s, 6H, 4(8)-$CH_3$ or 3(9)-$CH_3$], 1.84 [dd, *J* = 11.6 Hz, *J'* = 1.6 Hz, 2H, 5(7)-$H_b$] 1.99 [d, *J* = 10.8 Hz, 2H, 2(10)-$H_b$], 2.57 (m, 1H, 6-H), 3.88 (s, 3H, $OCH_3$), 6.58 (broad s, 1H, NH), 6.78-6.89 (cs, 4H, 3'-H, 4'-H, 5'-H, 6'-H), 7.37 (s, 1H, NH). $^{13}$C-NMR (100.6 MHz, $CDCl_3$) δ: 15.69 [$CH_3$, C3(9)-$\underline{C}H_3$ or C4(8)-$\underline{C}H_3$], 15.76 [$CH_3$, C4(8)-$\underline{C}H_3$ or C3(9)-$\underline{C}H_3$], 37.8 (CH, C6), 38.4 [$CH_2$, C5(7)], 41.3 [$CH_2$, C2(10)], 45.2 [C, C3(9) or C4(8)], 45.7 [C, C4(8) or C3(9)], 48.4 (C, C1), 55.8 ($CH_3$, $OCH_3$), 110.6 (CH, C3' or C6'), 112.8 (CH, C6' or C3'), 121.0 (CH, C4' or C5'), 121.1 (CH, C5' or C4'), 137.9 (C, C1'), 147.7 (C, C2'), 176.7 (C, CO). HRMS-ESI+ *m/z* $[M+H]^+$ calcd for $[C_{22}H_{31}N_2O_2]^+$: 355.2380, found: 355.2379. Elemental analysis: Calcd for $C_{22}H_{30}N_2O_2$: C 74.54, H 8.53, N 7.90. Found: C 74.26, H 8.49, N 7.70. HPLC-UV purity at 254 nm = 97.88%.

***N'*-(3-Chlorophenyl)-3,4,8,9-tetramethyltetracyclo[4.4.0.0$^{3,9}$.0$^{4,8}$]decane-1-carbohydrazide (ALT-P39).** 3,4,8,9-Tetramethyltetracyclo[4.4.0.0$^{3,9}$.0$^{4,8}$]decane-1-carbonyl chloride (85 mg, 0.34 mmol) was obtained following the general procedure **A** and, without further purification, was reacted with (3-chlorophenyl)hydrazine hydrochloride (66 mg, 0.37 mmol) in the presence of anh. triethylamine (93 µL, 0.67 mmol), following the general procedure **B**. The solvent was evaporated under vacuum and DCM (10 mL) was added. The organic phase was washed with 2 N HCl solution (2 x 5 mL), 2 N NaOH solution (2 x 5 mL) and brine (2 x 5 mL), dried over anhydrous $Na_2SO_4$, filtered, and concentrated under vacuum to give a residue. An analytical sample of the title compound was obtained by crystallization from DCM/Pentane (70 mg, 58% yield), mp 199-200 °C. IR (ATR) ν: 3288, 2947, 2920, 2864, 1670, 1644, 1596, 1479,

1467, 1419, 1385, 1371, 1307, 1285, 1264, 1247, 1224, 1206, 1156, 1123, 1097, 1071, 1017, 992, 919, 852, 818, 768, 698, 680, 648, 637, 606 $cm^{-1}$. $^1$H-NMR (400 MHz, $CDCl_3$) δ: 0.80 [dd, *J* = 11.6 Hz, *J'* = 2.0 Hz, 2H, 5(7)-$H_a$], 0.96 [d, *J* = 10.4 Hz, 2H, 2(10)-$H_a$], 0.97 [s, 6H, 3(9)-$CH_3$ or 4(8)-$CH_3$], 0.98 [s, 6H, 4(8)-$CH_3$ or 3(9)-$CH_3$], 1.84 [dd, *J* = 11.8 Hz, *J'* = 1.0 Hz, 2H, 5(7)-$H_b$] 1.99 [d, *J* = 10.8 Hz, 2H, 2(10)-$H_b$], 2.56 (m, 1H, 6-H), 6.23 (d, *J* = 3.4 Hz, 1H, NH), 6.70 (dd, *J* = 8.0 Hz, *J'* = 1.6 Hz, 1H, 6'-H), 6.80 (dd, *J* = 1.6 Hz, *J'* = 1.2 Hz, 1H, 2'-H), 6.85 (dd, *J* = 8.0 Hz, *J'* = 1.6 Hz, 1H, 4'-H), 7.13 (t, *J* = 8.0 Hz, 1H, 5'-H), 7.43 (d, *J* = 3.4 Hz, 1H, NH). $^{13}$C-NMR (100.6 MHz, $CDCl_3$) δ: 15.67 [$CH_3$, C3(9)-$\underline{C}H_3$ or C4(8)-$\underline{C}H_3$], 15.73 [$CH_3$, C4(8)-$\underline{C}H_3$ or C3(9)-$\underline{C}H_3$], 37.7 (CH, C6), 38.3 [$CH_2$, C5(7)], 41.3 [$CH_2$, C2(10)], 45.2 [C, C3(9) or C4(8)], 45.8 [C, C4(8) or C3(9)], 48.4 (C, C1), 112.O (CH, C6'), 113.8 (CH, C2'), 121.3 (CH, C4'), 130.4 (CH, C5'), 135.1 (C, C3'), 149.9 (C, C1'), 177.4 (C, CO). HRMS-ESI+ *m/z* $[M+H]^+$ calcd for $[C_{21}H_{28}ClN_2O]^+$: 359.1885, found: 359.1894. HPLC-UV purity at 254 nm = 98.87%.

***N'*-(4-Chlorophenyl)-3,4,8,9-tetramethyltetracyclo[4.4.0.0$^{3,9}$.0$^{4,8}$]decane-1-carbohydrazide (ALT-P40).** 3,4,8,9-Tetramethyltetracyclo[4.4.0.0$^{3,9}$.0$^{4,8}$]decane-1-carbonyl chloride (85 mg, 0.34 mmol) was obtained following the general procedure **A** and, without further purification, was reacted with (4-chlorophenyl)hydrazine hydrochloride (66 mg, 0.37 mmol) in the presence of anh. triethylamine (93 µL, 0.67 mmol), following the general procedure **B**. The solvent was evaporated under vacuum and DCM (10 mL) was added. The organic phase was washed with 2 N HCl solution (2 x 5 mL), 2 N NaOH solution (2 x 5 mL) and brine (2 x 5 mL) and dried over anhydrous $Na_2SO_4$, filtered, and concentrated under vacuum to give a residue. An analytical sample of the title compound was obtained by crystallization from DCM/Pentane (51 mg, 43% yield), mp 200-201 °C. IR (ATR) v: 3285, 2949, 2920, 2863, 1667, 1644, 1596, 1489, 1385, 1371, 1306, 1285, 1247, 1223, 1155, 1087, 1020, 992, 920, 852, 834, 818, 768, 699, 680, 637, 602 $cm^{-1}$. $^1$H-NMR (400 MHz, $CDCl_3$) δ: 0.80 [dd, *J* = 11.6 Hz, *J'* = 1.6 Hz, 2H, 5(7)-$H_a$], 0.95 [d, *J* = 11.6 Hz, 2H, 2(10)-$H_a$], 0.97 [s, 6H, 3(9)-$CH_3$ or 4(8)-$CH_3$], 0.98 [s, 6H, 4(8)-$CH_3$ or 3(9)-$CH_3$], 1.84 [d, *J* = 10.8 Hz, 2H, 5(7)-$H_b$] 1.99 [d, *J* = 10.8 Hz, 2H, 2(10)-$H_b$], 2.55 (m, 1H, 6-H), 6.17 (broad s, 1H, NH), 6.76 [d, *J* = 8.8 Hz, 2H, 2'(6')-H], 7.17 [d, *J* = 8.8 Hz, 2H, 3'(5')-H], 7.41 (broad s, 1H, NH). $^{13}$C-NMR (100.6 MHz, $CDCl_3$) δ: 15.67 [$CH_3$, C3(9)-$\underline{C}H_3$ or C4(8)-$\underline{C}H_3$], 15.74 [$CH_3$, C4(8)-$\underline{C}H_3$ or C3(9)-$\underline{C}H_3$], 37.7 (CH, C6), 38.3 [$CH_2$, C5(7)], 41.3 [$CH_2$, C2(10)], 45.2 [C, C3(9) or C4(8)], 45.8 [C, C4(8) or C3(9)], 48.4 (C, C1), 115.1 [CH, C2'(6')], 126.1 (C, C4'), 129.2 [CH, C3'(5')], 147.2 (C, C1'), 177.4 (C, CO). HRMS-ESI+ *m/z* $[M+H]^+$ calcd for $[C_{21}H_{28}ClN_2O]^+$: 359.1885, found: 359.1893. Elemental analysis: Calcd for $C_{21}H_{27}ClN_2O \cdot 0.2H_2O$: C 69.58, H 7.62, N 7.73. Found: C 69.77, H 7.49, N 7.70. HPLC-UV purity at 254 nm = 99.59%.

***N'*-(2,3-Dichlorophenyl)-3,4,8,9-tetramethyltetracyclo[4.4.0.0$^{3,9}$.0$^{4,8}$]decane-1-carbohydrazide (ALT-P41).** 3,4,8,9-Tetramethyltetracyclo[4.4.0.0$^{3,9}$.0$^{4,8}$]decane-1-carbonyl chloride (85 mg, 0.34 mmol) was obtained following the general procedure **A** and, without further purification, was reacted with (2,3-dichlorophenyl)hydrazine hydrochloride (78 mg, 0.37 mmol) in the presence of anh. triethylamine (93 µL, 0.67 mmol), following the general procedure **B**. The solved was evaporated and DCM (10 mL) was added. The organic phase was washed with 2 N HCl solution (2 x 5 mL), 2 N NaOH solution (2 x 5 mL) and brine (2 x 5 mL) and dried over anhydrous $Na_2SO_4$, filtered, and concentrated under vacuum to give a residue. An analytical sample of the title compound was obtained by crystallization from DCM/Pentane (91 mg, 69% yield), mp 237-238 °C. IR (ATR) ν: 3362, 3304, 2950, 2864, 1656, 1583, 1480, 1451, 1406, 1385, 1372, 1304, 1272, 1226, 1190, 1148, 1124, 1092, 1046, 1017, 920, 895, 852, 834, 769, 738, 701, 679, 607, 573, 565 cm$^{-1}$. $^1$H-NMR (400 MHz, $CDCl_3$) δ: 0.80 [dd, *J* = 12.4 Hz, *J'* = 2.0 Hz, 2H, 5(7)-H$_a$], 0.95 [d, *J* = 11.6 Hz, 2H, 2(10)-H$_a$], 0.97 [s, 6H, 3(9)-$CH_3$ or 4(8)-$CH_3$], 0.98 [s, 6H, 4(8)-$CH_3$ or 3(9)-$CH_3$], 1.84 [dd, *J* = 11.6 Hz, *J'* = 1.2 Hz, 2H, 5(7)-H$_b$] 1.99 [d, *J* = 12.4 Hz, 2H, 2(10)-H$_b$], 2.56 (m, 1H, 6-H), 6.53 (d, *J* = 3.2 Hz, 1H, NH), 6.76 (dd, *J* = 8.0 Hz, *J'* = 1.2 Hz, 2H, 6'-H), 6.99 (dd, *J* = 8.0 Hz, *J'* = 1.2 Hz, 2H, 4'-H), 7.08 (t, *J* = 8.0 Hz, 1H, 5'-H), 7.47 (d, *J* = 3.2 Hz, 1H, NH). $^{13}$C-NMR (100.6 MHz, $CDCl_3$) δ: 15.66 [$CH_3$, C3(9)-$\underline{C}H_3$ or C4(8)-$\underline{C}H_3$], 15.72 [$CH_3$, C4(8)-$\underline{C}H_3$ or C3(9)-$\underline{C}H_3$], 37.7 (CH, C6), 38.3 [$CH_2$, C5(7)], 41.3 [$CH_2$, C2(10)], 45.2 [C, C3(9) or C4(8)], 45.8 [C, C4(8) or C3(9)], 48.4 (C, C1), 111.5 (CH, C6'), 118.2 (C, C2'), 122,1 (CH, C4'), 127.8 (CH, C5'), 133.3 (C, C3'), 146.1 (C, C1'), 177.0 (C, CO). HRMS-ESI+ *m/z* $[M+H]^+$ calcd for $[C_{21}H_{27}Cl_2N_2O]^+$: 393.1495, found: 393.1505. Elemental analysis: Calcd for $C_{21}H_{26}Cl_2N_2O$: C 64.12, H 6.66, N 7.12. Found: C 63.96, H 6.59, N 7.06.

***N'*-(2,4-Dichlorophenyl)-3,4,8,9-tetramethyltetracyclo[4.4.0.0$^{3,9}$.0$^{4,8}$]decane-1-carbohydrazide (ALT-P42).** 3,4,8,9-Tetramethyltetracyclo[4.4.0.0$^{3,9}$.0$^{4,8}$]decane-1-carbonyl chloride (85 mg, 0.34 mmol) was obtained following the general procedure **A** and, without further purification, was reacted with (2,4-dichlorophenyl)hydrazine hydrochloride (78 mg, 0.37 mmol) in the presence of anh. triethylamine (93 µL, 0.67 mmol), following the general procedure **B**. The solved was evaporated and DCM (10 mL) was added. The organic phase was washed with 2 N HCl solution (2 x 5 mL), 2 N NaOH solution (2 x 5 mL) and brine (2 x 5 mL) and dried over anhydrous $Na_2SO_4$, filtered, and concentrated under vacuum to give a residue. An analytical sample of the title compound was obtained by crystallization from DCM/Pentane (72 mg, 55% yield), mp 209-210 °C. IR (ATR) ν: 3289, 2948, 2863, 1656, 1583, 1542, 1477, 1453, 1423, 1385, 1372, 1304, 1284, 1224, 1091, 1047, 1018, 922, 853, 803, 769, 738, 704, 681, 625, 605 cm$^{-1}$. $^1$H-

NMR (400 MHz, $CDCl_3$) δ: 0.80 [dd, *J* = 11.6 Hz, *J'* = 2.0 Hz, 2H, 5(7)-$H_a$], 0.95 [d, *J* = 11.6 Hz, 2H, 2(10)-$H_a$], 0.97 [s, 6H, 3(9)-$CH_3$ or 4(8)-$CH_3$], 0.98 [s, 6H, 4(8)-$CH_3$ or 3(9)-$CH_3$], 1.84 [dd, *J* = 11.6 Hz, *J'* = 1.0 Hz, 2H, 5(7)-$H_b$] 1.99 [d, *J* = 10.8 Hz, 2H, 2(10)-$H_b$], 2.56 (m, 1H, 6-H), 6.42 (d, *J* = 3.0 Hz, 1H, NH), 6.78 (d, *J* = 8.4 Hz, 2H, 6'-H), 7.12 (dd, *J* = 8.4 Hz, *J'* = 2.2 Hz, 2H, 5'-H), 7.29 (d, *J* = 2.2 Hz, 1H, 3'-H), 7.41 (d, *J* = 3.0 Hz, 1H, NH). $^{13}$C-NMR (100.6 MHz, $CDCl_3$) δ: 15.66 [$CH_3$, C3(9)-$\underline{C}H_3$ or C4(8)-$\underline{C}H_3$], 15.73 [$CH_3$, C4(8)-$\underline{C}H_3$ or C3(9)-$\underline{C}H_3$], 37.7 (CH, C6), 38.3 [$CH_2$, C5(7)], 41.3 [$CH_2$, C2(10)], 45.2 [C, C3(9) or C4(8)], 45.8 [C, C4(8) or C3(9)], 48.4 (C, C1), 114.5 (CH, C6'), 120.5 (C, C2'), 125.8 (CH, C4'), 127.9 (CH, C5'), 129.3 (CH, C3'), 143.3 (C, C1'), 177.1 (C, CO). HRMS-ESI+ *m/z* $[M+H]^+$ calcd for $[C_{21}H_{27}Cl_2N_2O]^+$: 393.1495, found: 393.1508. Elemental analysis: Calcd for $C_{21}H_{26}Cl_2N_2O\cdot 0.2H_2O$: C 63.54, H 6.70, N 7.06. Found: C 63.57, H 6.56, N 6.96. HPLC-UV purity at 254 nm = 99.24%.

***N'*-(2-Fluorophenyl)-3,4,8,9-tetramethyltetracyclo[4.4.0.0$^{3,9}$.0$^{4,8}$]decane-1-carbohydrazide (CB-P1).** 3,4,8,9-Tetramethyltetracyclo[4.4.0.0$^{3,9}$.0$^{4,8}$]decane-1-carbonyl chloride (161 mg, 0.64 mmol) was obtained following the general procedure **A** and, without further purification, was reacted with (2-fluorophenyl)hydrazine hydrochloride (114 mg, 0.70 mmol) in the presence of anhydrous triethylamine (200 µL, 1.28 mmol), following the general procedure **B**. The formed precipitate was filtered off and the organic layer was concentrated under vacuum, subsequently washed with 4 mL of a mixture of DCM/hexane (1/9), yielding the desired product as a beige solid (66 mg, 50% yield), mp 139 ˚C. IR (ATR) v: 3401, 3327, 3281, 2954, 2928, 2864, 1677, 1645, 1619, 1588, 1500, 1454, 1384, 1371, 1309, 1282, 1246, 1221, 1197, 1187, 1155, 1102, 1085, 1031, 918, 829, 740, 713, 635 cm$^{-1}$. $^1$H-NMR (400 MHz, $CDCl_3$) δ: 0.78 [dd, *J* = 10.8 Hz, *J'* = 2.5 Hz, 2H, 5(7)-$H_a$], 0.96 [m, 2H, 2(10)-$H_a$], 0.966 [s, 6H, 3(9)-$CH_3$ or 4(8)-$CH_3$], 0.972 [s, 6H, 4(8)-$CH_3$ or 3(9)-$CH_3$], 1.83 [dd, *J* = 11.6 Hz, *J'* = 1.5 Hz, 2H, 5(7)-$H_b$], 1.99 [d, *J* = 11.2 Hz, 2H, 2(10)-$H_b$], 2.55 (broad s, 1H, 6-H), 6.35 (broad s, 1H, NH), 6.82 (m, 1H, 4'-H), 6.88 (dt, *J* = 8.0 Hz, *J'* = 2.5 Hz, 2H, 6'-H), 6.97-7.04 (cs, 2H, 3'-H and 5'-H), 7.53 (broad s, 1H, NH). $^{13}$C-NMR (100.6 MHz, $CDCl_3$) δ: 15.66 [$CH_3$, C3(9)-$\underline{C}H_3$ or C4(8)-$\underline{C}H_3$], 15.71 [$CH_3$, C4(8)-$\underline{C}H_3$ or C3(9)-$\underline{C}H_3$], 37.7 (CH, C6), 38.3 [$CH_2$, C5(7)], 41.2 [$CH_2$, C2(10)], 45.2 [C, C3(9) or C4(8)], 45.7 [C, C4(8) or C3(9)], 48.4 (C, C1), 114.8 (CH, d, $J_{C\text{-}F}$ = 2.0 Hz, C6'), 115.4 (CH, d, $J_{C\text{-}F}$ = 18.1 Hz, C3'), 121.4 (CH, d, $J_{C\text{-}F}$ = 7.1 Hz, C4'), 124.6 (CH, d, $J_{C\text{-}F}$ = 3.0 Hz, C5'), 136.5 (C, d, $J_{C\text{-}F}$ = 10.1 Hz, C1'), 151.9 (C, d, $J_{C\text{-}F}$ = 241.4 Hz, C2'), 177.2 (C, CO). $^{19}$F-NMR (376.3 MHz, $CDCl_3$) δ: -133.5 (s, 1F, 1'-F). HRMS-ESI+ *m/z* $[M+H]^+$ calcd for $[C_{21}H_{28}FN_2O]^+$: 342.2180, found: 343.2177. HPLC purity at 220 nm= 97.29%.

***N'*-(2-Iodophenyl)-3,4,8,9-tetramethyltetracyclo[4.4.0.0$^{3,9}$.0$^{4,8}$]decane-1-carbohydrazide ALT-P35).** 3,4,8,9-Tetramethyltetracyclo[4.4.0.0$^{3,9}$.0$^{4,8}$]decane-1-carboxylic acid (143 mg, 0.61 mmol), (2-iodophenyl)hydrazine (170 mg, 0.91 mmol), *N*-ethyl-*N'*-(3-dimethylaminopropyl)carbodiimide hydrochloride (175 mg, 0.91 mmol), DIPEA (265 µL, 1.52 mmol) were solved in anh. DCM (2.5 mL) and stirred for 1.5 h. The solution formed was concentrated under vacuum. Purification in silica gel using as eluent ethyl acetate to ethylacetate/methanol mixture (1/99) gave the title compound as a white solid (168 mg, 61% yield), mp >200 °C (dec.). IR (ATR) ν: 3322, 2902, 2859, 1646, 1584, 1481, 1450, 1432, 1383, 1369, 1313, 1281, 1244, 1156, 1110, 1088, 1011, 801, 745, 715, 675 cm$^{-1}$. $^{1}$H-NMR (500 MHz, $CDCl_3$) δ: 0.81 [dd, *J* = 11.5 Hz, *J'* = 2.5 Hz, 2H, 5(7)-H$_a$], 0.97 [m, 2H, 2(10)-H$_a$], 0.98 [s, 6H, 3(9)-$CH_3$ or 4(8)-$CH_3$], 0.99 [s, 6H, 4(8)-$CH_3$ or 3(9)-$CH_3$], 1.85 [d, *J* = 11.5 Hz, *J'* = 1.0 Hz, 2H, 5(7)-H$_b$], 2.01 [d, *J* = 11.0 Hz, 2H, 2(10)-H$_b$], 2.59 (s, 1H, 6-H), 6.28 (broad s, 1H, NH), 6.65 (dt, *J* = 7.5 Hz, *J'* = 1.5 Hz, 5'-H), 6.79 (d, *J* = 7.5 Hz, 1H, 6'-H), 7.23 (td, *J* = 7.5 Hz, *J'* = 1 Hz, 1H, 4'-H), 7.43 (broad s, 1H, NH), 7.69 (dd, *J* = 7.5 Hz, *J'* = 1.5 Hz, 3'-H). $^{13}$C-NMR (125.7 MHz, $CDCl_3$) δ: 15.7 [$CH_3$, C3(9)-$CH_3$ or C4(8)-$CH_3$], 15.8 [$CH_3$, C4(8)-$CH_3$ or C3(9)- $CH_3$], 37.8 (CH, C6), 38.3 [$CH_2$, C5(7)], 41.3 [$CH_2$, C2(10)], 45.2 [C, C3(9) or C4(8)], 45.8 [C, C4(8) or C3(9)], 48.4 (C, C1), 84.0 (C, C2'), 113.6 (CH, C6'), 122.9 (CH, C5'), 129.4 (CH, C4'), 139.3 (CH, C3'), 147.5 (CH, C11'), 176.9 (C, CO). HRMS-ESI+ *m/z* $[M+H]^+$ calcd for $[C_{21}H_{26}IN_2O]^+$: 449.1095, found: 449.1083. HPLC purity at 220 nm = 95.80%.

**3,4,8,9-Tetramethyl-*N'*-(2-(trifluoromethyl)phenyl)tetracyclo[4.4.0.0$^{3,9}$.0$^{4,8}$]decane-1-carbohydrazide (MBX-47).** 3,4,8,9-Tetramethyltetracyclo[4.4.0.0$^{3,9}$.0$^{4,8}$]decane-1-carbonyl chloride (110 mg, 0.44 mmol) was obtained following the general procedure **A** and, without further purification, was reacted with 2-(trifluoromethyl)phenylhydrazine hydrochloride (103 mg, 0.48 mmol) in the presence of anh. triethylamine (122 µL, 0.88 mmol), following the general procedure **B**. The formed precipitate was filtered off and the organic layer was concentrated under vacuum, subsequently washed with 2 mL of a mixture of DCM/hexane (1/9), yielding the desired product as a beige solid (138 mg, 80% yield). mp 206 °C. IR (ATR) ν: 3326, 2956, 2875, 1658, 1618, 1587, 1489, 1317, 1276, 1246, 1168, 1107, 1062, 1034, 768, 756 cm$^{-1}$. $^{1}$H-NMR (400 MHz, $CDCl_3$) δ: 0.81 [dd, *J* = 11.6 Hz, *J'* = 2.4 Hz, 2H, 5(7)-H$_a$], 0.98 [s, 6H, 3(9)-$CH_3$ or 4(8)-$CH_3$], 0.99 [s, 6H, 4(8)-$CH_3$ or 3(9)-$CH_3$], 1.01 [broad s, 2H, 2(10)-H$_a$], 1.86 [dd, *J* = 12 Hz, *J'* = 1.6 Hz, 2H, 5(7)-H$_b$], 2.03 [d, *J* = 10.8 Hz, 2H, 2(10)-H$_b$], 2.60 (broad s, 1H, 6-H), 6.60 (broad s, 1H, NH), 6.95 (t, *J* = 7.6 Hz, 4'-H), 7.00 (d, *J* = 8.4 Hz, 1H, 6'-H), 7.32 (d, *J* = 3.2 Hz, 1H, NH), 7.41 (t, *J* = 8 Hz, 1H, 5'-H), 7.51 (d, *J* = 8 Hz, 1H, 3'-H). $^{13}$C-NMR (100.6 MHz, $CDCl_3$) δ: 15.67 [$CH_3$, C3(9)-$\underline{C}H_3$ or C4(8)-$\underline{C}H_3$], 15.74 [$CH_3$, C4(8)-$\underline{C}H_3$ or C3(9)-$\underline{C}H_3$], 37.7 (CH,

C6), 38.3 [$CH_2$, C5(7)], 41.3 [$CH_2$, C2(10)], 45.2 [C, C3(9) or C4(8)], 45.8 [C, C4(8) or C3(9)], 48.4 (C, C1), 113.7 (CH, C6'), 115.4 (C, q, $J_{C-F}$ = 30.2 Hz, C2'), 120.4 (CH, C4'), 124.7 (C, q, $J_{C-F}$ = 272.6 Hz, $\underline{C}F_3$), 126.7 (CH, q, $J_{C-F}$ = 6 Hz, C3'), 133.2 (CH, C5'), 146.2 (C, C1'), 177.1 (C, CO). HRMS-ESI+ *m/z* $[M+H]^+$ calcd for $[C_{22}H_{28}F_3N_2O]^+$: 393.2148, found: 393.2148. Elemental analysis: calcd for $C_{22}H_{27}F_3N_2O$: C 67.33, H 6.93, N 7.14. Found: C 67.05, H 7.08, N 6.97.

**3,4,8,9-Tetramethyl-*N'*-(2-(pentafluoro-$\Lambda^6$-sulfanyl)phenyl)tetracyclo[4.4.0.0$^{3,9}$.0$^{4,8}$]decane-1-carbohydrazide (ALT-P6).** 3,4,8,9-Tetramethyltetracyclo[4.4.0.0$^{3,9}$.0$^{4,8}$]decane-1-carbonyl chloride (135 mg, 0.53 mmol) was obtained following the general procedure **A** and, without further purification, was reacted with (2-pentafluorosulfanylphenyl)hydrazine hydrochloride (159 mg, 0.59 mmol) in the presence of anh. triethylamine (148 µL, 1.06 mmol), following the general procedure **B**. The brown suspension was evaporated under vacuum to dryness. The crude was purified by Combiflash® in silica gel using as eluent hexane to ethyl acetate/hexane mixture (1/9) to give a white solid (58 mg, 25% yield), mp 183 °C. IR (NaCl disk) ν: 3263, 2951, 2927, 2866, 1653, 1603, 1518, 1505, 1472, 1456, 1373, 841, 827, 750 $cm^{-1}$. $^1$H-NMR (400 MHz, $CDCl_3$) δ: 0.81 [dd, *J* = 11.6 Hz, *J'* = 2.4 Hz, 2H, 5(7)-$H_a$], 0.98 [s, 6H, 3(9)-$CH_3$ or 4(8)-$CH_3$], 0.99 [s, 6H, 4(8)-$CH_3$ or 3(9)-$CH_3$], 1.01 [broad s, 2H, 2(10)-$H_a$], 1.85 [dd, *J* = 12 Hz, *J'* = 1.2 Hz, 2H, 5(7)-$H_b$], 2.03 [d, *J* = 10.8 Hz, 2H, 2(10)-$H_b$], 2.60 (broad s., 1H, 6-H), 6.91-6.93 (c.s., 2H, 4'-H and NH), 7.07 (d, *J* = 8.4 Hz, 1H, 6'-H), 7.34-7.38 (c.s., 2H, 5'-H and NH), 7.67 (dd, *J* = 8.4 Hz, *J'* = 1.2 Hz, 3'-H). $^{13}$C-NMR (100.6 MHz, $CDCl_3$) δ: 15.7 [$CH_3$, C3(9)-$\underline{C}H_3$ or C4(8)-$\underline{C}H_3$], 15.8 [$CH_3$, C4(8)-$\underline{C}H_3$ or C3(9)-$\underline{C}H_3$], 37.7 (CH, C6), 38.3 [$CH_2$, C5(7)], 41.3 [$CH_2$, C2(10)], 45.2 [C, C3(9) or C4(8)], 45.8 [C, C4(8) or C3(9)], 48.5 (C, C1), 114.9 (CH, C6'), 119.9 (CH, C4'), 128.6 [m, C2'(3')], 132.7 (CH, C5'), 143.1 (C, C1'), 177.0 (C, CO). $^{19}$F-NMR (376.5 MHz, $CDCl_3$) δ: 66.4 (d, *J* = 148.8 Hz, 4 F, $S\underline{F_4}F$), 87.1(quint., *J* = 148.8 Hz, 1 F, $SF_4\underline{F}$). HRMS-ESI+ *m/z* $[M+H]^+$ calcd for $[C_{21}H_{28}F_5N_2OS]^+$: 451.1837, found: 451.1833. Elemental analysis: Calcd for $C_{21}H_{27}F_5N_2OS$: C 55.99, H 6.04, N 6.22. Found: C 56.03, H 6.33, N 6.28.

***N'*-(3',5'-Dichloropyridin-4'-yl)-3,4,8,9-tetramethyltetracyclo[4.4.0.0$^{3,9}$.0$^{4,8}$]decane-1-carbohydrazide (ALT-P1).** 3,4,8,9-Tetramethyltetracyclo[4.4.0.0$^{3,9}$.0$^{4,8}$]decane-1-carbonyl chloride (129 mg, 0.51 mmol) was obtained following the general procedure **A** and, without further purification, was reacted with 3,5-dichloro-4-hydrazineylpyridine (100 mg, 0.56 mmol) in the presence of anhydrous triethylamine (142 µL, 1.02 mmol), following the general procedure **B**. The formed precipitate was filtered off and the organic layer was concentrated under vacuum to give a yellow-light solid (55 mg, 27% yield). An analytical sample was obtained by crystallization from DMC/pentane, mp 170 °C. IR

(ATR) ν: 3337, 2951, 2864, 1658, 1567, 1488, 1479, 1455, 1404, 1385, 1371, 1303, 1280, 1238, 1206, 1171, 1087, 1039, 1020, 901, 843, 800, 729, 680, 666 $cm^{-1}$. $^{1}H$-NMR (400 MHz, DMSO-$d_6$) δ: 0.68 [dd, *J* = 11.2 Hz, *J'* = 2.8 Hz, 2H, 5(7)-$H_a$], 0.89 [d, *J* = 11.2 Hz, 2H, 2(10)-$H_a$], 0.93 [s, 12H, 3(9)-$CH_3$ and 4(8)-$CH_3$], 1.77 [dd, *J* = 11.6 Hz, *J'* = 1.2 Hz, 2H, 5(7)-$H_b$] 1.92 [d, *J* = 11.2 Hz, 2H, 2(10)-$H_b$], 2.52 (m, 1H, 6-H), 8.07 (broad s, 1H, NH), 8.23 [s, 2H, 2'(6')-H], 9.82 (broad s, 1H, NH). $^{13}C$-NMR (100.6 MHz, DMSO-$d_6$) δ: 15.45 [$CH_3$, C3(9)-$\underline{C}H_3$ or C4(8)-$\underline{C}H_3$], 15.52 [$CH_3$, C4(8)-$\underline{C}H_3$ or C3(9)-$\underline{C}H_3$], 36.8 (CH, C6), 37.7 [$CH_2$, C5(7)], 40.5 [$CH_2$, C2(10)], 44.5 [C, C3(9) or C4(8)], 44.8 [C, C4(8) or C3(9)], 47.6 (C, C1), 117.1 [C, C3'(5')], 147.3 (C, C4'), 148.1 [CH, C2'(6')], 174.7 (C, CO). HRMS-ESI+ *m/z* $[M+H]^+$ calcd for $[C_{20}H_{26}Cl_2N_3O]^+$: 394.1447, found: 394.1445. Elemental analysis: Calcd for $C_{20}H_{25}Cl_2N_3O \cdot 0.5H_2O$: C 59.56, H 6.50, N 10.42. Found: C 59.62, H 6.67, N 10.21. HPLC purity at 220 nm = 100%.

**3,4,8,9-Tetramethyl-*N'*-(quinoline-5'-yl)tetracyclo[4.4.0.0$^{3,9}$0$^{4,8}$]decane-1-carbohydrazide (ALT-P2).** 3,4,8,9-Tetramethyltetracyclo[4.4.0.0$^{3,9}$.0$^{4,8}$]decane-1-carbonyl chloride (485 mg, 1.92 mmol) was obtained following the general procedure **A** and, without further purification, was reacted with 5-hydrazineylquinoline dihydrochloride (534 mg, 2.30 mmol) in the presence of anh. triethylamine (533 μL, 3.84 mmol), following the general procedure **B**. The formed precipitate was filtered off and the organic layer was concentrated under vacuum. Purification by Combiflash® in silica gel using as eluent hexane to ethyl acetate/hexane mixture (8/2) gave a yellow solid (210 mg, 24% yield). Its dihydrochloride was obtained by adding an excess of HCl / dioxane to a solution of the compound in DCM, followed by evaporation to obtain the dihydrochloride salt as a red wax (269 mg) which was washed with pentane to obtain a red solid, mp >200 ˚C (dec.). IR (NaCl disk) ν: 3266, 2948, 2919, 2864, 1660, 1618, 1591, 1577, 1502, 1466, 1454, 1411, 1383, 1373, 1363, 1315, 1266, 1095, 791 $cm^{-1}$. $^{1}H$-NMR (400 MHz, $CDCl_3$) δ: 0.85 [dd, *J* = 11.6 Hz, *J'* = 1.6 Hz, 2H, 5(7)-$H_a$], 1.00 [s, 6H, 3(9)-$CH_3$ or 4(8)-$CH_3$], 1.02 [s, 6H, 4(8)-$CH_3$ or 3(9)-$CH_3$], 1.05 [d, *J* = 10.5 Hz, 2H, 2(10)-$H_a$], 1.90 [d, *J* = 12.4 Hz, 5(7)-$H_b$], 2.08 [d, *J* = 10.8 Hz, 2H, 2(10)-$H_b$], 2.62 (broad s., 1H, 6-H), 6.91 (d, *J* = 5.2 Hz, 1H, NH), 6.98 (d, *J* = 7.6 Hz, 1H, 8'-H), 7.35 (dd, *J* = 9.2 Hz, *J'* = 4.4 Hz, 1H, 3'-H), 7.41 (d, *J* = 5.2 Hz, 1H, NH), 7.58 (t, *J* = 8 Hz, 7'-H), 7.70 (d, *J* = 9.6 Hz, 1H, 6'-H), 8.30 (d, *J* = 9.2 Hz, 1H, 4'-H), 8.90 (d, *J* = 4.4 Hz, 1H, 2'-H). $^{13}C$-NMR (100.6 MHz, $CDCl_3$) δ: 15.7 [$CH_3$, C3(9)-$\underline{C}H_3$ or C4(8)-$\underline{C}H_3$], 15.8 [$CH_3$, C4(8)-$\underline{C}H_3$ or C3(9)-$\underline{C}H_3$], 37.9 (CH, C6), 38.4 [$CH_2$, C5(7)], 41.4 [$CH_2$, C2(10)], 45.3 [C, C3(9) or C4(8)], 45.8 [C, C4(8) or C3(9)], 48.6 (C, C1), 107.9 (CH, C8'), 119.7 (C, C4a'), 120.2 (CH, C3'), 123.0 (CH, C6'), 129.1 (CH, C4'), 129.7 (CH, C7'), 143.3 (C, C5'), 149.0 (C, C8a'), 150.4 (CH, C2'), 177.2 (C, CO). HRMS-ESI+ *m/z* $[M+H]^+$ calcd for $[C_{24}H_{30}N_3O]^+$: 376.2383, found: 376.2378.

Elemental analysis: Calcd for $C_{24}H_{29}N_3O \cdot 0.75H_2O$: C 74.10, H 7.90, N 10.80. Found: C 74.06, H 7.72, N 10.68. HPLC-UV purity at 254 nm = 98.58%.

***N'*-(6-Chloroquinolin-5-yl)-3,4,8,9-tetramethyltetracyclo[4.4.0.0$^{3,9}$.0$^{4,8}$]decane-1-carbohydrazide (ALT-P21).** 3,4,8,9-Tetramethyltetracyclo[4.4.0.0$^{3,9}$.0$^{4,8}$]decane-1-carbonyl chloride (171 mg, 0.68 mmol) was obtained following the general procedure **A** and, without further purification, was reacted with 6-chloro-5-hydrazineylquinoline dihydrochloride (198 mg, 0.74 mmol) in the presence of anh. triethylamine (188 µL, 1.36 mmol), following the general procedure **B**. The formed precipitate was filtered off and the organic layer was concentrated under vacuum. Purification by Combiflash® in silica gel using as eluent hexane to ethyl acetate/hexane mixture (4/6) gave the compound as a yellow solid (80 mg, 29% yield), mp 166 ˚C. IR (ATR) ν: 3448, 3310, 3215, 2950, 2862, 1667, 1648, 1588, 1574, 1490, 1465, 1450, 1385, 1371, 1362, 1312, 1286, 1182, 1130, 1123, 1086, 937, 918, 867, 846, 829, 813, 803, 778, 745, 685, 653, 633 cm$^{-1}$. $^{1}$H-NMR (400 MHz, $CDCl_3$) δ: 0.74 [dd, *J* = 11.6 Hz, *J'* = 2.8 Hz, 2H, 5(7)-H$_a$], 0.85 [d, *J* = 11.2 Hz, 2H, 2(10)-H$_a$], 0.93 [s, 12H, 3(4,8,9)-$CH_3$], 1.78 [dd, *J* = 11.6 Hz, *J'* = 1.2 Hz, 2H, 5(7)-H$_b$], 1.92 [d, *J* = 10.8 Hz, 2(10)-H$_b$], 2.43 (m, 1H, 6-H), 6.82 (d, *J* = 3.6 Hz, 1H, NH), 7.44 (dd, *J* = 8.8 Hz, *J'* = 4.4 Hz, 1H, 3'-H), 7.64 (d, *J* = 9.2 Hz, 7'-H), 7.77 (d, *J* = 3.6 Hz, 1H, NH), 7.81 (d, *J* = 9.2 Hz, 1H, 8'-H), 8.71 (d, *J* = 8.4 Hz, 1H, 4'-H), 8.89 (dd, *J* = 4 Hz, *J'* = 1.6 Hz, 1H, 2'-H). $^{13}$C-NMR (100.6 MHz, $CDCl_3$) δ: 15.6 [$CH_3$, C3(9)-$\underline{C}H_3$ or C4(8)-$\underline{C}H_3$], 15.7 [$CH_3$, C4(8)-$\underline{C}H_3$ or C3(9)-$\underline{C}H_3$], 37.7 (CH, C6), 38.2 [$CH_2$, C5(7)], 41.1 [$CH_2$, C2(10)], 45.1 [C, C3(9) or C4(8)], 45.7 [C, C3(9) or C4(8)], 48.5 (C, C1), 121.2 (CH, C3'), 123.0 (C, d, *J* = 8 Hz, C4a'), 126.7 (CH, C8'), 130.8 (CH, C7'), 131.4 (CH, C4'), 139.4 (C, C5'), 147.7 (C, C8a'), 150.5 (CH, C2'), 176.9 (C, CO). HRMS-ESI+ *m/z* $[M+H]^+$ calcd for $[C_{24}H_{29}ClN_3O]^+$: 410.1994, found: 410.1985. HPLC-UV purity at 254 nm = 96.19%.

**General procedure C for the synthesis of amides**

To a solution of the required amine (0.34 mmol, 1 eq) in DCM (2 mL) is added triethylamine (1.36 mmol, 4 eq) and acyl chloride (0.34 mmol, 1 eq), obtained using General procedure A. The resulting mixture is stirred for 2 h at room temperature and successively washed with water (2 x 3 mL), saturated aq. $NaHCO_3$ solution (2 x 2mL) and brine (2 mL). The organic layer is dried over anh. $Na_2SO_4$, filtered and concentrated under vacuum. The purification procedure is specified below for each compound.

**3,4,8,9-Tetramethyl-*N*-(quinolin-5-yl)tetracyclo[4.4.0.0$^{3,9}$.0$^{4,8}$]decane-1-carboxamide (ALT-P19).** 3,4,8,9-Tetramethyltetracyclo[4.4.0.0$^{3,9}$.0$^{4,8}$]decane-1-carbonyl chloride (86 mg, 0.34 mmol) was obtained following the general procedure **A**

and, without further purification was reacted with 5-aminoquinoline (49 mg, 0.34 mmol) in the presence of anh. triethylamine (189 μL, 1.36 mmol), following the general procedure **C**. Purification by Combiflash® in silica gel using as eluent hexane to ethyl acetate/hexane mixture (4/6) gave the title compound as a white solid (83 mg, 0.23 mmol, 68% yield). An analytical sample was obtained by crystallization from MeOH/$Et_2O$, 250 ˚C. IR (NaCl disk) ν: 3312, 2957, 2926, 2865, 1645, 1615, 1593, 1570, 1559, 1482, 1457, 1387, 1373, 1305, 1271, 1204, 1161, 1143, 1108, 806 $cm^{-1}$. $^1$H-NMR (400 MHz, $CDCl_3$) δ: 0.88 [dd, *J* = 11.2 Hz, *J'* = 2.4 Hz, 2H, 5(7)-$H_a$], 1.02 [s, 6H, 3(9)-$CH_3$ or 4(8)-$CH_3$], 1.04 [s, 6H, 4(8)-$CH_3$ or 3(9)-$CH_3$], 1.14 [d, *J* = 10.8 Hz, 2H, 2(10)-$H_a$], 1.96 [dd, *J* = 12 Hz, *J'* = 1.2 Hz, 5(7)-$H_b$], 2.14 [d, *J* = 10.8 Hz, 2H, 2(10)-$H_b$], 2.76 (broad s., 1H, 6-H), 7.44 (dd, *J* = 8.8 Hz, *J'* = 4.4 Hz, 1H, 3'-H), 7.64 (broad s., 1H, NH), 7.71 (t, *J* = 8 Hz, 7'-H), 7.91 (d, *J* = 8 Hz, 1H, 8'-H), 7.97 (d, *J* = 8.4 Hz, 1H, 6'-H), 8.13 (d, *J* = 8.4 Hz, 1H, 4'-H), 8.93 (dd, *J* = 4 Hz, *J'* = 1.6 Hz, 1H, 2'-H). $^{13}$C-NMR (100.6 MHz, $CDCl_3$) δ: 15.7 [$CH_3$, C3(9)-$\underline{C}H_3$ or C4(8)-$\underline{C}H_3$], 15.8 [$CH_3$, C4(8)-$\underline{C}H_3$ or C3(9)-$\underline{C}H_3$], 38.1 (CH, C6), 38.5 [$CH_2$, C5(7)], 41.9 [$CH_2$, C2(10)], 45.2 [C, C3(9) or C4(8)], 46.0 [C, C4(8) or C3(9)], 50.6 (C, C1), 121.1 (CH, C3'), 121.8 (CH, C8'), 123.1 (C, C4a'), 127.4 (CH, C6'), 129.5 (CH, C7'), 129.8 (CH, C4'), 132.7 (C, C5') 148.8 (C, C8a'), 150.5 (CH, C2'), 176.1 (C, CO). HRMS-ESI+ *m/z* $[M+H]^+$ calcd for $[C_{24}H_{29}N_2O]^+$: 361.2274, found: 361.2272. Elemental analysis: Calcd for $C_{24}H_{28}N_2O$: C 79.96, H 7.83, N 7.77. Found: C 79.94, H 7.89, N 7.98.

**3,4,8,9-Tetramethyl-*N*-(quinolin-5-ylmethyl)tetracyclo[4.4.0.0$^{3,9}$.0$^{4,8}$]decane-1-carboxamide (ALT-P20).** 3,4,8,9-Tetramethyltetracyclo[4.4.0.0$^{3,9}$.0$^{4,8}$]decane-1-carbonyl chloride (75 mg, 0.30 mmol) was obtained following the general procedure **A** and, without further purification was reacted with quinolin-5-ylmethanamine hydrochloride (58 mg, 0.30 mmol) in the presence of anh. triethylamine (169 μL, 1.20 mmol), following the general procedure **C**. Purification by Combiflash® in silica gel using as eluent hexane to ethyl acetate/hexane mixture (3.5/6.5) gave the desired compound as a white solid (45 mg, 0.12 mmol, 40% yield), mp 185 ˚C. IR (NaCl disk) ν: 3332, 2948, 2864, 1639, 1598, 1573, 1528, 1502, 1479, 1454, 1412, 1385, 1371, 1310, 1283, 1224, 1224, 1154, 1113, 1070, 1039, 802, 734, 706 $cm^{-1}$. $^1$H-NMR (400 MHz, $CDCl_3$) δ: 0.75 [dd, *J* = 11.2 Hz, *J'* = 2.4 Hz, 2H, 5(7)-$H_a$], 0.84 [d, *J* = 10.8 Hz, 2H, 2(10)-$H_a$], 0.92 [s, 6H, 3(9)-$CH_3$ or 4(8)-$CH_3$], 0.93 [s, 6H, 4(8)-$CH_3$ or 3(9)-$CH_3$], 1.79 [dd, *J* = 12 Hz, *J'* = 1.2 Hz, 2H, 5(7)-$H_b$], 1.91 [d, *J* = 10.8 Hz, 2H, 2(10)-$H_b$], 2.53 (m, 1H, 6-H), 4.92 (d, *J* = 5.2 Hz, 2H, $NHCH_2$), 5.81 (broad s, 1H, NH), 7.45 (dd, *J* = 8.4 Hz, *J'* = 4.4 Hz, 1H, 3'-H), 7.49 (d, *J* = 6.9 Hz, 1 H, 8'-H), 7.66 (m, 7'-H), 8.07 (d, *J* = 8.6 Hz, 1H, 6'-H), 8.43 (d, *J* = 8.6 Hz, 1H, 4'-H), 8.94 (dd, *J* = 4.2 Hz, *J'* = 1.7 Hz, 1H, 2'-H). $^{13}$C-NMR (100.6 MHz,

$CDCl_3$) δ: 15.66 [$CH_3$, C3(9)-$\underline{C}H_3$ or C4(8)-$\underline{C}H_3$], 15.74 [$CH_3$, C4(8)-$\underline{C}H_3$ or C3(9)-$\underline{C}H_3$], 37.9 (CH, C6), 38.5 [$CH_2$, C5(7)], 41.0 ($CH_2$, NH$\underline{C}H_2$), 41.6 [$CH_2$, C2(10)], 45.1 [C, C3(9) or C4(8)], 45.8 [C, C4(8) or C3(9)], 49.6 (C, C1), 121.6 (CH, C3'), 126.9 (C, C4a'), 127.1 (CH, C8'), 129.0 (CH, C7'), 130.1 (CH, C6'), 132.4 (CH, C4'), 134.9 (C) and 149.0 (C), (C5 and C8a'), 150.6 (CH, C2'), 176.9 (C, CO). HRMS-ESI+ *m/z* $[M+H]^+$ calcd for $[C_{25}H_{31}N_2O]^+$: 375.2431, found: 375.2429. Elemental analysis: calcd for $C_{25}H_{30}N_2O$: C 80.17, H 8.07, N 7.48. Found: C 80.18, H 8.03, N 7.46.

***N*-((3,4,8,9-Tetramethyltetracyclo[4.4.0.0$^{3,9}$.0$^{4,8}$]decan-1-yl)methyl)quinoline-5-carboxamide (ALT-P10).** The quinoline-5-carbonyl chloride hydrochloride (52 mg, 0.23 mmol) was obtained following the general procedure **A** and, without further purification, was reacted with (3,4,8,9-tetramethyltetracyclo[4.4.0.0$^{3,9}$.0$^{4,8}$]decan-1-yl)methanamine (51 mg, 0.23 mmol) in the presence of anh. triethylamine (128 µL, 0.92 mmol), following the general procedure **C**. The crude was crystallized from DCM/pentane yielding the title compound as a white solid (40 mg, 0.11 mmol, 48% yield), mp 173-174 ˚C. IR (NaCl disk) ν: 3225, 2942, 2908, 2860, 1663, 1647, 1624, 1595, 1570, 1558, 1534, 1496, 1466, 1453, 1408, 1368, 1320, 1269, 1220, 1189, 1146, 1108, 1062, 973, 837, 827, 795, $cm^{-1}$. $^1$H-NMR (400 MHz, $CDCl_3$) δ: 0.62 [d, *J* = 10.8 Hz, 2H, 2'(10')-$H_a$], 0.74 [d, *J* = 11.2 Hz, 2H, 5'(7')-$H_a$], 0.96 [s, 12H, 3'(9')-$CH_3$ and 4'(8')-$CH_3$], 1.71 [d, *J* = 11.2 Hz, 2H, 2'(10')-$H_b$], 1.77 [d, *J* = 11.2 Hz, 2H, 5'(7')-$H_b$], 2.19 (broad s, 1H, 6'-H), 3.61 (d, *J* = 6 Hz, 2H, $CH_2$NHCO), 6.11 (broad s, 1H, NH), 7.48 (dd, *J* = 8.8 Hz, *J'* = 4.4 Hz, 1H, 3-H), 7.67-7.73 [cs, 2H, 7(8)-H], 8.19 (d, *J* = 8 Hz, 1H, 6-H), 8.76 (d, *J* = 8.4 Hz, 1H, 4-H), 8.96 (dd, *J* = 4.2 Hz, *J'* = 1.7 Hz, 1H, 2-H). $^{13}$C-NMR (100.6 MHz, $CDCl_3$) δ: 15.7 [$CH_3$, C3'(9')-$\underline{C}H_3$ or C4'(8')-$\underline{C}H_3$], 16.0 [$CH_3$, C4'(8')-$\underline{C}H_3$ or C3'(9')-$\underline{C}H_3$], 37.5 (CH, C6'), 38.8 [$CH_2$, C2'(10')], 41.8 [$CH_2$, C5'(7')], 43.4 [C, C3'(9') or C4'(8')], 45.0 [C, C4'(8') or C3'(9')], 46.0 (C, C1'), 47.4 ($CH_2$, $\underline{C}H_2$NHCO), 122.1 (CH, C3), 125.1 [CH, C7(8)], 126.0 (C, C4a), 128.4 [CH, C8(7)], 132.3 (CH, C6), 134.4 (CH, C4), 135.2 (C) and 148.4 (C) (C5 and C8a), 151.1 (CH, C2), 168.7 (C, CO). HRMS-ESI+ *m/z* $[M+H]^+$ calcd for $[C_{25}H_{31}N_2O]^+$: 375.2431, found: 375.2426. Elemental analysis: Calcd for $C_{25}H_{30}N_2O·0.4H_2O$: C 78.66, H 8.13, N 7.34. Found: C 79.09, H 8.13, N 6.91. HPLC-UV purity at 254 nm = 98.73%.

***N*-(quinolin-5-yl)-2-(3,4,8,9-tetramethyltetracyclo[4.4.0.0$^{3,9}$.0$^{4,8}$]decan-1-yl)acetamide (ALT-P25).** 2-(3,4,8,9-Tetramethyltetracyclo[4.4.0.0$^{3,9}$.0$^{4,8}$]decan-1-yl)acetyl chloride (103 mg, 0.39 mmol) was obtained from 2-(3,4,8,9-tetramethyltetracyclo[4.4.0.0$^{3,9}$.0$^{4,8}$]decan-1-yl)acetic acid (see below for its synthetic details) following the general procedure **A** and, without further purification, was reacted with 5-aminoquinoline (56 mg, 0.39 mmol) in the presence of anh. triethylamine (216 µL, 1.56 mmol), following the general procedure **C**. Purification by Combiflash® in silica gel using as eluent hexane to ethyl acetate/hexane mixture (5/5) gave the title compound as a white solid (60 mg, 41% yield), mp 230 ˚C. IR (ATR) ν: 3222, 2959, 2941, 2911, 2860, 1668, 1645, 1624, 1596, 1538, 1496, 1466, 1409, 1367, 1321, 1269, 1236, 1222, 1198, 1147, 1063, 973, 842, 826, 794, 760, 746, 681 $cm^{-1}$. $^1$H-NMR (400 MHz, $CDCl_3$) δ: 0.71 [d, $J$ = 10.8 Hz, 2H, 2'(10')-$H_a$], 0.74 [d, $J$ = 11.2 Hz, 2H, 5'(7')-$H_a$], 0.95 [s, 12H, 3'(9')-$CH_3$ and 4'(8')-$CH_3$], 1.83 [d, $J$ = 11.6 Hz, 2H, 5'(7')-$H_b$], 1.91 [d, $J$ = 11.2 Hz, 2H, 2'(10')-$H_b$], 2.36 (broad s., 1H, 6'-H), 2.65 (s, 2H, $CH_2$), 7.44 (dd, $J$ = 8.8 Hz, $J'$ = 4.4 Hz, 1H, 3-H), 7.49 (broad s, 1H, NH), 7.71 (t, $J$ = 8 Hz, 1H, 7-H), 7.86 (d, $J$ = 7.6 Hz, 1H, 8-H), 7.99 (d, $J$ = 8.4 Hz, 1H, 6-H), 8.23 (d, $J$ = 8.4 Hz, 1H, 4-H), 8.93 (d, $J$ = 2.8 Hz, 1H, 2-H). $^{13}$C-NMR (100.6 MHz, $CDCl_3$) δ: 15.7 [$CH_3$, C3'(9')-$\underline{C}H_3$ or C4'(8')-$\underline{C}H_3$], 15.9 [$CH_3$, C4'(8')-$\underline{C}H_3$ or C3'(9')-$\underline{C}H_3$], 38.8 [$CH_2$, C5'(7')], 39.3 (CH, C6'), 41.2 (C, C1'), 44.3 [$CH_2$, C2'(10')], 44.8 [C, C3'(9') or C4'(8')], 46.1 [C, C3'(9') or C4'(8')], 47.0 ($CH_2$, N$\underline{C}H_2$), 121.2 (CH, C3), 122.2 (CH, C8), 123.2 (C, C4a), 127.6 (CH, C6), 129.5 (CH, C7), 130.4 (CH, C4), 132.8 (C, C5) 148.7 (C, C8a), 150.4 (CH, C2), 171.2 (C, CO).HRMS-ESI+ *m /z* $[M+H]^+$ calcd for $[C_{25}H_{31}N_2O]^+$: 375.2431, found: 375.2435. Elemental analysis: calcd for $C_{25}H_{30}N_2O·0.35H_2O$: C 78.85, H 8.13, N 7.36. Found: C 78.79, H 7.90, N 7.24. HPLC-UV purity at 220 nm = 98.57%.

**General procedure D for the synthesis of ureas**

Triphosgene (0.43 mmol, 0.5 eq) is added in a single portion to a solution of the required amine (0.86 mmol, 1 eq) in DCM (7 mL) and saturated aq. $NaHCO_3$ solution (4 mL). The resulting biphasic mixture was stirred at room temperature for 30 min. Then, the two phases were separated, and the organic layer was washed with brine, dried over anh. $Na_2SO_4$ and filtered. Evaporation under vacuum provides the isocyanate intermediate which was used in the next step without further purification.

Next, 5-aminoquinoline (1.21 mmol, 1.2 eq) is dissolved in anhydrous THF (12 mL) under argon and cooled to -60 °C on a dry ice in acetone bath. Afterwards *n*-butyllithium (1.21 mmol, 2.5 M in hexane, 1 eq) is added dropwise. The reaction mixture was then removed from the dry ice in acetone bath and tempered to 0 °C with an ice bath. The isocyanate (1.01 mmol, 1 eq) is added and then the mixture is stirred at room temperature overnight followed by the methanol addition to quench any reacted *n*-butyllithium. Purification by column chromatography provided the desired urea.

**1-(Quinolin-5-yl)-3-(3,4,8,9-tetramethyltetracyclo[4.4.0.0$^{3,9}$.0$^{4,8}$]decan-1-yl)urea (ALT-P22).** 3,4,8,9-Tetramethyltetracyclo[4.4.0.0$^{3,9}$.0$^{4,8}$]decan-1-amine hydrochloride (208 mg, 0.86 mmol) reacted with triphosgene (127 mg, 0.43 mmol) following general procedure **D** to give the corresponding isocyanate intermediate. Next, 5-aminoquinoline

(148 mg, 1.03 mmol) and *n*-butyllithium (344 µL, 2.5 M in hexane, 0.86 mmol) in THF were reacted with the isocyanate to yield a green solid. Purification by Combiflash® in silica gel using as eluent hexane to ethyl acetate/hexane mixture (3.5/6.5) followed by concentration under vacuum gave the title compound as a white solid (72 mg, 22% yield), mp 260 ˚C. IR (ATR) ν: 3287, 2945, 2862, 1668, 1633, 1597, 1597, 1538, 1478, 1455, 1422, 1361, 1308, 1291, 1243, 1226, 1094, 1072, 796 $cm^{-1}$. $^1$H-NMR (400 MHz, $CD_3OD$) δ: 0.81 [dd, *J* = 11.2 Hz, *J'* = 3.2 Hz, 2H, 5'(7')-$H_a$], 0.89 [d, *J* = 10.8 Hz, 2H, 2'(10')-$H_a$], 0.99 [s, 6H, 3'(9')-$CH_3$ or 4'(8')-$CH_3$], 1.00 [s, 6H, 4'(8')-$CH_3$ or 3'(9')-$CH_3$], 2.03 [dd, *J* = 12 Hz, *J'* = 1.2 Hz, 2H, 5'(7')-$H_b$], 2.23 [d, *J* = 10.8 Hz, 2H, 2'(10')-$H_b$], 2.23 (broad s, 1H, 6'-H), 7.55 (dd, *J* = 8.8 Hz, *J'* = 4.4 Hz, 1H, 3-H), 7.73 (dd, *J* = 8.4 Hz, *J'* = 4.4 Hz, 1H, 7-H), 7.79 (dt, *J* = 8.4 Hz, *J'* = 1.2 Hz, 1H, 6-H), 7.86 (dd, *J* = 7.6 Hz, *J'* = 1.2 Hz, 1H, 8-H), 8.50 (ddd, *J* = 8.4 Hz, *J'* = 1.6 Hz, *J''* = 0.8 Hz, 1H, 4-H), 8.84 (dd, *J* = 4.4 Hz, *J'* = 1.6 Hz, 1H, 2-H). $^{13}$C-NMR (100.6 MHz, $CD_3OD$) δ: 15.8 [$CH_3$, C3'(9')-$\underline{C}H_3$ or C4'(8')-$\underline{C}H_3$], 16.0 [$CH_3$, C4'(8')-$\underline{C}H_3$ or C3'(9')-$\underline{C}H_3$], 39.8 [$CH_2$, C5'(7')], 44.0 (CH, C6'), 44.3 [$CH_2$, C2'(10')], 45.7 [C, C3'(9') or C4'(8')], 45.9 [C, C4'(8') or C3'(9')], 58.4 (C, C1'), 120.9 (CH, C8), 121.9 (CH, C3), 124.3 (C, C4a), 124.7 (CH, C6), 131.0 (CH, C7), 132.8 (CH, C4), 136.4 (C, C5), 149.3 (C, C8a), 151.0 (CH, C2), 158.0 (C, CO). HRMS-ESI+ *m/z* $[M+H]^+$ calcd for $[C_{24}H_{30}N_3O]^+$: 376.2383, found: 376.2377. HPLC-UV purity at 254 nm = 98.53%.

**1-(Quinolin-5-yl)-3-((3,4,8,9-tetramethyltetracyclo[4.4.0.0$^{3,9}$.0$^{4,8}$]decan-1-yl)methyl)urea hydrochloride (ALT-P16).** (3,4,8,9-Tetramethyltetracyclo[4.4.0.0$^{3,9}$.0$^{4,8}$]decan-1-yl)methanamine hydrochloride (100 mg, 0.39 mmol) reacted with triphosgene (58 mg, 0.20 mmol) following general procedure **D** to give the corresponding isocyanate intermediate. Next, 5-aminoquinoline (67 mg, 0.47 mmol), and *n*-butyllithium (156 µL, 2.5 M in hexane, 0.39 mmol), in THF, were reacted with the isocyanate to yield a green solid. The crude was purified by Combiflash® in silica gel using as eluent hexane to ethyl acetate/hexane mixture (2.5/7.5). Its hydrochloride was obtained by adding an excess of HCl/$Et_2O$ to a solution of the amine in dichlorometane, followed by filtration of the resulting yellow precipitate (26 mg, 16% yield), mp 199 ˚C. IR (ATR) ν: 3290, 2944, 2862, 1668, 1651, 1590, 1538, 1455, 1422, 1384, 1361, 1309, 1291, 1226, 1163, 1116, 1072, 801 $cm^{-1}$. $^1$H-NMR (400 MHz, $CD_3OD$) δ: 0.64 [d, *J* = 10.8 Hz, 2H, 2'(10')-$H_a$], 0.75 [dd, *J* = 11.2 Hz, *J'* = 1.6 Hz, 2H, 5'(7')-$H_a$], 0.99 [s, 12H, 3'(4',8',9')-$CH_3$], 1.74 [d, *J* = 10.8 Hz, 2H, 2'(10')-$H_b$], 1.86 [d, *J* = 11.2 Hz, 2H, 5'(7')-$H_b$], 2.13 (broad s, 1H, 6'-H), 3.35 (s, 2H, $CH_2$NHCO), 7.88 (d, *J* = 8.8 Hz, 1H, 6-H), 8.06 (dd, *J* = 8.8 Hz, *J'* = 5.2 Hz, 1H, 3-H), 8.10 (t, *J* = 8 Hz, 1H, 7-H), 8.31 (d, *J* = 8 Hz, 1H, 8-H), 9.18 (d, *J* = 4 Hz, 1H, 2-H), 9.40 (m, 1H, 4-H). $^{13}$C-NMR (100.6 MHz,

$CD_3OD$) δ: 15.8 [$CH_3$, C3'(9')-$\underline{C}H_3$ or C4'(8')-$\underline{C}H_3$], 16.0 [$CH_3$, C4'(8')-$\underline{C}H_3$ or C3'(9')-$\underline{C}H_3$], 38.6 (CH, C6'), 39.7 [$CH_2$, C5'(7')], 42.6 [$CH_2$, C2'(10')], 44.5 [C, C3'(9') or C4'(8')], 46.0 [C, C4'(8') or C3'(9')], 47.0 (C, C1'), 48.3 ($CH_2$, $\underline{C}H_2$NHCO), 115.5 (CH, C6), 121.7 [CH, C3(8)], 124.7 (C, C4a), 136.9 (CH, C7), 138.8 (C, C5), 140.0 (C, C8a), 143.6 (CH) and 145.8 (CH) (C2 and C4), 158.1 (C, CO). HRMS-ESI+ *m/z* $[M+H]^+$ calcd for $[C_{25}H_{32}N_3O]^+$: 390.254, found: 390.2543. Elemental analysis: calcd for $C_{25}H_{31}N_3O \cdot HCl \cdot 2H_2O$: 64.99, H 7.85, N 9.09. Found: C 65.29, H 7.53, N 9.09. HPLC-UV purity at 254 nm = 97.70%.

**General procedure E for the synthesis of thioureas**

a) Synthesis of 5-isothiocyanatoquinoline. 1,1'-thiocarbonyldiimidazole (741 mg, 4.16 mmol) was added to a solution of 5-aminoquinoline (600 mg, 4.16 mmol) in DCM (3 mL) at 0 °C. The mixture was allowed to stir at room temperature for 2 h before the solvent was removed under vacuum. Purification by Combiflash® in silica gel using as eluent hexane to ethyl acetate/ hexane mixture (0.5/9.5) yielded a yellow solid (500 mg, 2.69 mmol, 65% yield) that was used in the next step. The spectroscopic data were identical to those previously published.[1]

b) 5-isothiocyanatoquinoline (0.64 mmol, 1 eq), synthesized as previously published, is added to a solution of the required amine (0.64 mmol, 1eq) in DCM (7 mL) and triethylamine (1.28 mmol, 2eq). The reaction is allowed to stir for 18 h at room temperature. After that, water (5 mL) is added and the phases are separated. The aq. layer is extracted with further DCM (10 mL) and the combined organic extracts are dried over anh. $Na_2SO_4$, filtered and concentrated under vacuum. The purification procedure is specified in each compound.

**1-(quinolin-5-yl)-3-(3,4,8,9-tetramethyltetracyclo[4.4.0.0$^{3,9}$.0$^{4,8}$]decan-1-yl)thiourea (ALT-P13).** From 3,4,8,9-tetramethyltetracyclo[4.4.0.0$^{3,9}$.0$^{4,8}$]decan-1-amine hydrochloride (120 mg, 0.50 mmol), 5-isothiocyanatoquinoline (93 mg, 0.50 mmol) and triethylamine (139 µL, 1.00 mmol) and following the general procedure **E**, a crude was obtained. Crystallization from DCM/pentane afforded the title compound as a white solid (130 mg, 66% yield), mp 212 °C. IR (ATR) ν: 3000-2800 (2969, 2943, 2916, 2863), 1650, 1587, 1574, 1532, 1519, 1491, 1456, 1405, 1370, 1334, 1266, 1213, 1187, 894, 817, 799, 751, 686, 667, 638 $cm^{-1}$. $^1$H-NMR (400 MHz, $CDCl_3$) δ: 0.62 [broad s., 2H, 2'(10')-$H_a$], 0.89-0.92 [complex signal, 14H, 3'(9')-$CH_3$ or 4'(8')-$CH_3$ and 5'(7')-$H_a$], 1.96 [d, *J* =

[1] Betschmann, P.; Carroll, W. A.; Ericsson, A. M.; Fix-Stenzel, S. R.; Friedman, M.; Hirst, G. C.; Josephsohn, N. S.; Pérez-Medrano, A.; Morytko, M. J.; Rafferty, P.; Chen, H. Piperazines as P2X7 antagonists. WO 2008/005368A2, Abbott Laboratories (Ex. 106).

11.6 Hz, 2H, 5'(7')-$H_b$], 2.29 [d, *J* = 10.8 Hz, 2H, 2'(10')-$H_b$], 2.79 (broad s., 1H, 6'-H), 6.11 (broad s., 1H, NH), 7.49-7.54 [complex signal, 2H, 3(8)-H], 7.75 (dd, *J* = 8.4 Hz, *J'* = 7.6 Hz, 7-H), 7.84 (broad s., 1H, NH), 8.13 (d, *J* = 8.6 Hz, 1H, 6-H), 8.36 (d, *J* = 8.5 Hz, 1H, 4-H), 8.84 (dd, *J* = 4.4 Hz, *J'* = 1.6 Hz, 1H, 2-H). $^{13}$C-NMR (100.6 MHz, $CDCl_3$) δ: 15.5 [$CH_3$, C3'(9')-<u>C</u>$H_3$ or C4'(8')-<u>C</u>$H_3$], 15.6 [$CH_3$, C4'(8')-<u>C</u>$H_3$ or C3'(9')-<u>C</u>$H_3$], 39.1 [$CH_2$, C5'(7')], 41.3 (CH, C6'), 43.4 [$CH_2$, C2'(10')], 44.5 [C, C3'(9') or C4'(8')], 45.1 [C, C4'(8') or C3'(')], 60.7 (C, C1'), 122.2 (CH, C8), 125.5 [CH, C3(4a)], 129.4 (CH, C7), 130.2 (CH, C6), 131.5 [CH, C4(5)], 149.2 (C, C8a), 151.5 (CH, C2), 181.0 (C, CS). HRMS-ESI+ *m/z* $[M+H]^+$ calcd for $[C_{24}H_{30}N_3S]^+$: 392.2155, found: 392.2158. Elemental analysis: calcd for $C_{24}H_{29}N_3S\cdot0.25H_2O$: 72.78, H 7.51, N 10.61. Found: C 73.11, H 7.37, N 10.39. HPLC-UV purity at 254 nm = 97.16%.

**1-(quinolin-5-yl)-3-((3,4,8,9-tetramethyltetracyclo[4.4.0.0$^{3,9}$.0$^{4,8}$]decan-1-yl)methyl)thiourea (ALT-P14).** From (3,4,8,9-tetramethyltetracyclo[4.4.0.0$^{3,9}$.0$^{4,8}$]decan-1-yl)methanamine hydrochloride (61 mg, 0.24 mmol), 5-isothiocyanatoquinoline (44 mg, 0.24 mmol) and triethylamine (70 µL, 0.48 mmol) and following the general procedure for the synthesis of thiourea, a crude was obtained. Crystallization from DCM/pentane afforded a white solid (87 mg, 89% yield), mp: 220 ˚C. IR (ATR) ν: 3126, 3000-2800 (2965, 2940, 2909, 2859), 1651, 1614, 1597, 1575, 1544, 1524, 1450, 1454, 1321, 1267, 1245, 1226, 1190, 1108, 1096, 1058, 1034, 1009, 891, 865, 800, 760, 739, 687, 675 cm$^{-1}$. $^1$H-NMR (400 MHz, $CDCl_3$) δ: 0.30 [d, *J* = 11.2 Hz, 2H, 2'(10')-$H_a$], 0.48 [d, *J* = 12.8 Hz, 2H, 5'(7')-$H_a$], 0.81 [s, 6H, 3'(9')-C<u>H</u>$_3$ or 4'(8')-C<u>H</u>$_3$], 0.84 [s, 6H, 4'(8')-C<u>H</u>$_3$ or 3'(9')-C<u>H</u>$_3$], 1.32 [d, *J* = 11.2 Hz, 4H, 2'(5',7',10')-$H_b$], 1.80 (broad s., 1H, 6'-H), 3.63 (s, 2H, C<u>H</u>$_2$NHCS), 5.68 (s, 1H, NH), 7.51 (dd, *J* = 8.4 Hz, *J'* = 4 Hz, 1H, 3-H), 7.55 (d, *J* = 7.2 Hz, 1H, 8-H), 7.79 (dd, *J* = 8.7 Hz, *J'* = 7.2 Hz, 1H, 7-H), 7.92 (s, 1H, NH), 8.18 (d, *J* = 8.4 Hz, 1H, 6-H), 8.41 (d, *J* = 8.4 Hz, 1H, 4-H), 9.00 (dd, *J* = 4.4 Hz, *J'* = 1.6 Hz, 1H, 2-H). $^{13}$C-NMR (100.6 MHz, $CDCl_3$) δ: 15.6 [$CH_3$, C3'(9')-<u>C</u>$H_3$ or C4'(8')-<u>C</u>$H_3$], 15.8 [$CH_3$, C4'(8')-<u>C</u>$H_3$ or C3'(9')-<u>C</u>$H_3$], 37.0 (CH, C6'), 38.2 [$CH_2$, C5'(7')], 41.6 [$CH_2$, C2'(10')], 42.9 [C, C3'(9') or C4'(8')], 44.8 [C, C4'(8') or C3'(9')], 45.7 (C, C1'), 53.2 ($CH_2$, <u>C</u>$H_2$NHCS), 122.5 (CH, C3), 125.7 (C, C4a), 126.0 (CH, C8), 129.4 (CH, C7), 130.8 (C, C6), 131.5 (CH, C4), 131.7 (C, C5), 149.3 (C, C8a), 151.7 (CH, C2), 182.2 (C, CS). HRMS-ESI+ *m/z* $[M+H]^+$ calcd for $[C_{25}H_{32}N_3S]^+$: 406.2311, found: 406.2309. Elemental analysis: calcd for $C_{25}H_{31}N_3S\cdot0.25H_2O$: 73.38, H 7.73, N 10.27. Found: C 73.15, H 7.48, N 10.15.

**2-cyano-1-(quinolin-5-yl)-3-((3,4,8,9-tetramethyltetracyclo[4.4.0.0$^{3,9}$.0$^{4,8}$]decan-1-yl)methyl)guanidine (ALT-P7).** (3,4,8,9-Tetramethyltetracyclo[4.4.0.0$^{3,9}$.0$^{4,8}$]decan-1-yl)methanamine hydrochloride (83 mg, 0.32 mmol) and phenyl-*N'*-cyano-*N*-(quinolin-5-

yl)carbamimidate (70 mg, 0.24 mmol) were suspended in 2-propanol (6 mL) and heated at reflux overnight. The reaction was cooled down and kept at 0 °C for 12 h. The precipitate formed was filtered off under vacuum yielding a yellow solid (50 mg, 50% yield), mp 259 ˚C. IR (ATR) v: 2965, 2942, 2861, 2175, 1650, 1589, 1566, 1451, 1426, 1373, 1357, 1340, 1310, 1243, 1129, 1084, 1058, 1084, 1058, 918, 828, 804, 774, 669, 652 $cm^{-1}$. $^1$H-NMR (400 MHz, $CDCl_3$) δ: 0.47 [d, *J* = 10.8 Hz, 2H, 2'(10')-$H_a$], 0.61 [d, *J* = 11.2 Hz, 2H, 5'(7')-$H_a$], 0.90 [s, 6H, 3'(9')-C<u>H</u>$_3$ or 4'(8')-C<u>H</u>$_3$], 0.91 [s, 6H, 4'(8')-C<u>H</u>$_3$ or 3'(9')-C<u>H</u>$_3$], 1.51 [d, *J* = 11.6 Hz, 2H, 2'(10')-$H_b$], 1.64 [d, *J* = 11.2 Hz, 2H, 5'(7')-$H_b$], 1.99 (s, 1H, 6'-H), 3.29 (s, 2H, C<u>H</u>$_2$NHC), 6.82 (s, 1H, NH), 7.45 (d, *J* = 7.2 Hz, 1H, 8-H), 7.59 (dd, *J* = 8.4 Hz, *J'* = 4 Hz, 1H, 3-H), 7.77 (t, *J* = 7.6 Hz, 1H, 7-H), 7.98 (d, *J* = 8.8 Hz, 1H, 6-H), 8.29 (d, *J* = 8.4 Hz, 1H, 4-H), 8.94 (d, *J* = 3.2 Hz, 1H, 2-H), 9.26 (broad s, 1H, NH). $^{13}$C-NMR (100.6 MHz, $CDCl_3$) δ: 15.5 [$CH_3$, C3'(9')-<u>C</u>$H_3$ or C4'(8')-<u>C</u>$H_3$], 15.7 [$CH_3$, C4'(8')-<u>C</u>$H_3$ or C3'(9')-<u>C</u>$H_3$], 36.4 (CH, C6'), 38.1 [$CH_2$, C5'(7')], 41.0 [$CH_2$, C2'(10')], 43.6 [C, C3'(9') or C4'(8')], 44.3 [C, C4'(8') or C3'(9')], 45.1 (C, C1'), 47.7 ($CH_2$, <u>C</u>$H_2$NCN), 117.2 (C, CN), 121.5 (CH, C3), 125.1 [(C, C4a) or (CH, C8)], 125.2 [(CH, C8) and (C, C4a)], 128.1 (CH, C6), 129.2 (CH, C7), 131.4 (CH, C4), 133.5 (C, C5), 148.4 (C, C8a), 150.7 (CH, C2), 159.6 (C, NCN). HRMS-ESI+ *m/z* $[M+H]^+$ calcd for $[C_{26}H_{32}N_5]^+$: 414.2652, found: 414.2649. Elemental analysis: calcd for $C_{26}H_{31}N_3S\cdot 0.75H_2O$: 73.12, H 7.67, N 16.40. Found: C 72.86, H 7.30, N 16.42.

**Synthesis of 2-(3,4,8,9-tetramethyltetracyclo[4.4.0.0$^{3,9}$.0$^{4,8}$]decan-1-yl)acetic acid, ALT-560, from known 3,4,8,9-tetramethyltetracyclo[4.4.0.0$^{3,9}$.0$^{4,8}$]decane-1-carboxylic acid.** a. Synthesis of the diazoketone (**146**). To an ice-water cooled solution of 3,4,8,9-tetramethyltetracyclo[4.4.0.0$^{3,9}$.0$^{4,8}$]decane-1-carboxylic acid (260 mg, 1.11 mmol) in DCM (721 mL) was added DMF (3 drops), followed by oxalyl chloride (130 µL, 1.52 mmol). The reaction mixture was allowed to stir at 0 °C for 6 h, and the solvent was evaporated under reduced pressure to leave crude acid chloride that was used directly. This was redissolved in a 1:1 solution of MeCN and THF (4 mL) and added dropwise to an ice-water cooled solution of trimethylsilyldiazomethane (0.8 mL, 2M in hexane, 1.60 mmol) and triethylamine (193 µL, 1.39 mmol) in a 1:1 solution of MeCN and THF (4 mL). The resulting yellow reaction mixture was allowed to warm to ambient temperature overnight. The solvent was removed under reduced pressure and the residue redissolved in ethyl acetate (5 mL) and washed with water (5 mL), saturated aq. $NaHCO_3$ solution (5 mL), and brine (5 mL). The organic phase was dried over anh. $Na_2SO_4$ and concentrated under vacuum. Purification by Combiflash® in silica gel using as eluent hexane to ethyl acetate/hexane mixture (1/9) gave the desired compound as a yellow solid (218 mg, 76% yield). $^1$H-NMR (400 MHz, $CDCl_3$) δ: 0.77 [dd, *J* = 11.2 Hz, *J'* = 2.5

Hz, 2H, 5(7)-$H_a$], 0.87 [d, *J* = 11.3 Hz, 2H, 2(10)-$H_a$], 0.96 [s, 12H, 3(9)-$CH_3$ and 4(8)-$CH_3$], 1.78 [dd, *J* = 11.7 Hz, *J'* = 1.3 Hz, 2H, 5(7)-$H_b$], 1.87 [d, *J* = 11.2 Hz, 2H, 2(10)-$H_b$], 2.56 (broad s, 1H, 6-H).

b) Synthesis of the methylester (**ALT-511**). The diazoketone (218 mg, 0.84 mmol) was dissolved in methanol (8 mL) and placed in an ultrasound bath, and a solution of silver benzoate (48 mg, 0.21 mmol) in triethylamine (466 µL, 3.36 mmol) was added dropwise and the mixture was sonicated for 1 h. The methanol was removed by evaporation and the residue dissolved in ethyl acetate (5 mL) and washed with saturated aq. $NaHCO_3$ solution (1x 5 mL), citric acid (2M, 1 x 5 mL), brine (1 x 5 mL), dried over anh. $Na_2SO_4$, and concentrated under vacuum. Purification by Combiflash® in silica gel using as eluent hexane to ethyl acetate/hexane mixture (0.5/9.5) gave the desired ester as a yellow oil (178 mg, 81% yield). IR (NaCl disk) ν: 2949, 2920, 2864, 1698, 1456, 1419, 1383, 1371, 1272, 1248, 1136, 1118, 1083, 1025, 938 $cm^{-1}$. $^1$H-NMR (400 MHz, $CDCl_3$) δ: 0.54 [d, *J* = 10.8 Hz, 2H, 2(10)-$H_a$], 0.66 [dd, *J* = 10.8 Hz, *J'* = 2.8 Hz, 2H, 5(7)-$H_a$], 0.92 [s, 6H, 3(9)-$CH_3$ or 4(8)-$CH_3$], 0.93 [s, 6H, 4(8)-$CH_3$ or 3(9)-$CH_3$], 1.74-1.78 [complex signal, 4H, 2(5,7,10)-$H_b$], 2.14 (m, 1H, 6-H), 2.44 (s, 2H, $CH_2CO$), 3.67 (s, 3H, $OCH_3$). $^{13}$C-NMR (100.6 MHz, $CDCl_3$) δ: 15.7 [$CH_3$, C3(9)-<u>C</u>$H_3$ or C4(8)-<u>C</u>$H_3$], 15.9 [$CH_3$, C4(8)-<u>C</u>$H_3$ or C3(9)-<u>C</u>$H_3$], 38.8 [$CH_2$, C2(10)], 39.2 (CH, C6), 40.5 (C, C1), 43.6 ($CH_2$, <u>C</u>$H_2$CO), 44.0 [$CH_2$, C5(7)], 44.7 [C, C3(9) or C4(8)], 45.9 [C, C4(8) or C3(9)], 51.3 ($CH_3$, O<u>C</u>$H_3$), 173.4 (C, CO).

c) 2-(3,4,8,9-tetramethyltetracyclo[4.4.0.0$^{3,9}$.0$^{4,8}$]decan-1-yl)acetic acid, **ALT-560**. A mixture of ester **147** (178 mg, 0.68 mmol) in a 40% methanol solution of KOH (2 mL) was heated to reflux for 2 h. Water (2 mL) was added and the reaction was refluxed for 3 h. The reaction mixture was allowed to cool down to room temperature and the methanol was removed under vacuum. Water (3 mL) was added to the residue and the solution acidified with conc. HCl until pH = 1. The aq. layer was extracted with DCM (3 x 3 mL) and the combined organics were dried over anh. $Na_2SO_4$, filtered and concentrated under reduced pressure to give a white solid (158 mg, 94% yield), mp 142 °C. IR (NaCl disk) ν: 2948, 2922, 2865, 1737, 1479, 1455, 1434, 1385, 1372, 1272, 1224, 1213, 1172, 1132, 1108, 1081 $cm^{-1}$. $^1$H-NMR (400 MHz, $CDCl_3$) δ: 0.57 [d, *J* = 11.2 Hz, 2H, 2(10)-$H_a$], 0.68 [dd, *J* = 11.2 Hz, *J'* = 2.8 Hz, 2H, 5(7)-$H_a$], 0.926 [s, 6H, 3(9)-$CH_3$ or 4(8)-$CH_3$], 0.934 [s, 6H, 4(8)-$CH_3$ or 3(9)-$CH_3$], 1.78 [dd, *J* = 11.6 Hz, *J'* = 1.2 Hz, 2H, 2(10)-$H_b$], 1.81 [d, *J* = 10.8 Hz, 2H, 5(7)-$H_b$], 2.17 (m, 1H, 6-H), 2.49 (s, 2H, $CH_2CO$). $^{13}$C-NMR (100.6 MHz, $CDCl_3$) δ: 15.7 [$CH_3$, C3(9)-<u>C</u>$H_3$ or C4(8)-<u>C</u>$H_3$], 15.9 [$CH_3$, C4(8)-<u>C</u>$H_3$ or C3(9)-<u>C</u>$H_3$], 38.8 [$CH_2$, C2(10)], 39.2 (CH, C6), 40.3 (C, C1), 43.5 ($CH_2$, CO<u>C</u>$H_2$), 44.0 [$CH_2$, C5(7)], 44.7 [C, C3(9) or C4(8)], 45.9 [C, C3(9) or C4(8)], 178.5 (C, CO).

**Radiochemistry: synthesis and characterization of [$^{11}$C]UB-CB-P3**

[$^{11}$C]$CH_4$ was directly produced in an IBA Cyclone 18/9 cyclotron by proton irradiation of a $N_2$/5% $H_2$ gas mixture and transferred to a GE TRACERlab™ $FX_{CPro}$ synthesizer (GE Healthcare, Waukesha, WI, USA). Within the synthesizer, [$^{11}$C]$CH_3I$ was generated using the gas-phase method.[2] At the end of the process, [$^{11}$C]$CH_3I$ was distilled under a continuous helium flow (20 mL/min) and directed into a stainless steel 2 mL loop, precharged with a solution of the appropriate desmethyl precursor, UB-CB-P4 (1 mg), dissolved in DMSO (100 µL) in the presence of 4 M NaOH (2 µL). The mixture was allowed to react for 6 min at room temperature. The crude product was then purified by semi-preparative high-performance liquid chromatography (HPLC) using a Mediterranea Sea 18 column (250 × 10 mm, 5 µm). The mobile phase consisted of 10 mM ammonium formate (pH 3.9)/acetonitrile (70/30, v/v), delivered at a flow rate of 5 mL/min. The desired fraction (retention time: 10–10.5 min) was collected and reformulated using a C18 cartridge (Sep-Pak C18 Plus Light Cartridge, Waters). The product was eluted with 1 mL of absolute ethanol into the final vial for dose formulation with 0.9% saline solution. Chemical and radiochemical purity were assessed by analytical radio-HPLC (Agilent 1200 Series, USA) equipped with a radio detector (Gabi, Elysia-Raytest, Germany) and a variable wavelength UV detector (Agilent, USA) connected in series. Separation was performed on an Agilent Zorbax Eclipse XDB-C18 column (150 × 4.6 mm, 5 µm) using a gradient mobile phase of solvent A (10 mM ammonium formate, pH 3.9) and solvent B (acetonitrile): 95% A to 50% A (2 min), to 5% A (15 min), held to 20 min, then returned to 95% A (22 min), at a flow rate of 1 mL/min. UV absorbance was monitored at 220 nm. The identity of [$^{11}$C]UB-CB-P3 (retention time = 10 min) was confirmed by co-elution with the non-radioactive reference standard.

The octanol-water partition coefficient (log*P*) was determined by adding 1.48 MBq of the radioligand to an Eppendorf tube containing 1.5 mL each of ultrapure water and n-octanol. The mixture was vortexed for 5 min and then centrifuged for 5 min at room temperature. Subsequently, 1 mL aliquots from each phase were collected separately, and radioactivity was quantified using a γ-counter.

[2] Larsen, P., Ulin J., Dahlstrom K. & Jensen M. Synthesis of [$^{11}$C]iodomethane by iodination of [$^{11}$C]methane. *Appl. Radiat. Isot.* **48**, 153–157 (1997).

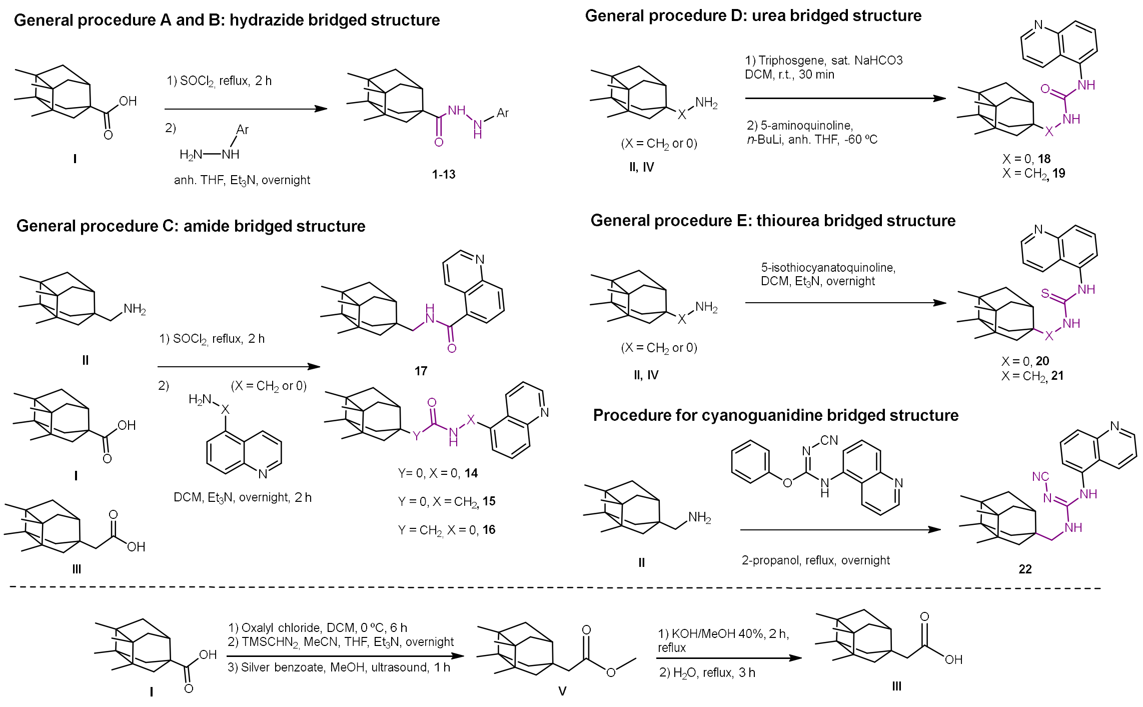


**Supplementary Figure 1.** General procedures for the synthesis of the new derivatives.

$^1$H NMR (400 MHz, DMSO-$d_6$)

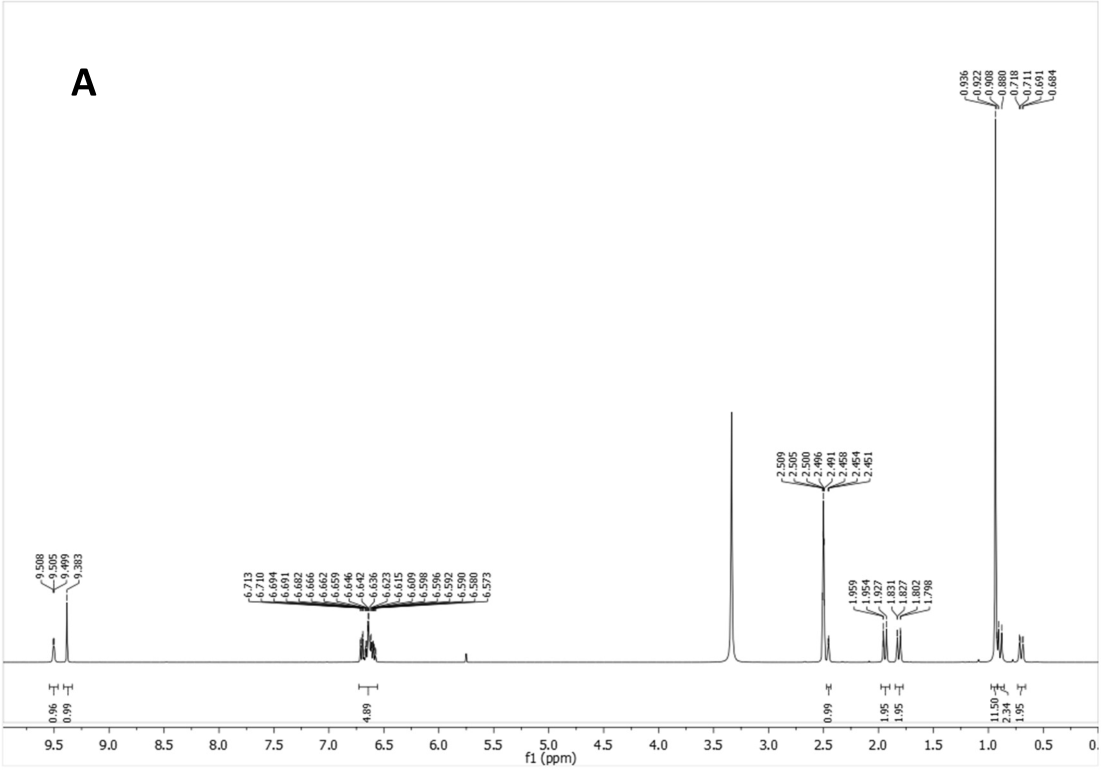


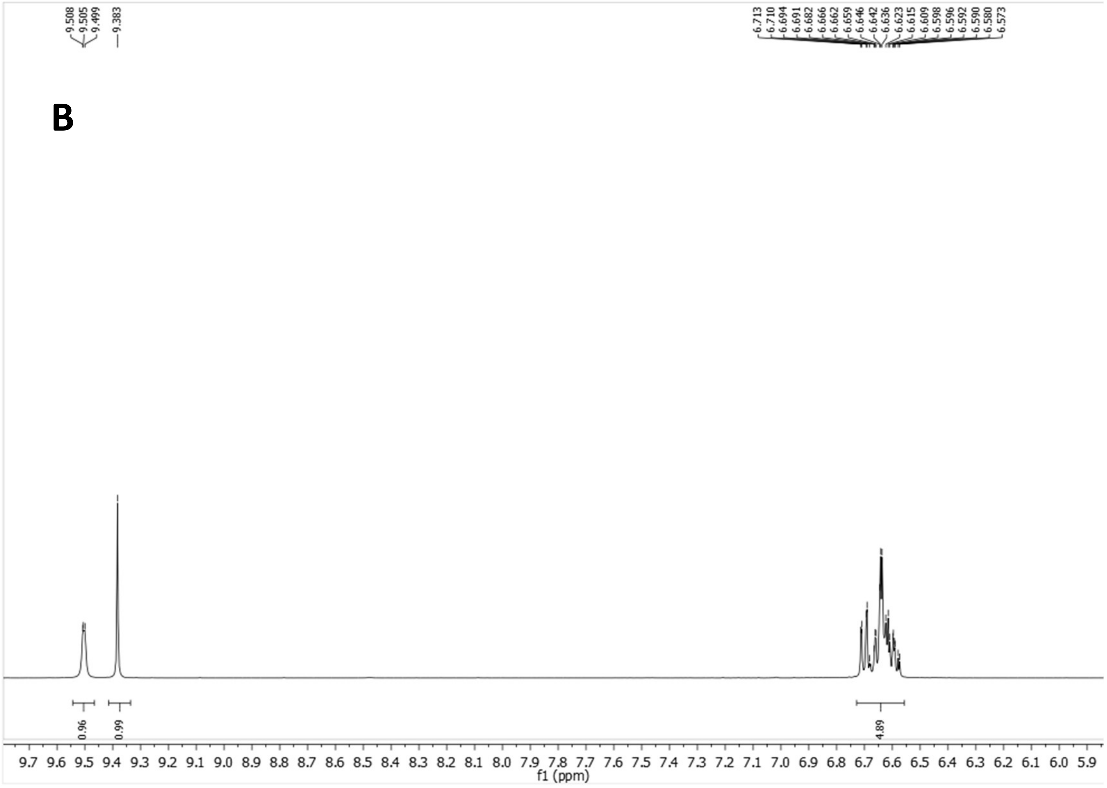

$^{13}$C-NMR (100.6 MHz, DMSO-$d_6$)

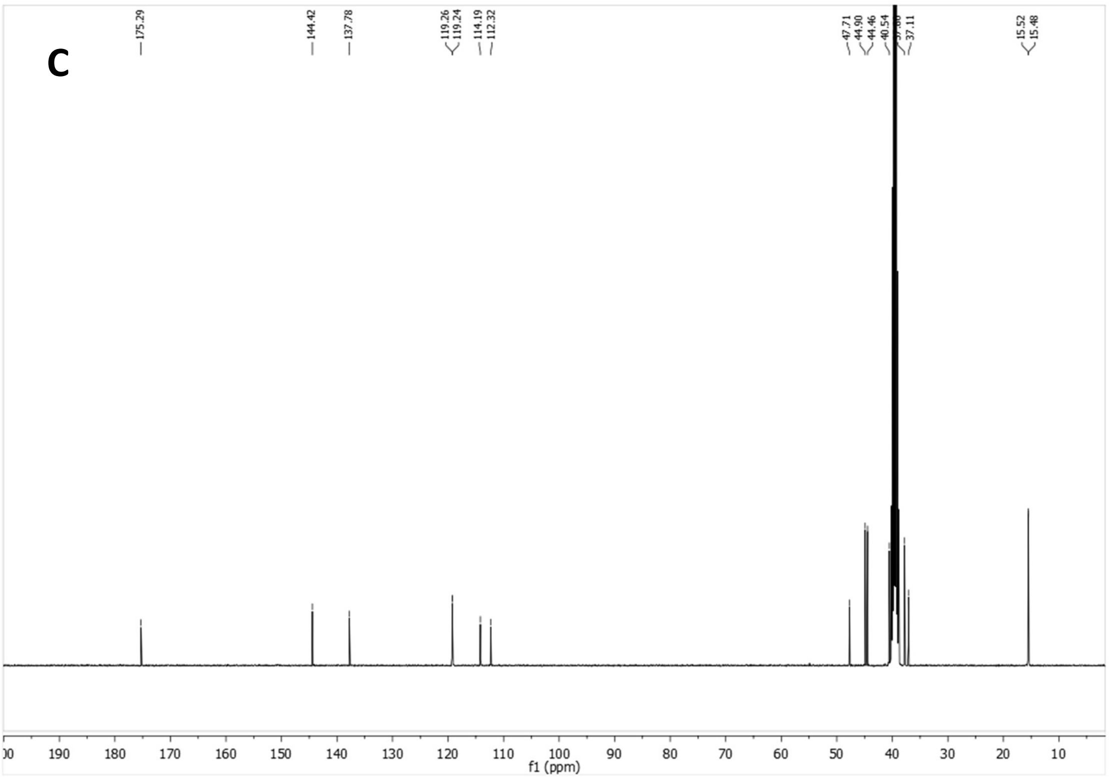


HSQC

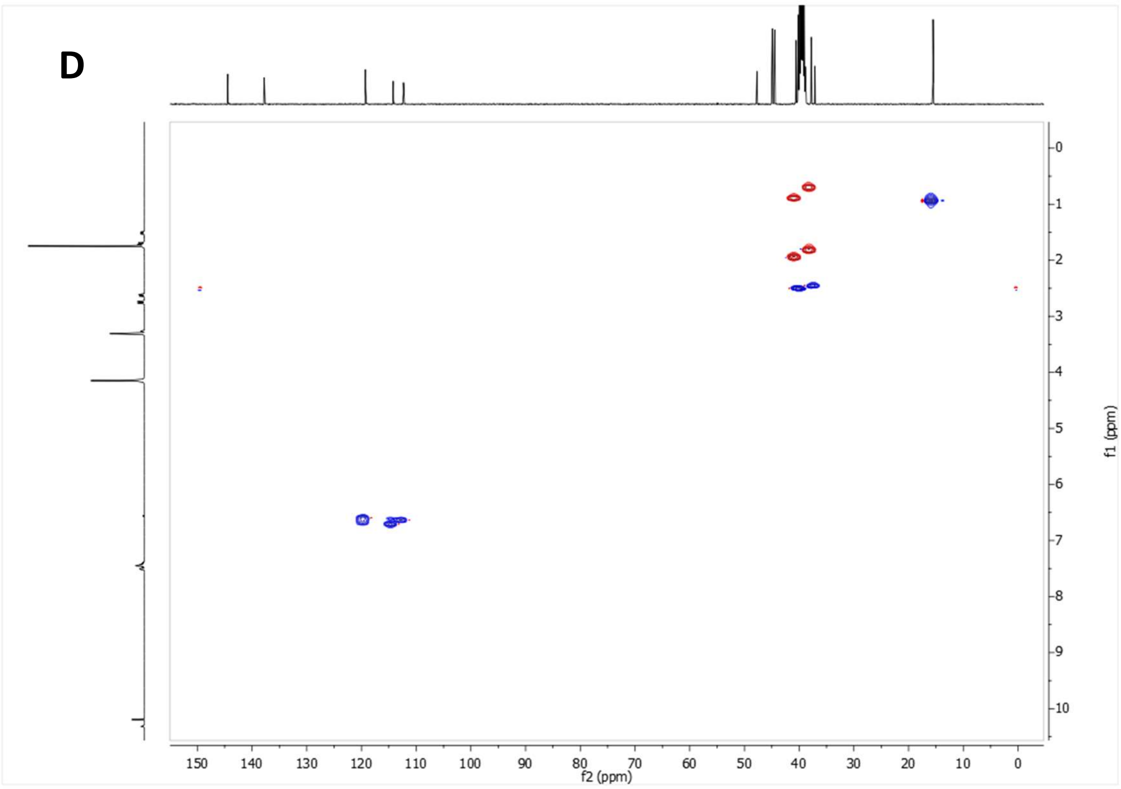


**Supplementary Figure 2:** *N'*-(2-Hydroxyphenyl)-3,4,8,9-tetramethyltetracyclo[4.4.0.0$^{3,9}$.0$^{4,8}$]decane-1-carbohydrazide (CB-P4). (A-B) $^{1}$H NMR spectrum. (C) $^{13}$C NMR spectrum. (D) HSQC spectrum.

$^1H$ NMR (400 MHz, $CDCl_3$)

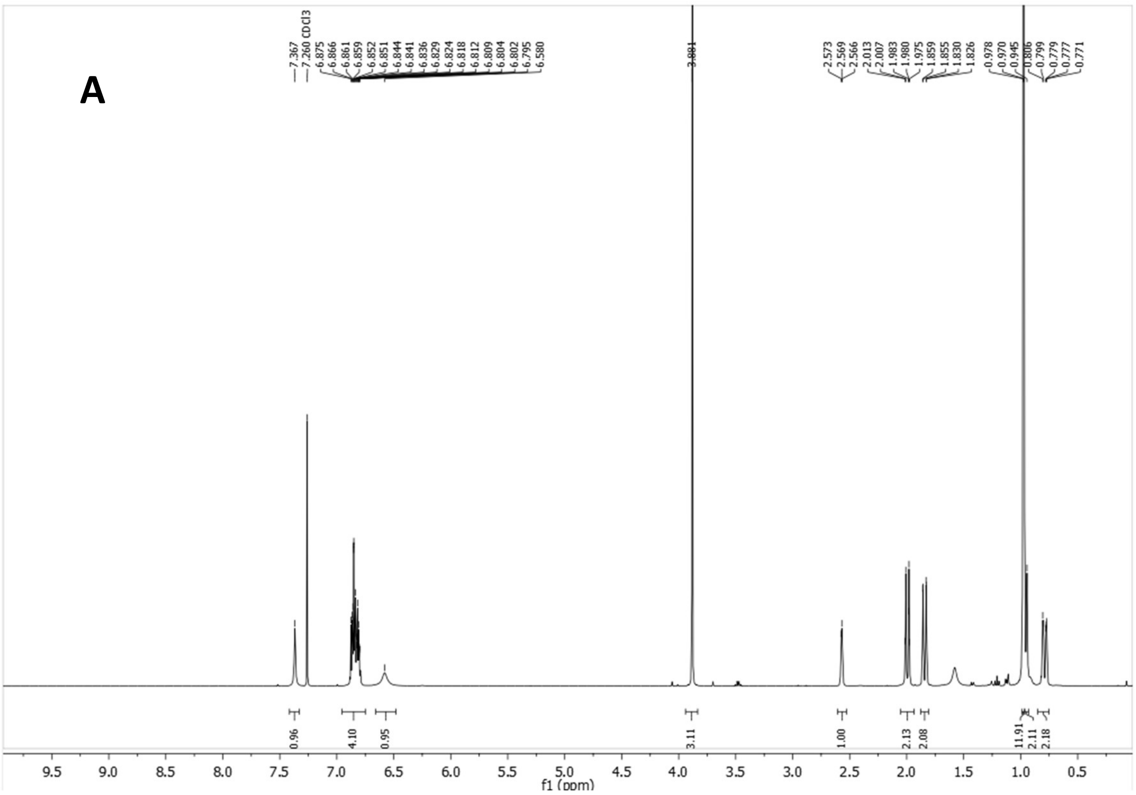

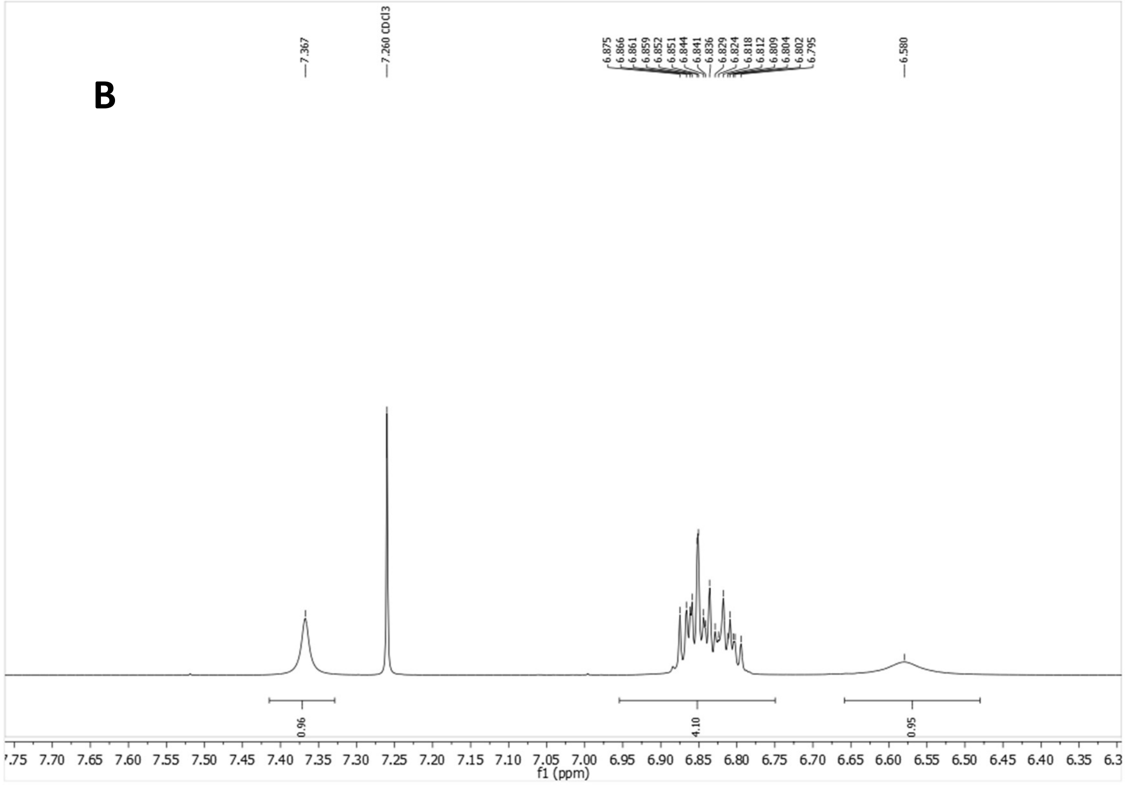

[13]C-NMR (100.6 MHz, )

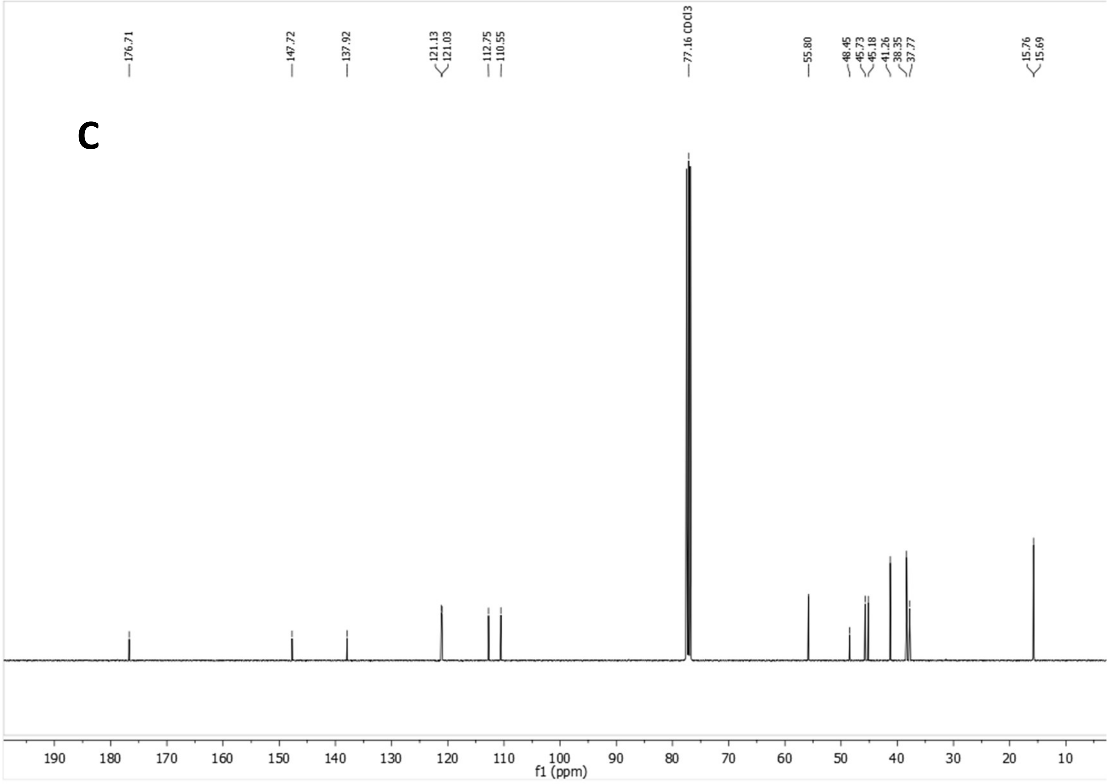


HSQC

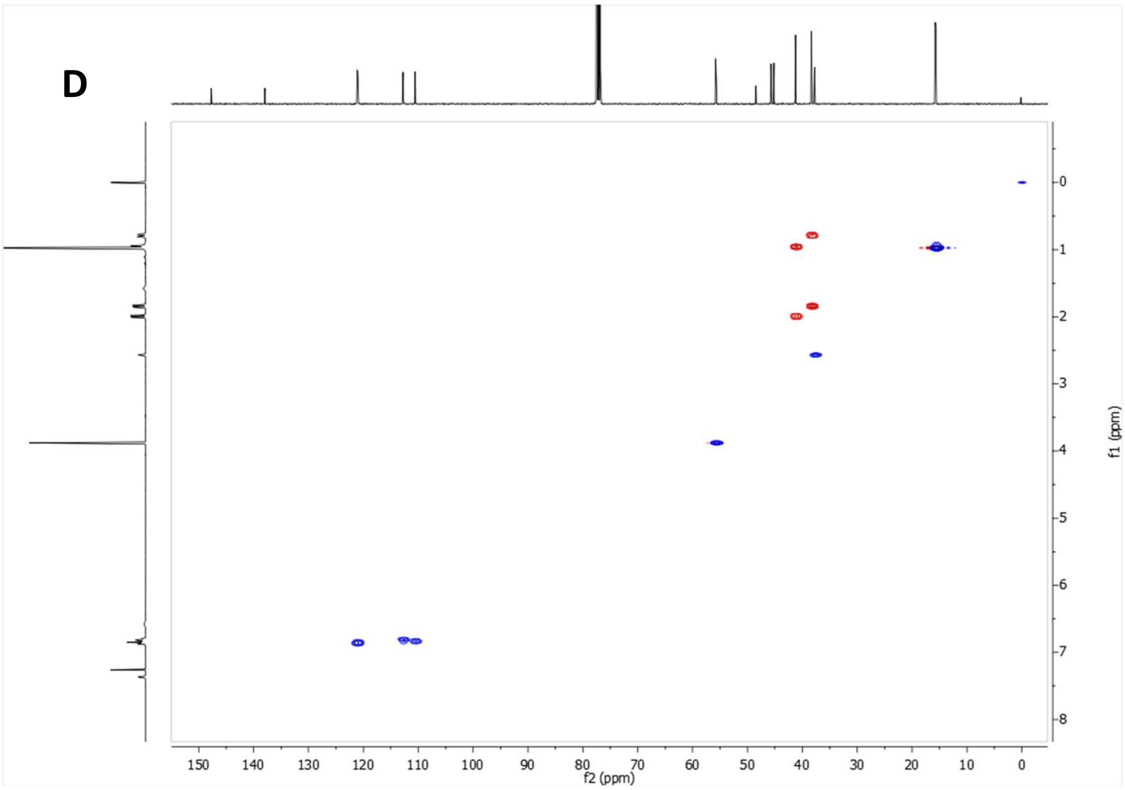


**Supplementary Figure 3:** *N'*-(2-Methoxyphenyl)-3,4,8,9-tetramethyltetracyclo[4.4.0.0$^{3,9}$.0$^{4,8}$]decane-1-carbohydrazide (CB-P3). (A-B) $^{1}$H NMR spectrum. (C) $^{13}$C NMR spectrum. (D) HSQC spectrum.

$^1H$ NMR (400 MHz, $CDCl_3$)

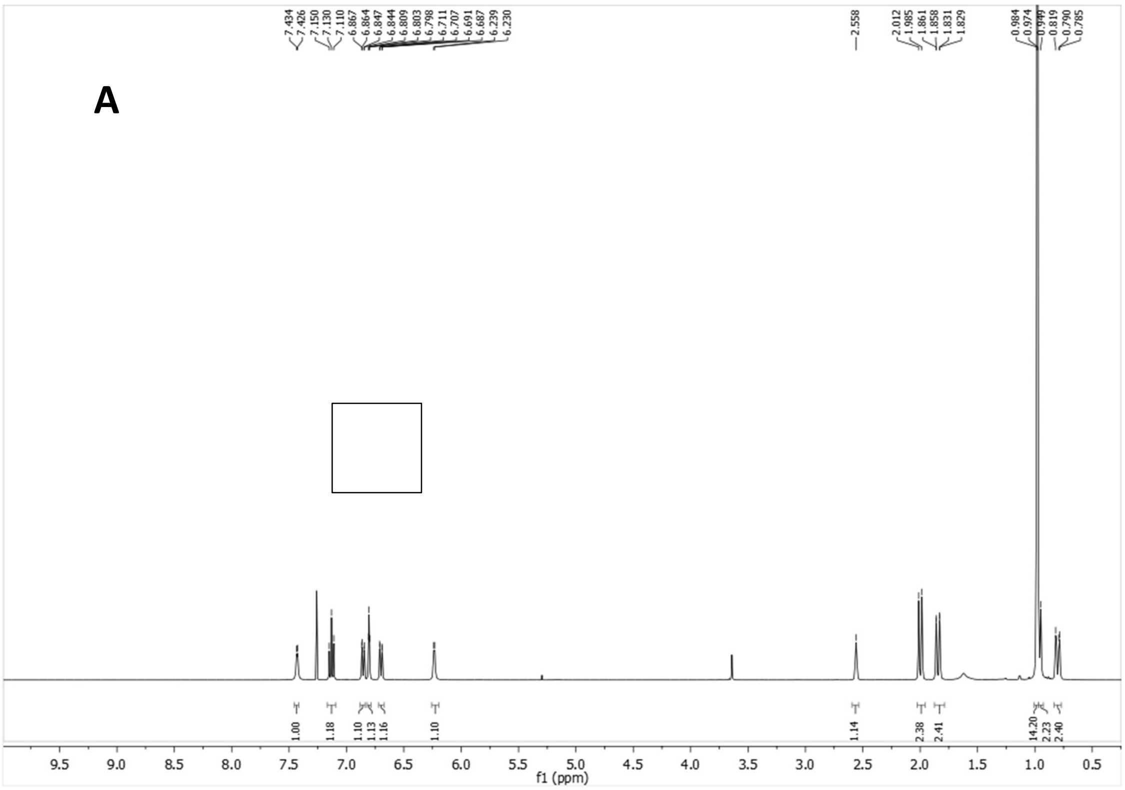


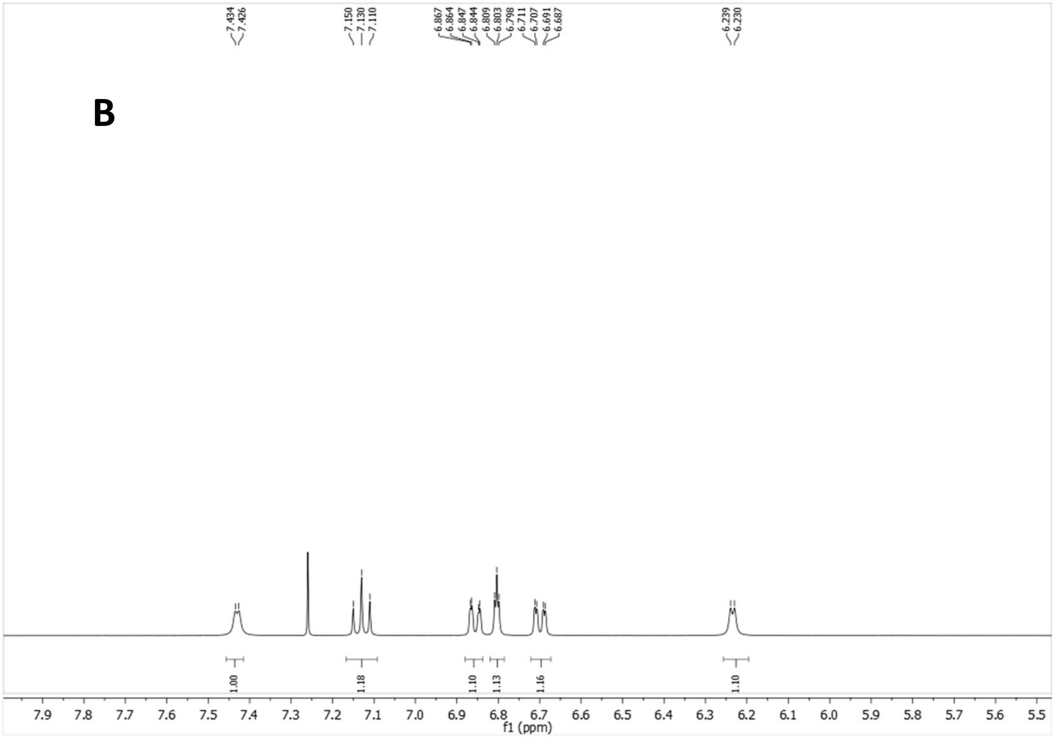

$^{13}C$-NMR (100.6 MHz, $CDCl_3$)

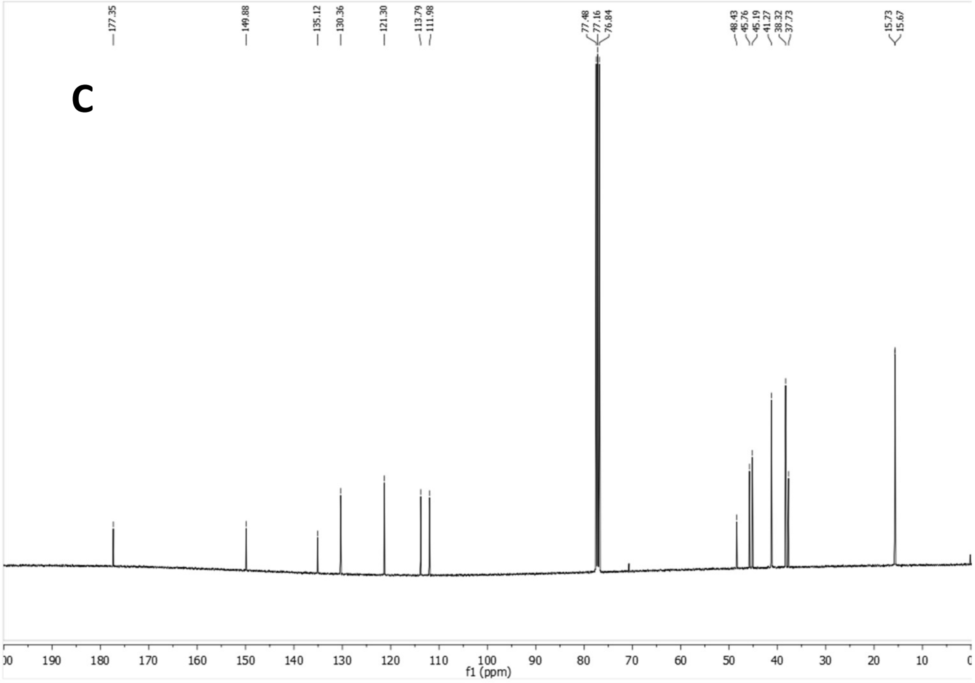


HSQC

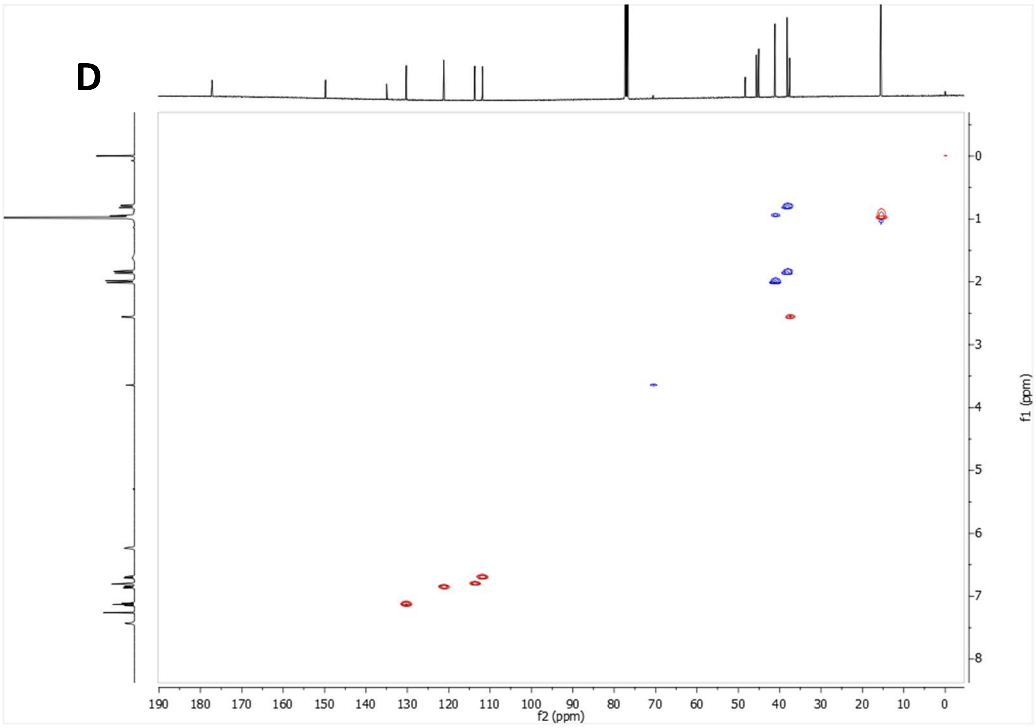


**Supplementary Figure 4:** *N'*-(3-Chlorophenyl)-3,4,8,9-tetramethyltetracyclo[4.4.0.0$^{3,9}$.0$^{4,8}$]decane-1-carbohydrazide (ALT-P39**).** (A-B) $^{1}H$ NMR spectrum. (C) $^{13}C$ NMR spectrum. (D) HSQC spectrum.

$^1H$ NMR (400 MHz, $CDCl_3$)

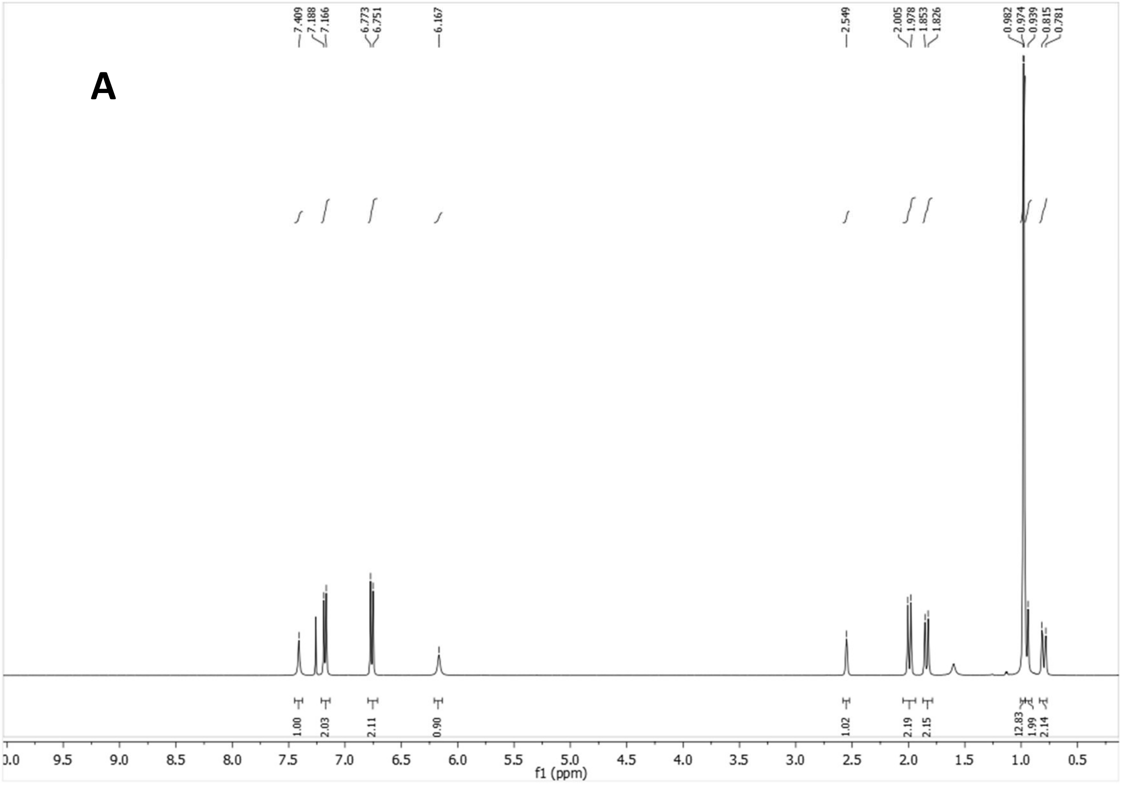


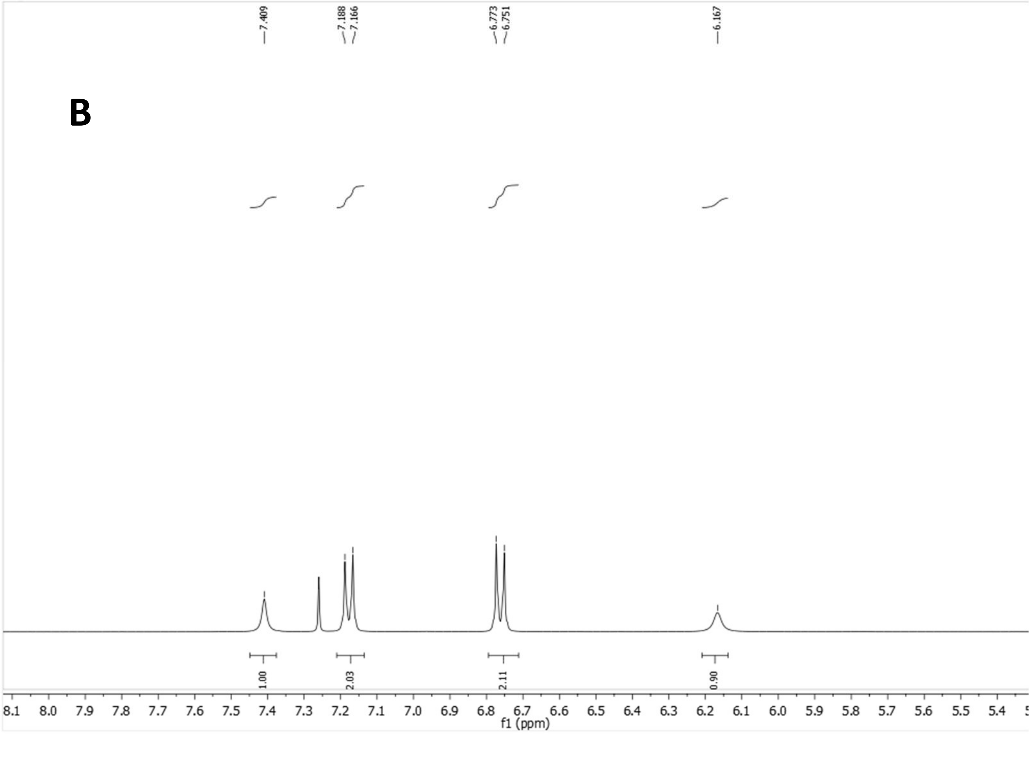

$^{13}$C-NMR (100.6 MHz, $CDCl_3$)

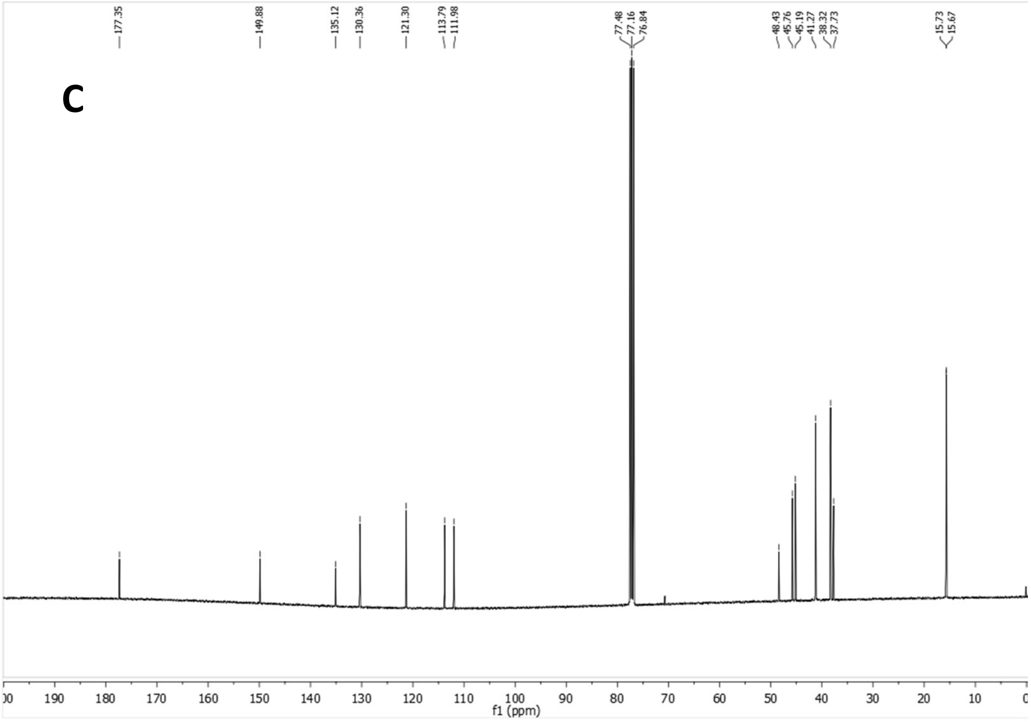


HSQC

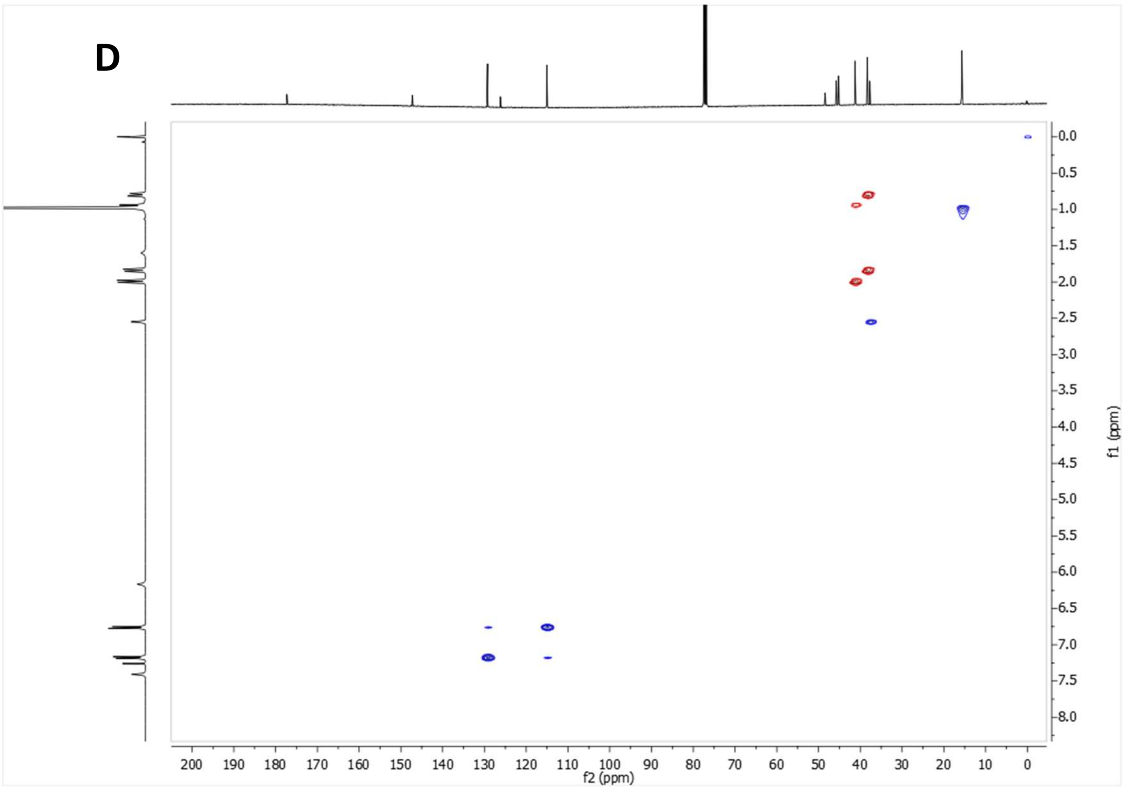


**Supplementary Figure 5:** *N'*-(4-Chlorophenyl)-3,4,8,9-tetramethyltetracyclo[4.4.0.0$^{3,9}$.0$^{4,8}$]decane-1-carbohydrazide (ALT-P40). (A-B) $^{1}$H NMR spectrum. (C) $^{13}$C NMR spectrum. (D) HSQC spectrum.

$^1H$ NMR (400 MHz, $CDCl_3$)

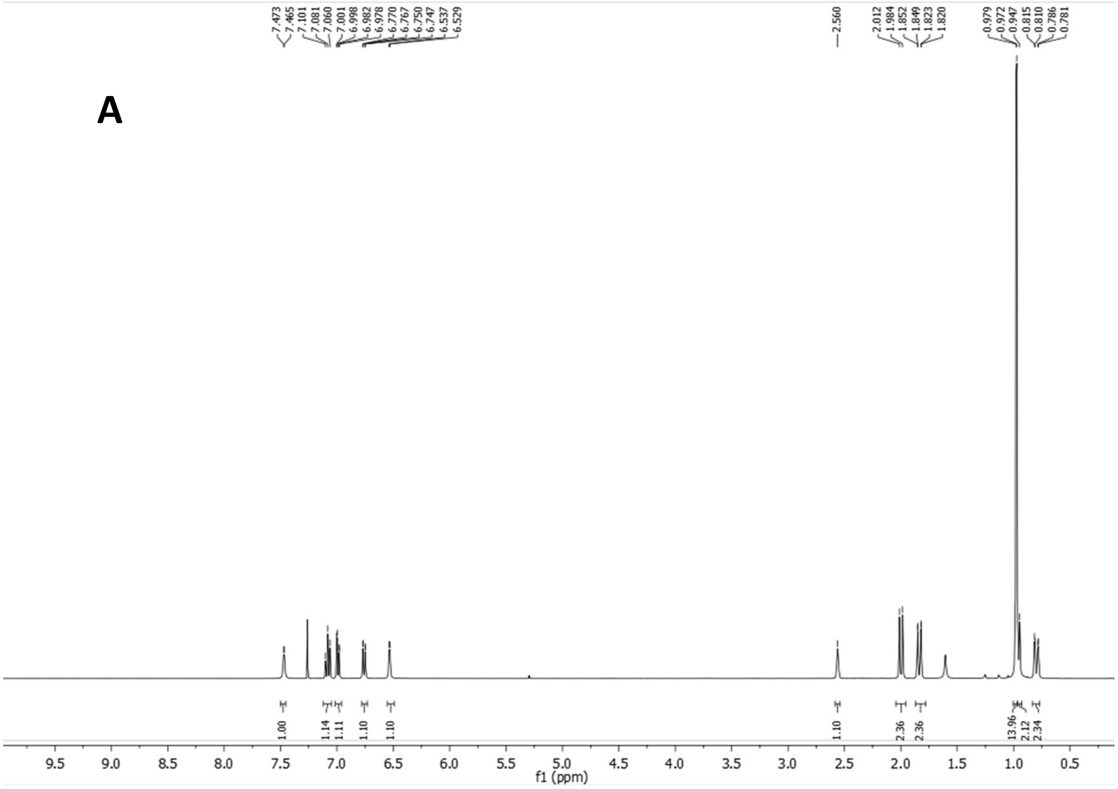

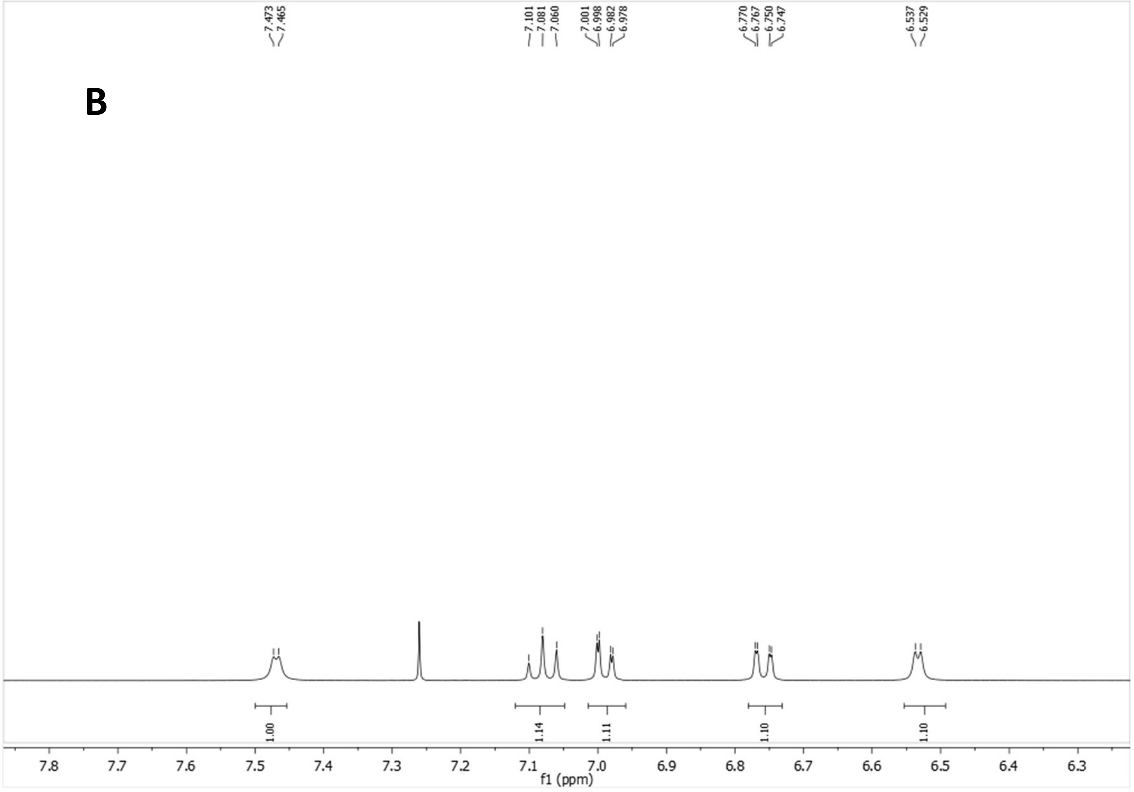

$^{13}$C-NMR (100.6 MHz, $CDCl_3$)

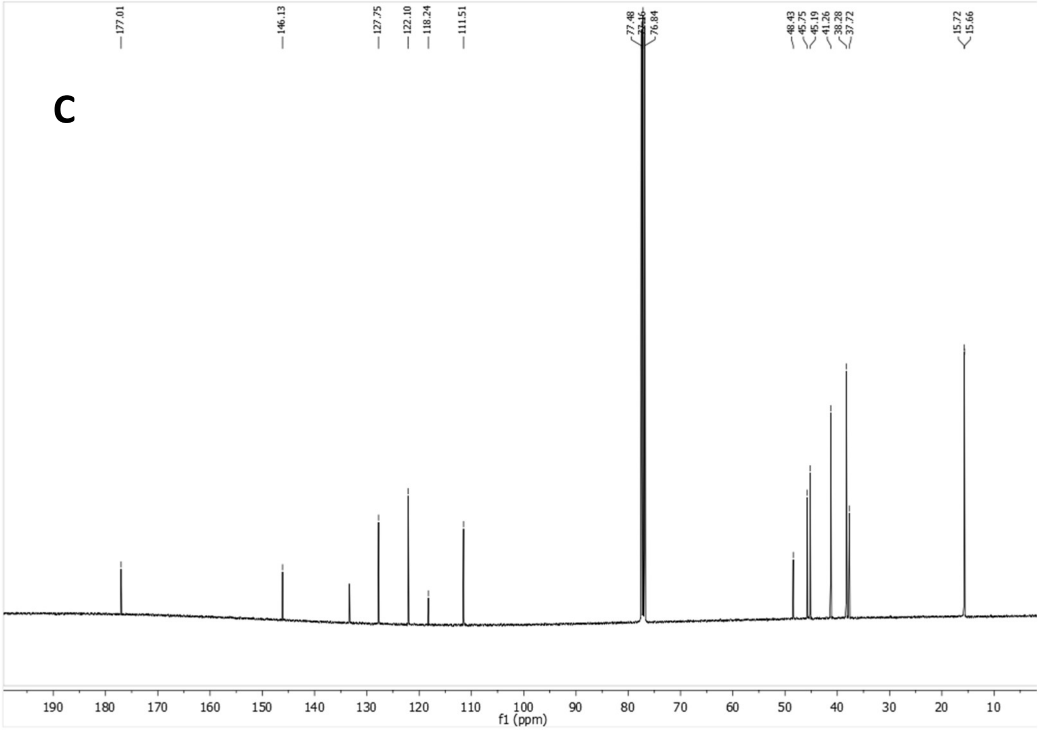


HSQC

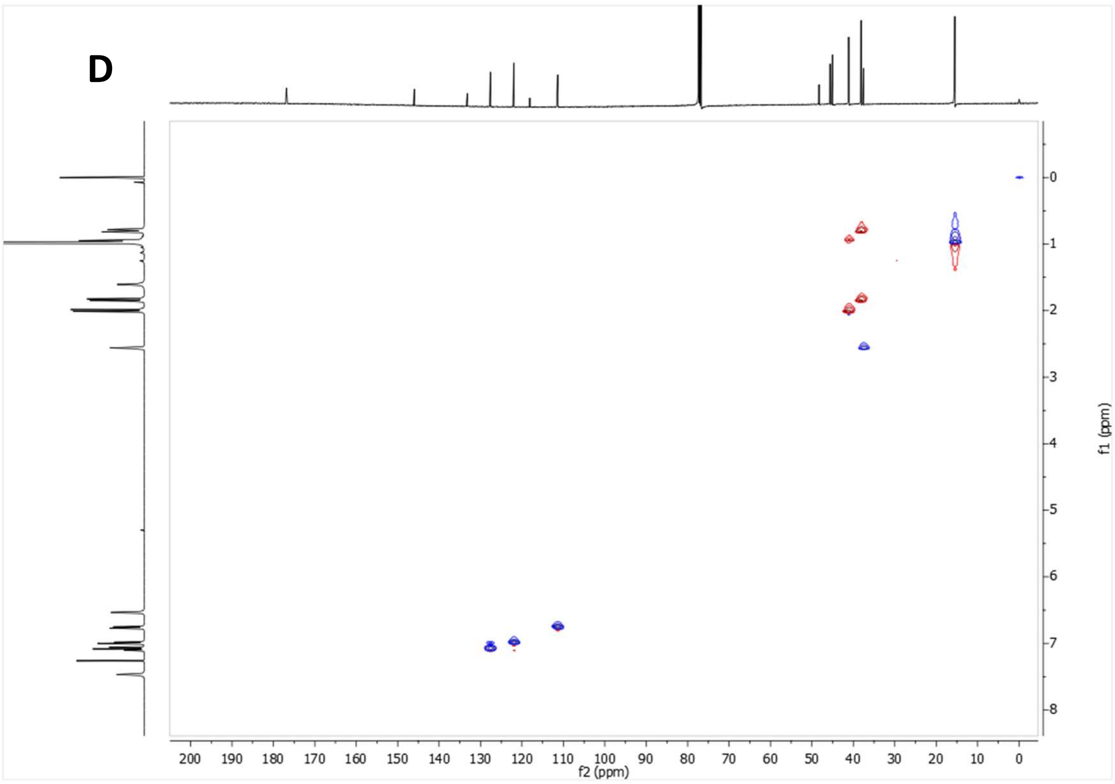


**Supplementary Figure 6:** *N'*-(2,3-Dichlorophenyl)-3,4,8,9-tetramethyltetracyclo[4.4.0.0$^{3,9}$.0$^{4,8}$]decane-1-carbohydrazide (ALT-P41). (A-B) $^{1}$H NMR spectrum. (C) $^{13}$C NMR spectrum. (D) HSQC spectrum.

[1]H NMR (400 MHz, $CDCl_3$)

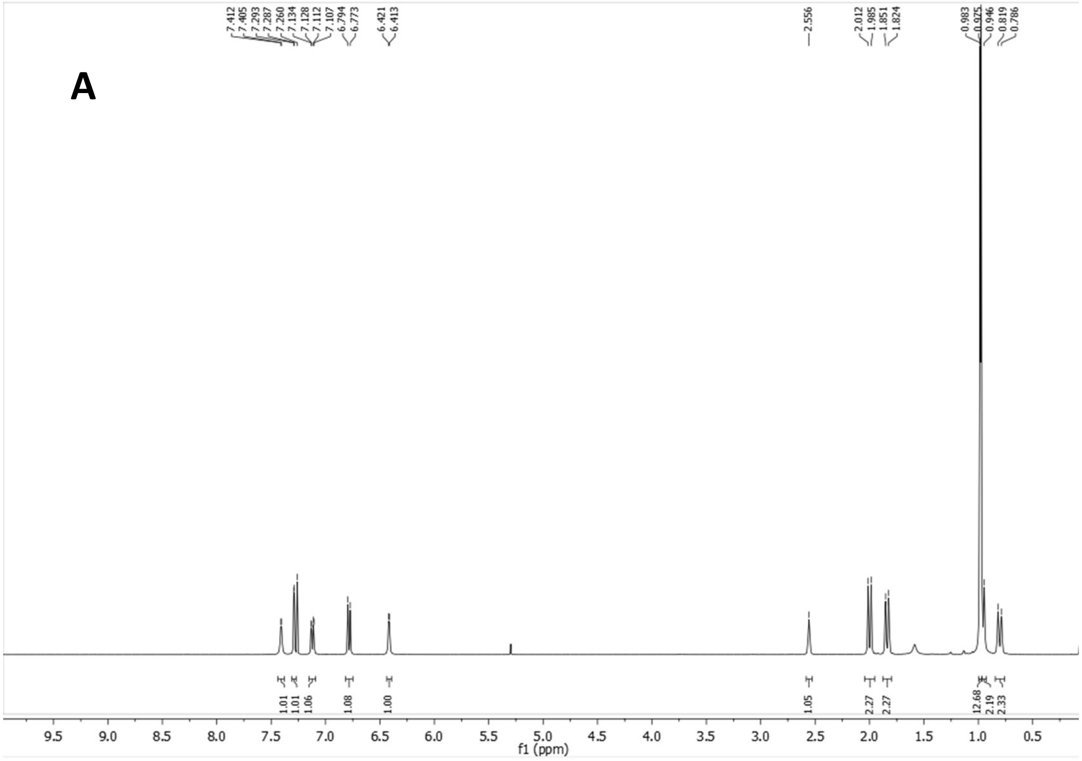


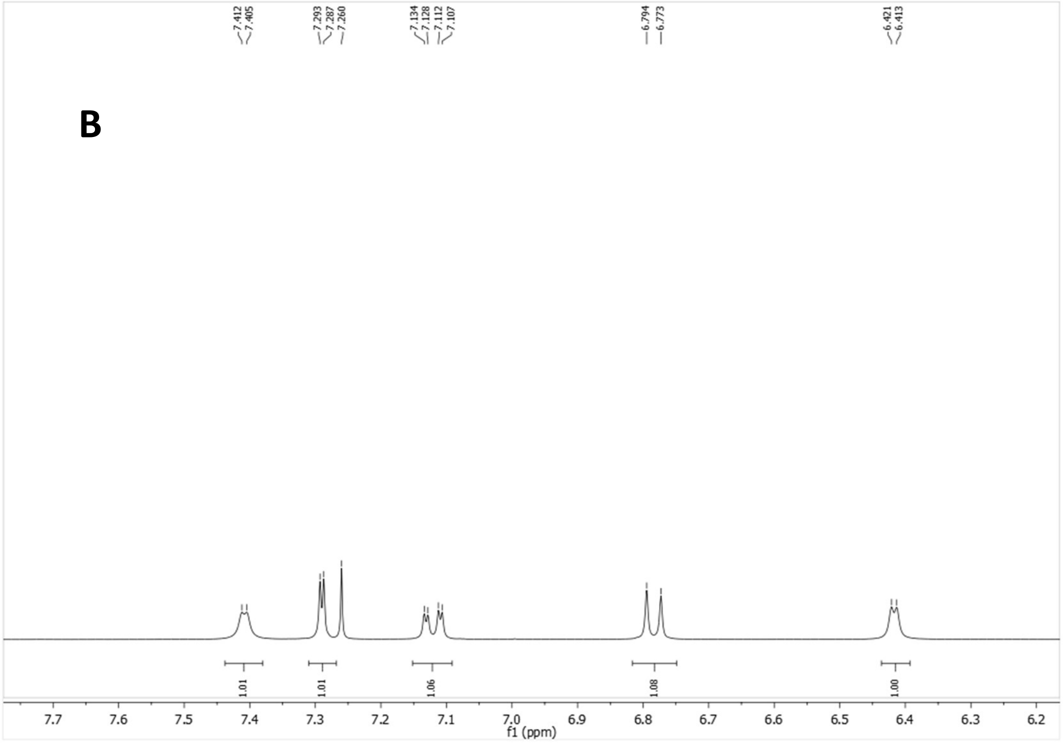

$^{13}C$-NMR (100.6 MHz, $CDCl_3$)

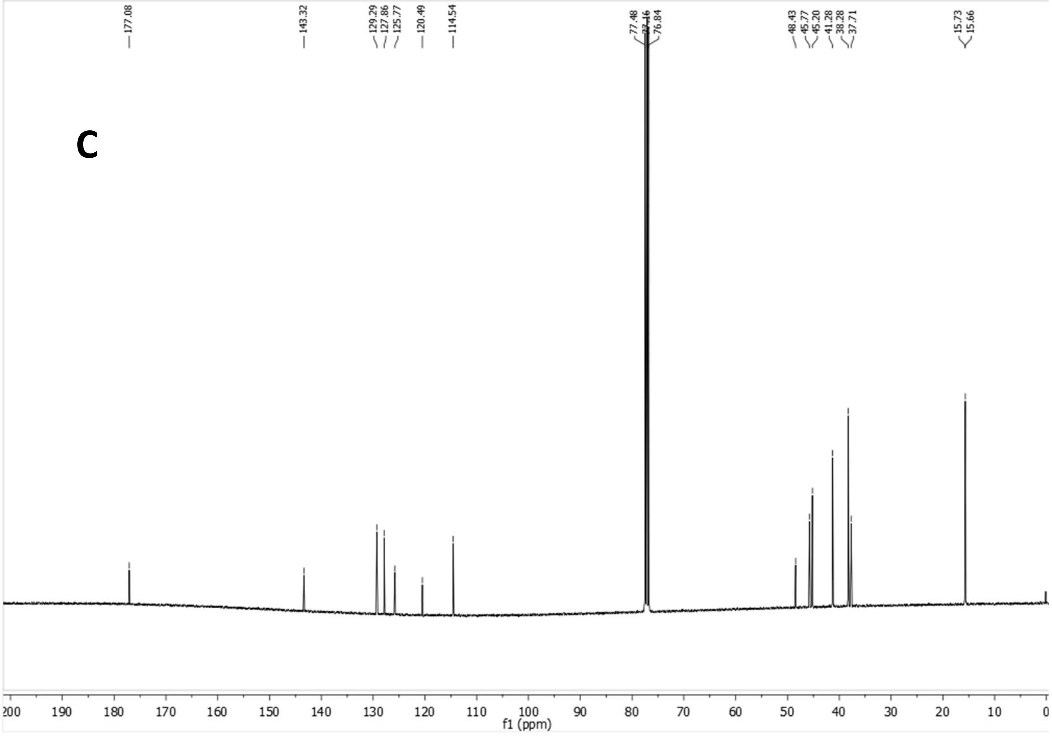


HSQC

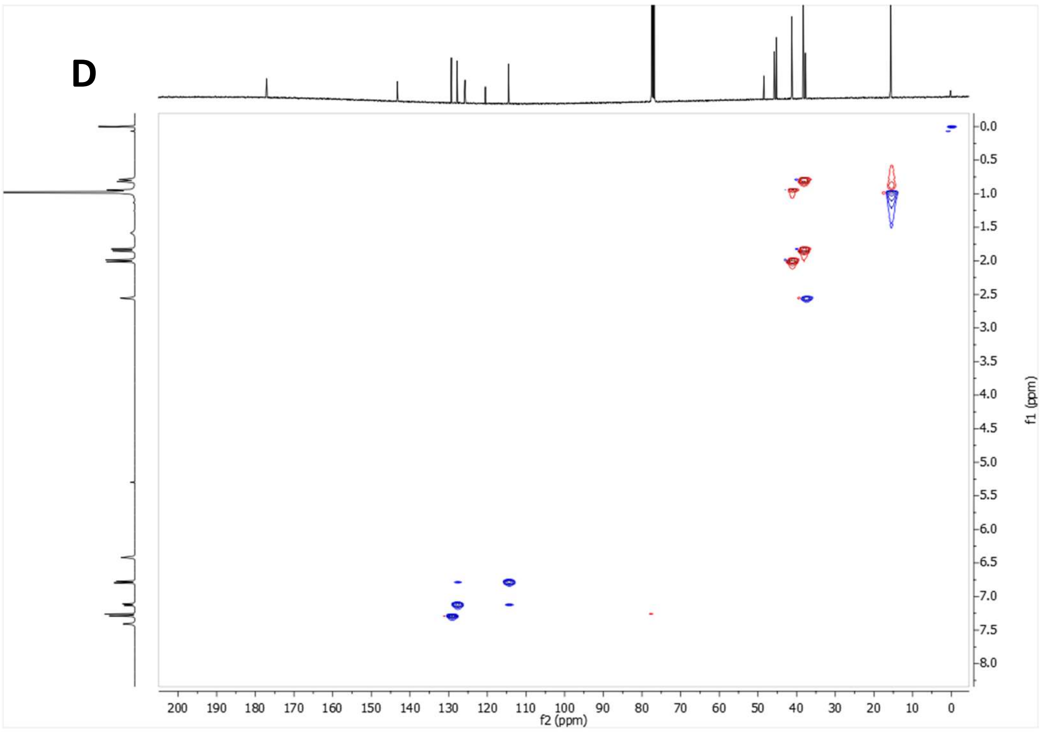


**Supplementary Figure 7:** *N'*-(2,4-Dichlorophenyl)-3,4,8,9-tetramethyltetracyclo[4.4.0.0$^{3,9}$.0$^{4,8}$]decane-1-carbohydrazide (ALT-P42). (A-B) $^{1}H$ NMR spectrum. (C) $^{13}C$ NMR spectrum. (D) HSQC spectrum.

$^1H$ NMR (400 MHz, $CDCl_3$)

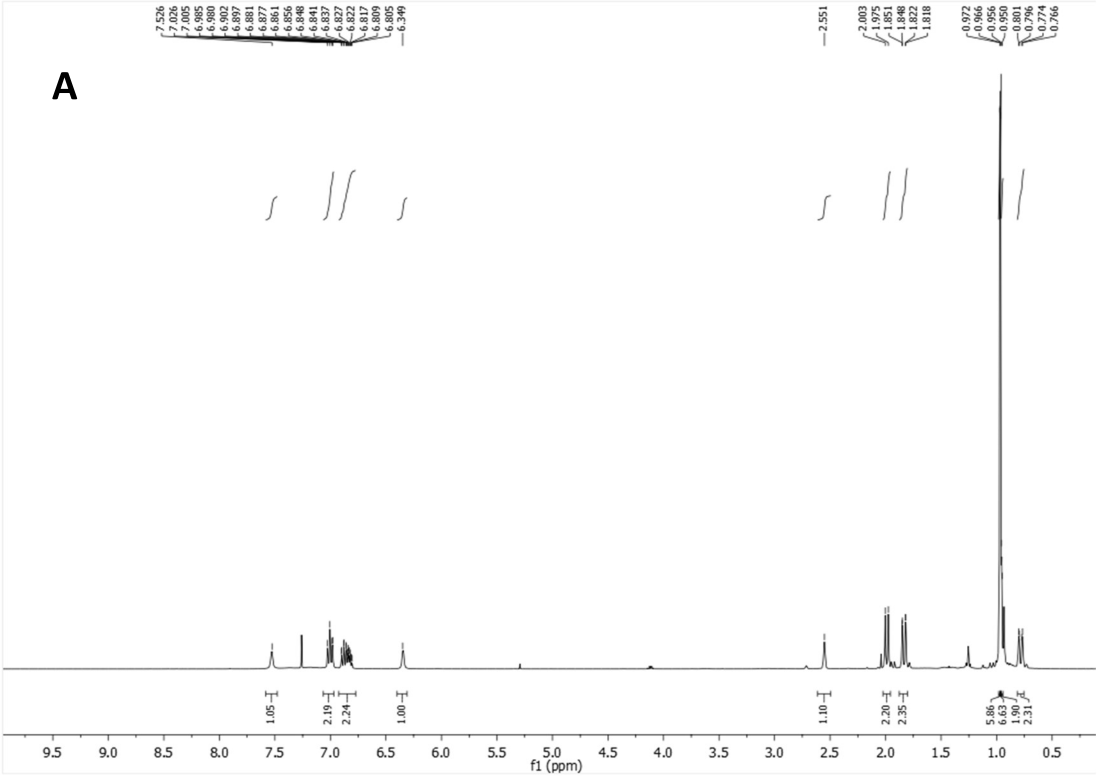


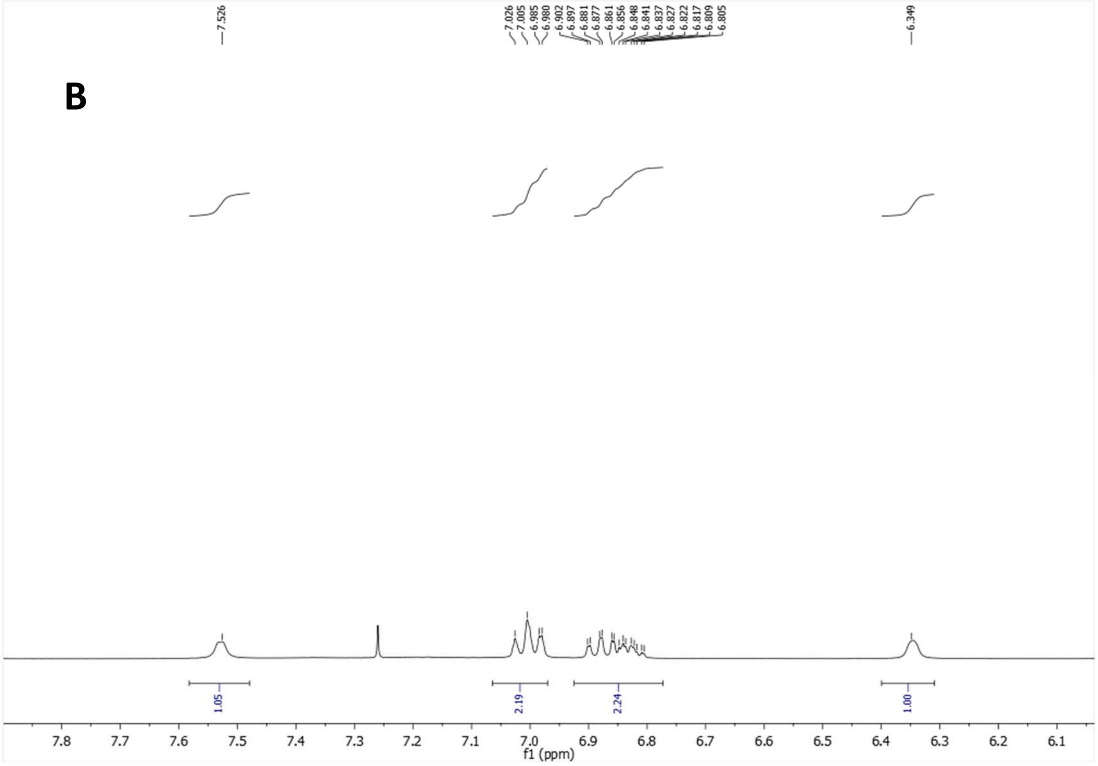

$^{13}C$-NMR (100.6 MHz, $CDCl_3$)

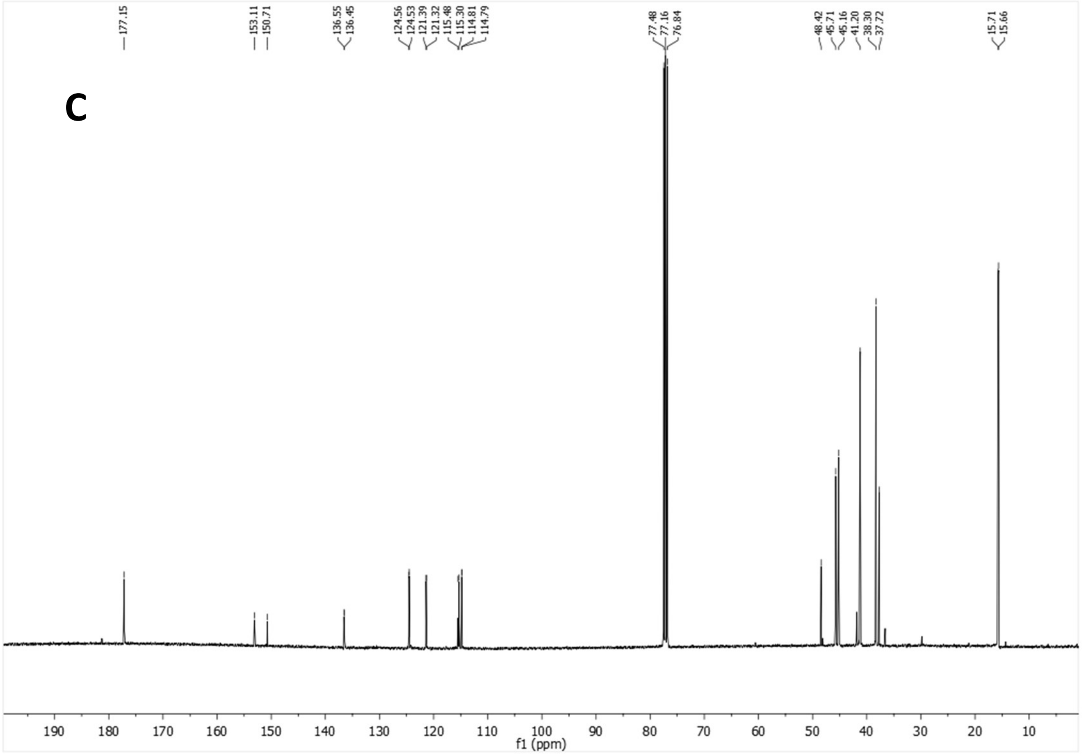


HSQC

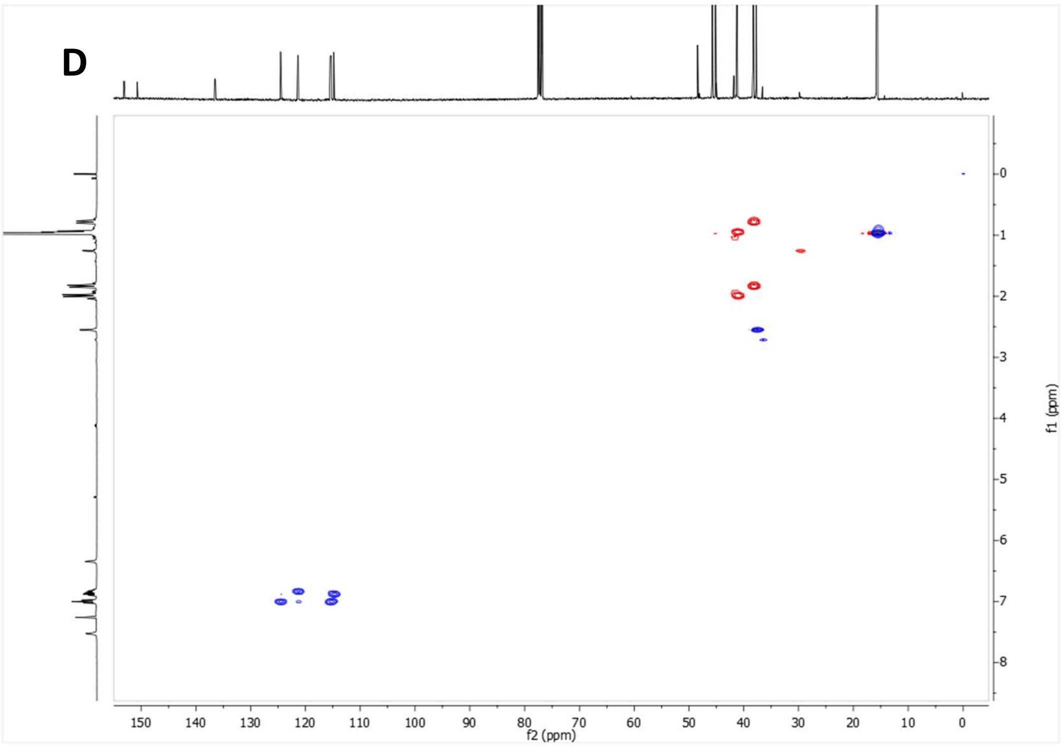

$^{19}F$-NMR (376.5 MHz, $CDCl_3$)

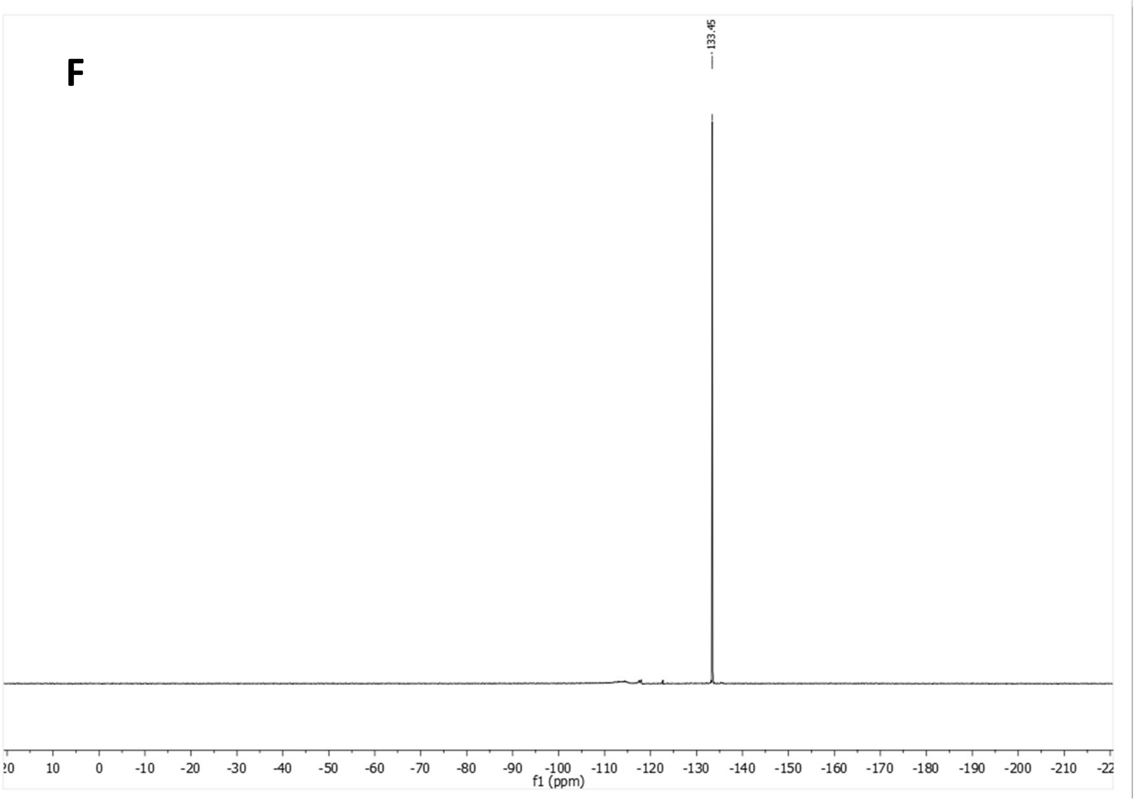


**Supplementary Figure 8:** *N'*-(2-Fluorophenyl)-3,4,8,9-tetramethyltetracyclo[4.4.0.0$^{3,9}$.0$^{4,8}$]decane-1-carbohydrazide (CB-P1). (A-B) $^{1}H$ NMR spectrum. (C) $^{13}C$ NMR spectrum. (D) HSQC spectrum. (F) $^{19}F$ spectrum.

$^1$H NMR (400 MHz, $CDCl_3$)

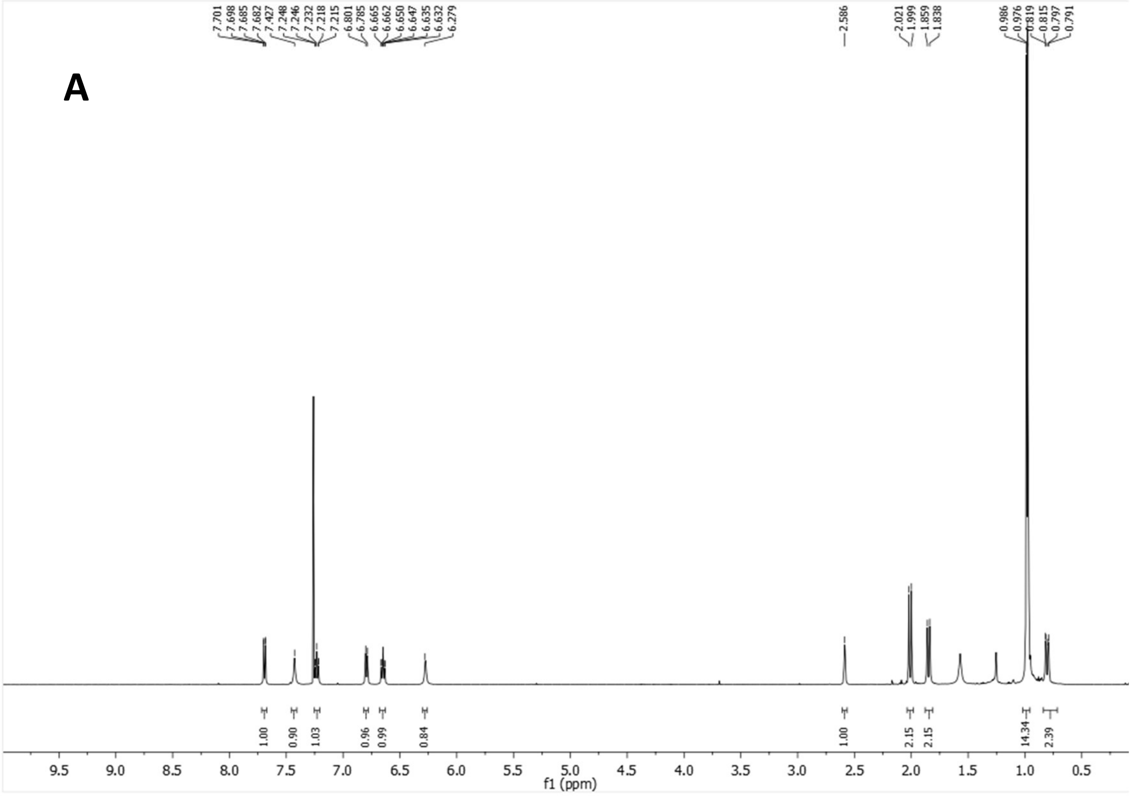


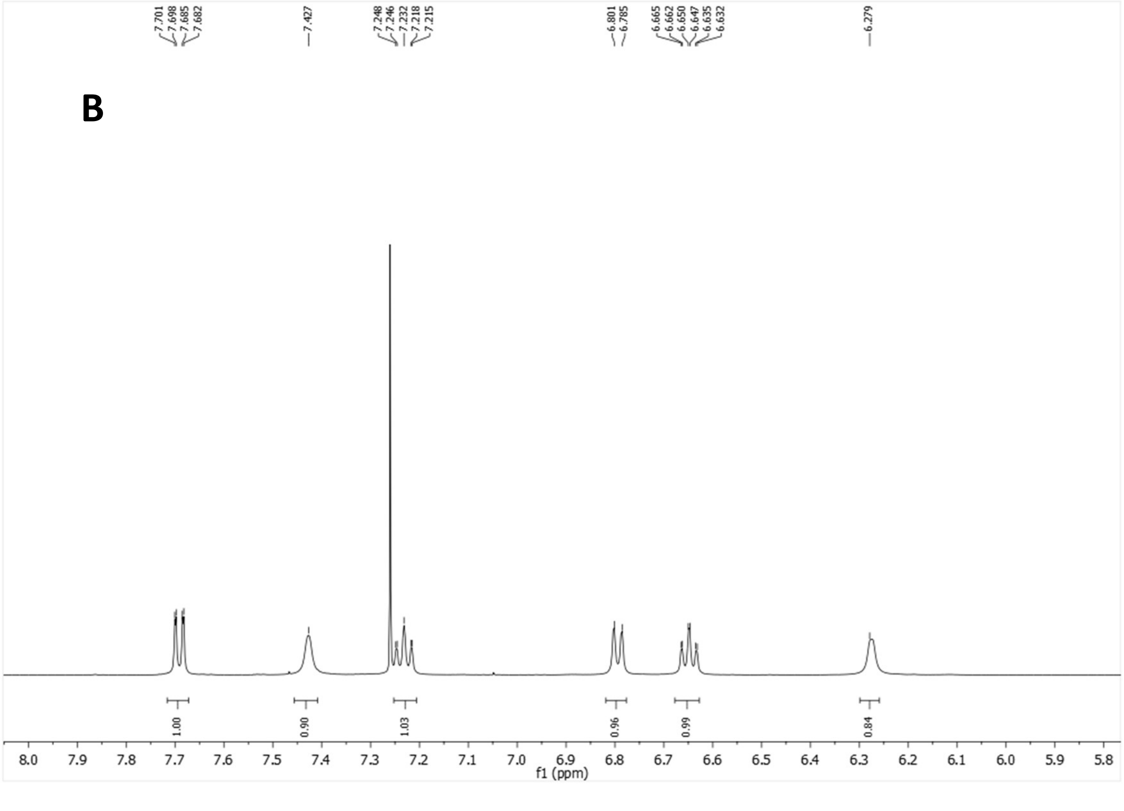

$^{13}$C-NMR (100.6 MHz, $CDCl_3$)

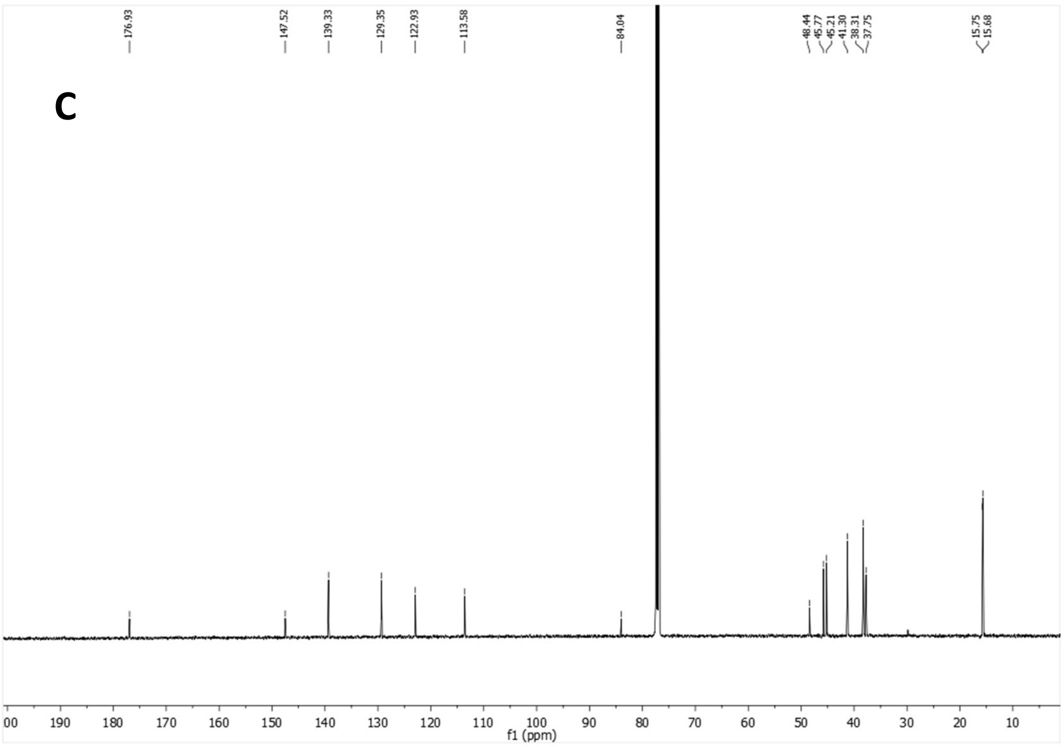


HSQC

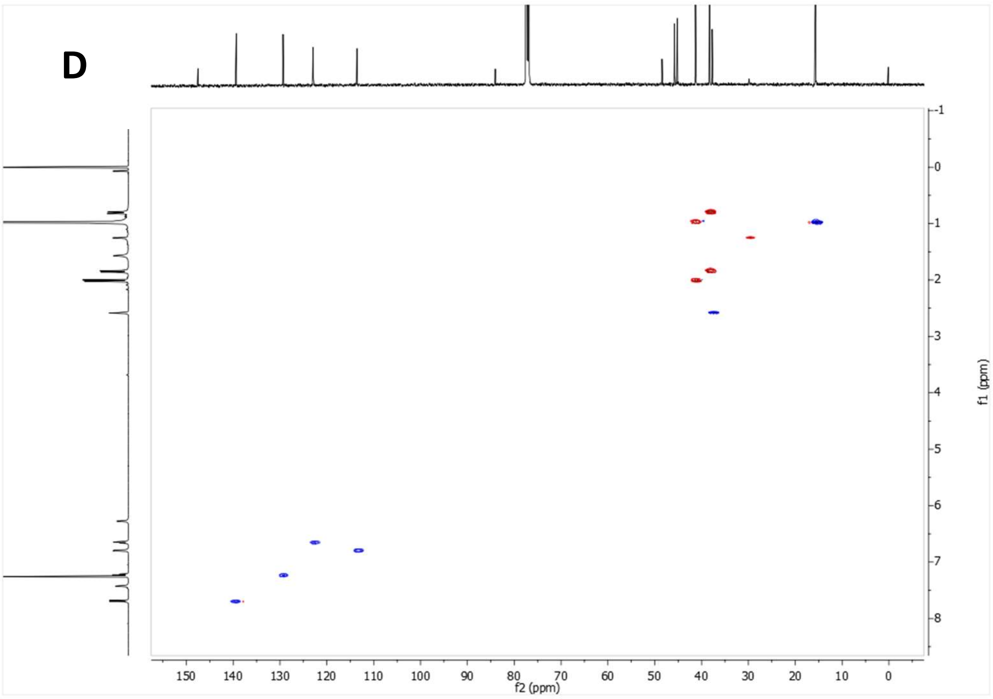


**Supplementary Figure 9:** *N'*-(2-Iodophenyl)-3,4,8,9-tetramethyltetracyclo[4.4.0.0$^{3,9}$.0$^{4,8}$]decane-1-carbohydrazide ALT-P35). (A-B) $^{1}$H NMR spectrum. (C) $^{13}$C NMR spectrum. (D) HSQC spectrum.

[1]H NMR (400 MHz, $CDCl_3$)

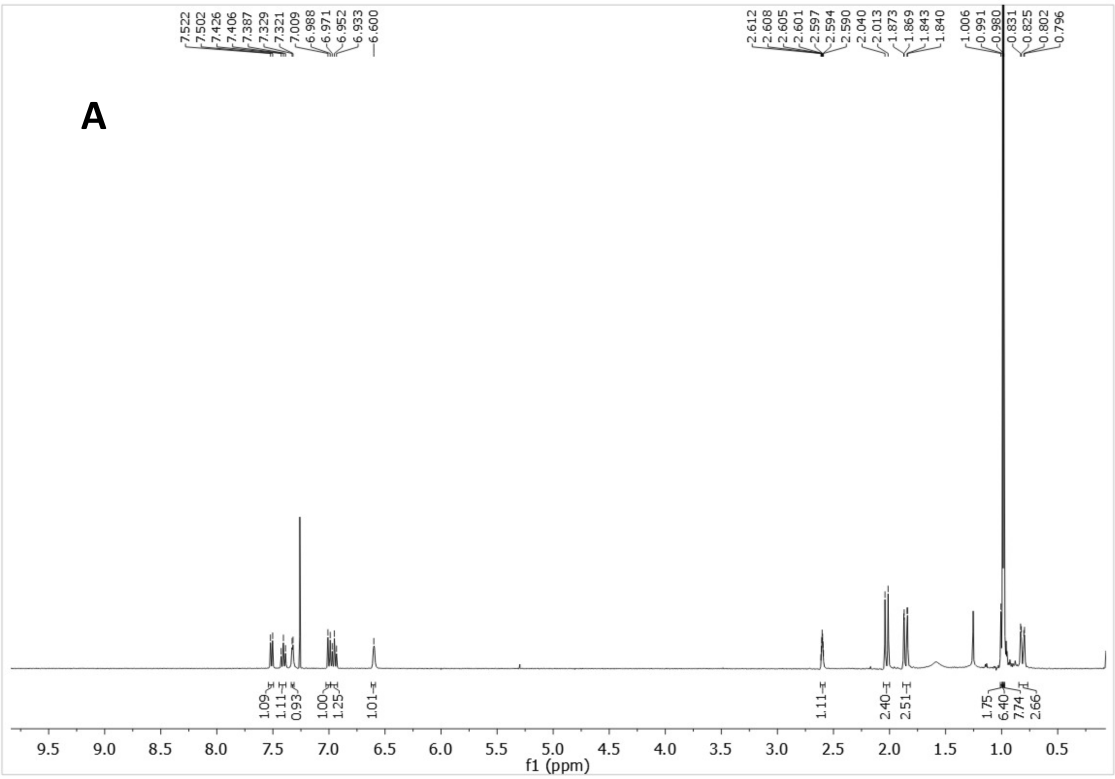

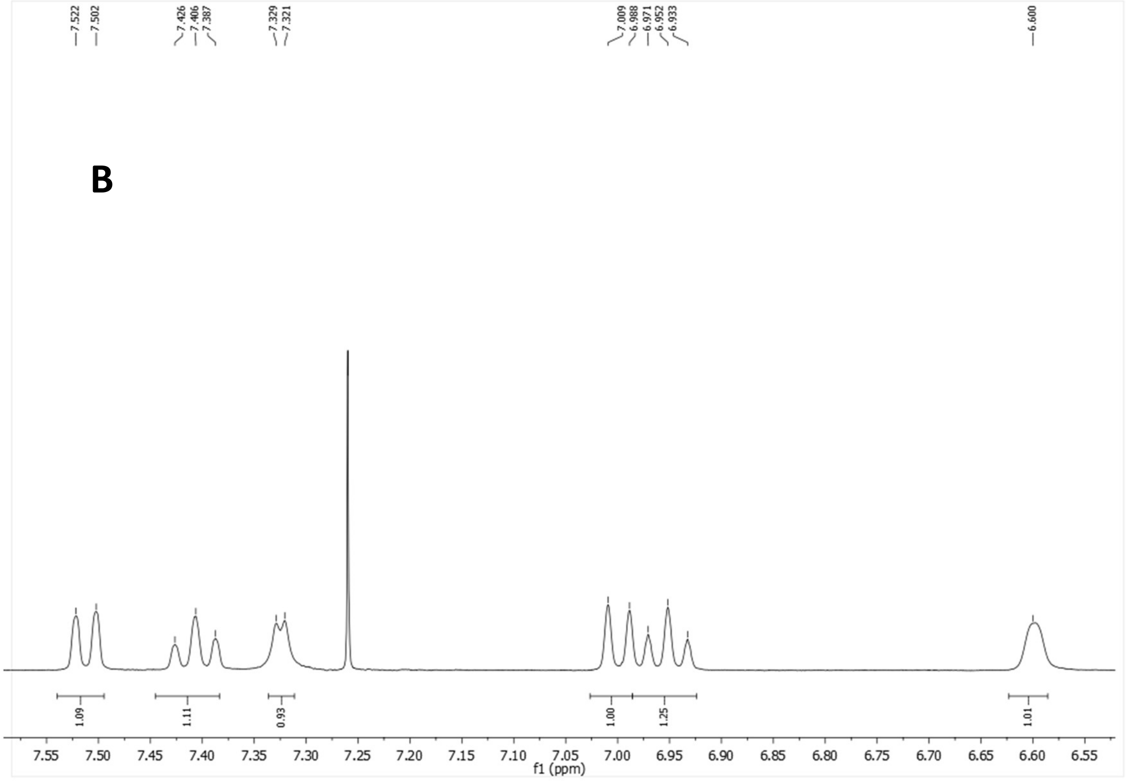

$^{13}C$-NMR (100.6 MHz, $CDCl_3$)

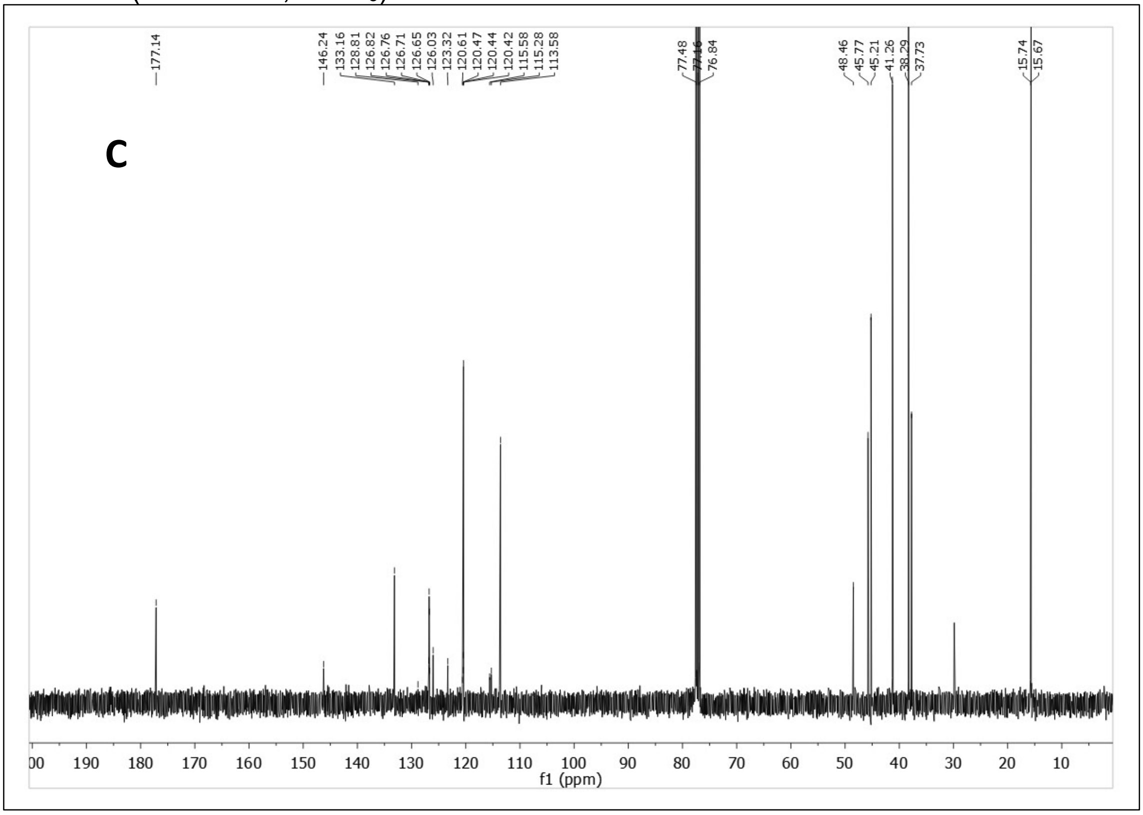


HSQC

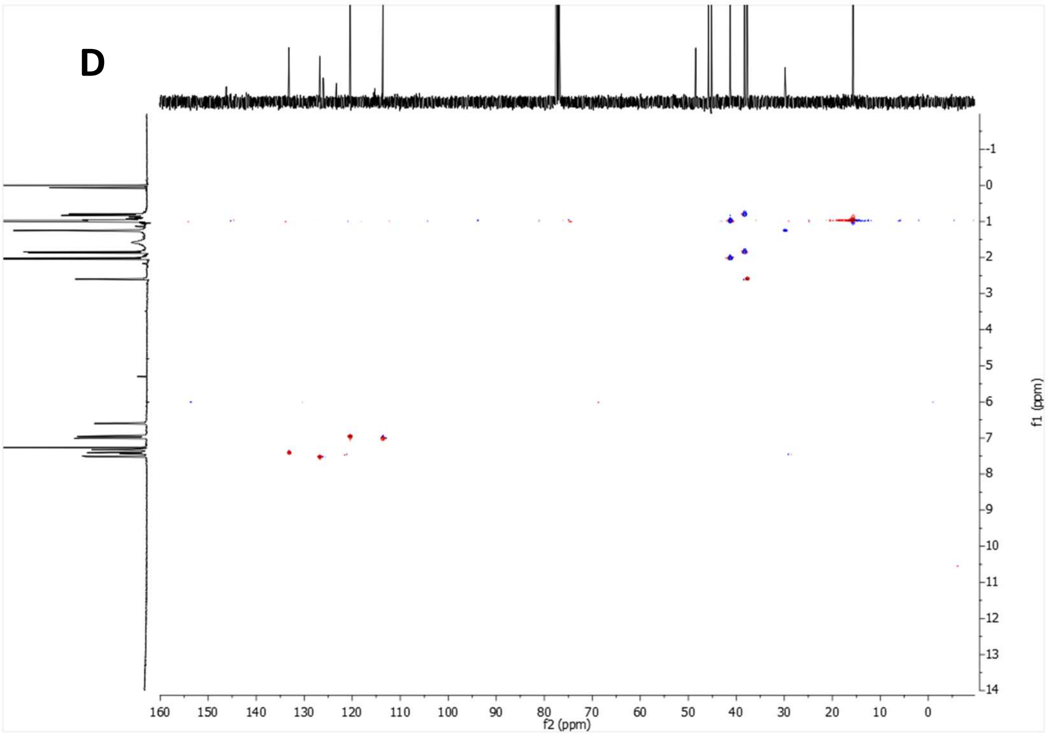


**Supplementary Figure 10:** 3,4,8,9-tetramethyl--(2-(trifluoromethyl)phenyl)tetracyclo[4.4.0.0$^{3,9}$.0$^{4,8}$]decane-1-carbohydrazide, MBX-47 (A-B) $^{1}H$ NMR spectrum. (C) $^{13}C$ NMR spectrum. (D) HSQC spectrum.

$^1H$ NMR (400 MHz, $CDCl_3$)

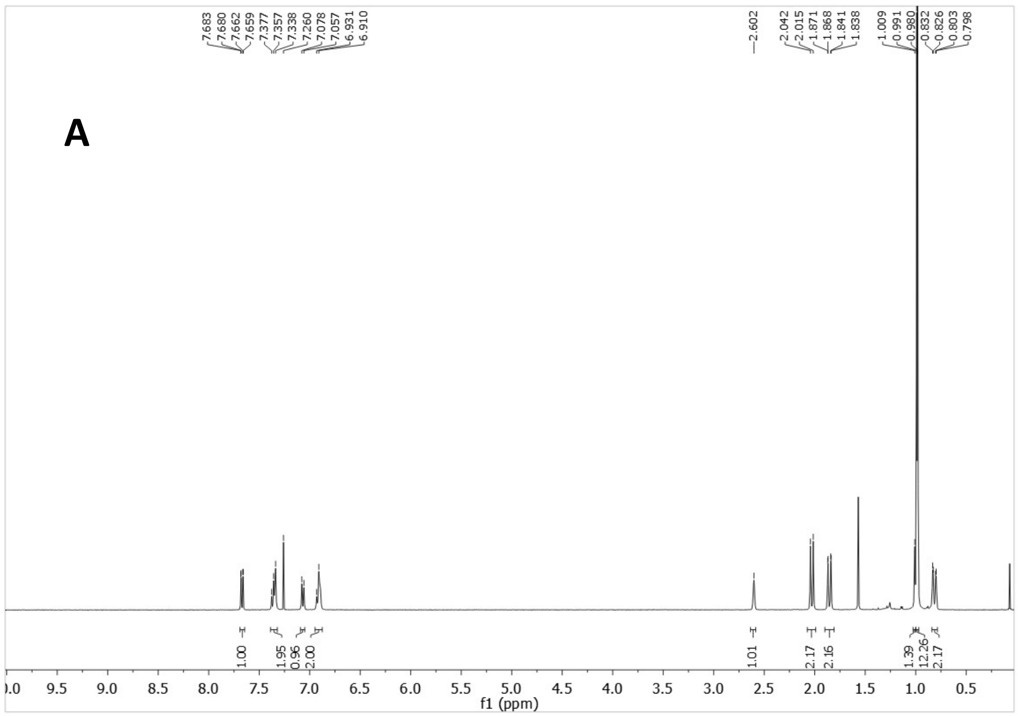

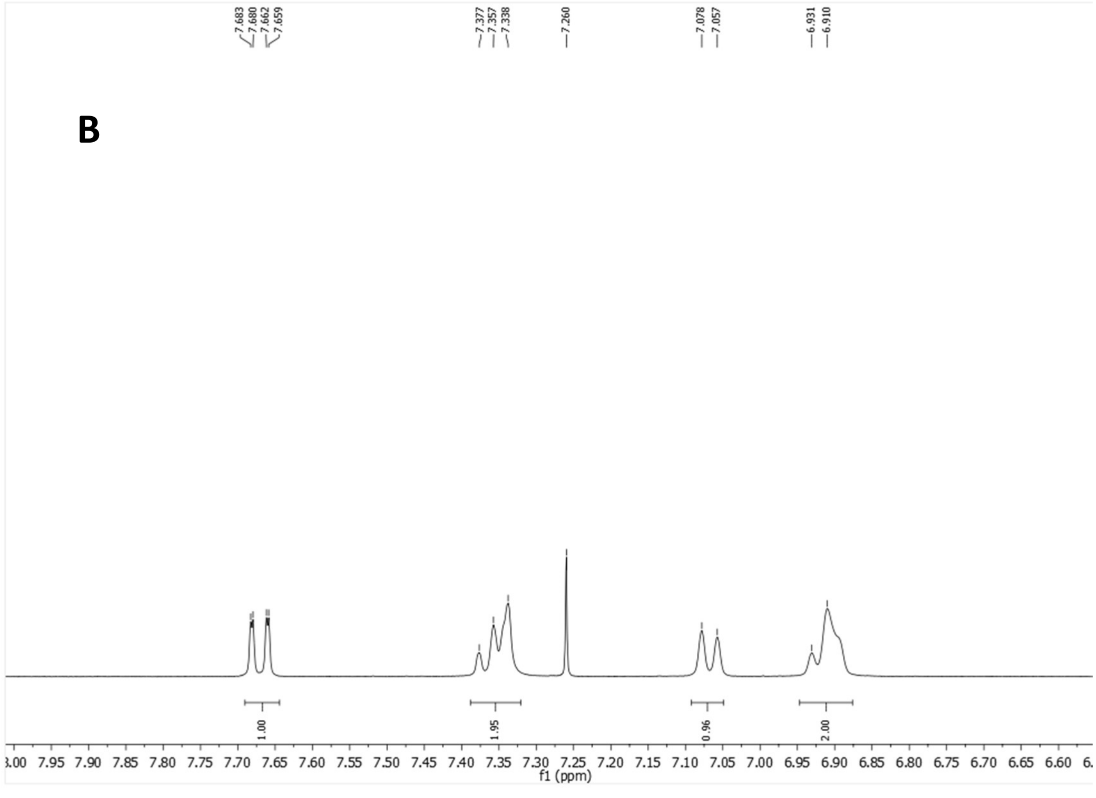

$^{13}C$-NMR (100.6 MHz, $CDCl_3$)

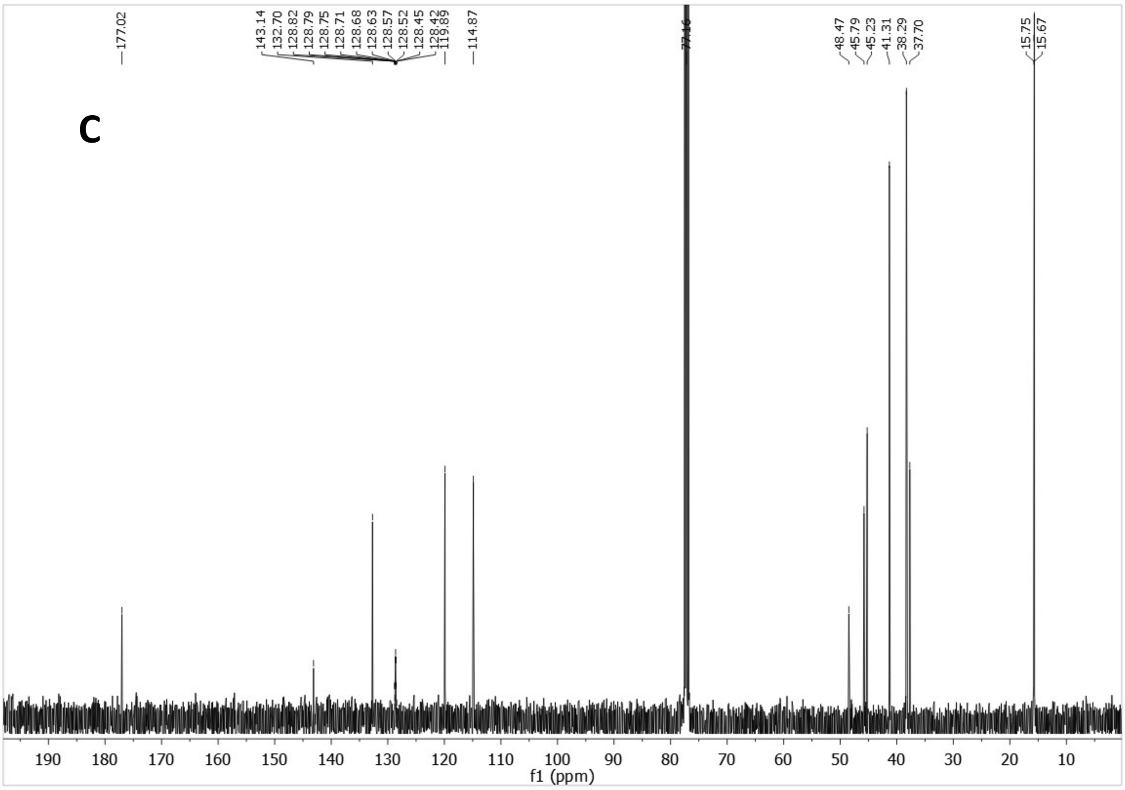


HSQC

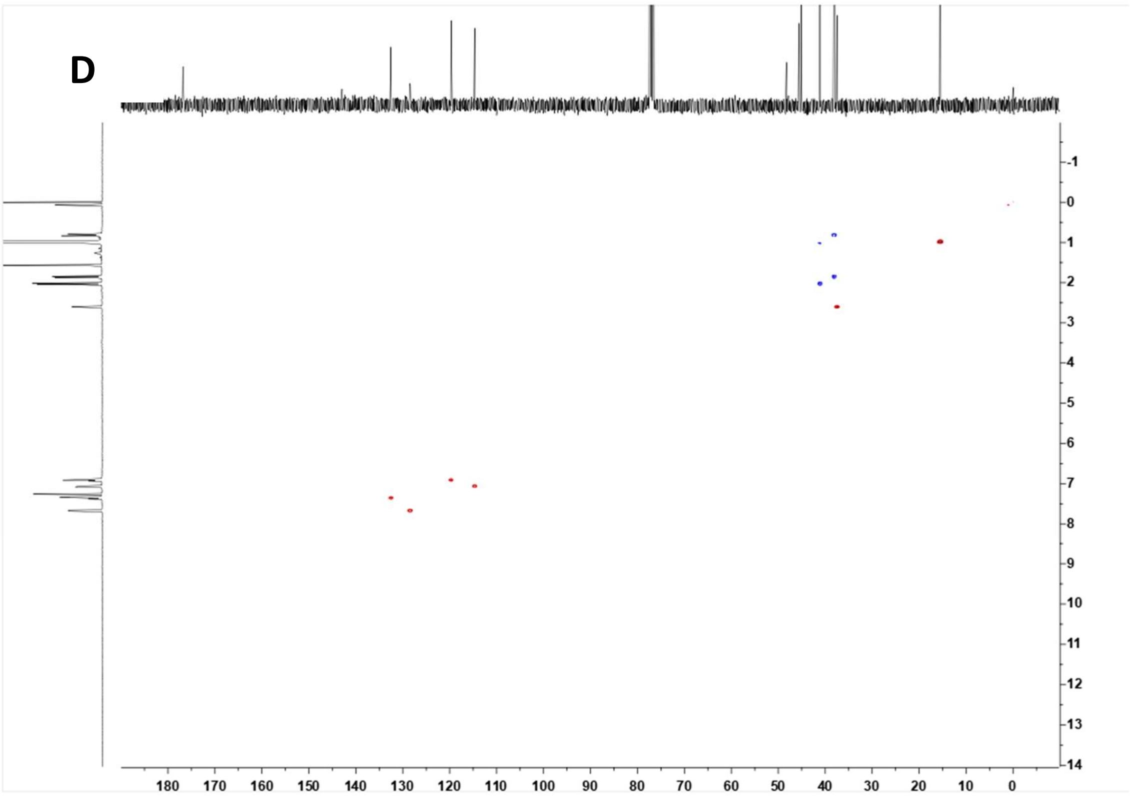

$^{19}$F-NMR (376.5 MHz, $CDCl_3$)

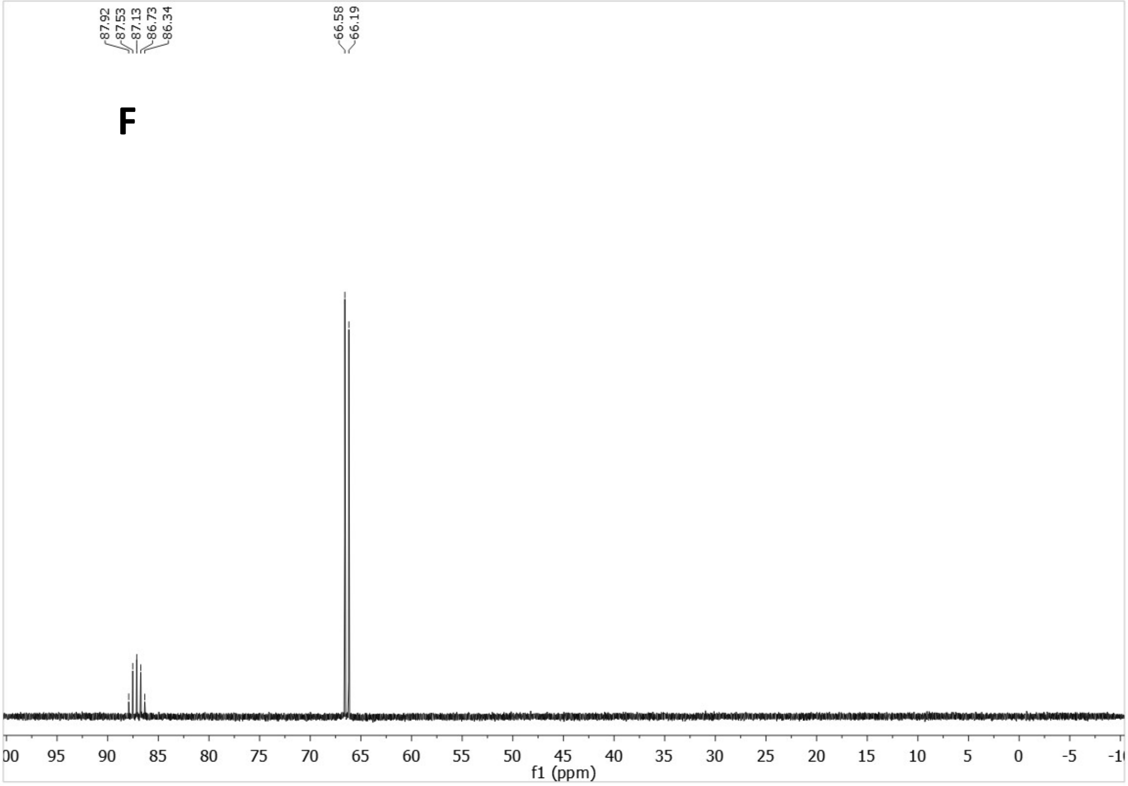


**Supplementary Figure 11:** 3,4,8,9-tetramethyl-*N′*-(2-(pentafluoro-$\lambda^6$-sulfanyl)phenyl)tetracyclo[4.4.0.0$^{3,9}$.0$^{4,8}$]decane-1-carbohydrazide, ALT-P6 (A-B) $^{1}$H NMR spectrum. (C) $^{13}$C NMR spectrum. (D) HSQC spectrum. (F) $^{19}$F spectrum.

[1]H NMR (400 MHz, DMSO-$d_6$)

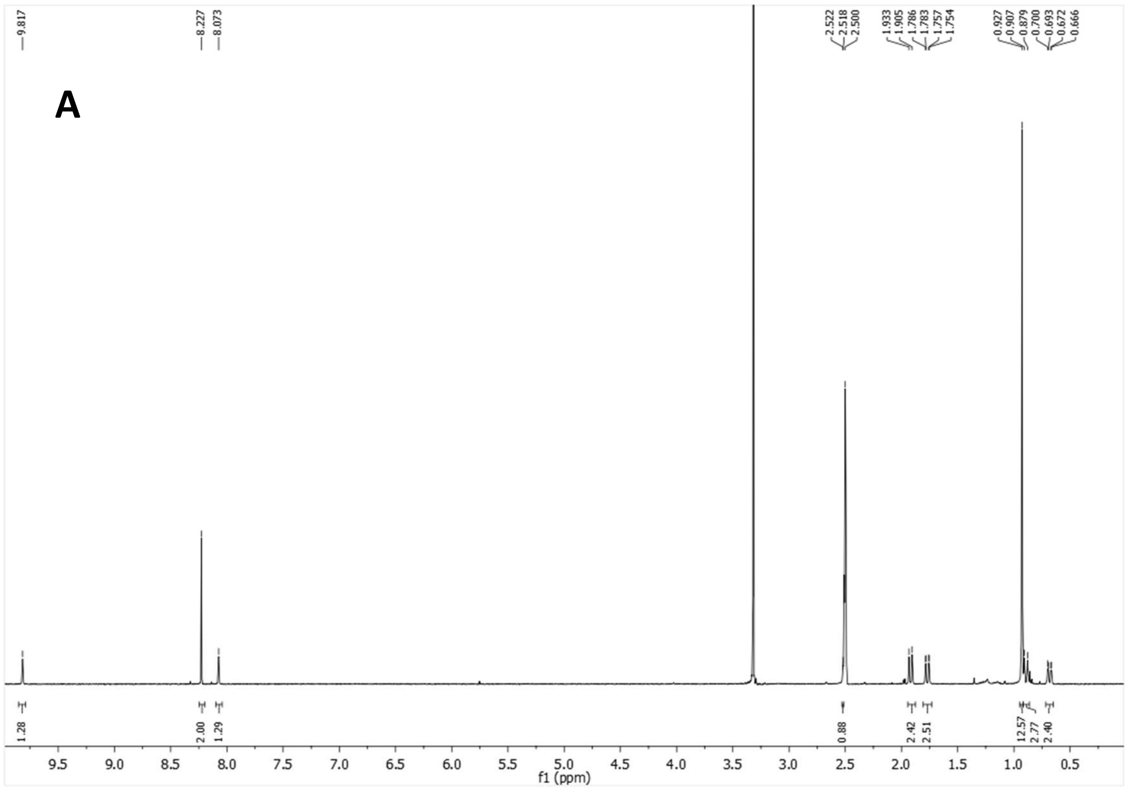


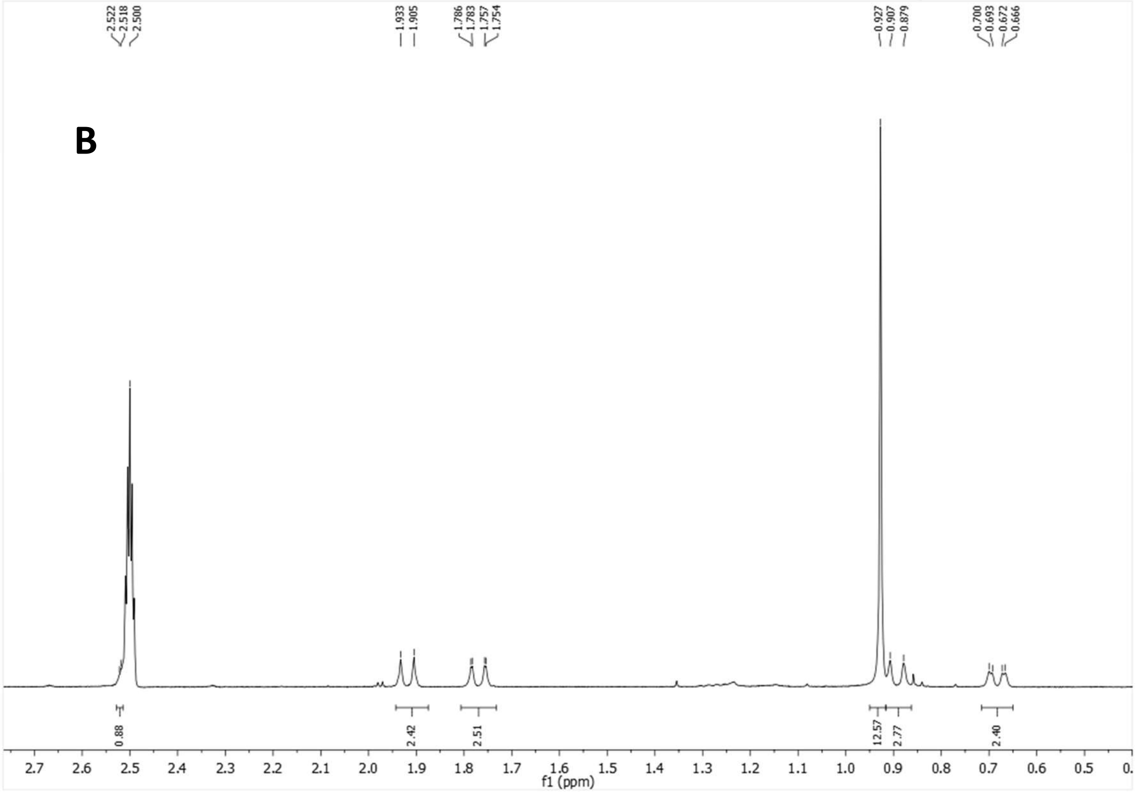

$^{13}$C-NMR (100.6 MHz, DMSO-$d_6$)

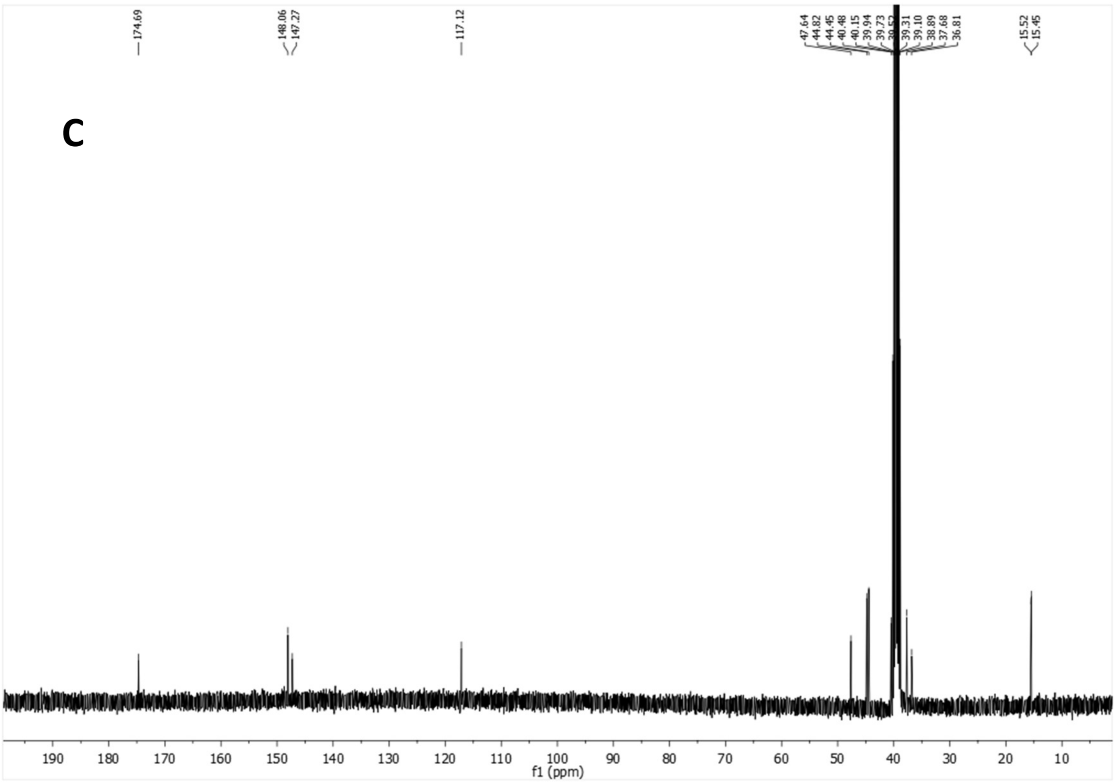


HSQC

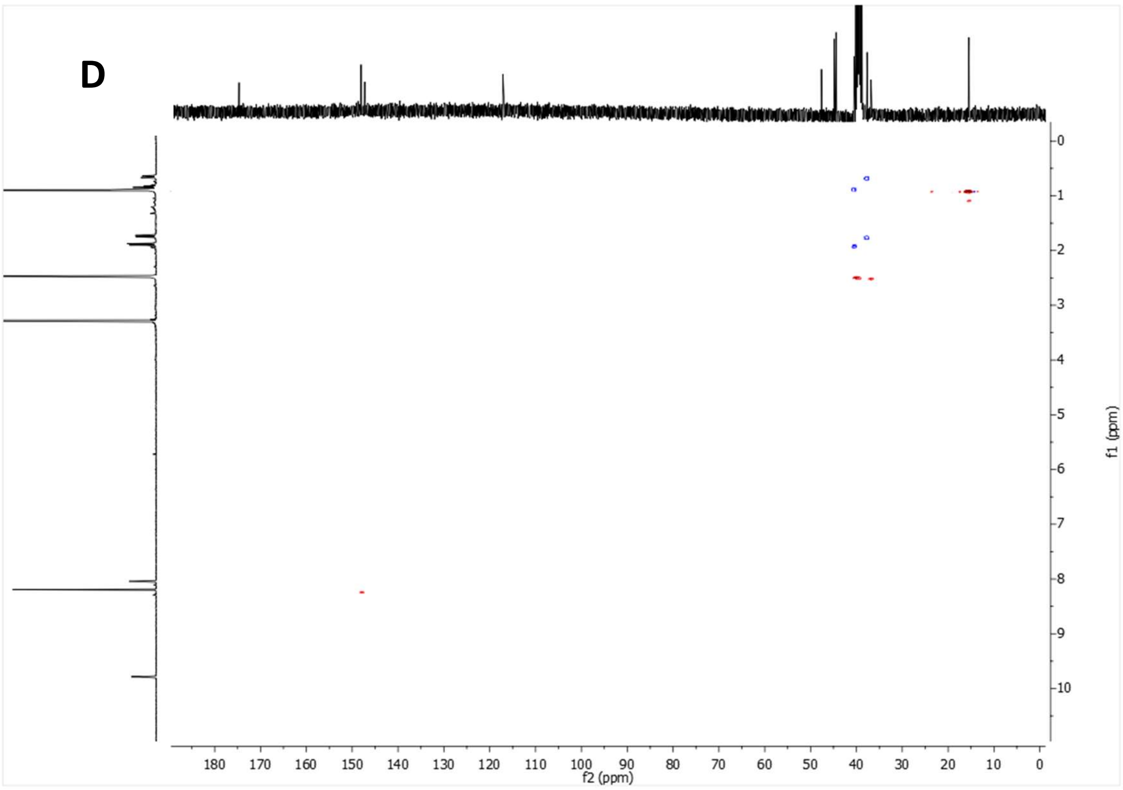


**Supplementary Figure 12:** *N'*-(3',5'-Dichloropyridin-4'-yl)-3,4,8,9-tetramethyltetracyclo[4.4.0.0$^{3,9}$.0$^{4,8}$]decane-1-carbohydrazide, ALT-P1 (A-B) $^1$H NMR spectrum. (C) $^{13}$C NMR spectrum. (D) HSQC spectrum.

$^1H$ NMR (400 MHz, $CDCl_3$)

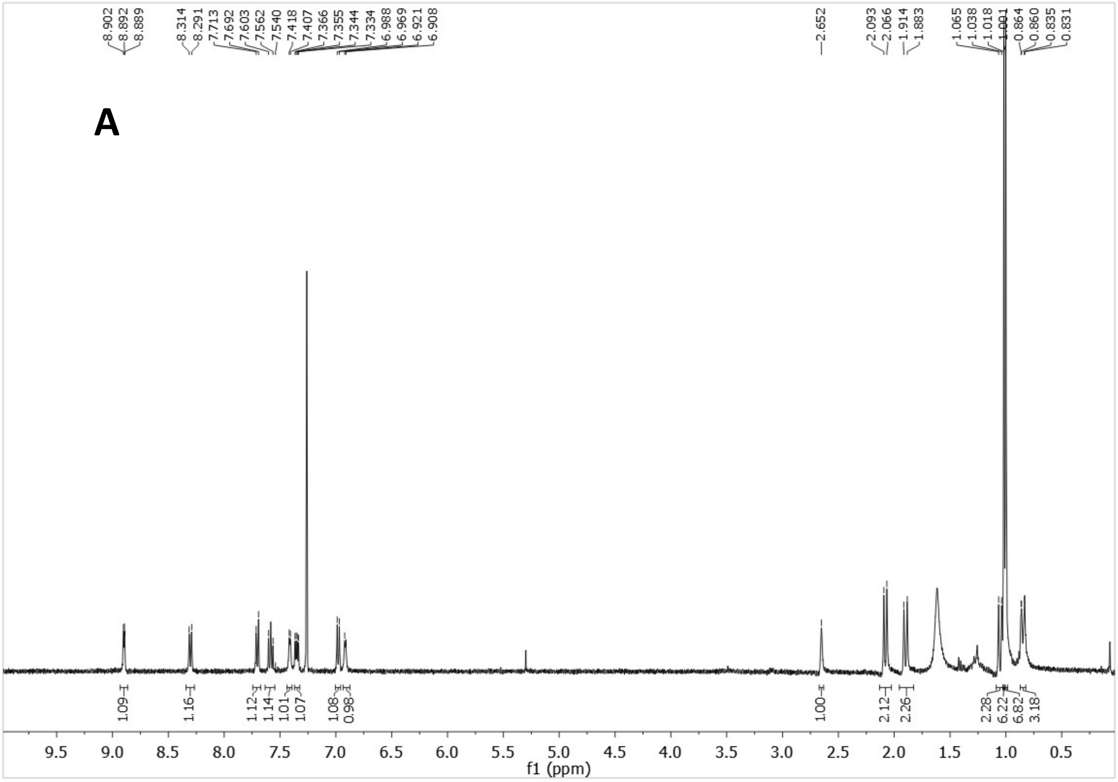


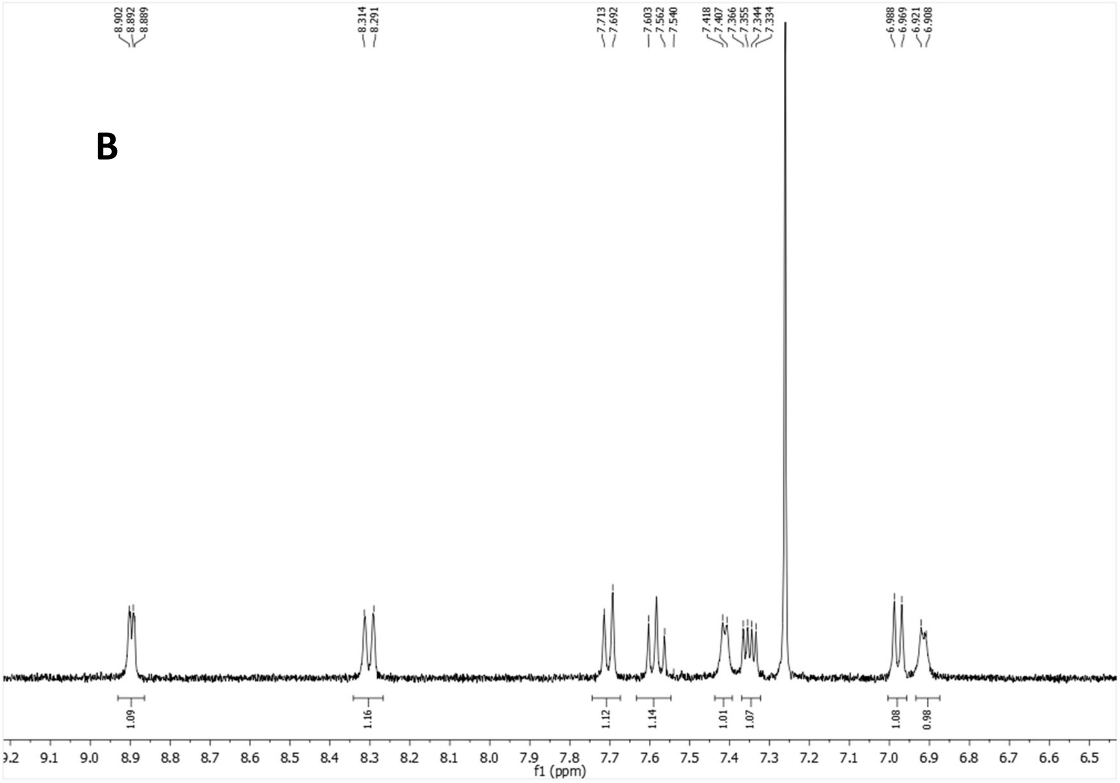

$^{13}$C-NMR (100.6 MHz, $CDCl_3$)

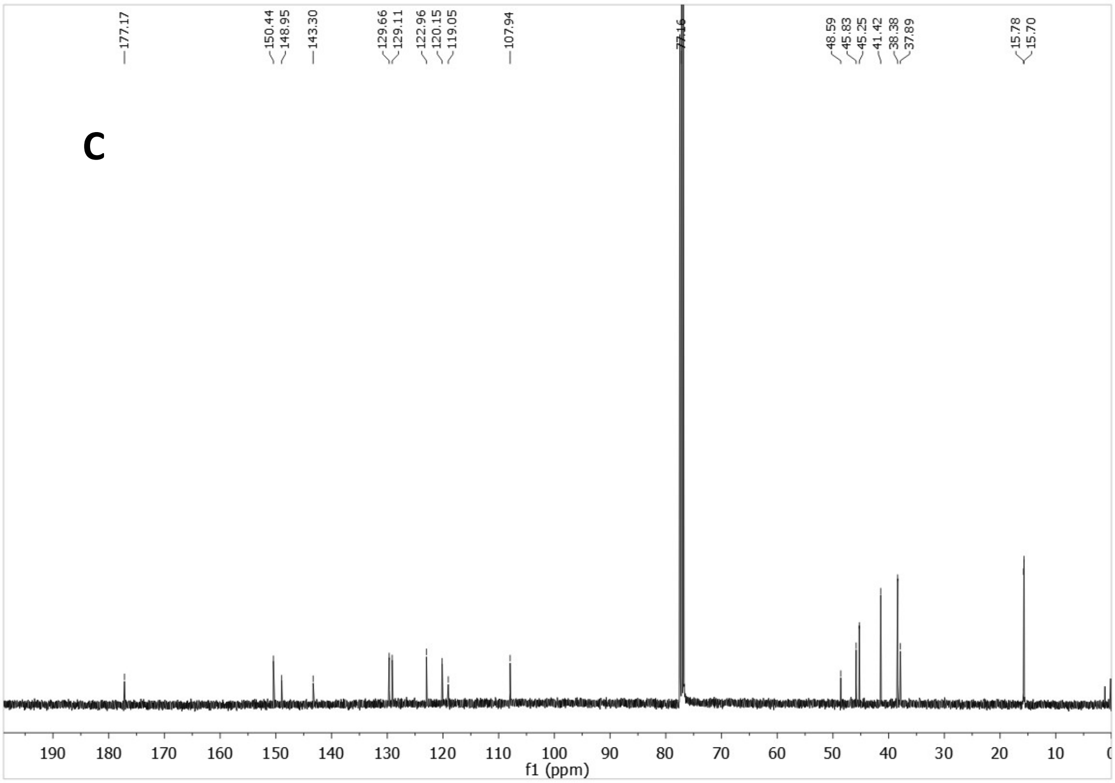


HSQC

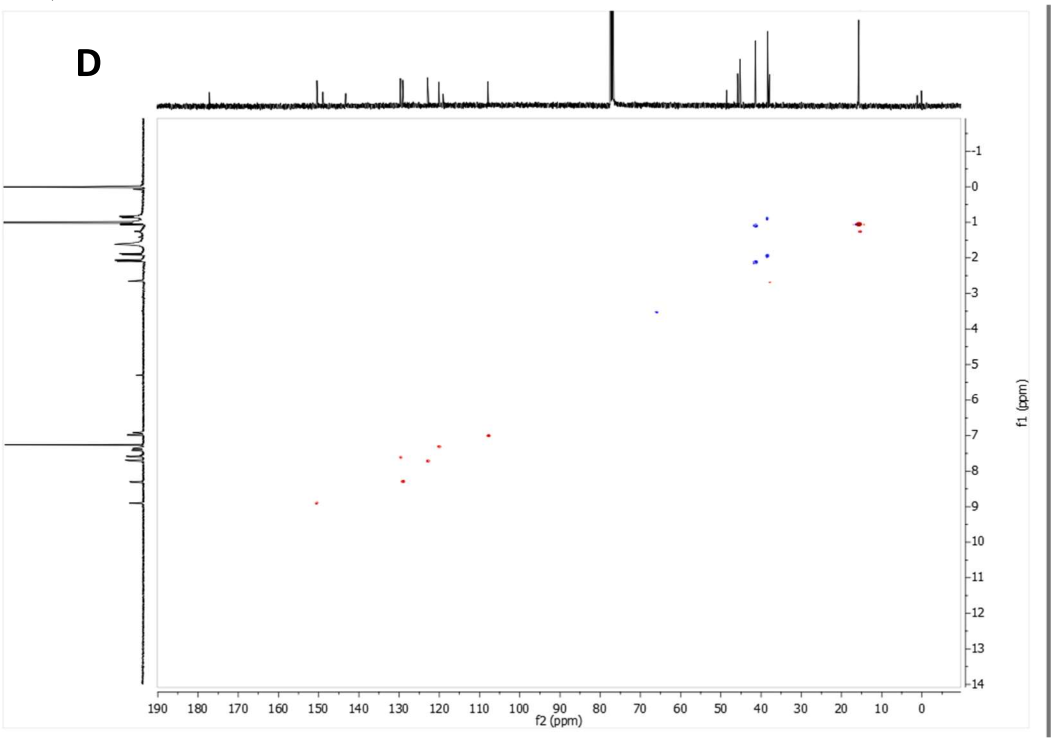


**Supplementary Figure 13:** 3,4,8,9-tetramethyl-*N'*-(quinoline-5'-yl)tetracyclo[4.4.0.0$^{3,9}$0$^{4,8}$]decane-1-carbohydrazide, ALT-P2 (A-B) $^{1}$H NMR spectrum. (C) $^{13}$C NMR spectrum. (D) HSQC spectrum.

[1]H NMR (400 MHz, $CDCl_3$)

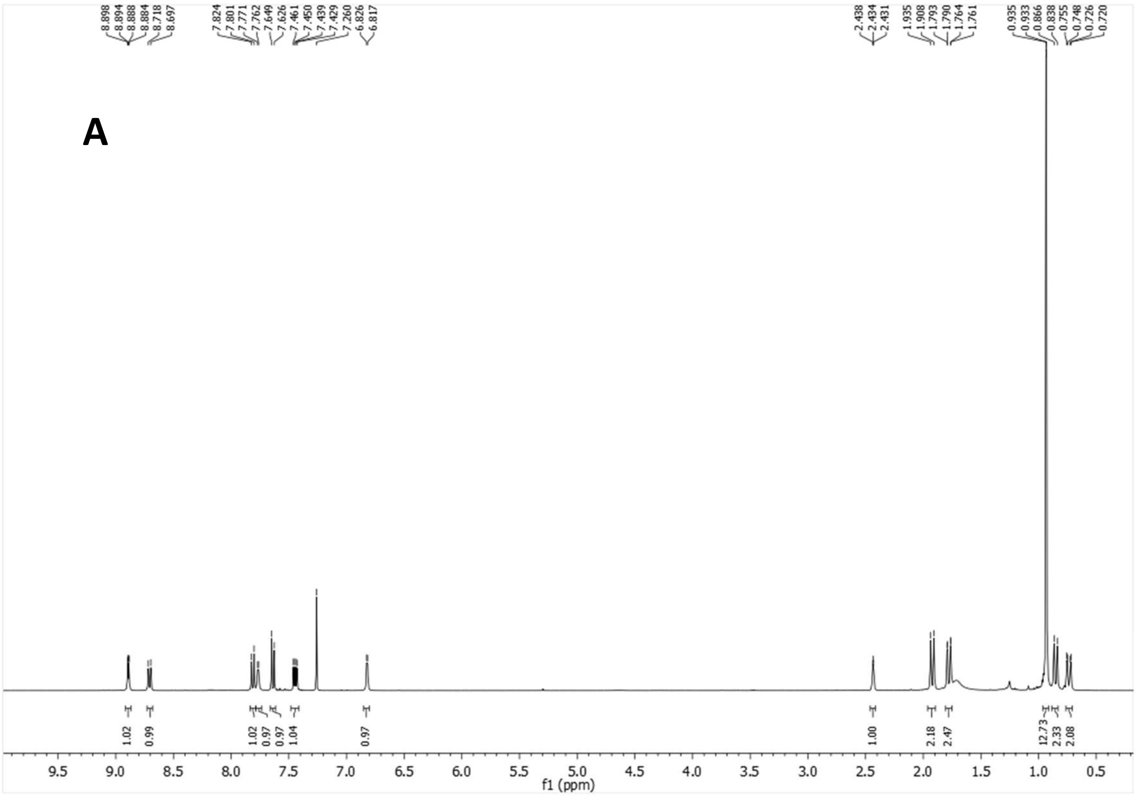

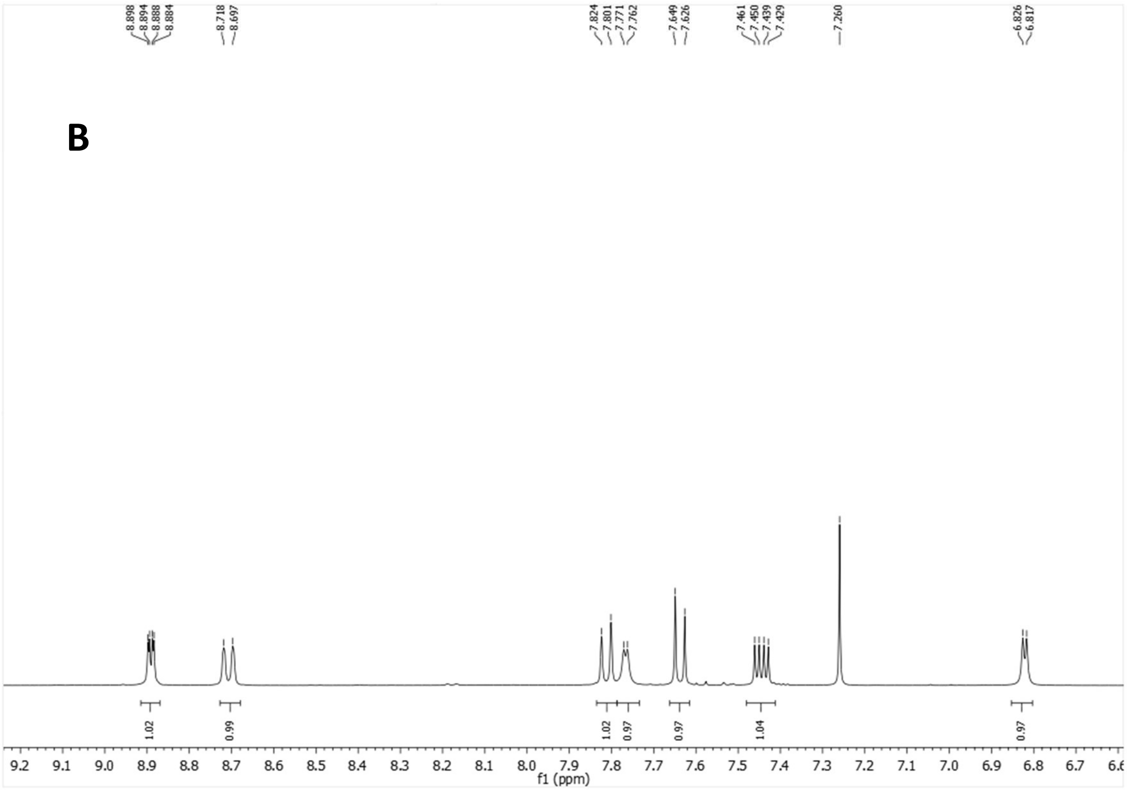

$^{13}$C-NMR (100.6 MHz, $CDCl_3$)

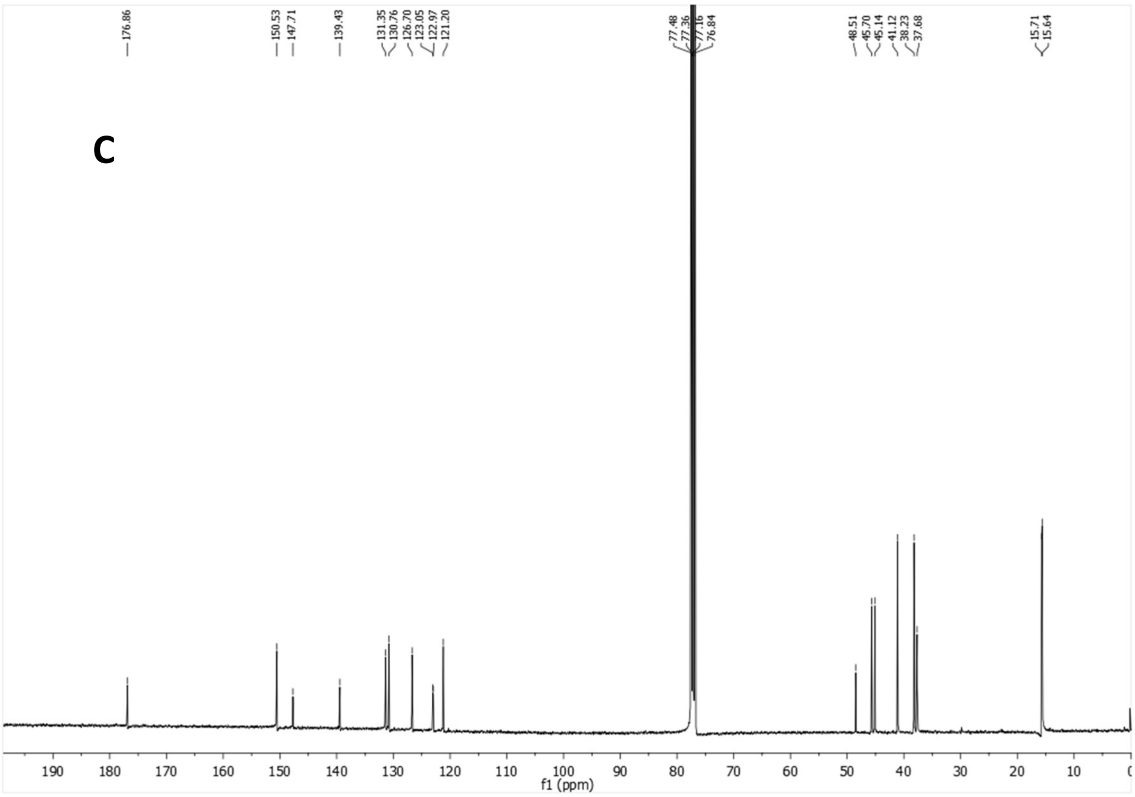


HSQC

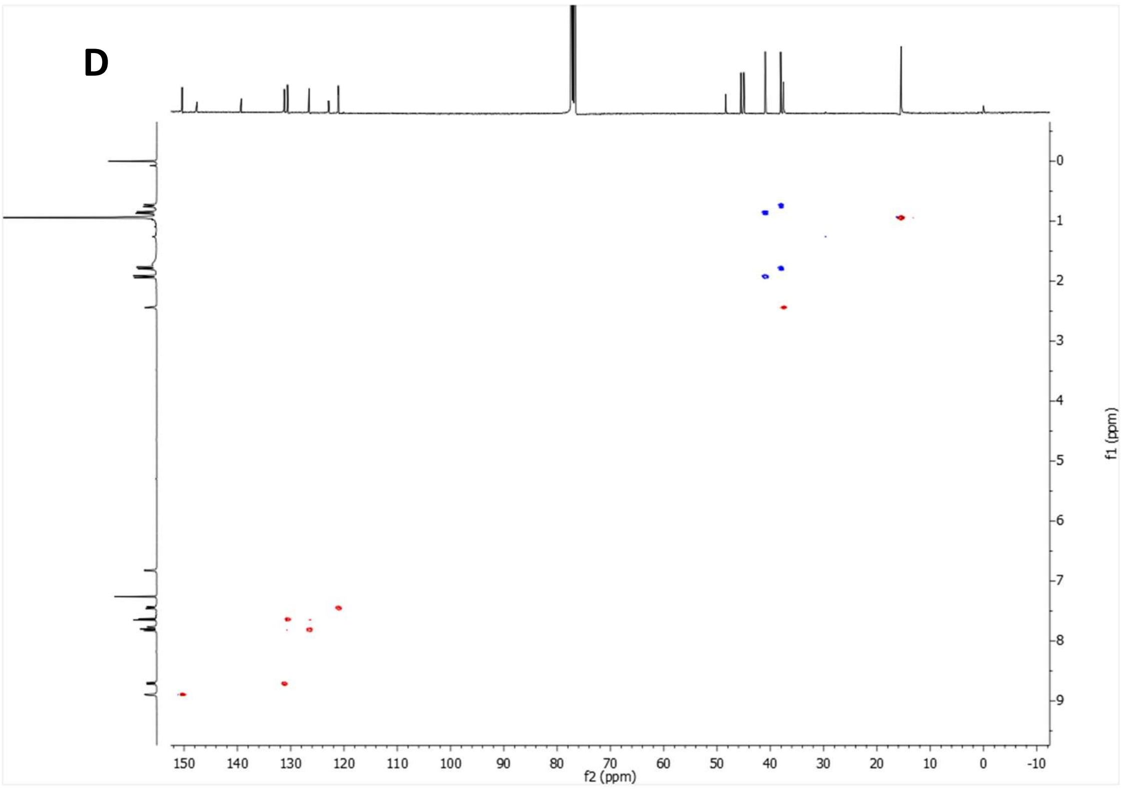


**Supplementary Figure 14:** *N′*-(6-chloroquinolin-5-yl)-3,4,8,9-tetramethyltetracyclo[4.4.0.0$^{3,9}$.0$^{4,8}$]decane-1-carbohydrazide, ALT-P21 (A-B) $^{1}$H NMR spectrum. (C) $^{13}$C NMR spectrum. (D) HSQC spectrum.

$^1H$ NMR (400 MHz, $CDCl_3$)

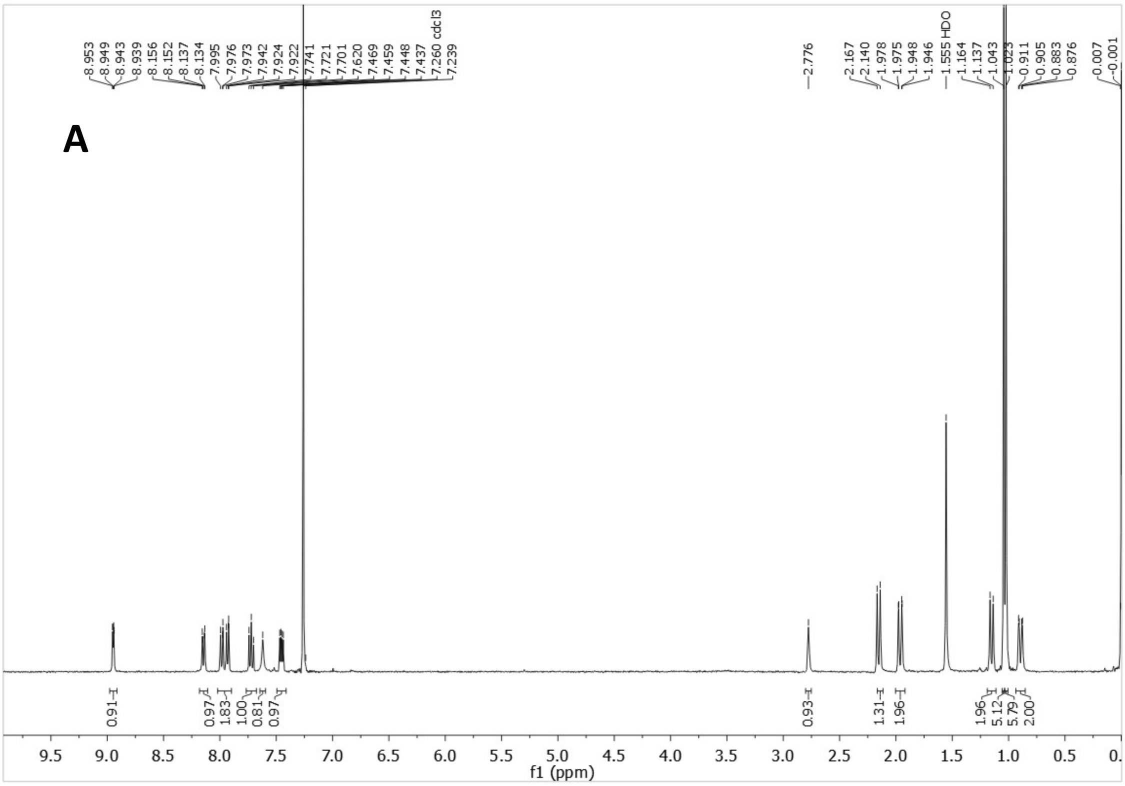


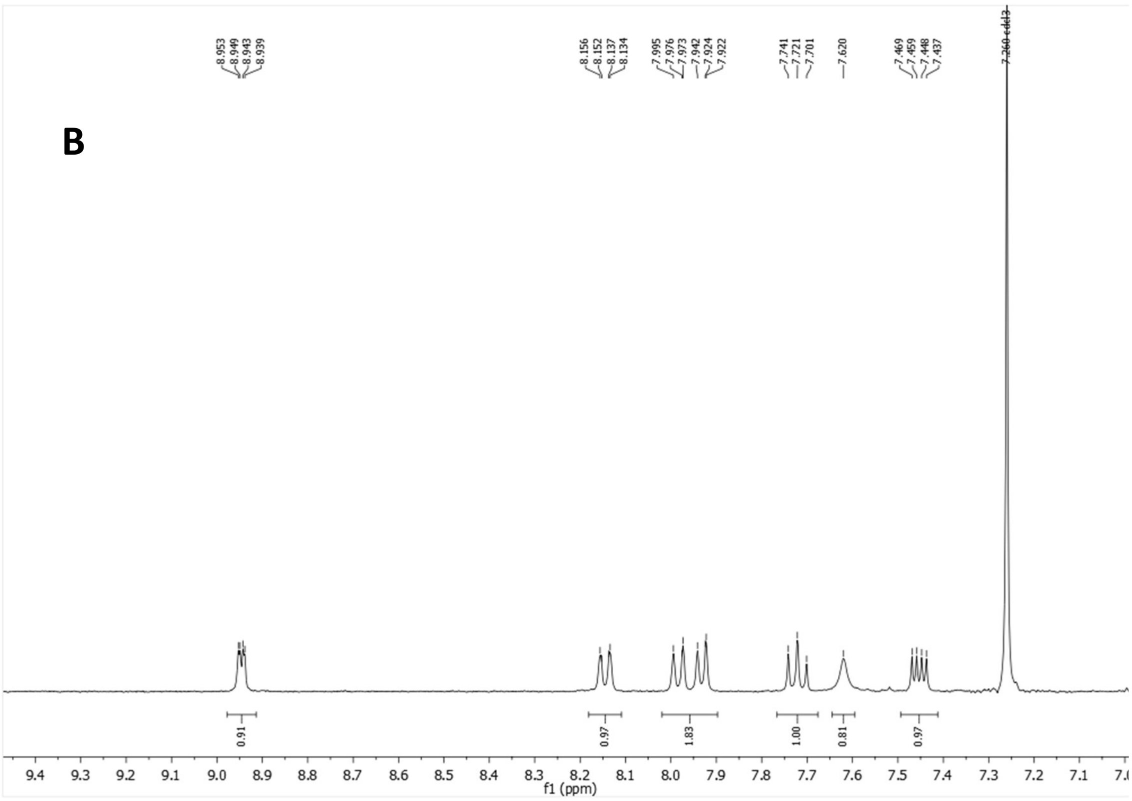

$^{13}C$-NMR (100.6 MHz, $CDCl_3$)

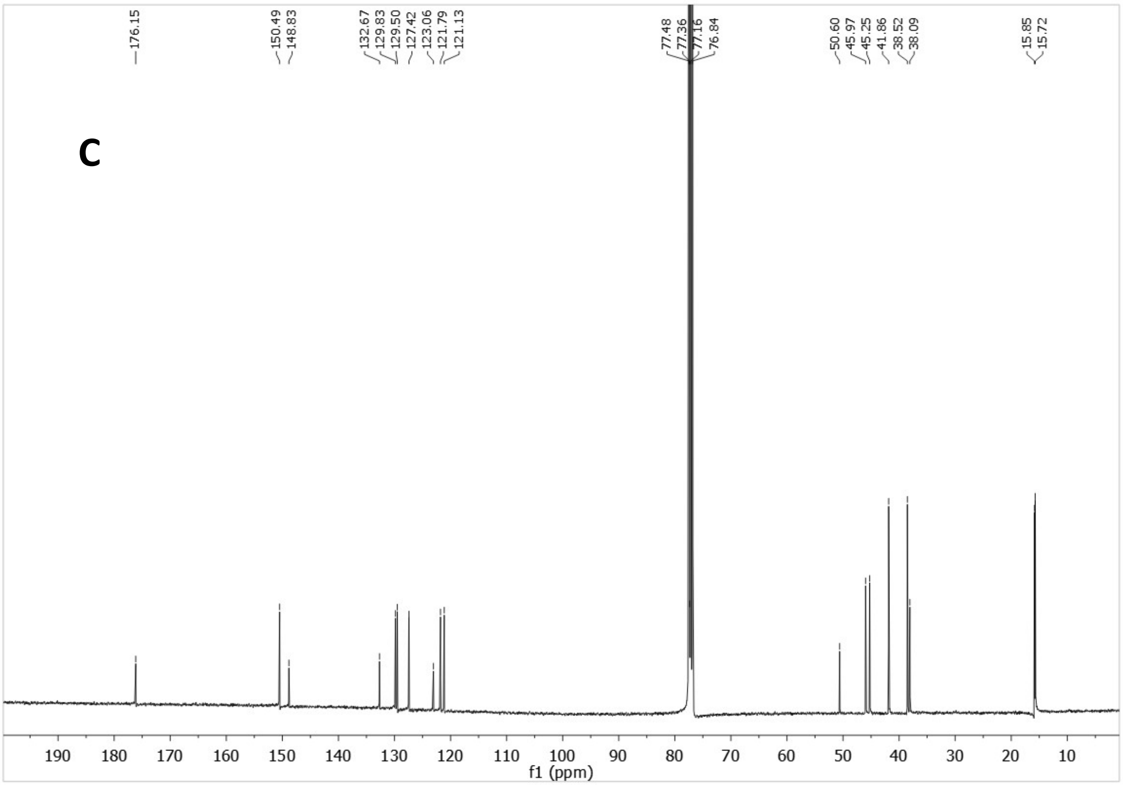


HSQC

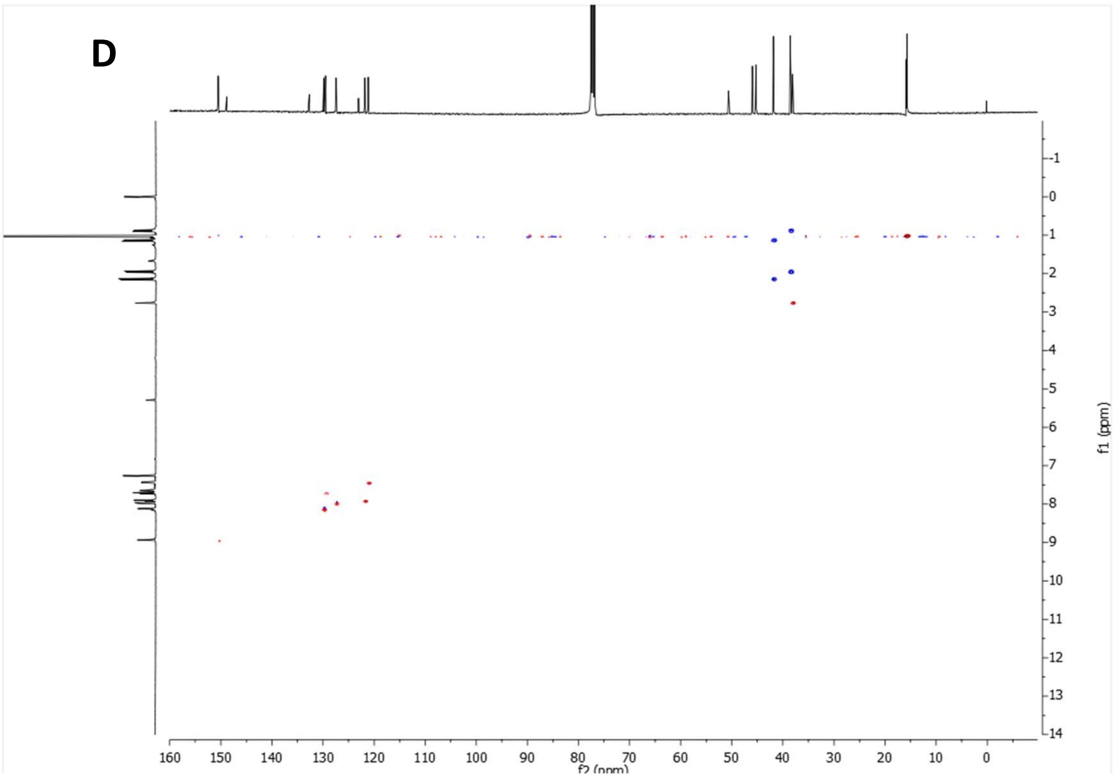


**Supplementary Figure 15:** 3,4,8,9-tetramethyl-*N*-(quinolin-5-yl)tetracyclo[4.4.0.0$^{3,9}$.0$^{4,8}$]decane-1-carboxamide, ALT-P19 (A-B) $^{1}H$ NMR spectrum. (C) $^{13}C$ NMR spectrum. (D) HSQC spectrum.

<sup></sup>

$^1$H NMR (400 MHz, $CDCl_3$)

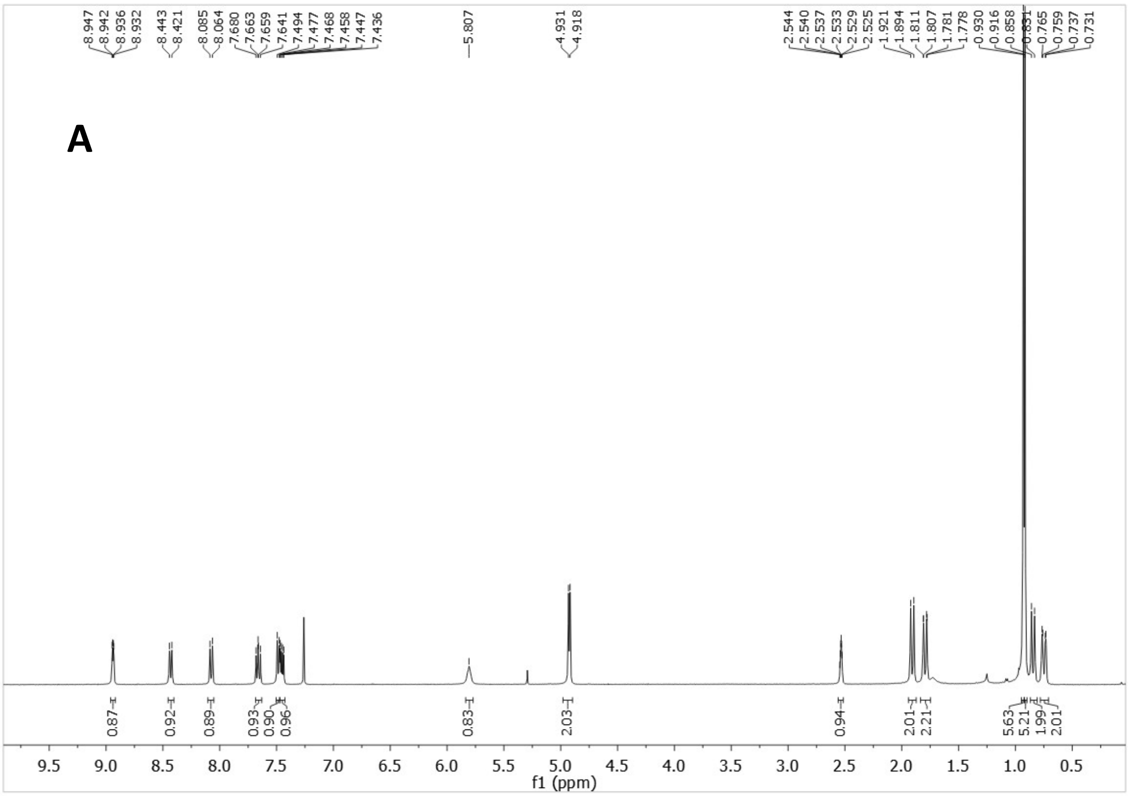


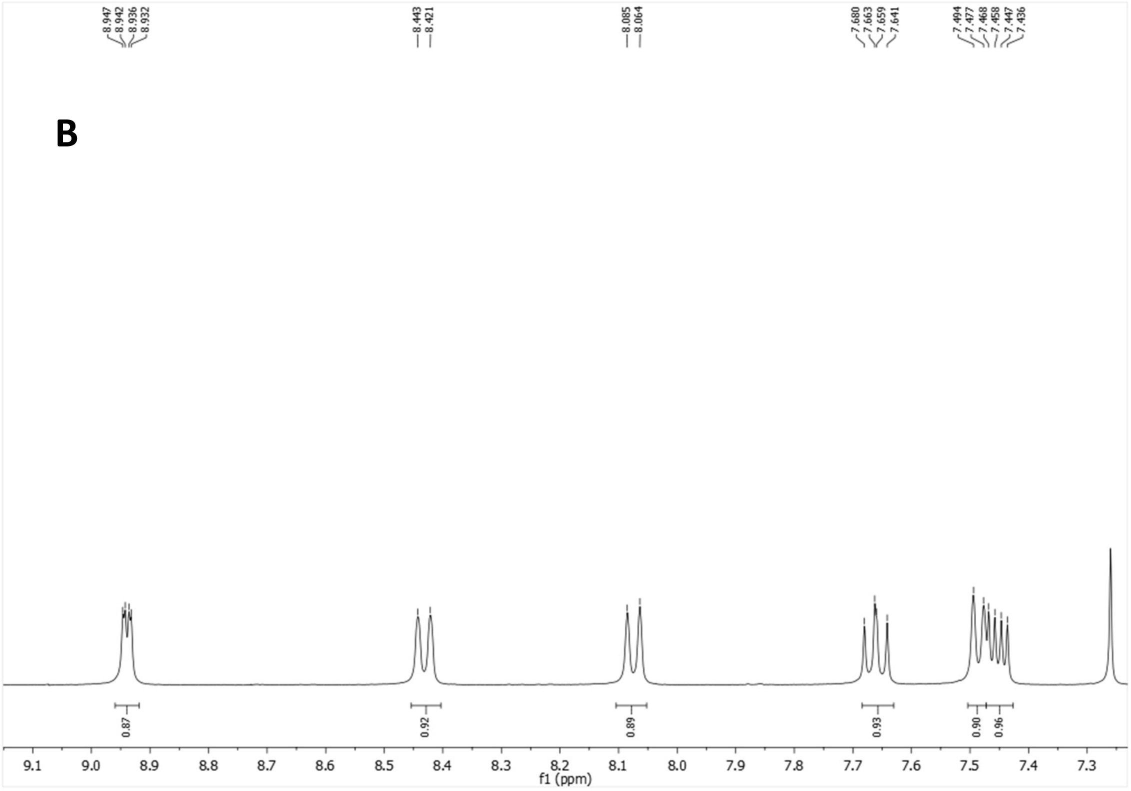

$^{13}$C-NMR (100.6 MHz, $CDCl_3$)

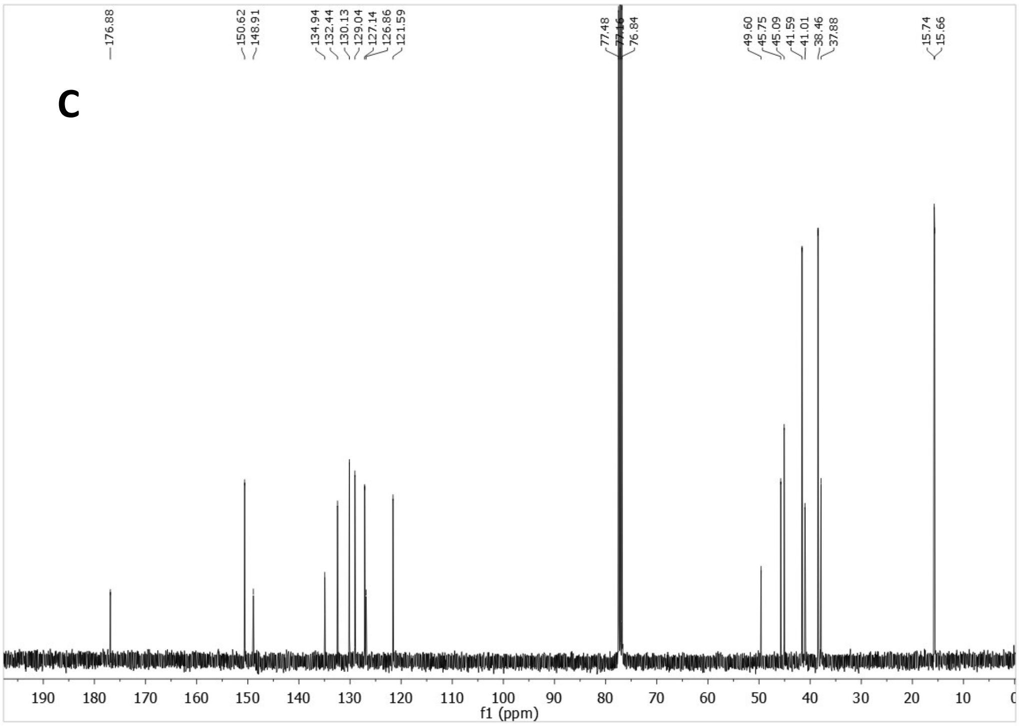


HSQC

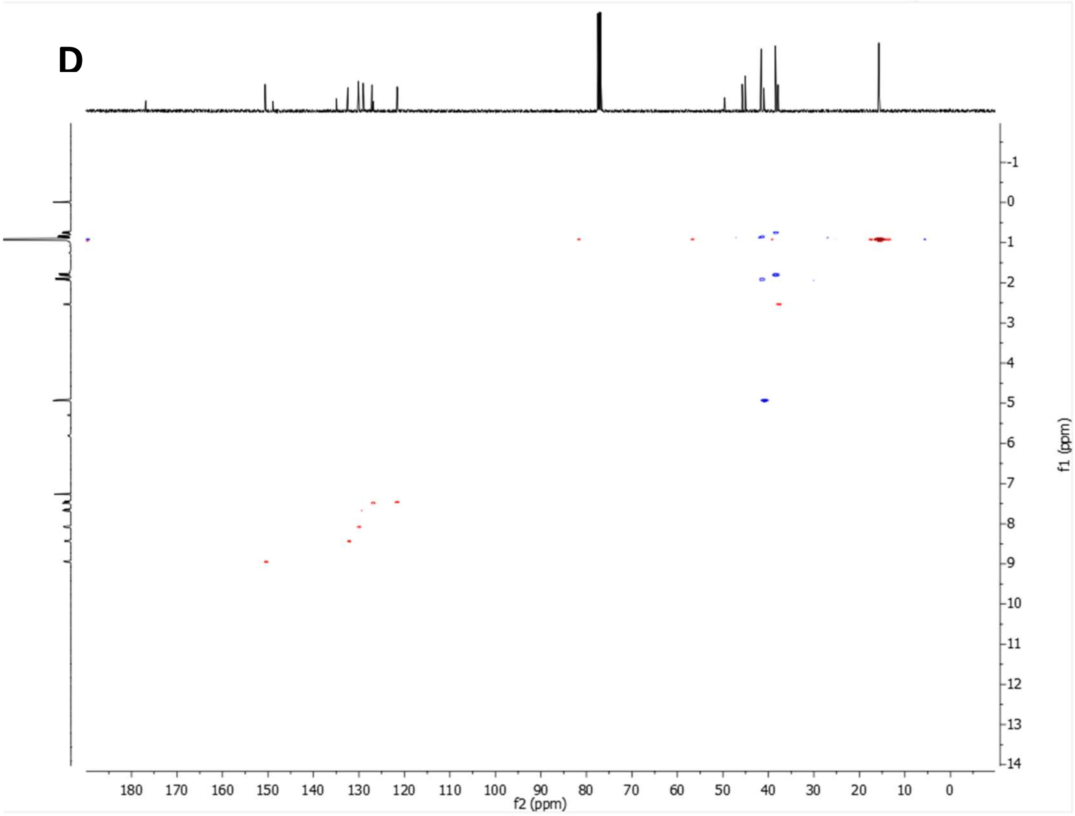


**Supplementary Figure 16:** 3,4,8,9-tetramethyl-*N*-(quinolin-5-ylmethyl)tetracyclo[4.4.0.0$^{3,9}$.0$^{4,8}$]decane-1-carboxamide, P20 (A-B) $^{1}$H NMR spectrum. (C) $^{13}$C NMR spectrum. (D) HSQC spectrum.

[1]H NMR (400 MHz, $CDCl_3$)

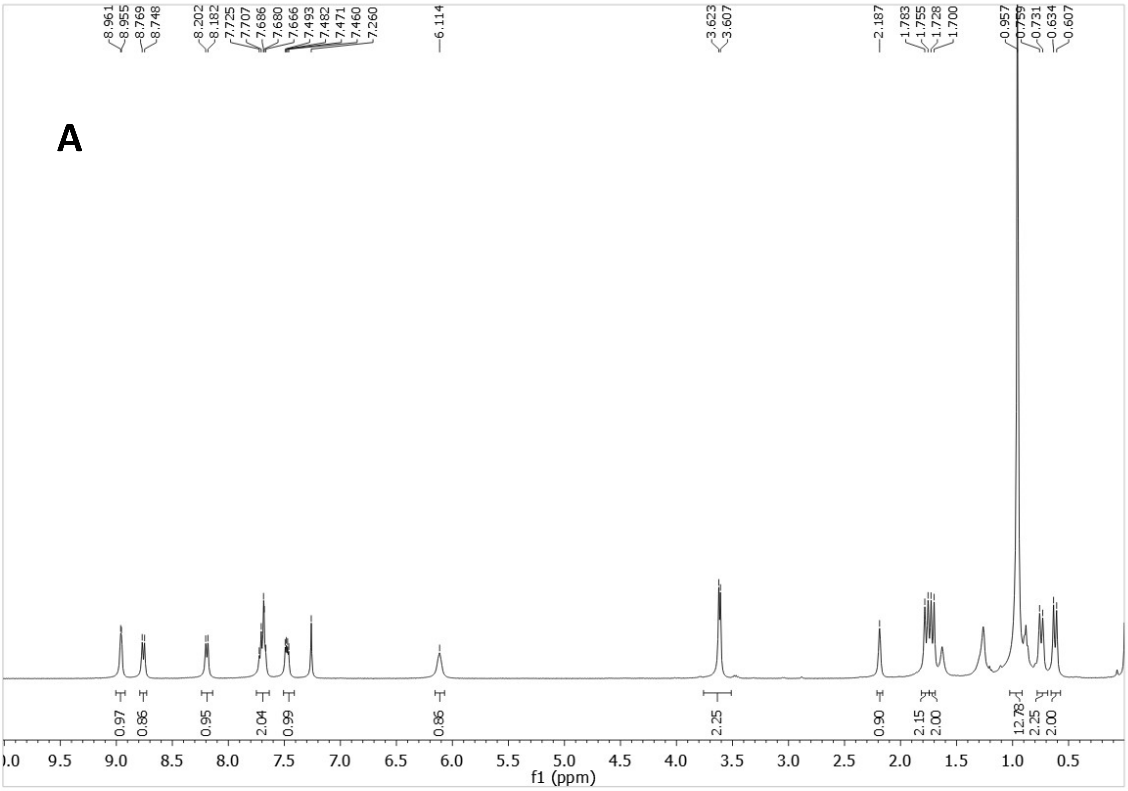


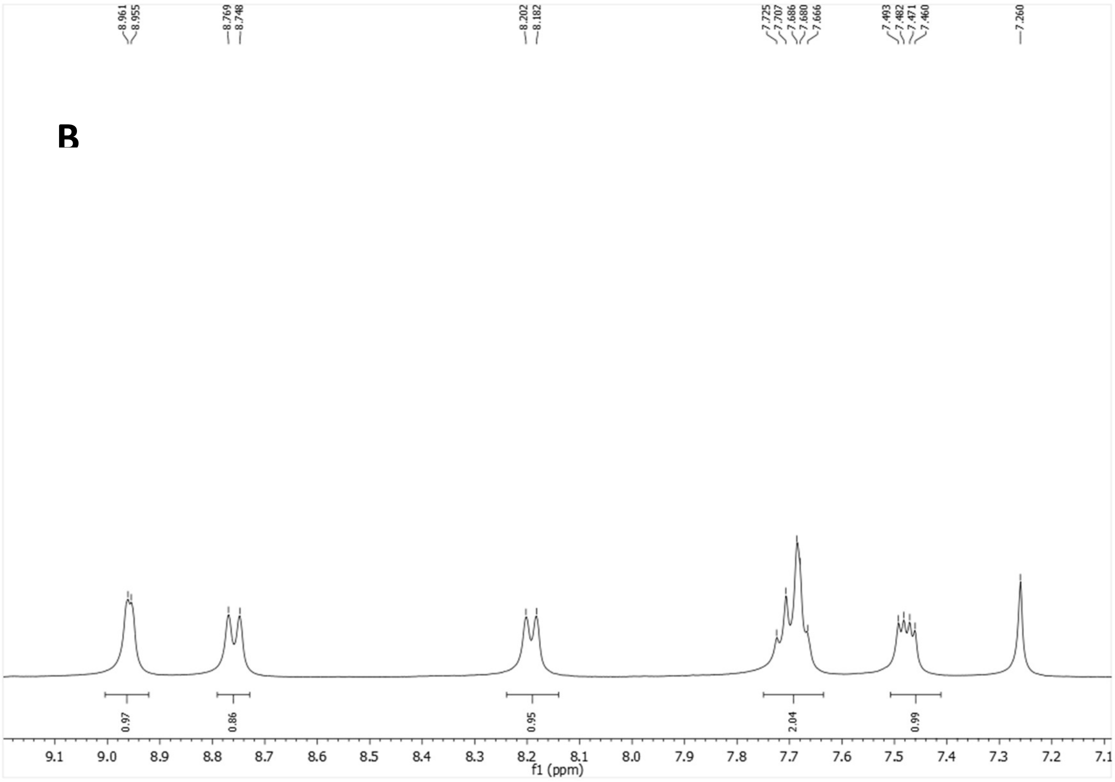

$^{13}$C-NMR (100.6 MHz, $CDCl_3$)

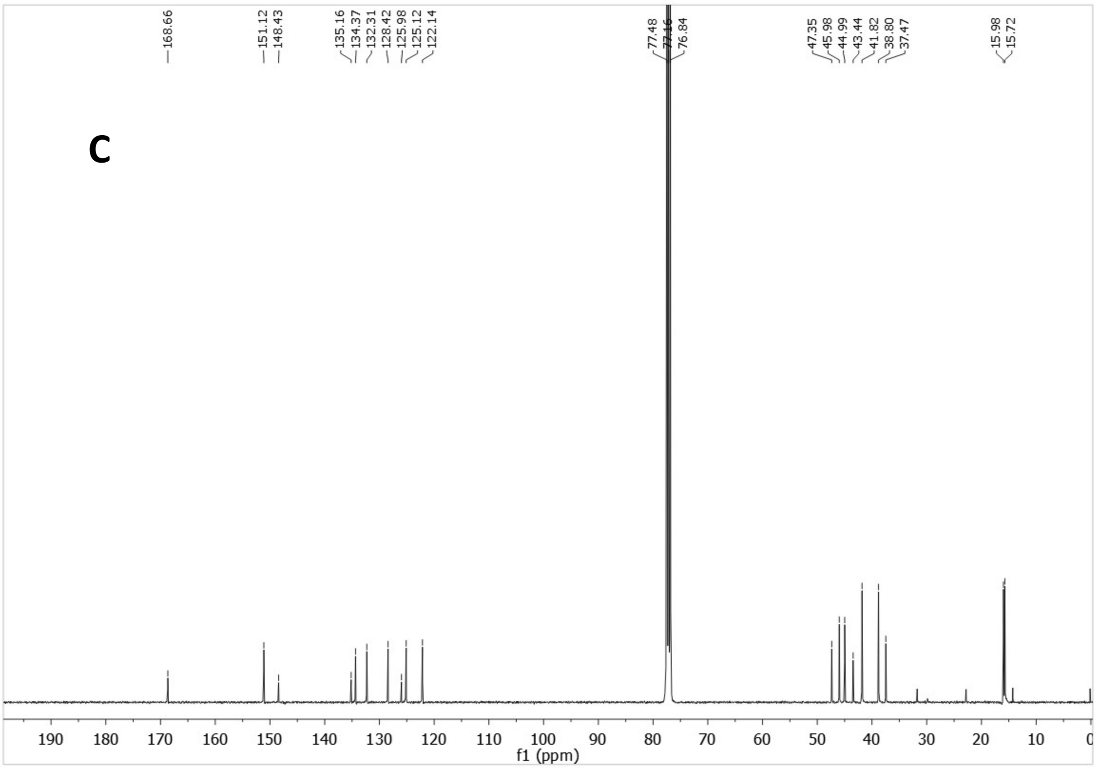


HSQC

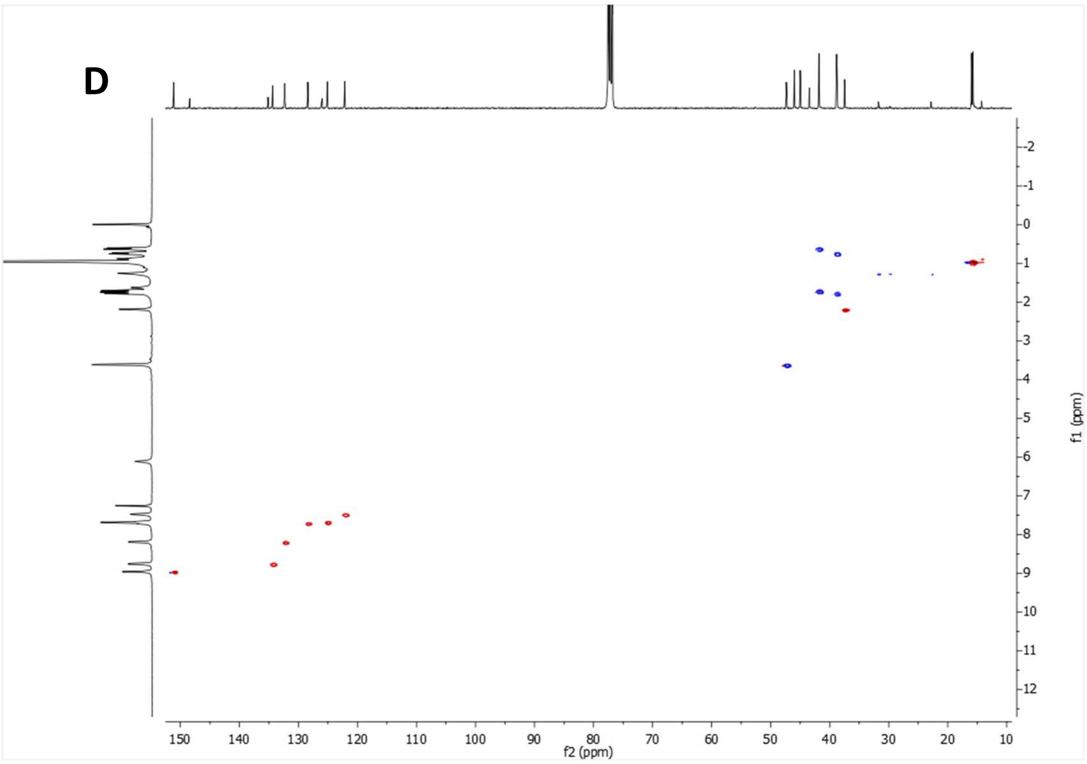


**Supplementary Figure 17:** *N*-((3,4,8,9-tetramethyltetracyclo[4.4.0.0$^{3,9}$.0$^{4,8}$]decan-1-yl)methyl)quinoline-5-carboxamide, ALT-P10 (A-B) $^{1}$H NMR spectrum. (C) $^{13}$C NMR spectrum. (D) HSQC spectrum.

$^1H$ NMR (400 MHz, $CDCl_3$)

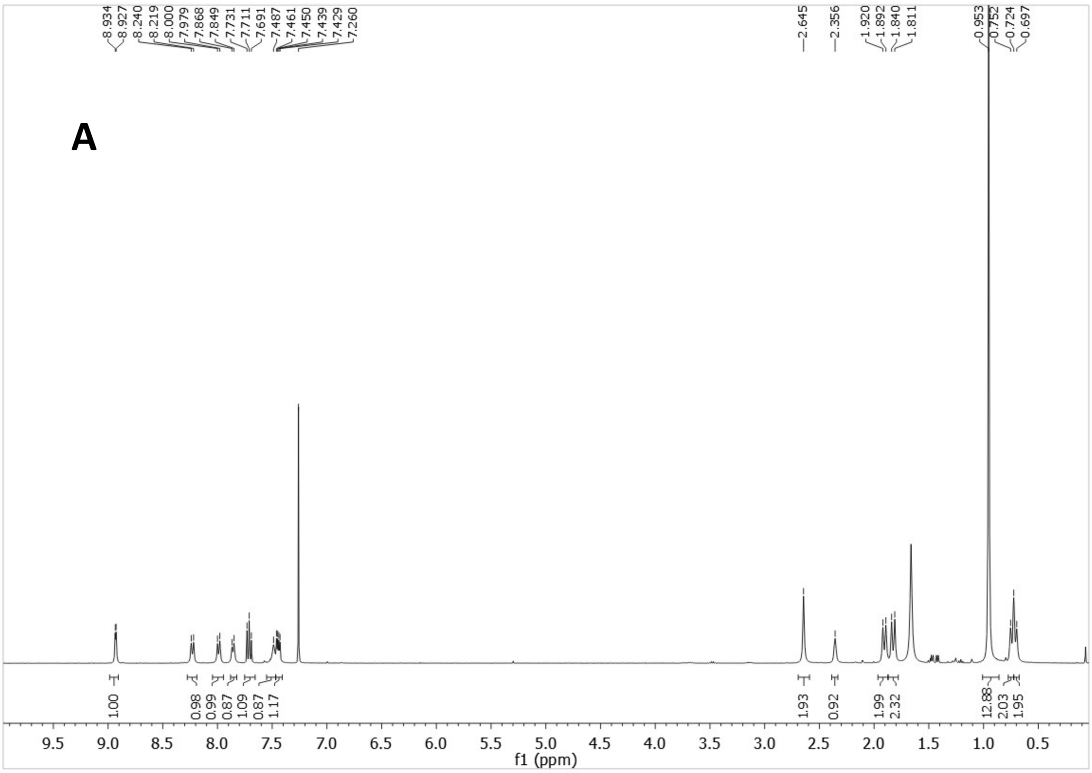

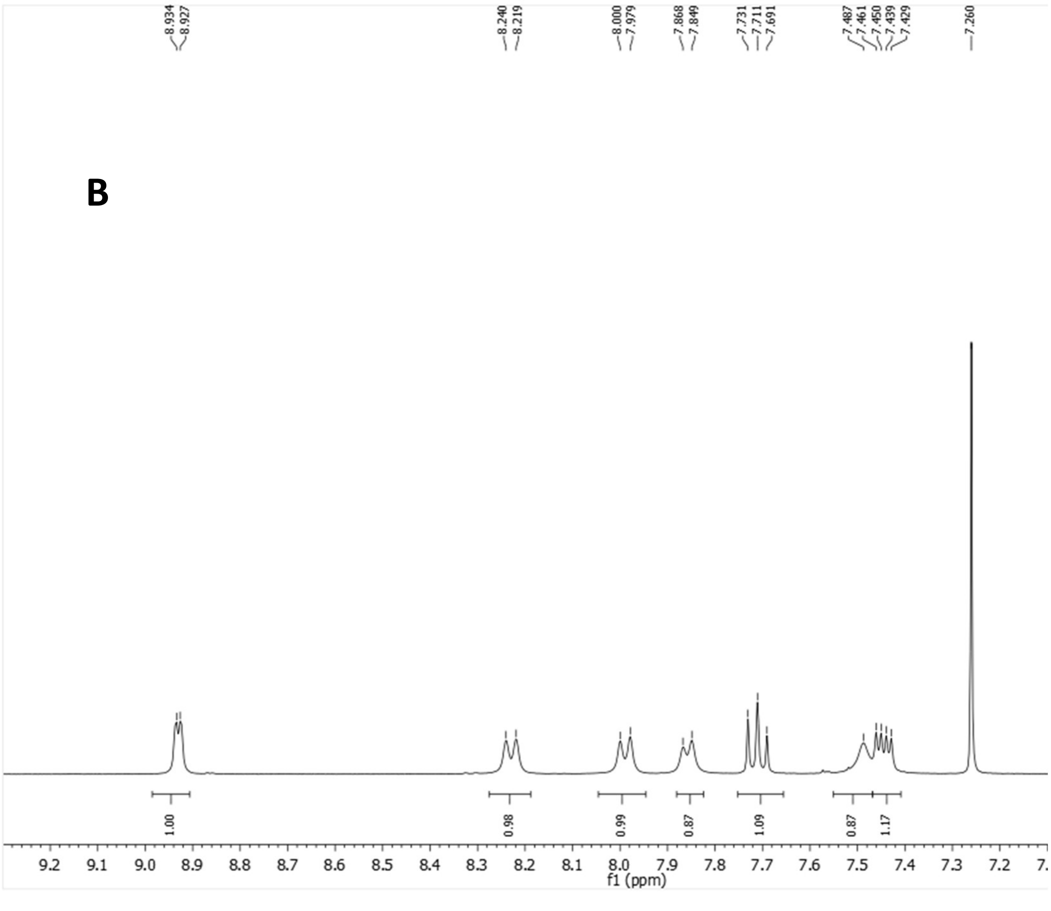

$^{13}$C-NMR (100.6 MHz, $CDCl_3$)

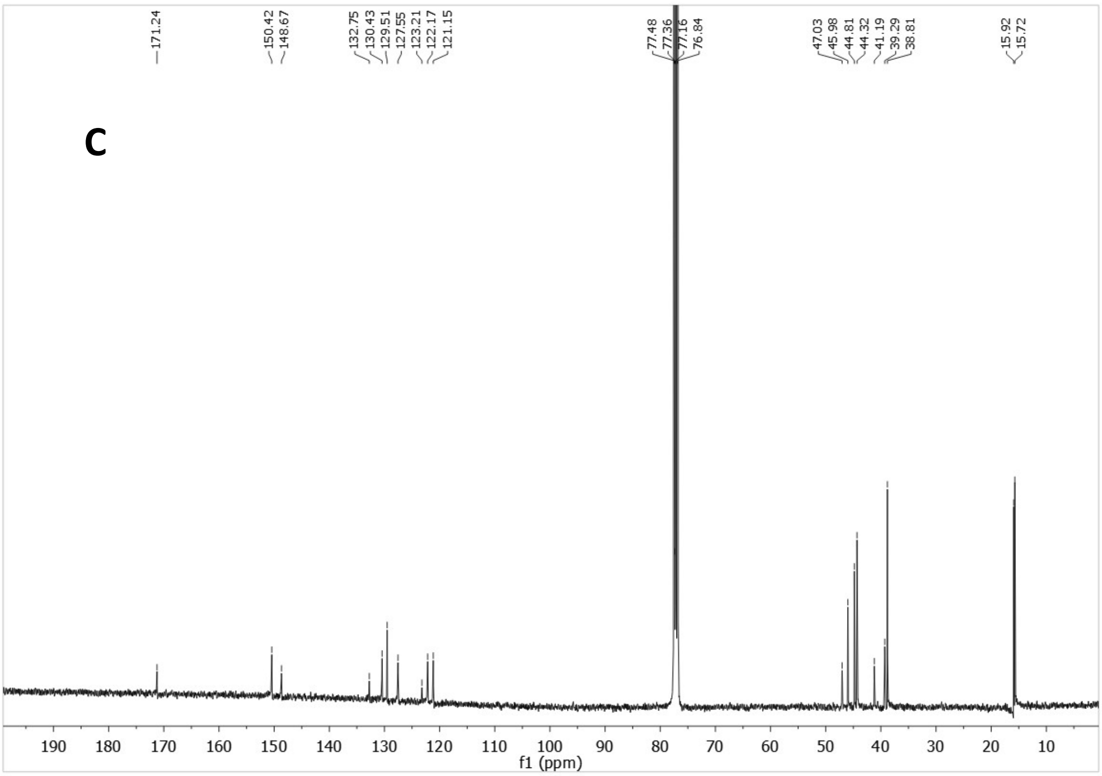


HSQC

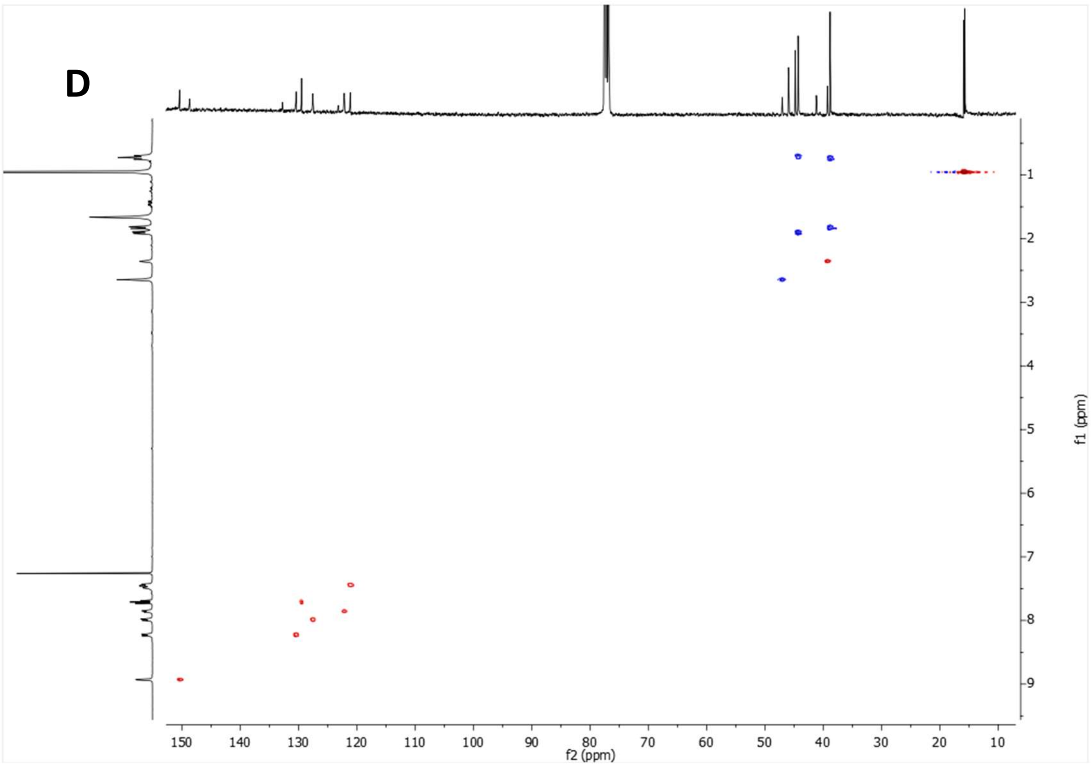


**Supplementary Figure 18:** *N*-(quinolin-5-yl)-2-(3,4,8,9-tetramethyltetracyclo[4.4.0.0$^{3,9}$.0$^{4,8}$]decan-1-yl)acetamide, ALT-P25 (A-B) $^{1}$H NMR spectrum. (C) $^{13}$C NMR spectrum. (D) HSQC spectrum.

$^{1}$H NMR (400 MHz, MeOD)

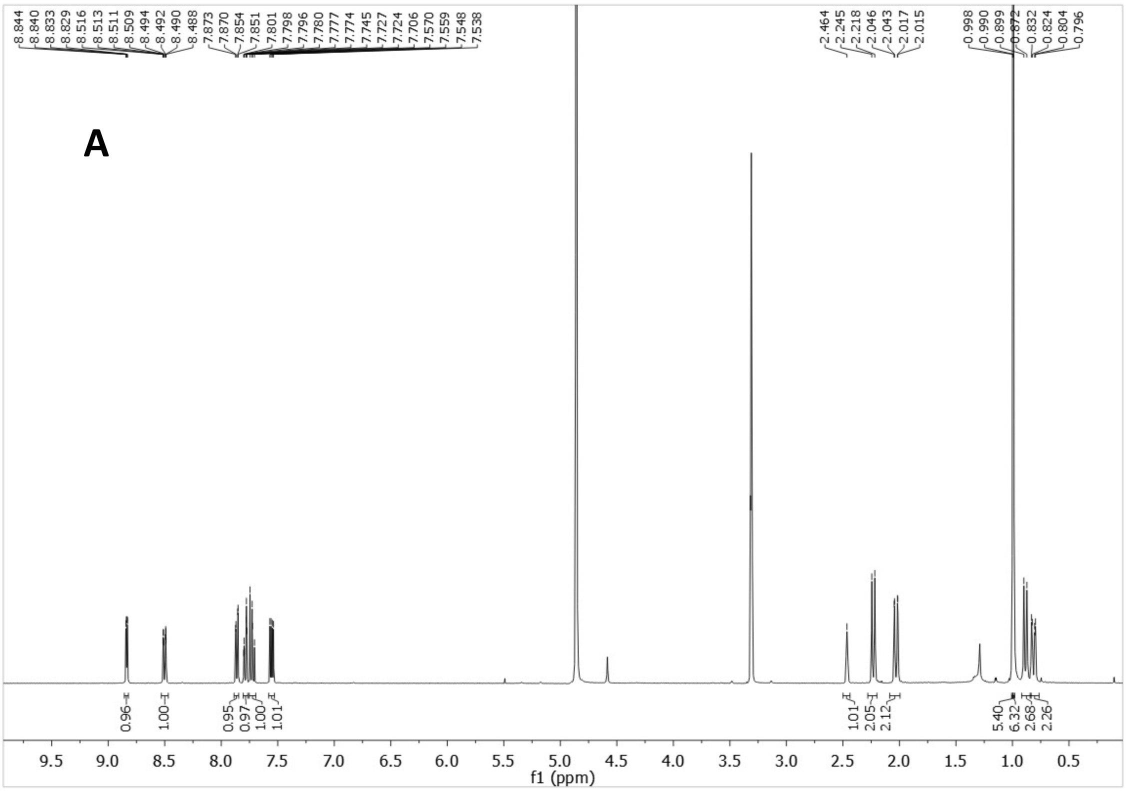

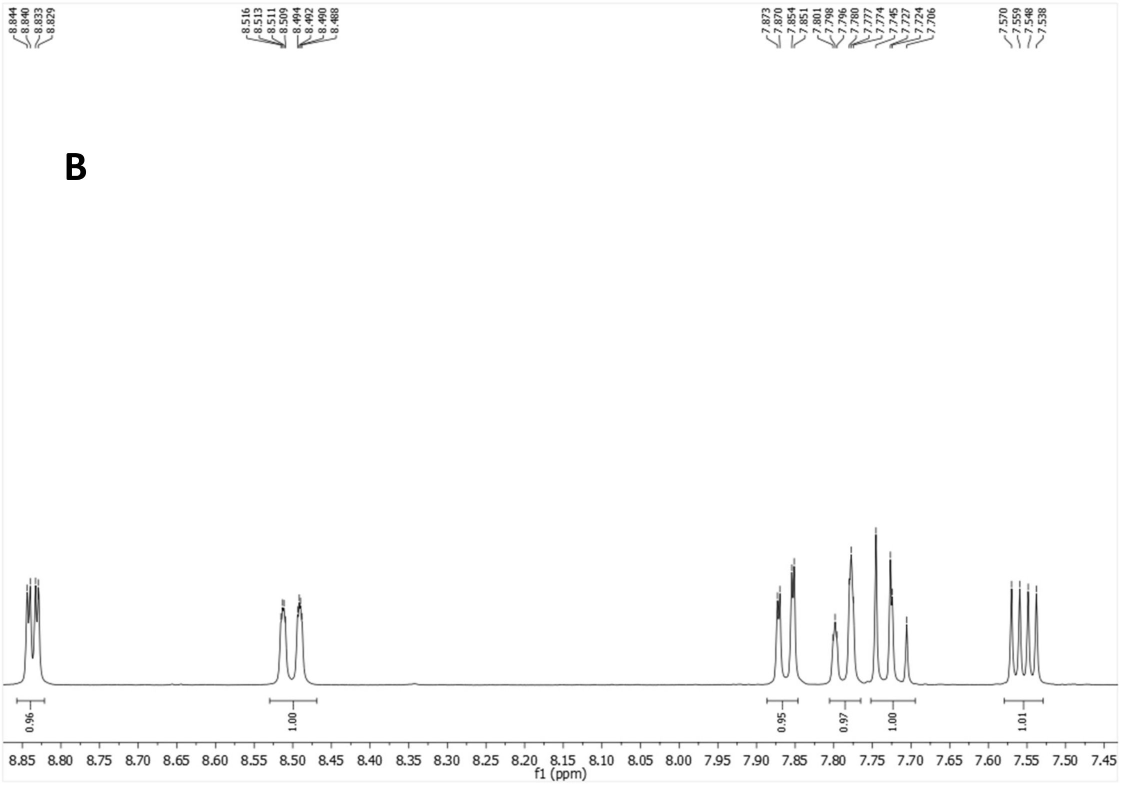

$^{13}$C-NMR (100.6 MHz, MeOD)

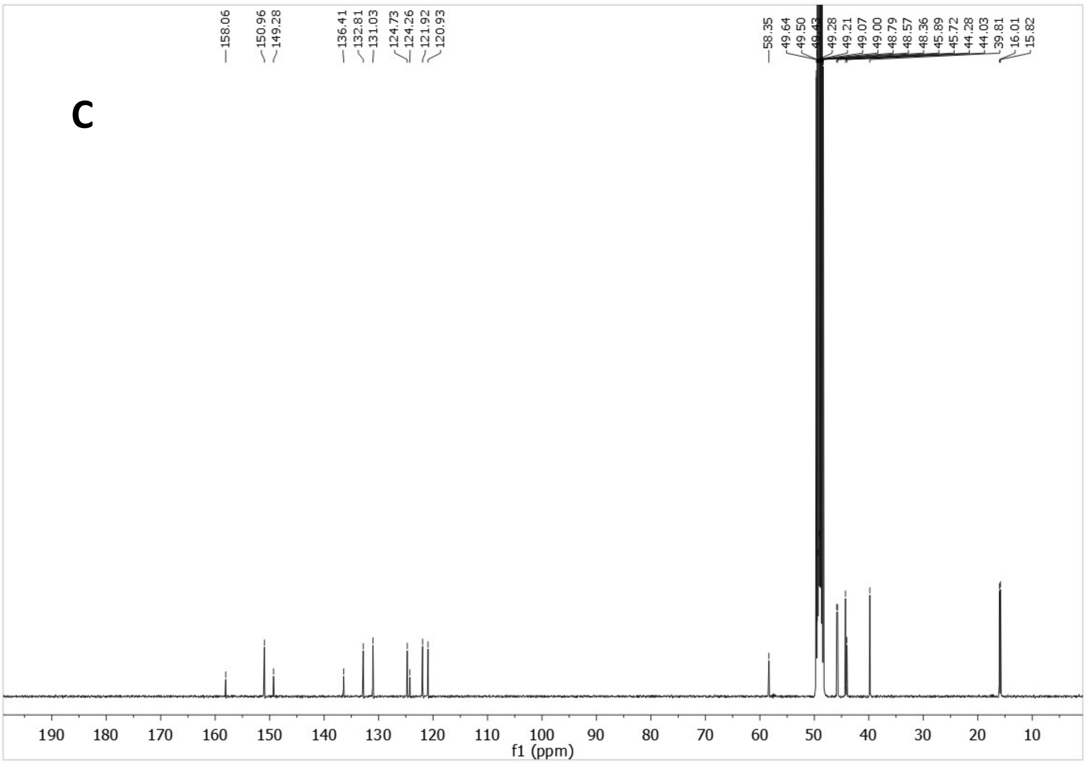


HSQC

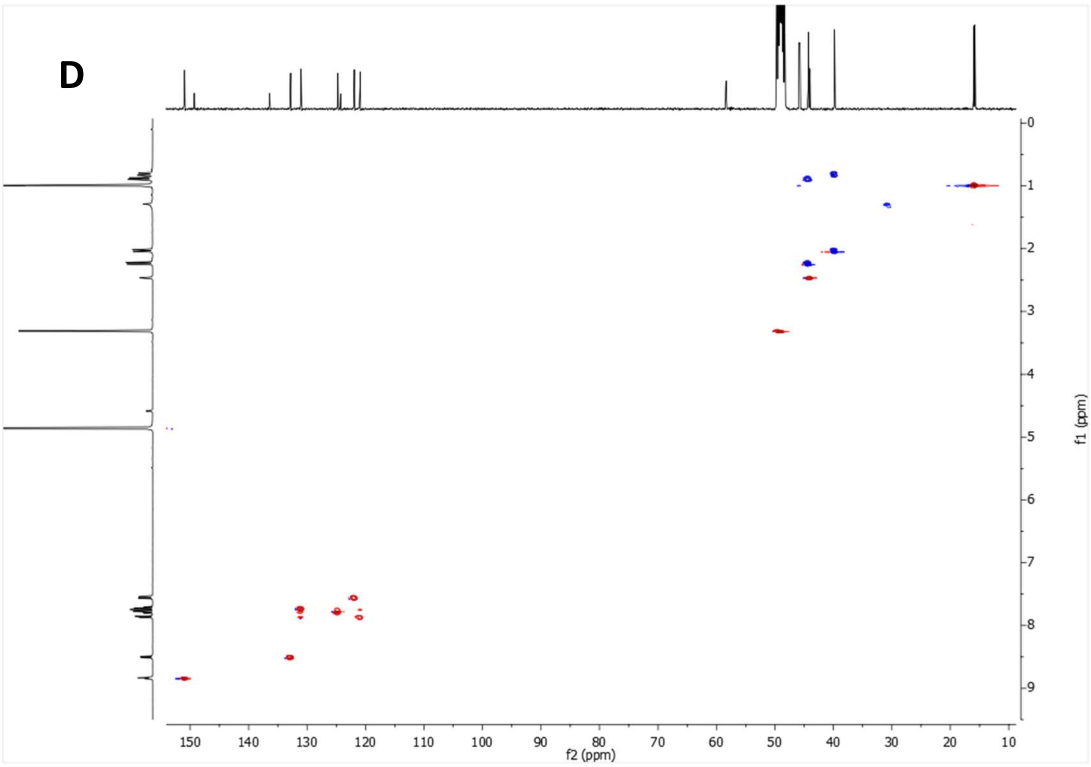


**Supplementary Figure 19:** 1-(quinolin-5-yl)-3-(3,4,8,9-tetramethyltetracyclo[4.4.0.0$^{3,9}$.0$^{4,8}$]decan-1-yl)urea, ALT-P22 (A-B) $^{1}$H NMR spectrum. (C) $^{13}$C NMR spectrum. (D) HSQC spectrum.

$^1$H NMR (400 MHz, MeOD)

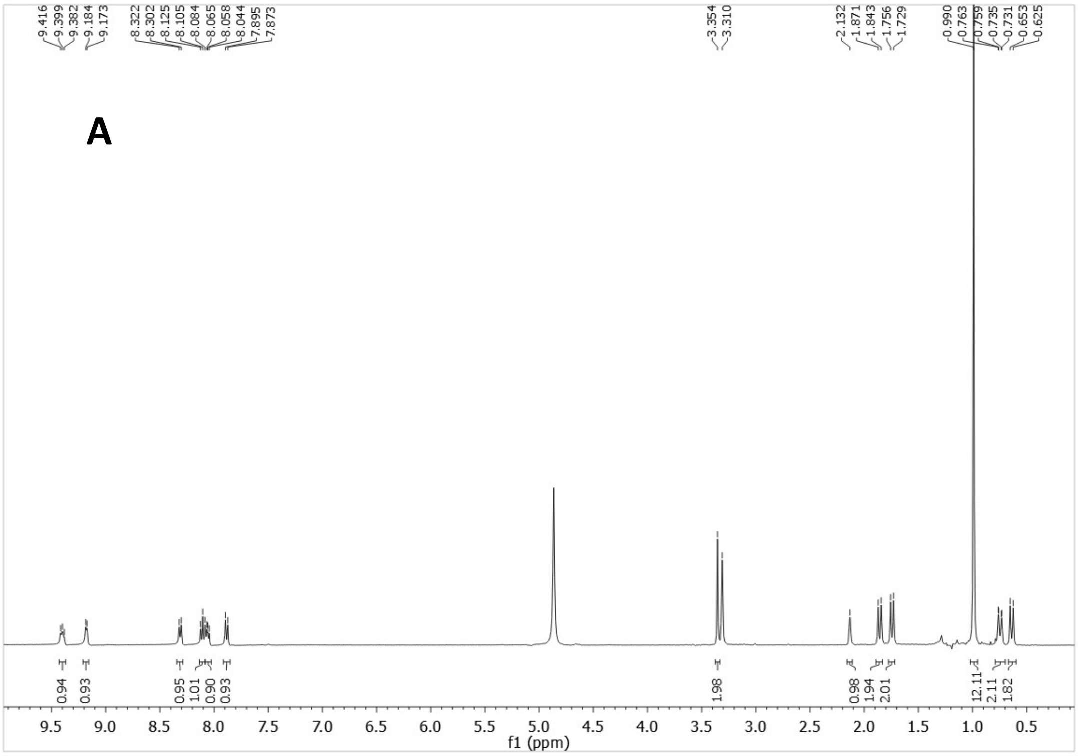


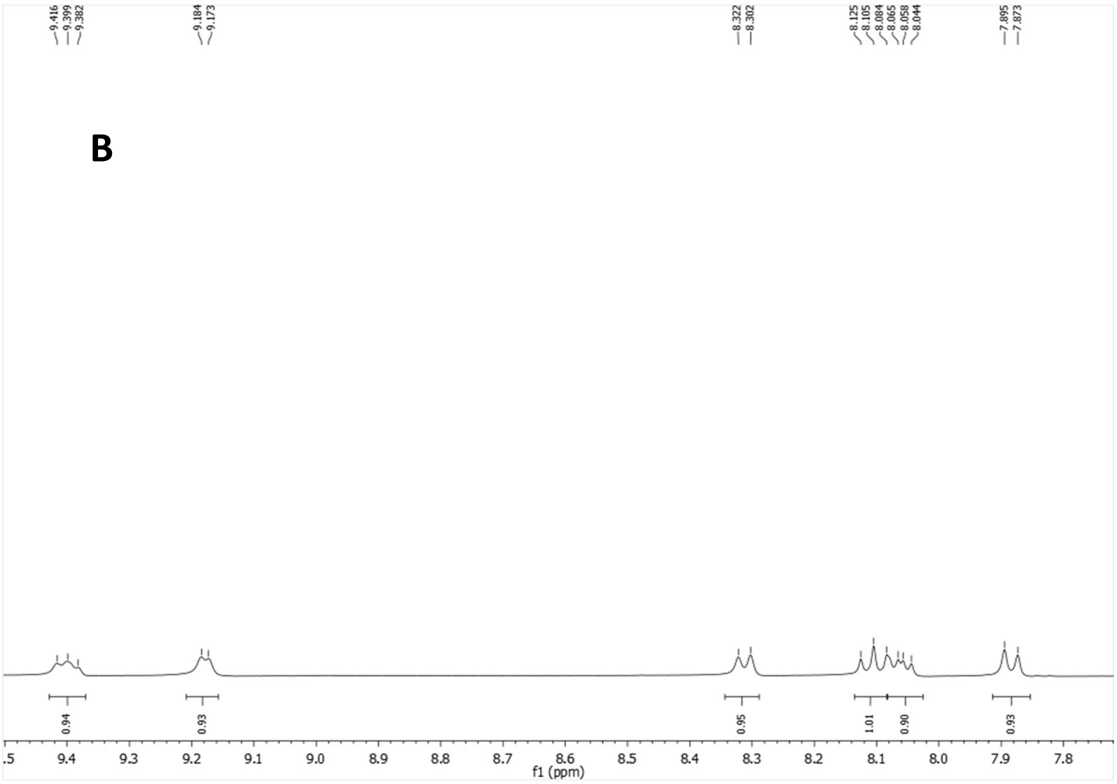

$^{13}$C-NMR (100.6 MHz, MeOD)

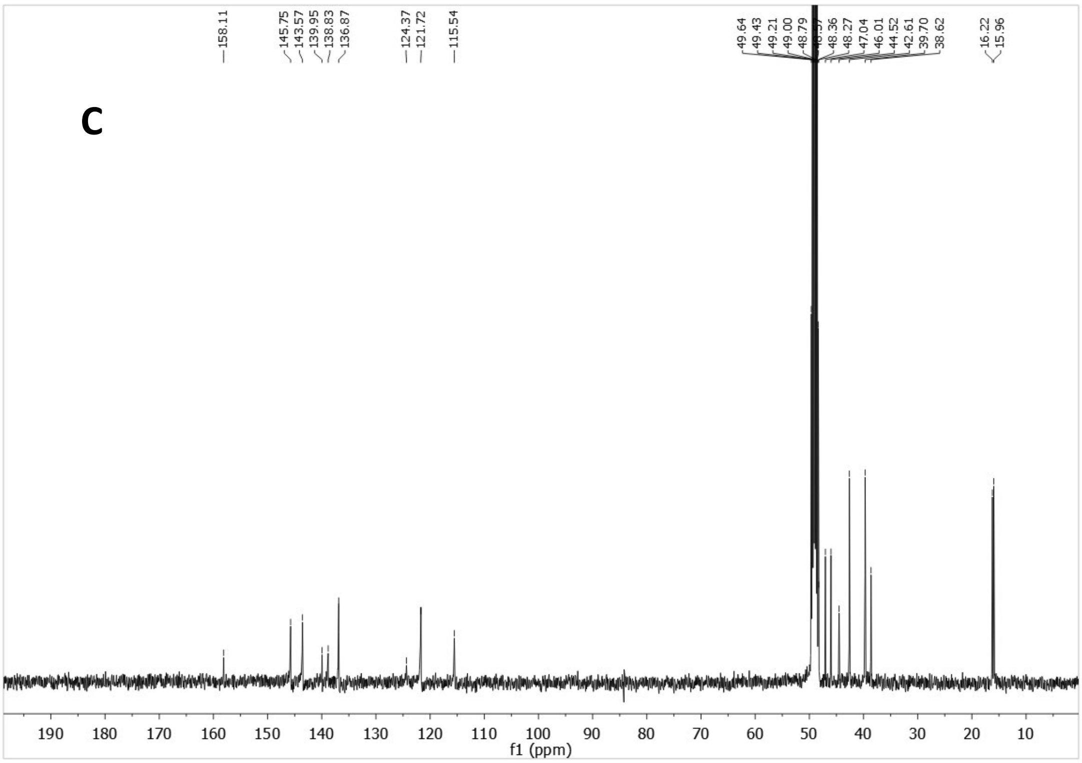


HSQC

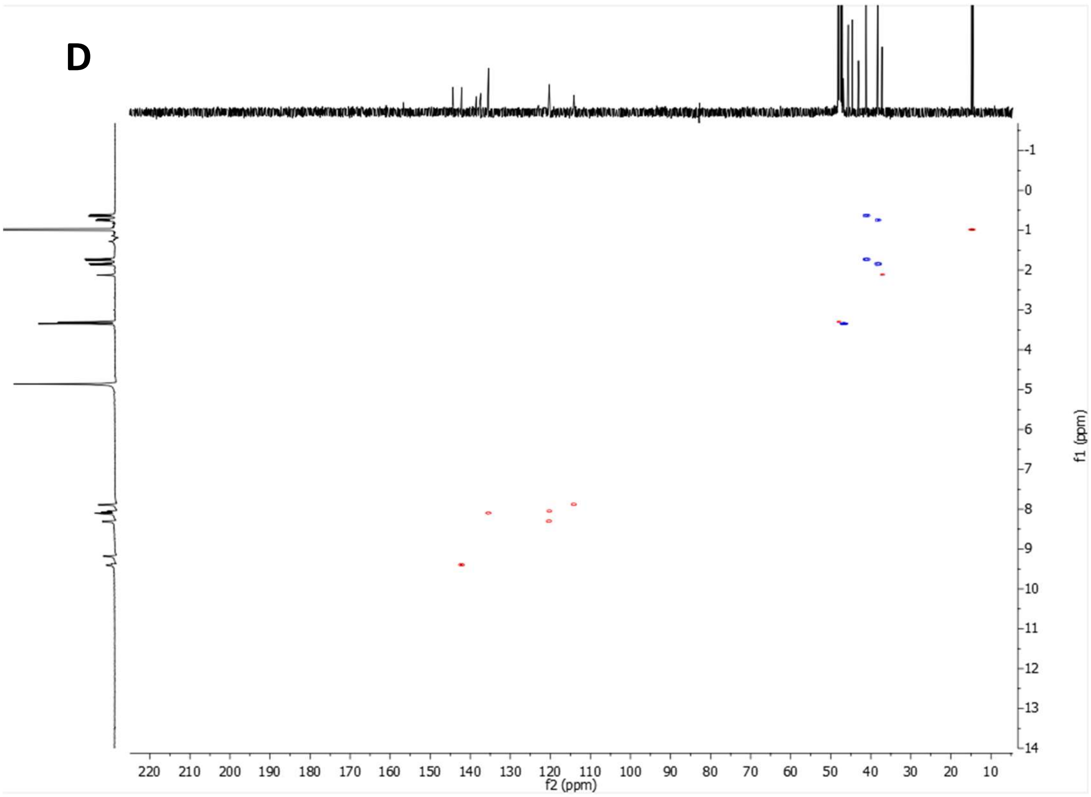


**Supplementary Figure 20:** 1-(quinolin-5-yl)-3-((3,4,8,9-tetramethyltetracyclo[4.4.0.0$^{3,9}$.0$^{4,8}$]decan-1-yl)methyl)urea hydrochloride, ALT-P16 (A-B) $^{1}$H NMR spectrum. (C) $^{13}$C NMR spectrum. (D) HSQC spectrum.

$^1H$ NMR (400 MHz, $CDCl_3$)

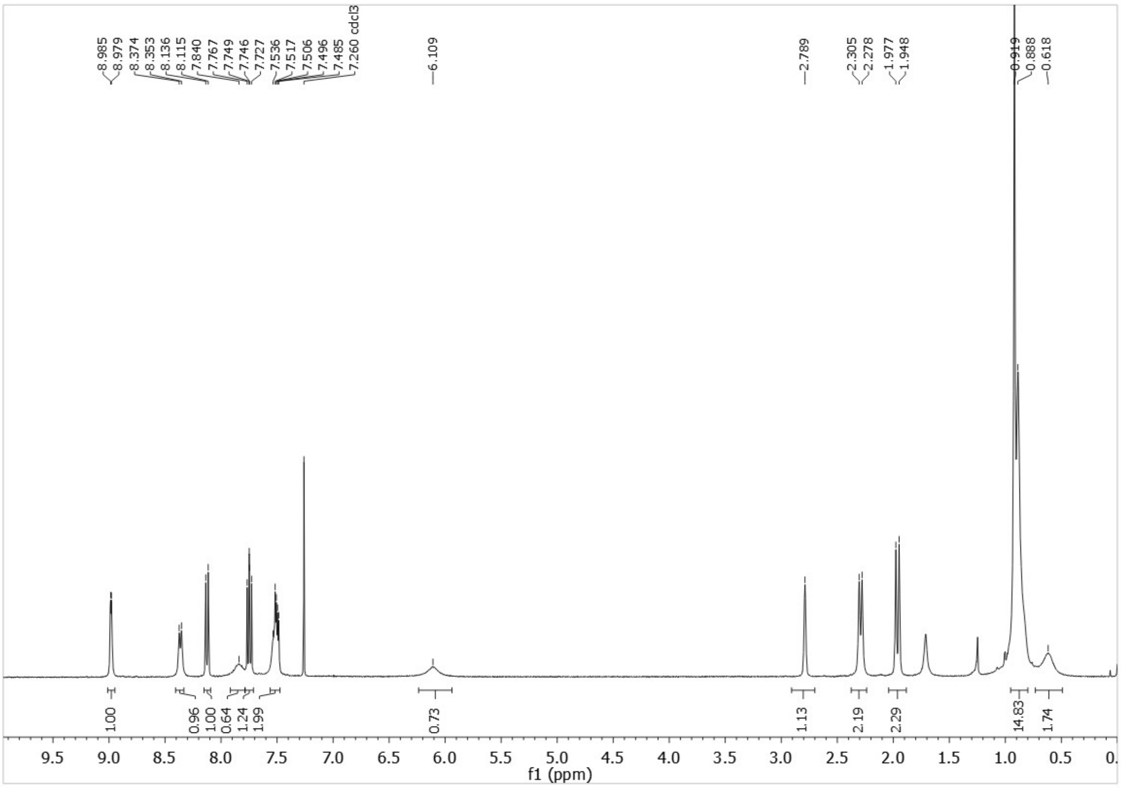


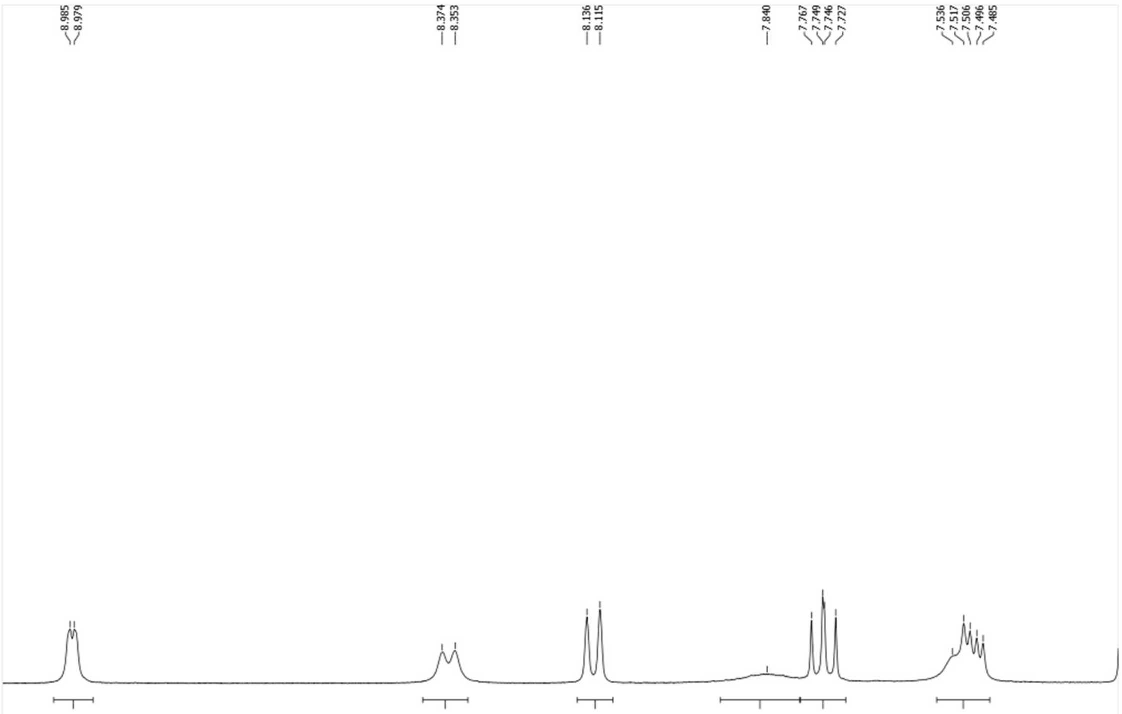

$^{13}C$-NMR (100.6 MHz, $CDCl_3$)

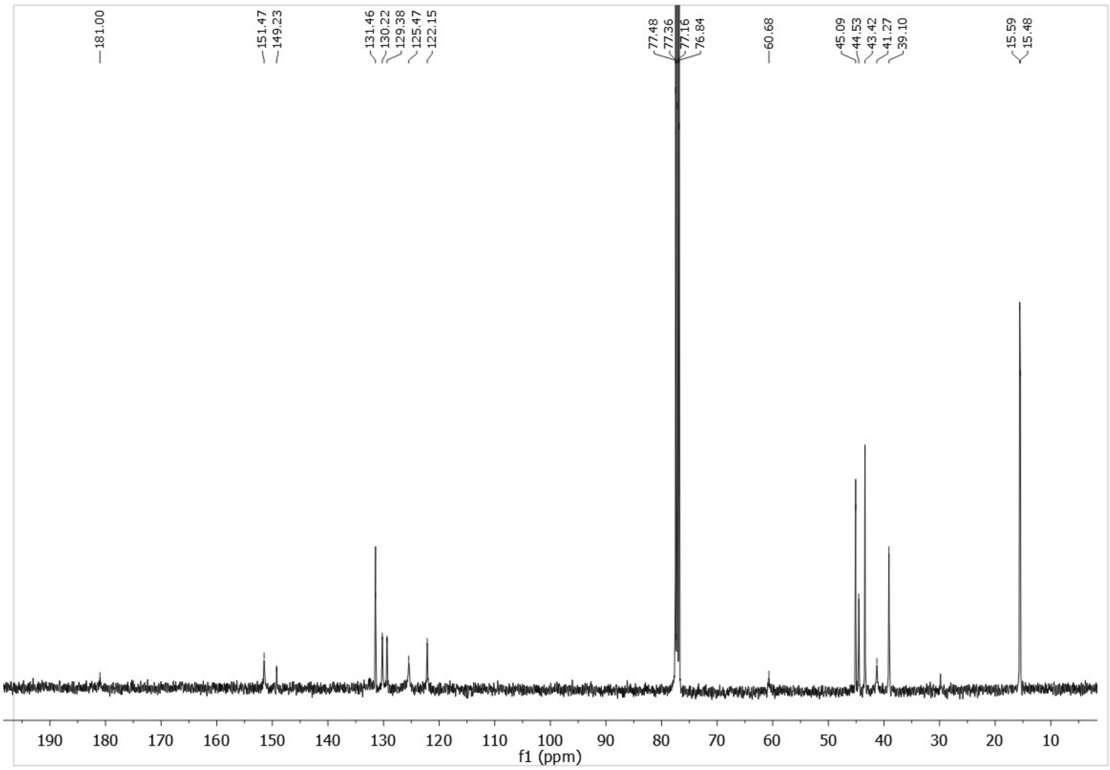


HSQC

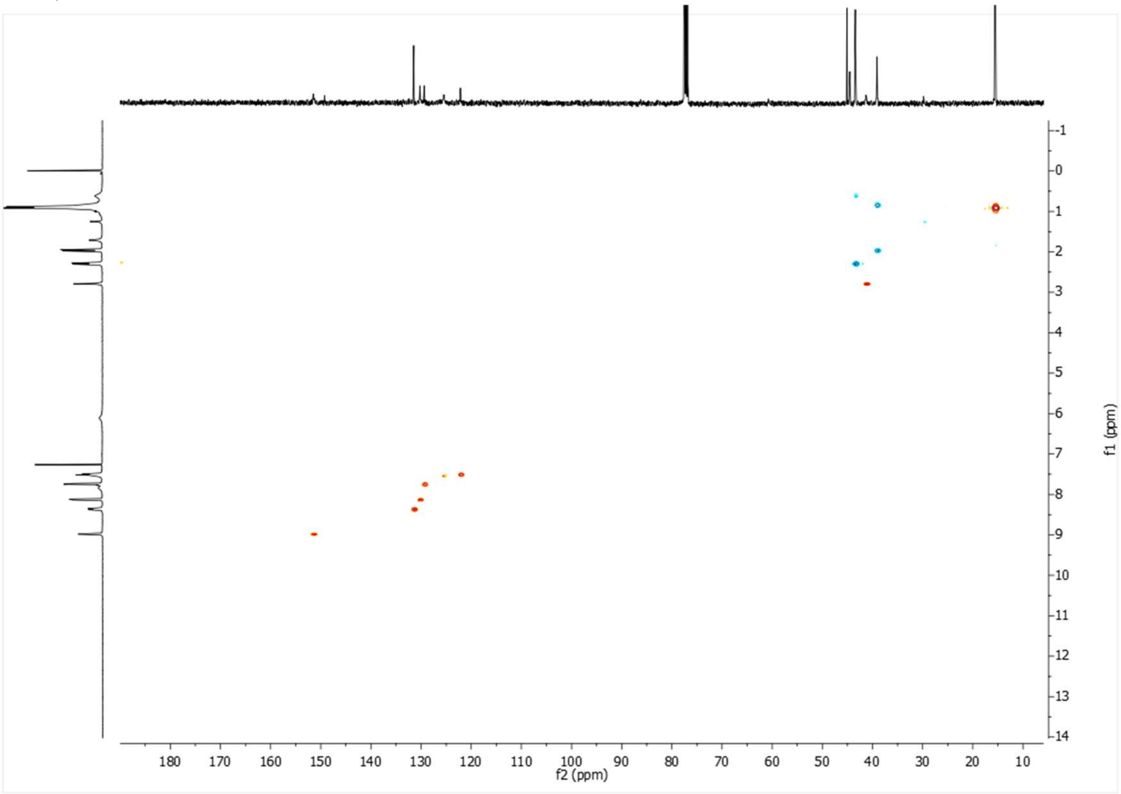


**Supplementary Figure 21:** 1-(quinolin-5-yl)-3-(3,4,8,9-tetramethyltetracyclo[4.4.0.0$^{3,9}$.0$^{4,8}$]decan-1-yl)thiourea, ALT-P13 (A-B) $^{1}H$ NMR spectrum. (C) $^{13}C$ NMR spectrum. (D) HSQC spectrum.

$^1H$ NMR (400 MHz, $CDCl_3$)

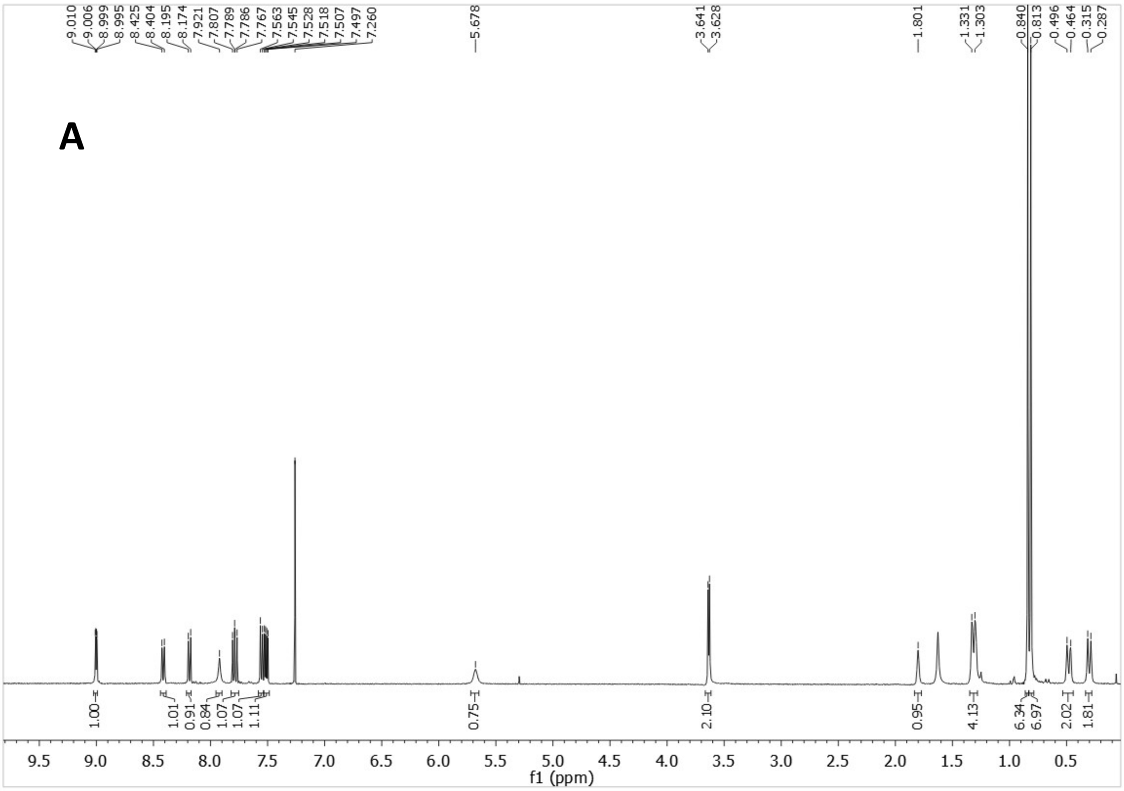


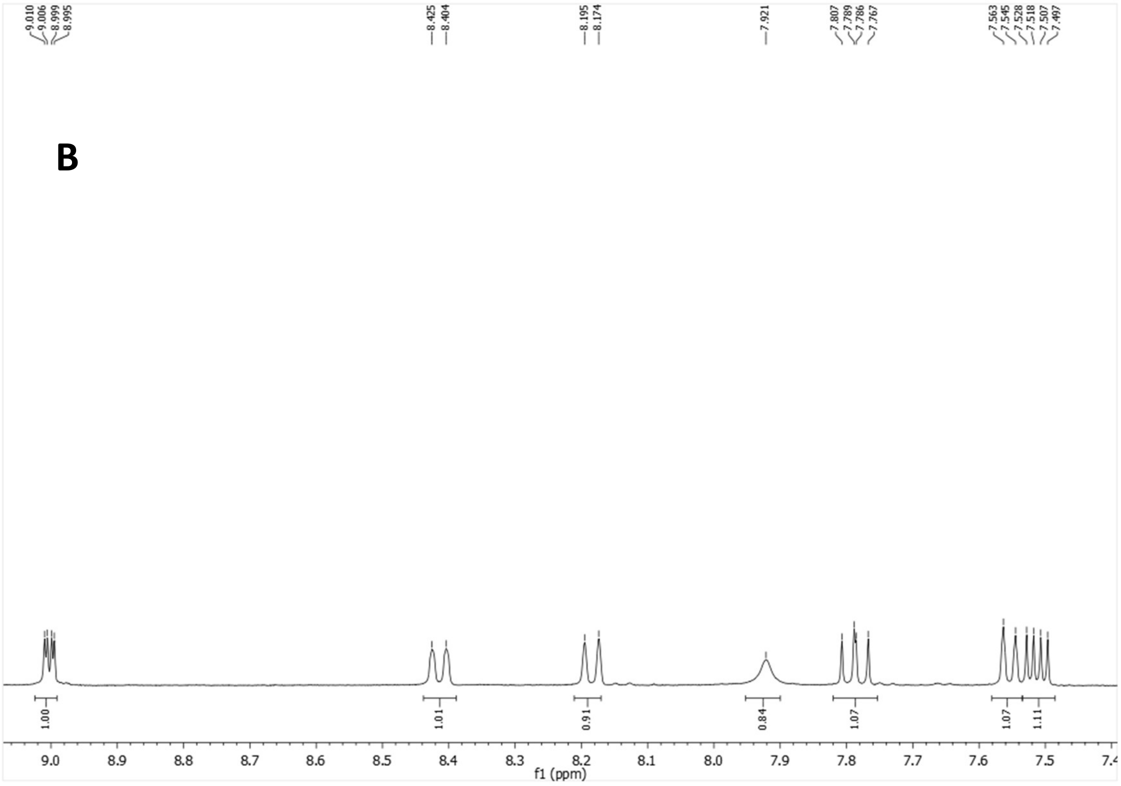

[13]C-NMR (100.6 MHz, $CDCl_3$)

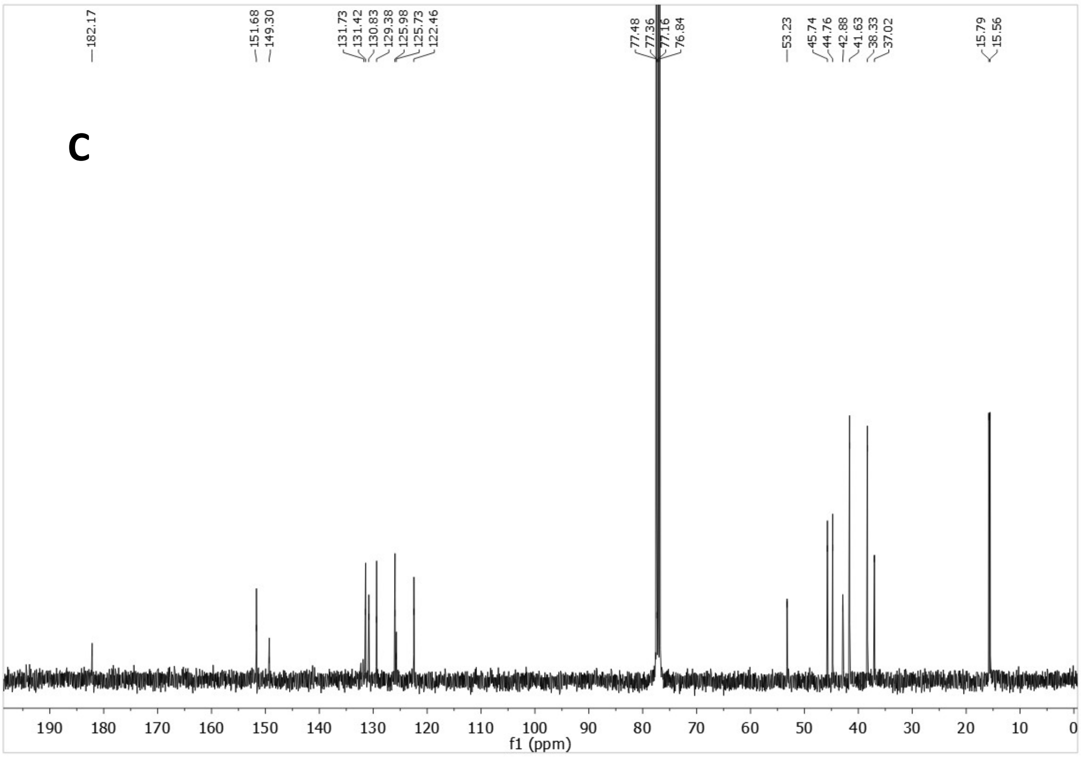


HSQC

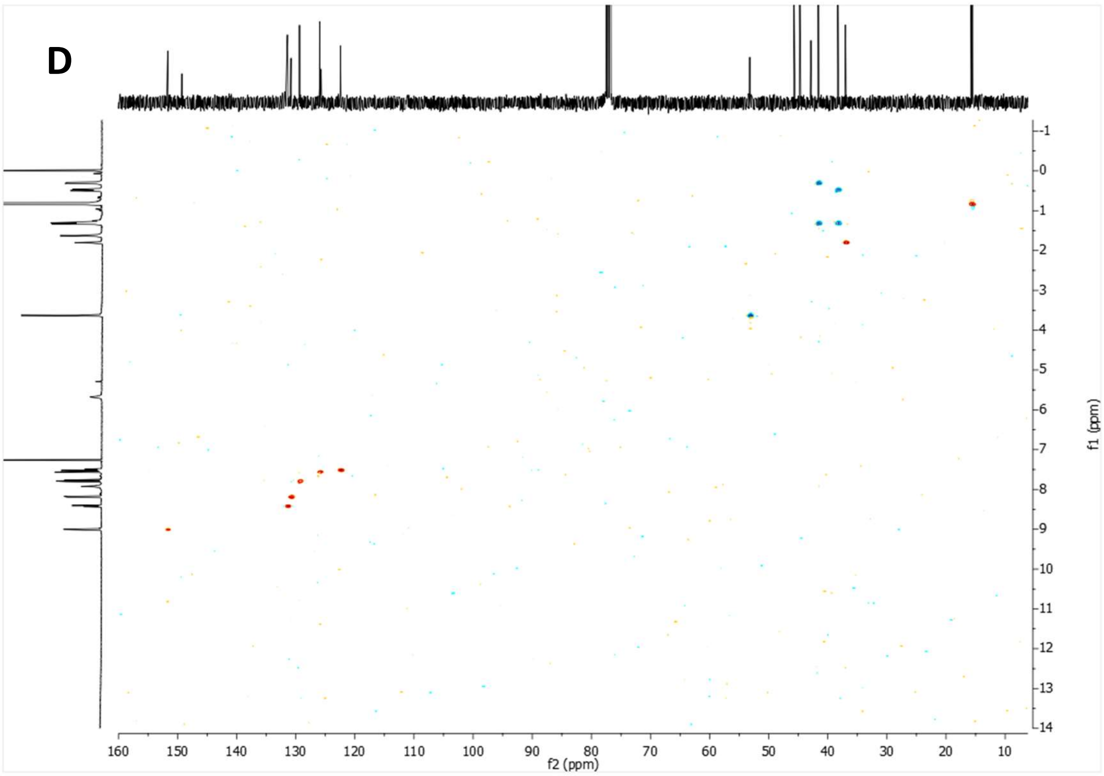


**Supplementary Figure 22:** 1-(quinolin-5-yl)-3-((3,4,8,9-tetramethyltetracyclo[4.4.0.0$^{3,9}$.0$^{4,8}$]decan-1-yl)methyl)thiourea, ALT-P14 (A-B) $^{1}$H NMR spectrum. (C) $^{13}$C NMR spectrum. (D) HSQC spectrum.

[1]H NMR (400 MHz, DMSO)

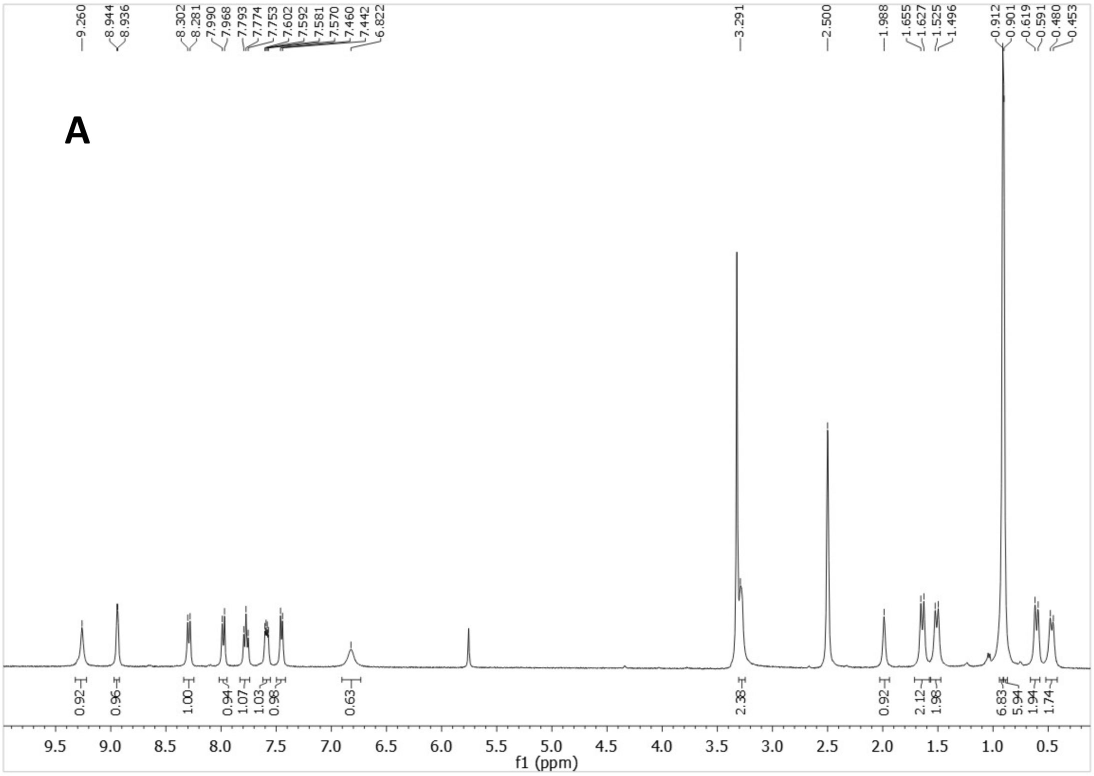

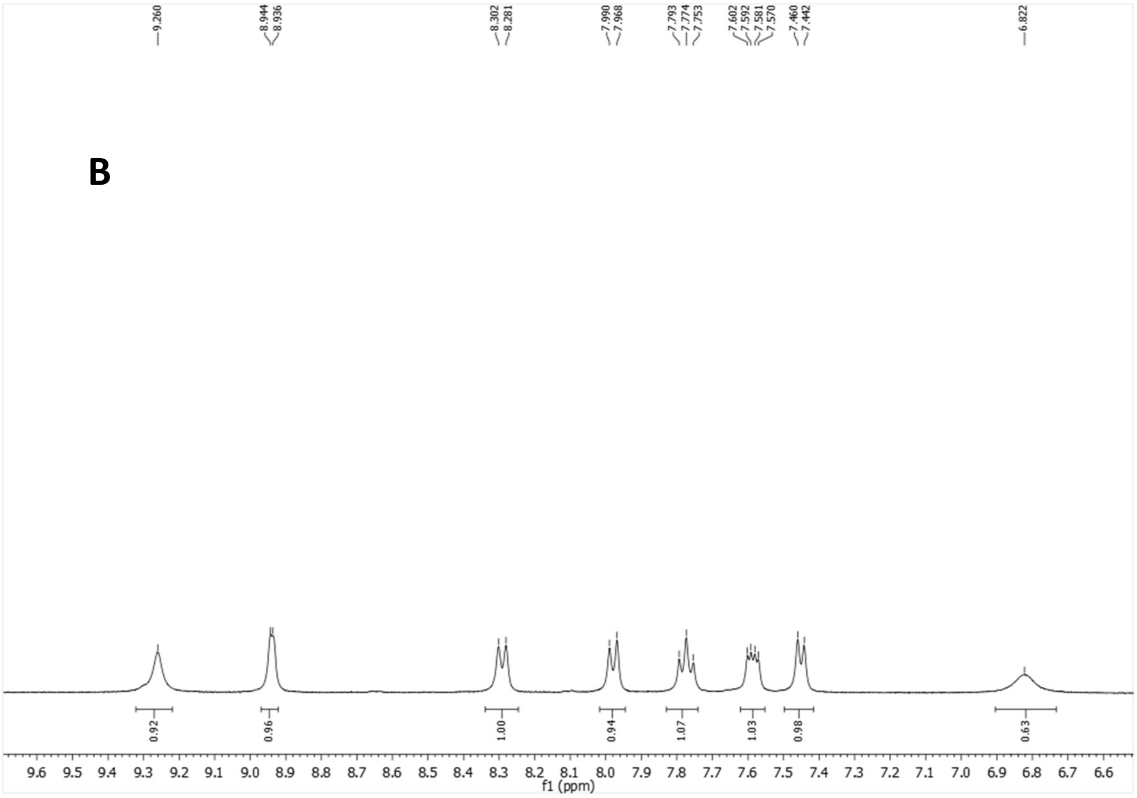

$^{13}C$-NMR (100.6 MHz, DMSO)

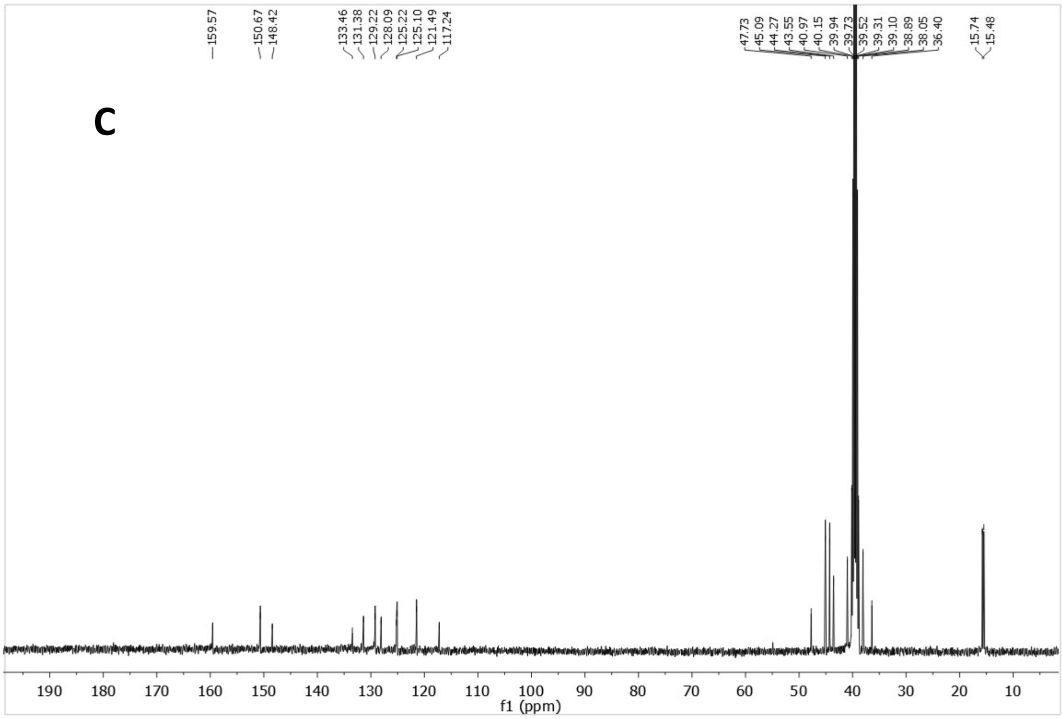


HSQC

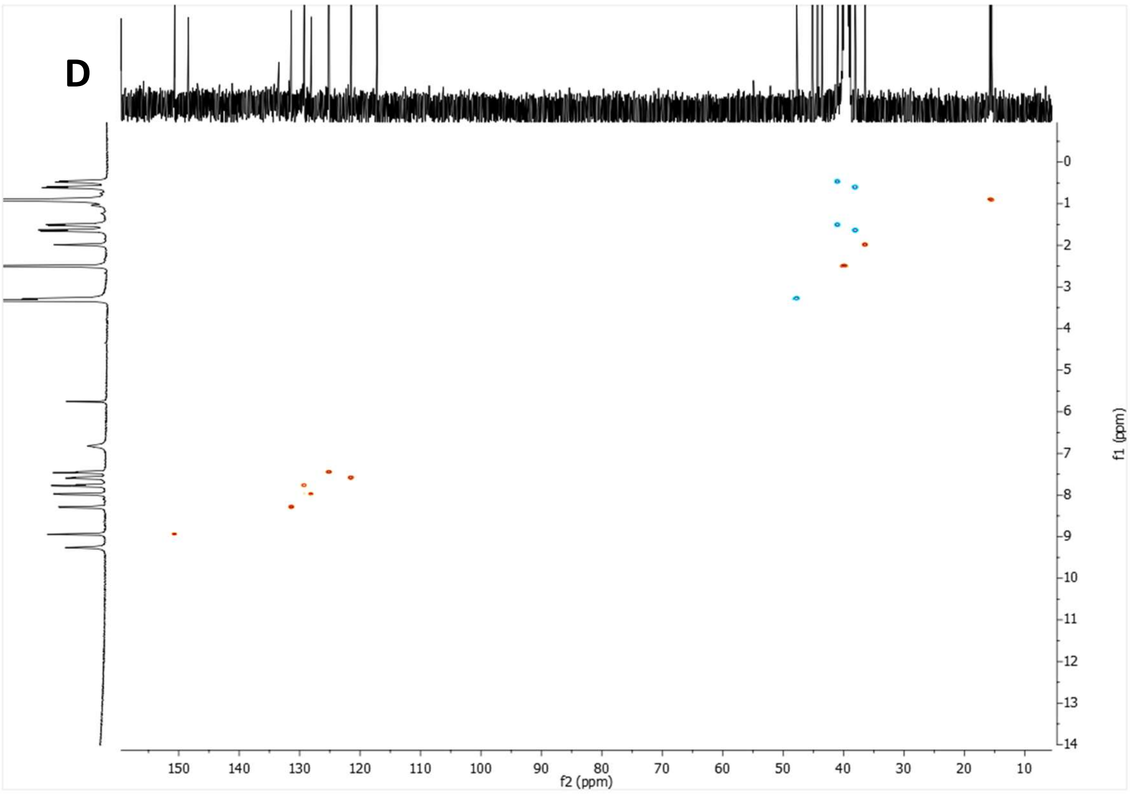


**Supplementary Figure 23:** 2-cyano-1-(quinolin-5-yl)-3-((3,4,8,9-tetramethyltetracyclo[4.4.0.0$^{3,9}$.0$^{4,8}$]decan-1-yl)methyl)guanidine, ALT-P7 (A-B) $^{1}H$ NMR spectrum. (C) $^{13}C$ NMR spectrum. (D) HSQC spectrum.

[1]H NMR (400 MHz, $CDCl_3$)

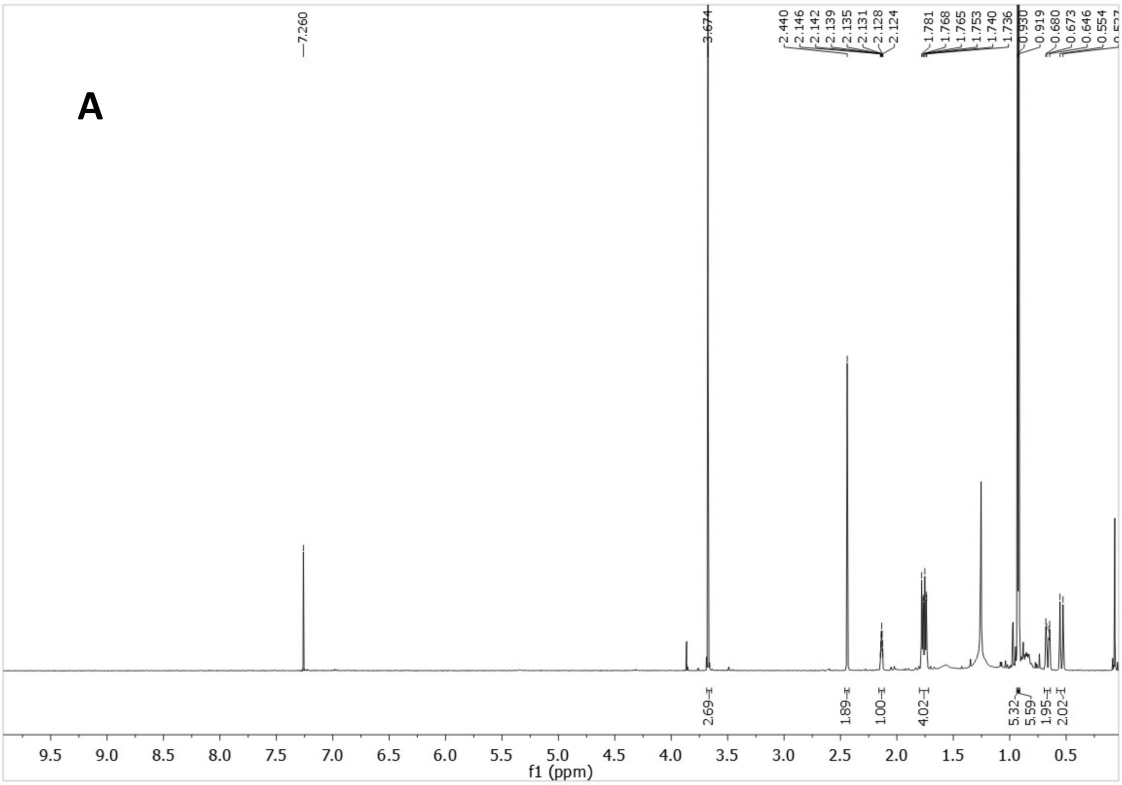


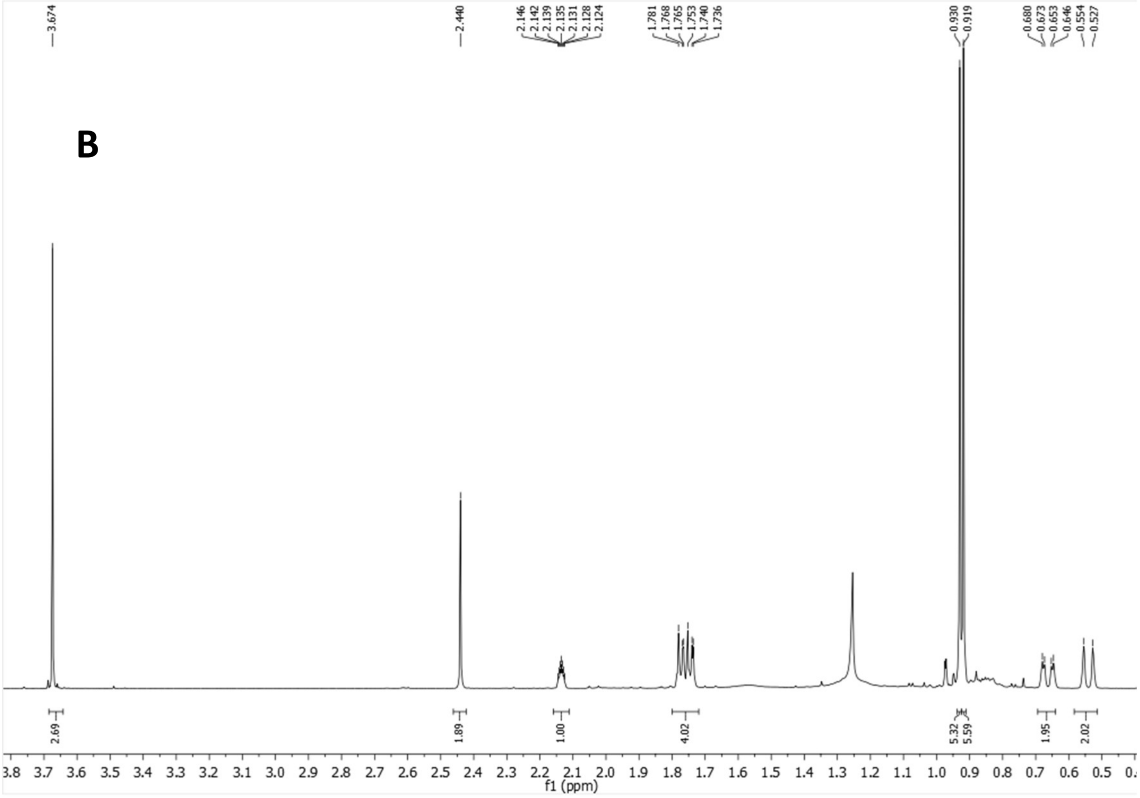

$^{13}$C-NMR (100.6 MHz, $CDCl_3$)

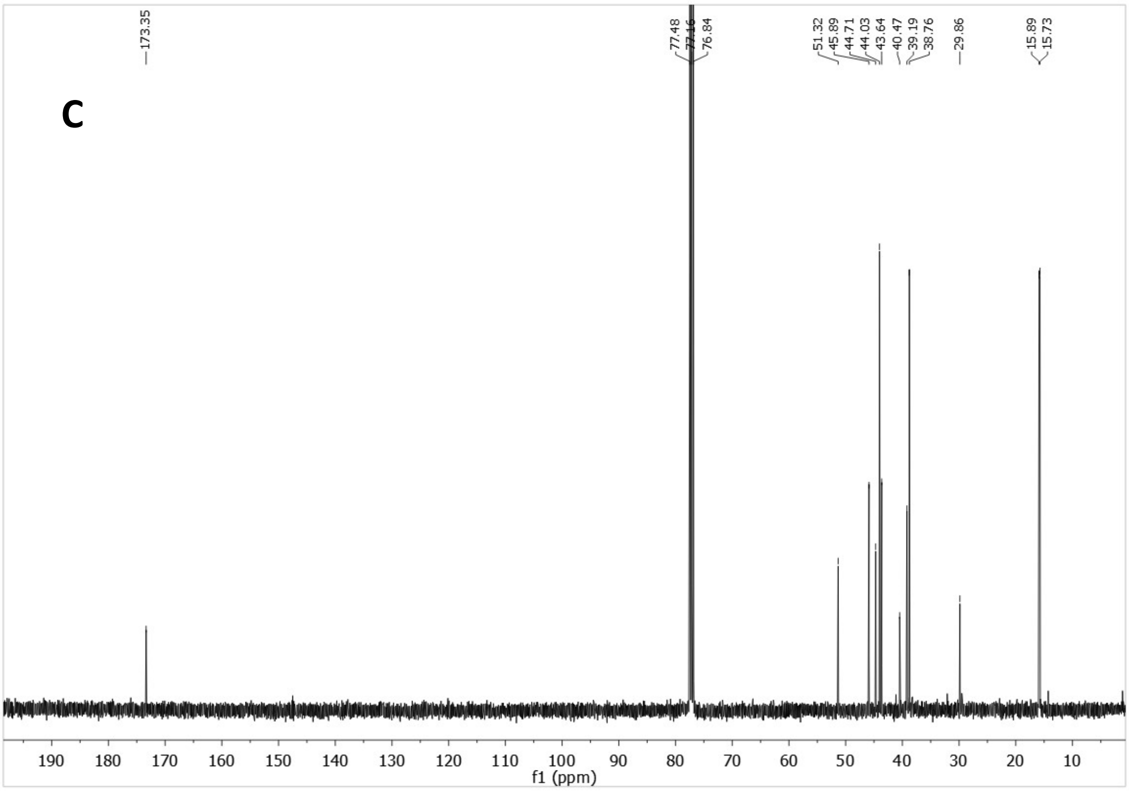


HSQC

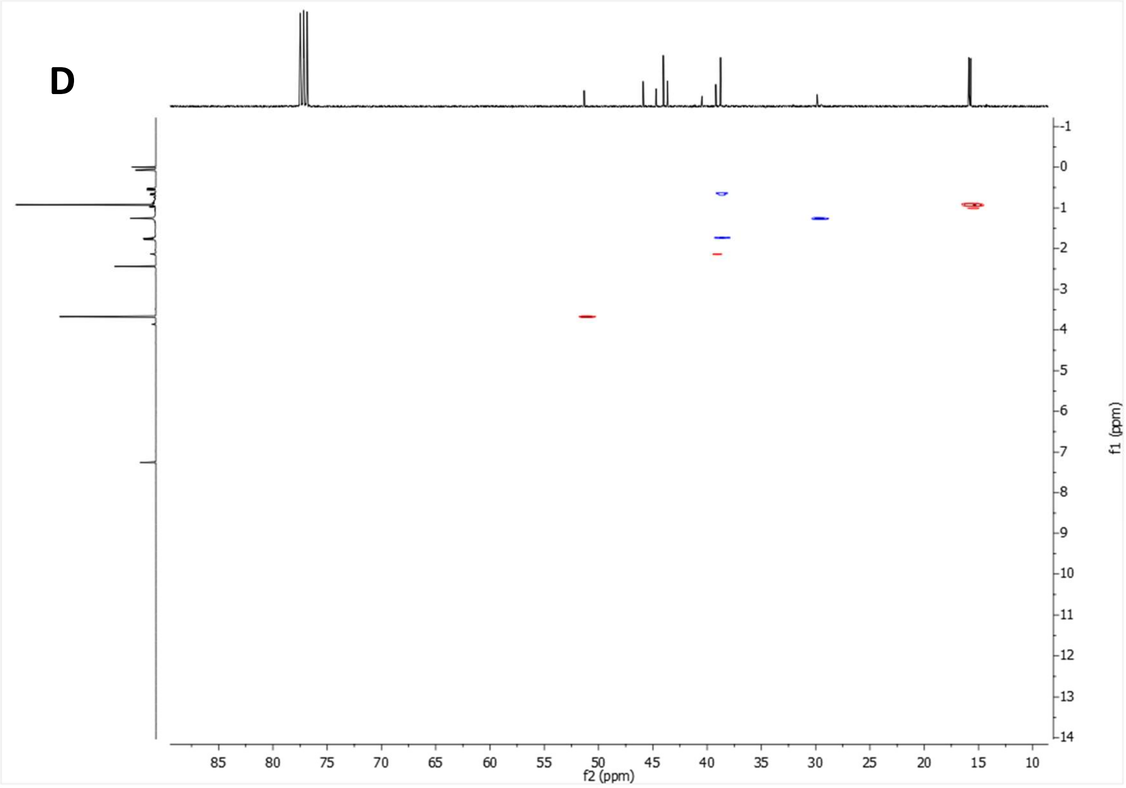


**Supplementary Figure 24:** methyl 2-(3,4,8,9-tetramethyltetracyclo[4.4.0.0$^{3,9}$.0$^{4,8}$]decan-1-yl)acetate, ALT-511 (A-B) $^{1}$H NMR spectrum. (C) $^{13}$C NMR spectrum. (D) HSQC spectrum.

$^{1}H$ NMR (400 MHz, $CDCl_3$)

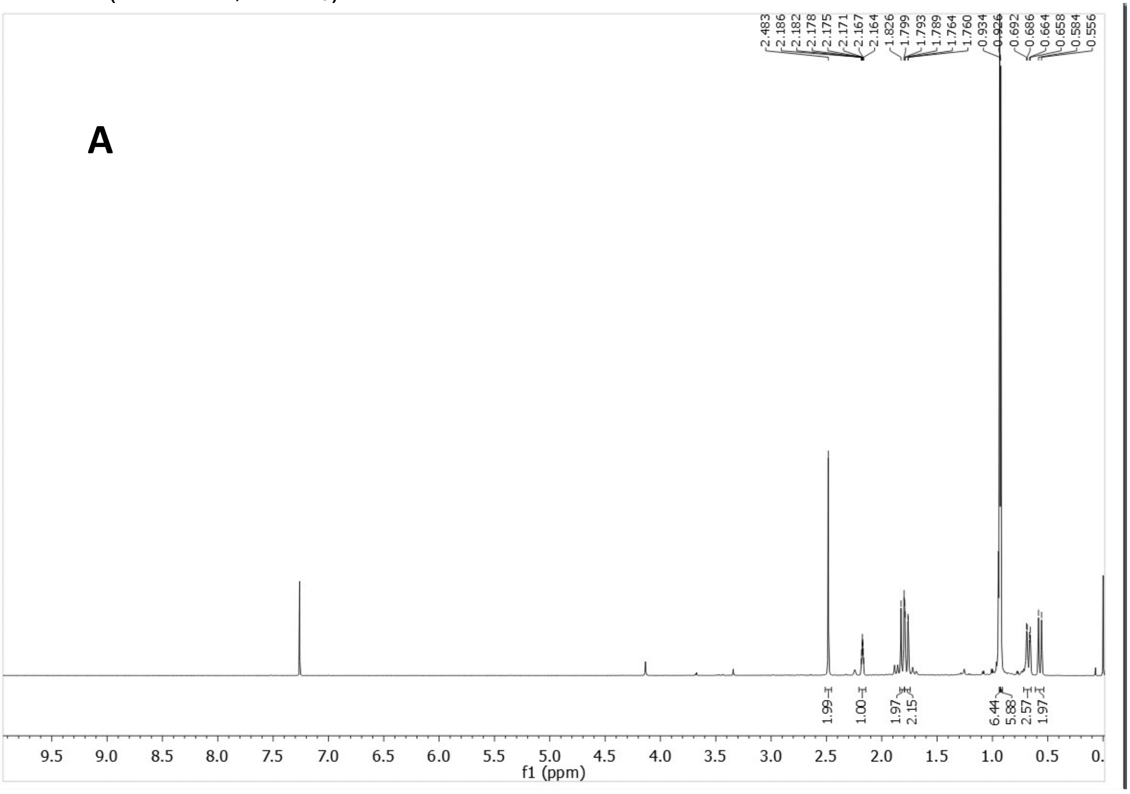


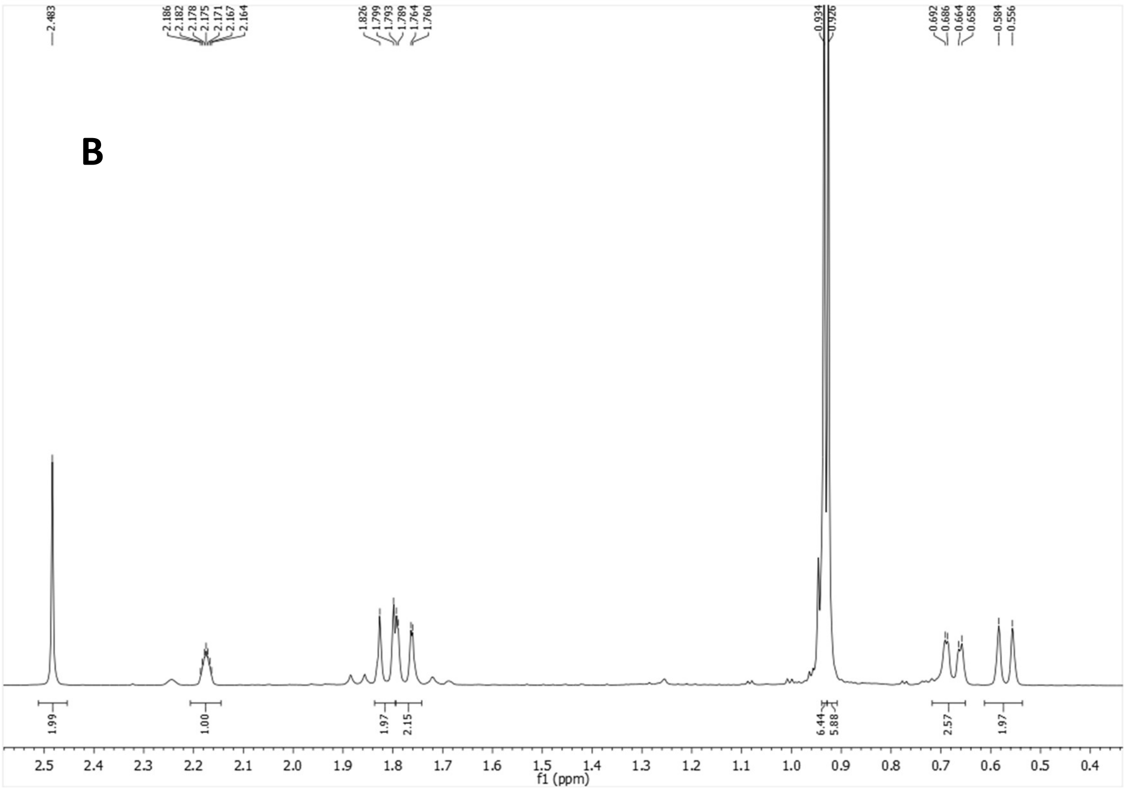

$^{13}C$-NMR (100.6 MHz, $CDCl_3$)

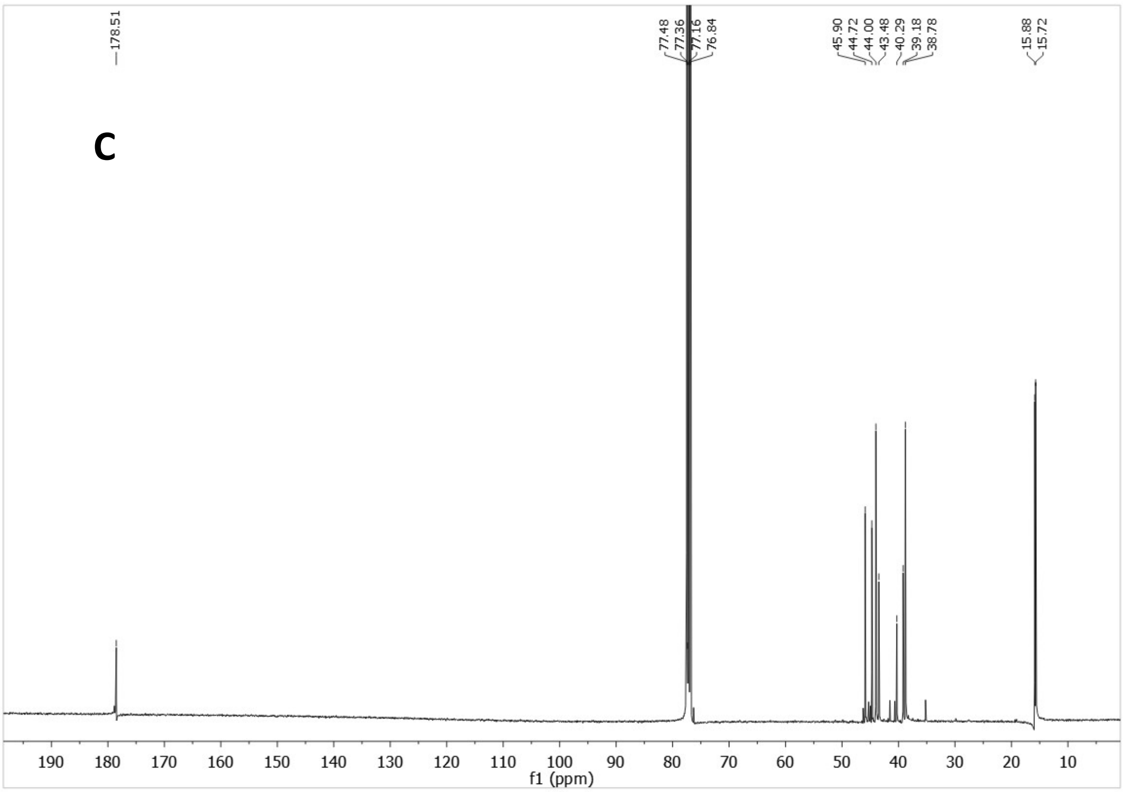


HSQC

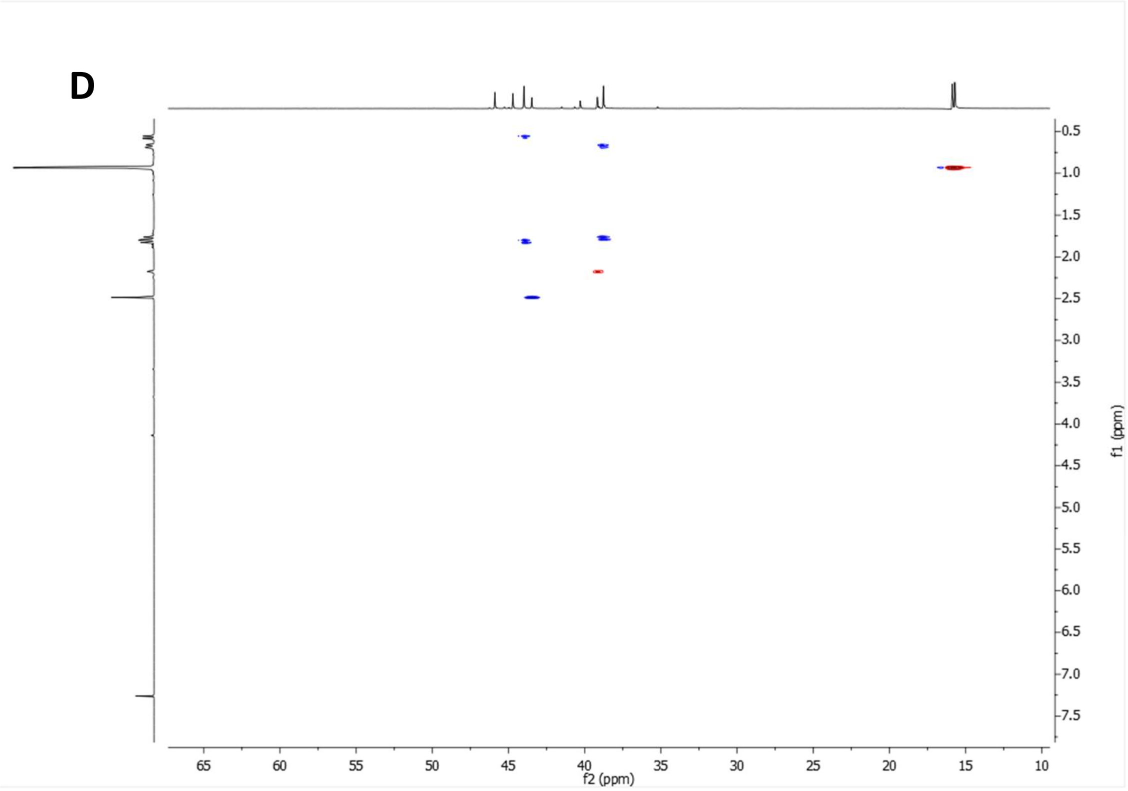


**Supplementary Figure 25:** 2-(3,4,8,9-tetramethyltetracyclo[4.4.0.0$^{3,9}$.0$^{4,8}$]decan-1-yl)acetic acid, ALT-560 (C) $^{13}C$ NMR spectrum. (D) HSQC spectrum.

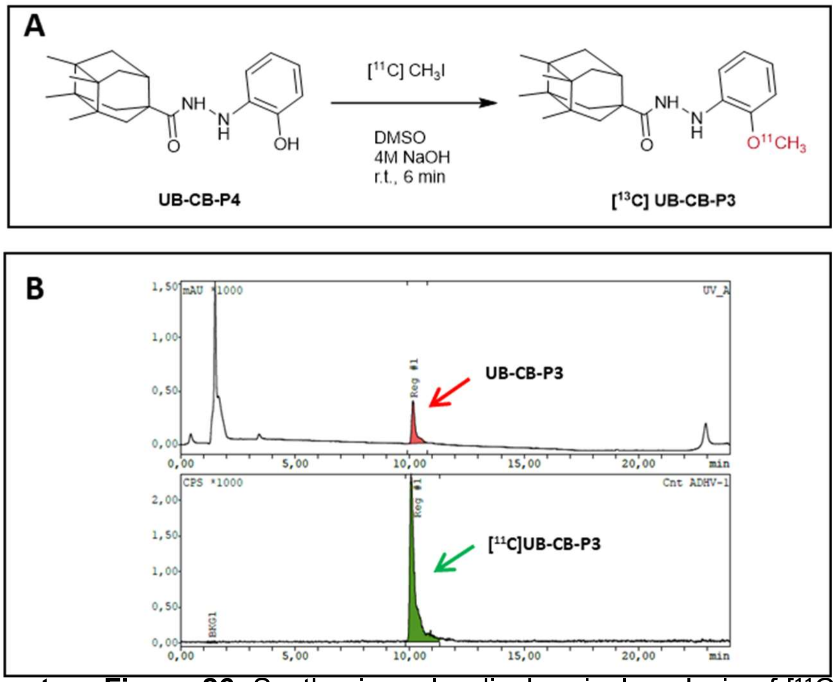


**Supplementary Figure 26.** Synthesis and radiochemical analysis of [11C]UB-CB-P3. **(A)**: Radiosynthesis of [11C]UB-CB-P3 by [11C]$CH_3$I methylation of the desmethyl precursor UB-CB-P4 in DMSO with NaOH at room temperature for 6 min. **(B)** Representative analytical HPLC chromatograms showing the UV trace (upper panel, red peak, UB-CB-P4 reference standard) and the corresponding radiochromatogram (lower panel, green peak, [11C]UB-CB-P3) with co-elution at ~10 min, confirming product identity.

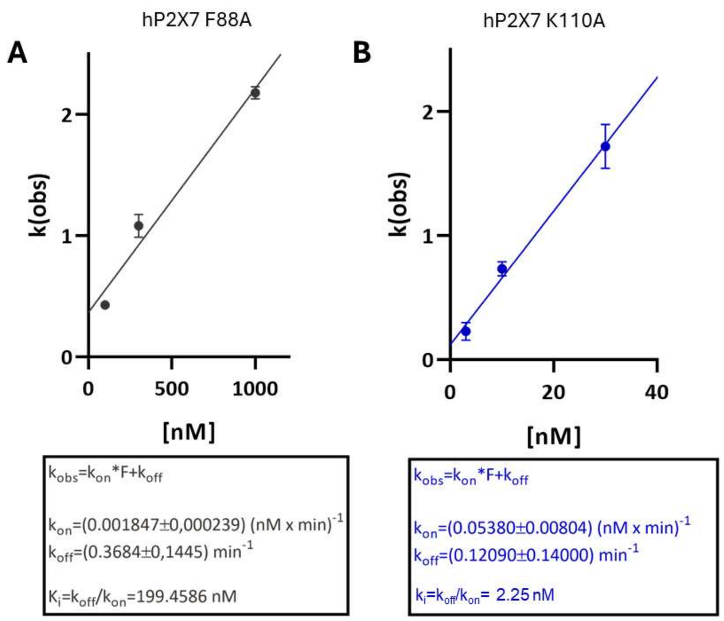


**Supplementary Figure 27. Kinetic analysis of UB-ALT-P2 binding to WT and mutants hP2X7R.** (**A–B**) Dependence of the k(obs) on UB-ALT-P2 concentration for hP2X7R-F88A (**A**) and hP2X7R-K110A. Observed on-rates kobs were determined by fitting the data to the function: % response = (100-Plateau)*exp(-kobs*time)+Plateau. The kobs were then plotted against the respective antagonist concentration (F). Assuming the simplest case of receptor-ligand kinetics, a 1:1 binding model, the formulas kobs=kon*F+koff and Ki=koff/kon were used to obtain an estimate for the theoretical off-rate constant koff (y-intercept) and the corresponding Ki value.

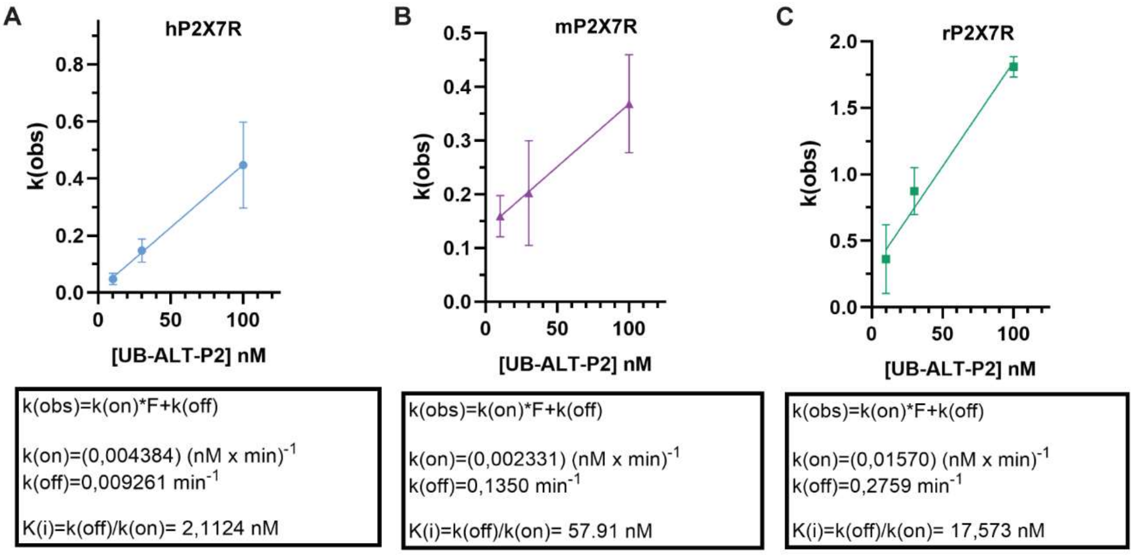


**Supplementary Figure 28. Kinetic analysis of UB-ALT-P2 binding to P2X7R across species.** (**A–C**) Dependence of the k(obs) on UB-ALT-P2 concentration for human (hP2X7R, **A**), mouse (mP2X7R, **B**), and rat (rP2X7R, **C**) receptors. Observed on-rates kobs were determined by fitting the data to the function: % response = (100-Plateau)*exp(-kobs*time)+Plateau. The kobs were then plotted against the respective antagonist concentration (F). Assuming the simplest case of receptor-ligand kinetics, a 1:1 binding model, the formulas kobs=kon*F+koff and Ki=koff/kon were used to obtain an estimate for the theoretical off-rate constant koff (y-intercept) and the corresponding Ki value.

**Supplementary Table 1:** HPLC–UV chromatogram of the new series of compounds.

**Blanc 220 nm**

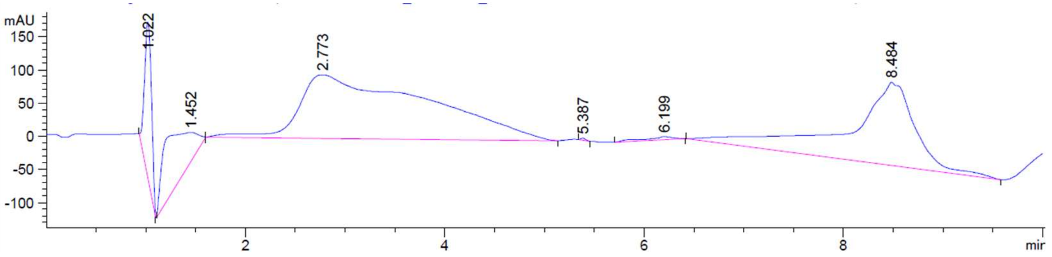


Signal 1: DAD1 A, Sig=220,4 Ref=360,100

| Peak # | RetTime [min] | Type | Width [min] | Area [mAU*s] | Height [mAU] | Area % |
|---|---|---|---|---|---|---|
| 1 | 1.022 | BB | 0.0761 | 1132.14441 | 235.24574 | 6.4975 |
| 2 | 1.452 | BB | 0.4586 | 1594.91003 | 44.03558 | 9.1533 |
| 3 | 2.773 | BB | 1.1466 | 8590.39941 | 95.76094 | 49.3010 |
| 4 | 5.387 | BB | 0.0493 | 9.84798 | 3.08270 | 0.0565 |
| 5 | 6.199 | BB | 0.2424 | 90.74776 | 4.67540 | 0.5208 |
| 6 | 8.484 | BB | 0.5732 | 6006.33984 | 125.30324 | 34.4709 |
| Totals : | | | | 1.74244e4 | 508.10360 | |

**Blanc 254 nm**

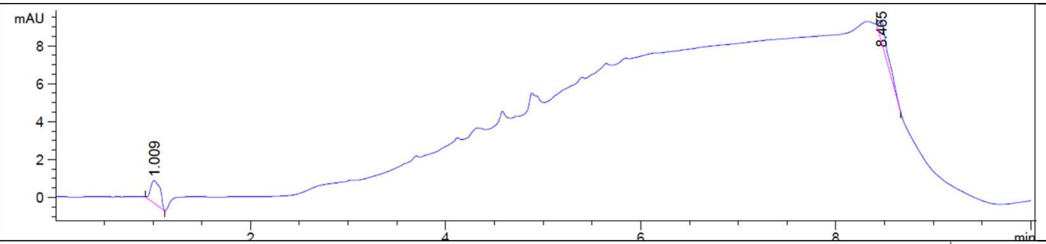


Signal 2: DAD1 B, Sig=254,4 Ref=360,100

| Peak # | RetTime [min] | Type | Width [min] | Area [mAU*s] | Height [mAU] | Area % |
|---|---|---|---|---|---|---|
| 1 | 1.009 | BB | 0.1022 | 8.31069 | 1.17909 | 53.3178 |
| 2 | 8.465 | BB | 0.0943 | 7.27639 | 1.05744 | 46.6822 |
| Totals : | | | | 15.58708 | 2.23653 | |

**CB-P4**

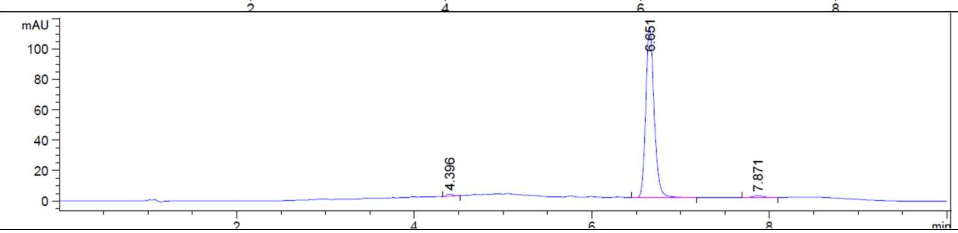


Signal 2: DAD1 B, Sig=254,4 Ref=360,100

| Peak # | RetTime [min] | Type | Width [min] | Area [mAU*s] | Height [mAU] | Area % |
|---|---|---|---|---|---|---|
| 1 | 4.396 | BB | 0.0774 | 6.61909 | 1.34483 | 0.8810 |
| 2 | 6.651 | BB | 0.1001 | 734.28741 | 112.53796 | 97.7373 |
| 3 | 7.871 | BB | 0.1246 | 10.38003 | 1.27997 | 1.3816 |
| Totals : | | | | 751.28654 | 115.16275 | |

**CB-P3**

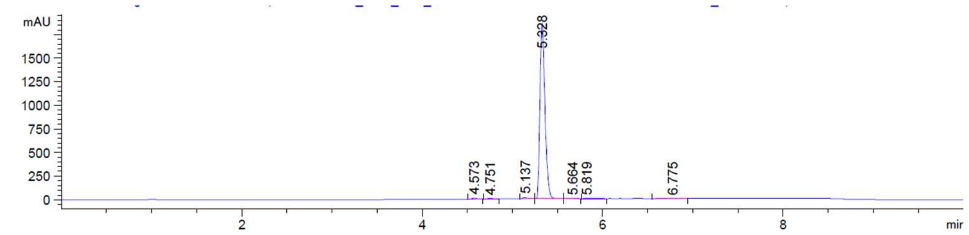


Signal 2: DAD1 B, Sig=254,4 Ref=360,100

| Peak # | RetTime [min] | Type | Width [min] | Area [mAU*s] | Height [mAU] | Area % |
|---|---|---|---|---|---|---|
| 1 | 4.573 | BB | 0.0586 | 38.95277 | 10.19177 | 0.5369 |
| 2 | 4.751 | BB | 0.0547 | 26.42298 | 7.56506 | 0.3642 |
| 3 | 5.137 | BB | 0.0562 | 48.70778 | 12.85499 | 0.6714 |
| 4 | 5.328 | BB | 0.0565 | 7101.17285 | 1860.66724 | 97.8822 |
| 5 | 5.664 | BB | 0.1052 | 12.00163 | 1.91203 | 0.1654 |
| 6 | 5.819 | BB | 0.0896 | 14.34217 | 2.21258 | 0.1977 |
| 7 | 6.775 | BB | 0.0745 | 13.21696 | 2.54866 | 0.1822 |
| Totals : | | | | 7254.81714 | 1897.95233 | |

**ALT-P39**

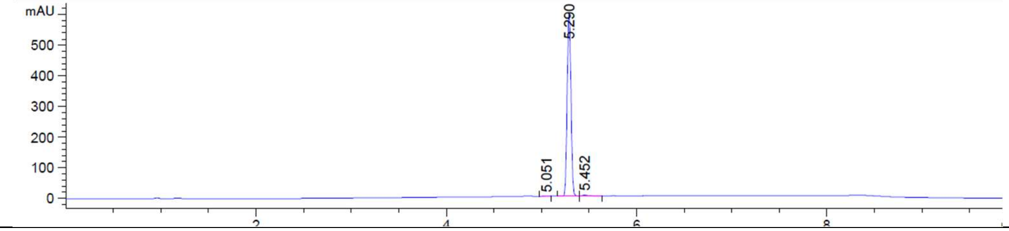


Signal 2: DAD1 B, Sig=254,4 Ref=360,100

| Peak # | RetTime [min] | Type | Width [min] | Area [mAU*s] | Height [mAU] | Area % |
|---|---|---|---|---|---|---|
| 1 | 5.051 | BB | 0.0660 | 5.72237 | 1.51784 | 0.3375 |
| 2 | 5.290 | BB | 0.0447 | 1676.61206 | 598.65588 | 98.8765 |
| 3 | 5.452 | BB | 0.0576 | 13.32887 | 3.40958 | 0.7861 |
| Totals : | | | | 1695.66330 | 603.58331 | |

**ALT-P40**

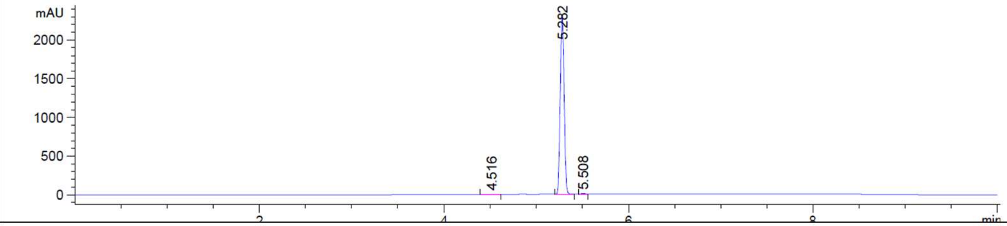


Signal 2: DAD1 B, Sig=254,4 Ref=360,100

| Peak # | RetTime [min] | Type | Width [min] | Area [mAU*s] | Height [mAU] | Area % |
|---|---|---|---|---|---|---|
| 1 | 4.516 | BB | 0.0572 | 5.75199 | 1.48371 | 0.0834 |
| 2 | 5.282 | BB | 0.0464 | 6871.26367 | 2329.24341 | 99.5955 |
| 3 | 5.508 | BB | 0.0433 | 22.15838 | 8.26883 | 0.3212 |
| Totals : | | | | 6899.17404 | 2338.99595 | |

**ALT-P42**

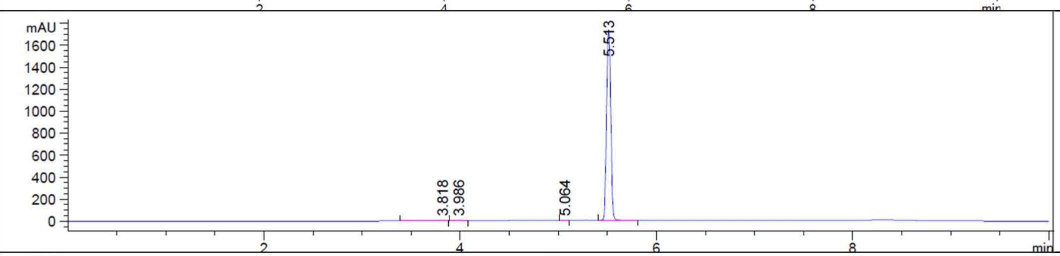


Signal 2: DAD1 B, Sig=254,4 Ref=360,100

| Peak # | RetTime [min] | Type | Width [min] | Area [mAU*s] | Height [mAU] | Area % |
|---|---|---|---|---|---|---|
| 1 | 3.818 | BB | 0.0503 | 15.42529 | 4.69344 | 0.3017 |
| 2 | 3.986 | BB | 0.0553 | 17.66102 | 4.99380 | 0.3454 |
| 3 | 5.064 | BB | 0.0424 | 5.56740 | 2.13684 | 0.1089 |
| 4 | 5.513 | BB | 0.0460 | 5073.99561 | 1739.42664 | 99.2440 |
| Totals : | | | | 5112.64932 | 1751.25071 | |

**CB-P1**

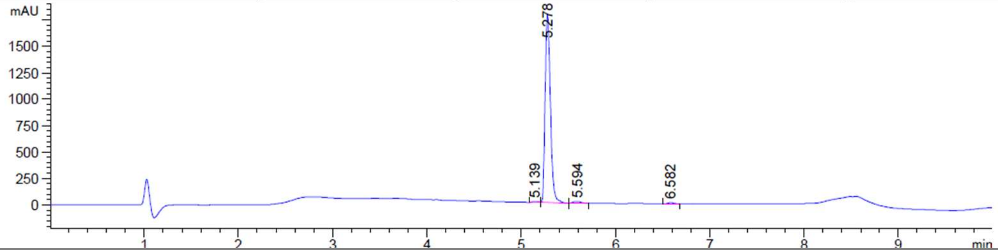


Signal 1: DAD1 A, Sig=220,4 Ref=360,100

| Peak # | RetTime [min] | Type | Width [min] | Area [mAU*s] | Height [mAU] | Area % |
|---|---|---|---|---|---|---|
| 1 | 5.139 | BB | 0.0617 | 26.50827 | 6.22025 | 0.3785 |
| 2 | 5.278 | BB | 0.0584 | 6813.46533 | 1788.79175 | 97.2945 |
| 3 | 5.594 | BB | 0.0973 | 110.71208 | 18.10458 | 1.5809 |
| 4 | 6.582 | BB | 0.0634 | 52.24287 | 12.84591 | 0.7460 |
| Totals : | | | | 7002.92855 | 1825.96249 | |

**ALT-P35**

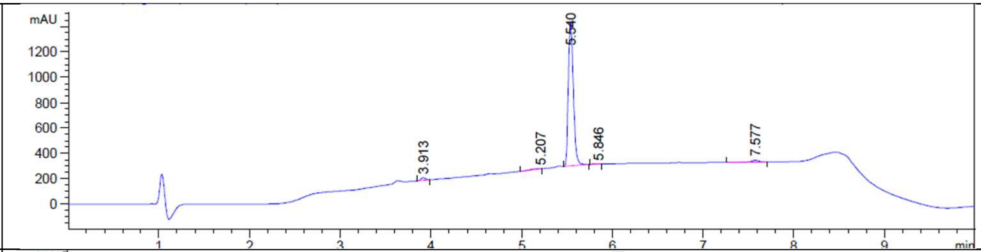


Signal 1: DAD1 A, Sig=220,4 Ref=360,100

| Peak # | RetTime [min] | Type | Width [min] | Area [mAU*s] | Height [mAU] | Area % |
|---|---|---|---|---|---|---|
| 1 | 3.913 | BB | 0.0565 | 76.42796 | 20.97732 | 1.6842 |
| 2 | 5.207 | BB | 0.2158 | 30.47289 | 1.74486 | 0.6715 |
| 3 | 5.540 | BB | 0.0584 | 4347.38086 | 1143.03516 | 95.8018 |
| 4 | 5.846 | BB | 0.0981 | 8.19845 | 1.11039 | 0.1807 |
| 5 | 7.577 | BB | 0.0815 | 75.41255 | 14.30258 | 1.6618 |

Totals : 4537.89271 1181.17030

**ALT-P1**

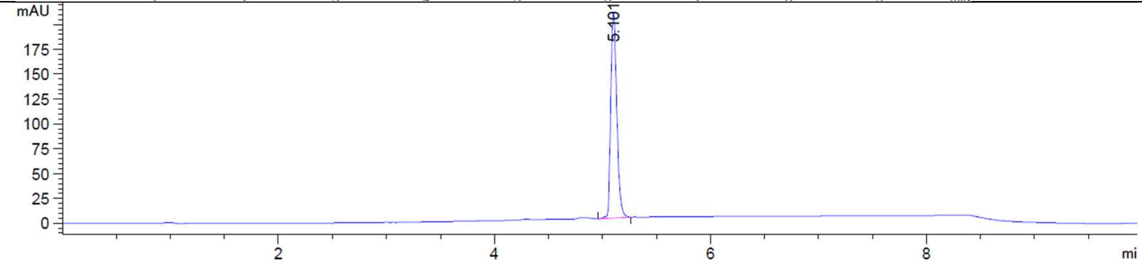


Signal 2: DAD1 B, Sig=254,4 Ref=360,100

| Peak # | RetTime [min] | Type | Width [min] | Area [mAU*s] | Height [mAU] | Area % |
|---|---|---|---|---|---|---|
| 1 | 5.101 | BB | 0.0583 | 790.72723 | 208.28522 | 100.0000 |

Totals : 790.72723 208.28522

**ALT-P2**

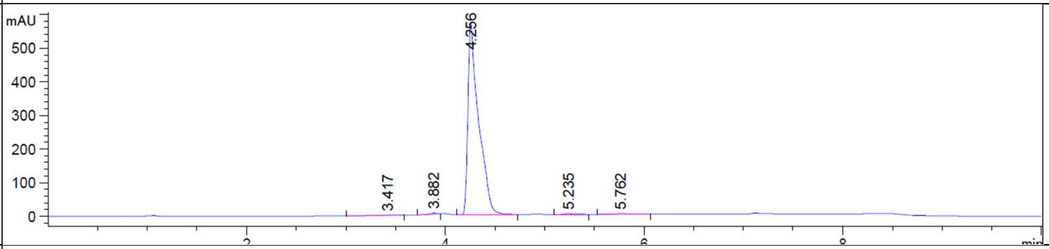


Signal 2: DAD1 B, Sig=254,4 Ref=360,100

| Peak # | RetTime [min] | Type | Width [min] | Area [mAU*s] | Height [mAU] | Area % |
|---|---|---|---|---|---|---|
| 1 | 3.417 | BB | 0.1440 | 13.54820 | 1.23497 | 0.3129 |
| 2 | 3.882 | BB | 0.1007 | 16.49966 | 2.57604 | 0.3811 |
| 3 | 4.256 | BB | 0.1008 | 4268.82031 | 573.45184 | 98.5881 |
| 4 | 5.235 | BB | 0.1112 | 12.80617 | 1.75411 | 0.2958 |
| 5 | 5.762 | BB | 0.1425 | 18.28262 | 1.71387 | 0.4222 |

**ALT-P21**

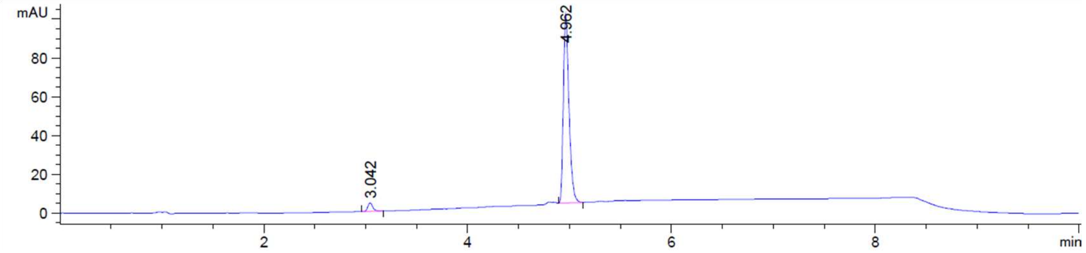


Signal 2: DAD1 B, Sig=254,4 Ref=360,100

| Peak # | RetTime [min] | Type | Width [min] | Area [mAU*s] | Height [mAU] | Area % |
|---|---|---|---|---|---|---|
| 1 | 3.042 | BB | 0.0517 | 14.85983 | 4.36331 | 3.8092 |
| 2 | 4.962 | BB | 0.0589 | 375.24487 | 97.50396 | 96.1908 |

Totals : 390.10470 101.86727

| | | |
|---|---|---|
| **ALT-P10** | 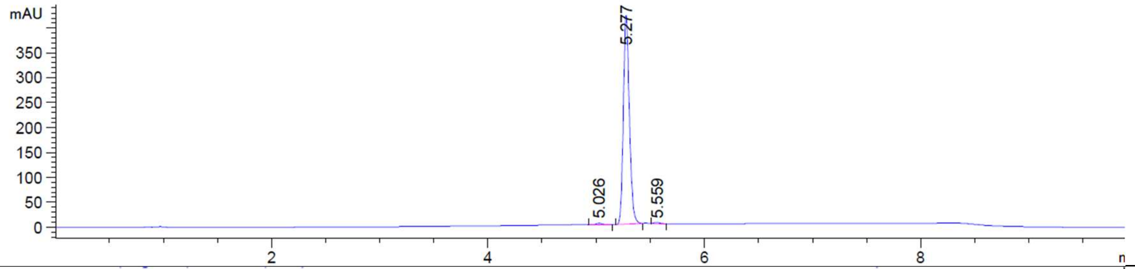 | Signal 2: DAD1 B, Sig=254,4 Ref=360,100 |
| **ALT-P25** | 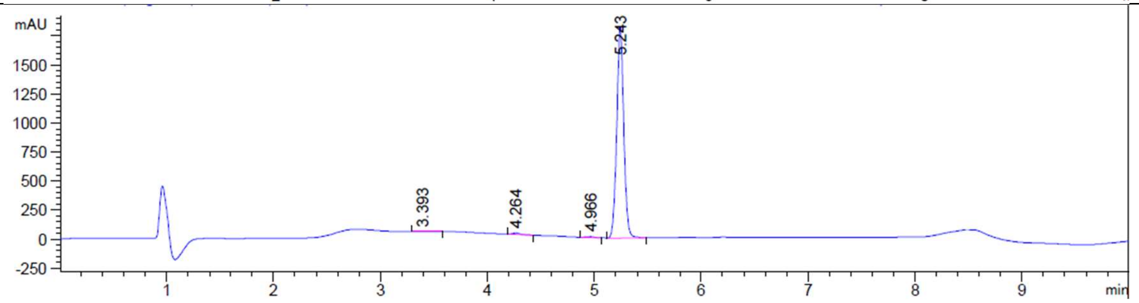 | Signal 1: DAD1 A, Sig=220,4 Ref=360,100 |
| **ALT-P22** | 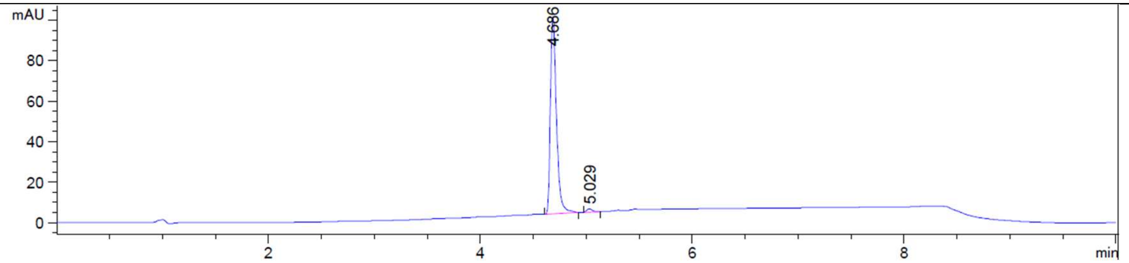 | Signal 2: DAD1 B, Sig=254,4 Ref=360,100 |

ALT-P10 — Signal 2: DAD1 B, Sig=254,4 Ref=360,100

| Peak # | RetTime [min] | Type | Width [min] | Area [mAU*s] | Height [mAU] | Area % |
|---|---|---|---|---|---|---|
| 1 | 5.026 | BB | 0.0571 | 10.69952 | 2.76950 | 0.6504 |
| 2 | 5.277 | BB | 0.0590 | 1624.28662 | 421.46146 | 98.7322 |
| 3 | 5.559 | BB | 0.0554 | 10.15696 | 2.86523 | 0.6174 |
| Totals : | | | | 1645.14310 | 427.09619 | |

ALT-P25 — Signal 1: DAD1 A, Sig=220,4 Ref=360,100

| Peak # | RetTime [min] | Type | Width [min] | Area [mAU*s] | Height [mAU] | Area % |
|---|---|---|---|---|---|---|
| 1 | 3.393 | BB | 0.1229 | 9.34516 | 1.00137 | 0.1106 |
| 2 | 4.264 | BB | 0.0762 | 68.36827 | 13.25361 | 0.8093 |
| 3 | 4.966 | BB | 0.0743 | 42.68462 | 8.53469 | 0.5052 |
| 4 | 5.243 | BB | 0.0692 | 8327.87012 | 1826.80884 | 98.5749 |
| Totals : | | | | 8448.26817 | 1849.59851 | |

ALT-P22 — Signal 2: DAD1 B, Sig=254,4 Ref=360,100

| Peak # | RetTime [min] | Type | Width [min] | Area [mAU*s] | Height [mAU] | Area % |
|---|---|---|---|---|---|---|
| 1 | 4.686 | BB | 0.0597 | 382.74762 | 97.67065 | 98.5320 |
| 2 | 5.029 | BB | 0.0552 | 5.70262 | 1.61463 | 1.4680 |
| Totals : | | | | 388.45024 | 99.28528 | |

**ALT-P16**

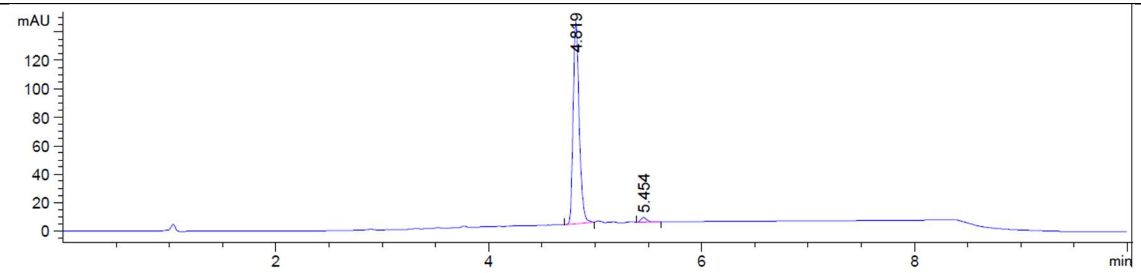

Signal 2: DAD1 B, Sig=254,4 Ref=360,100

| Peak # | RetTime [min] | Type | Width [min] | Area [mAU*s] | Height [mAU] | Area % |
|---|---|---|---|---|---|---|
| 1 | 4.819 | BB | 0.0599 | 556.24750 | 141.46063 | 97.7019 |
| 2 | 5.454 | BB | 0.0602 | 13.08360 | 3.30039 | 2.2981 |
| Totals : | | | | 569.33110 | 144.76103 | |

**ALT-P13**

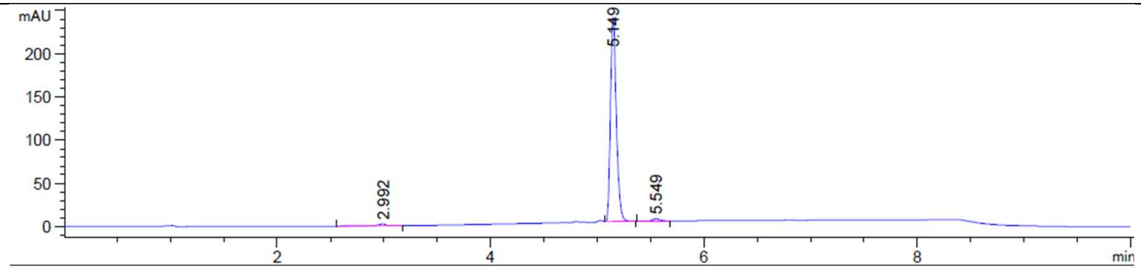

Signal 2: DAD1 B, Sig=254,4 Ref=360,100

| Peak # | RetTime [min] | Type | Width [min] | Area [mAU*s] | Height [mAU] | Area % |
|---|---|---|---|---|---|---|
| 1 | 2.992 | BB | 0.0816 | 12.67725 | 2.12385 | 1.3639 |
| 2 | 5.149 | BB | 0.0583 | 903.05365 | 238.02818 | 97.1570 |
| 3 | 5.549 | BB | 0.0750 | 13.74782 | 2.71693 | 1.4791 |
| Totals : | | | | 929.47872 | 242.86897 | |

**Supplementary Table 2:** Cryo-EM collection, refinement, and validation statistics.

| | UB-MBX-47 hP2X7R (EMD-74135) (PDB: 9ZF7) | UB-ALT-P1 hP2X7R (EMD-74139) (PDB: 9ZFB) | UB-ALT-P2 hP2X7R (EMDB-74136) (PDB: 9ZF8) | UB-ALT-P2 mP2X7R (EMD-74138) (PDB: 9ZFA) | UB-ALT-P2 rP2X7R (EMD-74134) (PDB: 9ZF6) |
|---|---|---|---|---|---|
| **Data collection and processing** | | | | | |
| Magnification (kx) | 130 | 130 | 130 | 130 | 130 |
| Voltage (kV) | 300 | 300 | 300 | 300 | 300 |
| Electron exposure (e–/Å$^2$) | 46 | 43 | 42 | 45 | 43 |
| Movie frames | 50 | 50 | 43 | 50 | 48 |
| Defocus range (μm) | -0.9 to -1.4 | -1.0 to -1.5 | -0.7 to -1.4 | -0.9 to -1.5 | -0.9 to -1.4 |
| Pixel size (Å) | 0.6485 | 0.647 (0.3235 super-res) | 0.6483 (0.3242 super-res) | 0.647 (0.3235 super-res) | 0.6483 (0.3243 super-res) |
| Symmetry imposed | C3 | C3 | C3 | C3 | C3 |
| Initial micrographs (no.) | 23,876 | 17,388 | 21,863 | 17,244 | 11,520 |
| Final micrographs used (no.) | 22,817 | 14,677 | 19,873 | 15,307 | 10,747 |
| Initial particle images (no.) | 2,691,280 | 1,570,109 | 3,297,698 | 1,728,998 | 4,040,409 |
| Final particle images (no.) | 89,484 | 178,063 | 364,964 | 422,475 | 287,449 |
| Map resolution (Å) FSC threshold | 2.81 (0.143) | 2.94 (0.143) | 2.56 (0.143) | 2.26 (0.143) | 2.58 (0.143) |
| Map resolution range (Å) | 1.6 to 28 | 2.5 to 27 | 1.4 to 24 | 1.4 to 9.4 | 1.4 to 27 |
| **Refinement** | | | | | |
| Initial model used (PDB code) | 9E3P | 9E3P | 9E3P | 9E3Q | 8TR8 |
| Model resolution (Å) FSC threshold | 2.78 (0.143) | 2.90 (0.143) | 2.52 (0.143) | 2.24 (0.143) | 2.56 (0.143) |
| Map sharpening *B* factor (Å$^2$) | 94.8 | 88.3 | 92.0 | 69.1 | 88.2 |
| **Model composition** | | | | | |
| Non-hydrogen atoms | 14,085 | 14013 | 14,152 | 13,876 | 13,639 |
| Protein Residues | 1626 | 1632 | 1626 | 1620 | 1,623 |
| Ligands | 30 | 30 | 30 | 19 | 24 |
| Waters | 159 | 132 | 250 | 291 | 178 |
| *B* factors (Å$^2$) | | | | | |
| Protein | 24.6/126/55.6 | 12.0/74.0/39.0 | 7.52/127/59.7 | 20.7/73.1/35.1 | 42.1/81.4/54.1 |
| Ligand | 38.0/108/74.8 | 16.2/75.2/34.9 | 13.7/122/46.4 | 27.4/54.6/32.9 | 48.7/71.8/55.8 |
| Nucleotide | 11.5/11.5/11.5 | 56.5/56.5/56.5 | 104/104/104 | 31.7/31.7/31.7 | 55.8/55.8/55.8 |
| Water | 30.0/60.3/45.8 | 11.0/55.8/19.8 | 11.6/90.2/24.0 | 24.9/46.7/31.6 | 42.0/58.6/50.3 |
| R.m.s. deviations | | | | | |
| Bond lengths (Å) | 0.009 (0) | 0.009 (0) | 0.005 (0) | 0.007 (0) | 0.012 (0) |
| Bond angles (°) | 1.136 (0) | 0.891 (0) | 0.816 (0) | 0.759 (0) | 0.946 (0) |
| Validation | | | | | |
| MolProbity score | 1.46 | 1.30 | 0.99 | 1.22 | 1.43 |
| Clash score | 4.42 | 4.25 | 2.18 | 4.37 | 4.55 |
| Poor rotamers (%) | 0.00 | 0 | 0.00 | 0.00 | 0.00 |
| Ramachandran plot | | | | | |
| Favored (%) | 96.36 | 97.52 | 98.21 | 98.34 | 96.76 |
| Allowed (%) | 3.64 | 2.48 | 1.79 | 1.66 | 3.24 |
| Disallowed (%) | 0 | 0 | 0 | 0 | 0 |

**Supplementary Table 3.** Cellular cytotoxicity in HEL, HeLa, Vero and MT4 cell lines.

| | Cellular Cytotoxicity | | | |
|---|---|---|---|---|
| | **HEL cells $CC_{50}$ (µM)** | **HeLa cells $CC_{50}$ (µM)** | **Vero cells $CC_{50}$ (µM)** | **MT4 cells $CC_{50}$ (µM)** |
| **UB-MBX-46** | ≥100 | ≥100 | ≥100 | >50 |
| **UB-MBX-47** | ≥100 | ≥100 | ≥100 | >50 |
| **UB-ALT-P1** | ≥100 | ≥100 | ≥100 | >50 |
| **UB-ALT-P2** | ≥100 | ≥100 | ≥100 | >50 |

$CC_{50}$ values were determined after 72 h of compound exposure using a standard cell viability assay. All compounds displayed $CC_{50}$ ≥ 50 µM in all tested lines, indicating low intrinsic cytotoxicity across both epithelial- and lymphoid-derived cell lines.

**Supplementary Table 4**. Permeability (*Pe* $10^{-6}$ cm $s^{-1}$) in the PAMPA-BBB assay from 14 commercial drugs and the assayed compounds (three different experiments in triplicate) and predictive penetration in the CNS.

| Compound | Bibliography value[38] | Experimental value (n = 3) ± S.D. | ***CNS Prediction*** |
|---|---|---|---|
| Verapamil | 16.0 | 26.4 ± 0.7 | N/A |
| Testosterone | 17.0 | 25.8 ± 0.5 | N/A |
| Corticosterone | 5.1 | 6.7 ± 0.1 | N/A |
| Clonidine | 5.3 | 6.5 ± 0.05 | N/A |
| Ofloxacin | 0.8 | 0.1 ± 0.07 | N/A |
| Lomefloxacin | 0.0 | 0.8 ± 0.03 | N/A |
| Progesterone | 9.3 | 16.8 ± 0.3 | N/A |
| Promazine | 8.8 | 13.8 ± 0.3 | N/A |
| Imipramine | 13.0 | 12.5 ± 0.2 | N/A |
| Hydrocortisone | 1.9 | 1.4 ± 0.05 | N/A |
| Piroxicam | 2.5 | 2.0 ± 0.08 | N/A |
| Desipramine | 12.0 | 17.8 ± 0.1 | N/A |
| Cimetidine | 0.0 | 0.7 ± 0.03 | N/A |
| Norfloxacin | 0.1 | 0.8 ± 0.05 | N/A |
| UB-MBX-46 | N/A | 7.5 ± 0.1 | CNS+ |
| UB-MBX-47 | N/A | 8.1 ± 0.3 | CNS+ |
| UB-ALT-P1 | N/A | 14.5 ± 1.45 | CNS+ |
| UB-ALT-P2 | N/A | 16.5 ± 0,85 | CNS+ |

**Supplementary Table 5:** Species-specific amino acid differences at key positions of the P2X7 receptor classical allosteric pocket.

| **Residue position** | **95 (± 5 aa)** | **108 (± 5 aa)** | **312 (± 5 aa)** |
|---|---|---|---|
| **Human P2X7R** | TADYT**F**PLQGN | QGNSF**F**VMTNF | RTLIK**V**FGIRF |
| **Mouse P2X7R** | TADYT**F**PLQGN | QGNSF**Y**VMTNF | RTLIK**A**FGIRF |
| **Rat P2X7R** | TADYT**L**PLQGN | QGNSF**Y**VMTNY | RTLIK**A**FGVRF |

Short sequence fragments (±5 residues) surrounding positions 95, 108, and 312 are shown for human, mouse and rat P2X7 receptors. Residue numbering is based on the human P2X7R sequence. The highlighted residues indicate species-specific substitutions within the classical allosteric pocket that modulate ligand binding.

**Supplementary Table 6:** Primary and secondary antibodies used for protein level determination by Western blotting in the *in vivo* UB-ALT-P2 study in 5xFAD murine model.

| Antibody | Host | Source/Catalog | WB dilution |
|---|---|---|---|
| **GAPDH** | Mouse | Millipore/MAB374 | 1:5000 |
| **Tau Total** | Mouse | Invitrogen/AHB0042 | 1:1000 |
| **p-Tau (Ser404)** | Rabbit | Invitrogen/44758G | 1:1000 |
| **AT8** | Rabbit | Pierce Rockford | 1:1000 |
| **SOD1** | Sheep | Calbiochem/574597 | 1:1000 |
| **Goat-anti-rabbit HRP conjugated** | | Biorad/170-6515 | 1:5000 |
| **Goat-anti-mouse HRP conjugated** | | Biorad/170-5047 | 1:5000 |

**Supplementary Table 7:** Primers and probes used in qPCR studies for the in vivo UB-ALT-P2 assay.

| Target | Product size (bp) | Forward primer (5'-3') | Reverse primer (5'-3') |
|---|---|---|---|
| ***β-actin*** | **190** | CAACGAGCGGTTCCGAT | GCCACAGGTTCCATACCCA |
| ***Il-6*** | **189** | ATCCAGTTGCCTTCTTGGGACTGA | TAAGCCTCCGACTTGTGAAGTGGT |
| ***Il-1β*** | **179** | ACAGAATATCAACCAACAAGTGATATTCTC | GATTCTTTCCTTTGAGGCCCA |
